# Tracing the evolution of aneuploid cancers by multiregional sequencing with CRUST

**DOI:** 10.1101/2020.11.11.376467

**Authors:** Subhayan Chattopadhyay, Jenny Karlsson, Anders Valind, Natalie Andersson, David Gisselsson

**Affiliations:** Division of Clinical Genetics, Department of Laboratory Medicine, Lund University, Lund, Sweden; Department of Pediatrics, Skåne University Hospital, Lund, Sweden; Division of Oncology and Pathology, Department of Clinical Sciences, Lund University, Lund, Sweden; Clinical Genetics and Pathology, Laboratory Medicine, Lund University Hospital, Skåne Healthcare Region, Lund, Sweden

**Keywords:** clonal deconvolution, subclonal reconstruction, multiregional sequencing, phylogeny

## Abstract

Clonal deconvolution of mutational landscapes is crucial to understand the evolutionary dynamics of cancer. Two limiting factors for clonal deconvolution that have remained unresolved are variation in tumor cell fraction (TCF) and chromosomal copy number across different samples of the same tumor. We developed a semi-supervised algorithm that tracks variant calls through multi-sample spatiotemporal tumor data. While normalizing allele frequencies based on TCF it also adjusts for copy number changes at clonal deconvolution. Absent à priori copy number data, it renders *in silico* copy number estimations from bulk sequences. Using published and simulated tumor sequences, we reliably segregated clonal/subclonal variants even at low sequencing depth (∼50x). Given at least one pure tumor sample (>70% TCF), we could normalize and deconvolve paired samples down to a TCF of 40%. This renders a reliable clonal reconstruction well adapted to multi-regionally sampled solid tumor, which are often aneuploid and contaminated by non-cancer cells.

## Introduction

Genetic diversification during tumorigenesis and disease progression is governed by Darwinian principles acting on the level of single cells. A concerted effort has been dispensed in recent years to unravel the mechanisms of evolutionary dynamics in cancer. Next generation sequencing across cancer types has confirmed that intratumor heterogeneity through phylogenetic branching is a common scenario[1], although the relative contributions from clonal selection versus neutral evolution in this process remain a matter of debate[2, 3].We recently demonstrated that intratumor genetic heterogeneity can result as a product of different evolutionary trajectories specific to the spatiotemporal localization of cells residing in a tumor[4]. Although all such cells are popularly believed to be neutrally evolved progenies of a common ancestor, depending on oncogenicity of the mutations acquired, some daughter cells observe a greater fitness advantage than those with neutral mutations. This pattern of divergent evolution can be observed by interrogating bulk sequencing data from tumors. As the genetic landscape in solid tumors often varies geographically within the same cancer, comprehensive reconstruction of tumor phylogeny requires multi-regional analysis and subclonal deconvolution[5, 6]. Such a deconvolution process leverages the relative abundance of a mutation across samples represented by variant allele frequencies (VAFs)[7]. Besides being very loosely defined, designation of clonality of a mutant variant depends on when in tumor development it emerges, its effect on cellular fitness, and on the spatial architecture of the tumor. A subclonal mutation is suggestively defined as any lesion that emerges out of a clone and does not observe extensive positive selection[8]. However, subclonal populations do accumulate mutations in known ‘driver’ genes, sometimes even emerging as events of convergent evolution[9, 10]. A subclonal lesion thus may represent itself in a fraction of the tumor cells, and its relative abundance often vary among samples even if these are acquired from a tumor at the same time. Hence subclonal mutations may become regionally fixed (present in all tumor cells) and thus appear clonal. Systematic bioinformatic approaches are critical to resolve such complex scenarios.

Clonal deconvolution is generally attempted using unsupervised clustering. It determines subclonal populations with distributional assumption on the variant read count[7]. Most methods assume that in a repertoire of clones and subclones, the relative abundances of variants resemble a binomial and beta-binomial admixture[11–13]. Thus, a Dirichlet finite mixture can segregate several clonal populations with distinct shapes and scales. Following this dogma, one can expect the clonal mutations to be distributed with markedly higher mean relative frequency than subclonal progenies. However, even with a modest rate of silent substitutions, many passenger mutations accumulate in subclonal populations within a few generations presenting a gradually regressing heavy tail of private mutations[13]. Furthermore, samples collected from a tumor at different locations or stages of progression can contain a remarkably varied proportion of normal cells from adjacent non-cancerous tissue. As a result, the VAF distribution of clonal mutations of one sample can mimic that of subclonal mutations of another, purer sample. Variants can also end up with higher (or lower) than expected relative abundance if residing in chromosomal regions affected by copy number changes. As copy number alterations can appear as both clonal and subclonal lesions, they can significantly complicate clonal deconvolution based on VAF distributions[14].

Here, we intended to solve the problem of how VAF values are influenced by purity and copy number with an analysis suite for clonal deconvolution named CRUST (**C**lonal **R**econstruction of t**U**mors with **S**patio**-T**emporal sampling). It can classify clonal and subclonal mutations from bulk sequencing data of multi-sampled tumor tissue. CRUST will probabilistically rescale VAFs from samples with low tumor cell fractions (TCFs) contrasting against the sample with the highest purity. The program is additionally built to integrate data from precise copy number estimations such as those from single nucleotide polymorphism (SNP array) data and produce allelic composition specific clonality predictions. In absence of SNP array data, CRUST can estimate copy number profiles given sequencing summaries from tumor and constitutional genome. While these processes need to have mathematical rigor by parameterizing stochastic assumption on the distribution of clonal variants, we also recognized the need of a semi-supervised algorithm to reconcile a purely traditional data driven approach and a user driven heuristic pattern recognition for clonal deconvolution. The user thus will be able to actively curate the semi-supervised clustering process prior to the deconvolution based on visual input. CRUST also allows sample specific user driven readjustments in deconvolution post analysis. We were able to demonstrate in clinical samples and in simulated tumor biopsies that in presence of at least one relatively pure sample, without user intervention, CRUST can rectify clonality estimates for samples with compromising purity that would otherwise be heavily biased towards a prediction of subclonality. Furthermore, we demonstrated that CRUST increases the resolution of clonal deconvolution for aneuploid tumors by taking the influence of chromosomal copy numbers into account. With curation provided by the user and proper consideration to the temporal fluctuations in appearance of each genetic lesion, CRUST can thus help reconstruct the most likely phylogenetic history of a tumor.

## Results

### A semi-supervised approach to clonal deconvolution

The primary functionality of CRUST is in clonal deconvolution from a substrate of sequencing summaries of single nucleotide variants. As a first example, we demonstrate this on a hypothetical tetraploid tumor where mutations are present in either one or three out of the four available homologous chromosomes (**Figure 1A**). The simulated tumor is represented by eight biopsies (inbuilt data *test.dat*). With a set of samples from the same tumor obtained at different locations and/or time points, CRUST deconvolves each variant to a predicted clonal or sub-clonal status, calibrating clonality assignment against given parameters on allele-specific copy number status and sample purity **(Figure 1B-C; Methods: Quick user guide)**. It realigns the frequency distribution across samples with probabilistic quotient normalization. Hereafter the distribution is queried to fit into an optimum number of clusters based on statistics comparing loss of information (**Supplementary Figure 1**). With the copy number analysis, sequence variants from a single tumor are analyzed separately for each allelic configuration (1+1, 1+2, 2+0 etc.), where CRUST visualizes the predicted clonal/subclonal assignments for all spatiotemporal samples. The subclonal estimation process is based on semi-supervised cluster determination. It verifies the optimal solution first without user input; next the user is given opportunity to override the unsupervised solution after visual inspection of the expected subclonality (**Figure 1D I-II)** to retain provision for a biologically derived deconvolution assessment, if needed. In addition, subclonality assignment can be altered for specific samples post-prediction, a feature useful in presence of compromising purity or inter-sample heterogeneity with respect to the complexity of chromosomal alterations (e.g., chromothripsis and whole genome doubling). In this hypothetical case, CRUST thus correctly assigns mutations present in 1/4 and 3/4 alleles to both clonal and subclonal states, while a cluster-based deconvolution without accounting for copy number may assign all mutations present in 1/4 alleles to the subclonal stratum.

**Figure 1.**
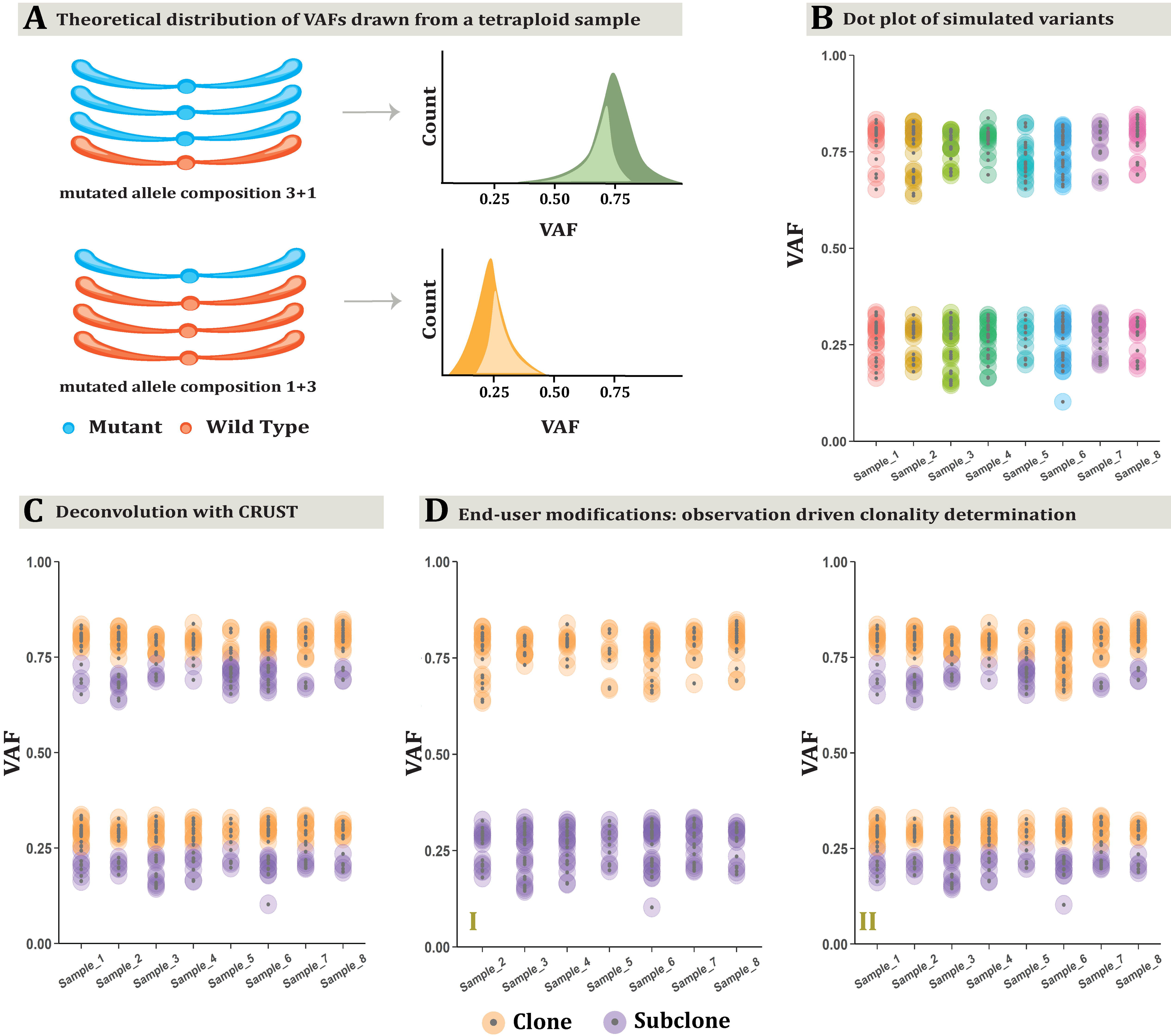
Clonal deconvolution of a simulated tumor genome. A tetraploid tumor is simulated where all samples adhere to an allelic composition of 3+1 or 1+3 (**A**). This makes the expected VAF distribution bimodal with corresponding peaks at frequencies 0.25 (i.e., 1/4) and 0.75 (i.e., 3/4). There are eight samples representing different biopsies. CRUST first displays a dot plot of the VAFs pertaining to all samples (**B**). Given provision for a purely estimation driven approach, it predicts clonality from the optimum number of clusters determined without supervision. This results in a deconvolution independent of the user suggested input (**C**). A user can decide to opt for a semi-supervised approach instead if the optimum number of clusters predicted is dissimilar to a biologically expected deconvolution, for example prior knowledge from single cell karyotyping or sequencing. In this example the default optimization is given with 4 clusters as seen above (two clonal and two subclonal). In (**D I)** however, the user chooses to fit a 2-cluster deconvolution resulting in a prediction of one clonal and one subclonal cluster. The predictions can also be modulated post-hoc for individual samples **(D II)**. Over the default optimum prediction, for *Sample_6* a user has here chosen to fit a 3-cluster deconvolution that picks up two clusters attributed to the clonal population (at allelic compositions 1+3 and 3+1) and one subclonal.

### Performance testing of scaling with simulation

To assess the accuracy of CRUST-based deconvolution across varied purity and sequencing coverage, we simulated tumor samples under three distinct assumptions (**Figure 2A; Supplementary Figure 2**). Here, the frequency distribution of variants queried from low depth calls were left-tail heavy although the pure distribution is expected to follow a beta-binomial distribution. Extending from a one-parametric power law function[13], we modelled the reduction in variability biased towards the left tail with a log-exponent function and simulated an admixture of clonal and subclonal variants assuming a balanced background copy number. The model assumptions indicated what the varying tumor cell fractions (TCF) could affect; first, the mean i.e., the cluster centroid of the VAF distribution; second, variance i.e., the scale of the distribution and lastly, both mean and variance. Accuracy of prediction was measured by comparing expected clonality based on the simulation assumption and the predicted clonality inferred by CRUST with the Jaccard index. For all three model assumptions, 1500 simulations were performed respectively to generate two samples in each iteration. TCFs were sequentially modified for each iteration to have produced 100 variants for each sample. When only the variance of the distributions were varied, the deconvolution and prediction accuracy broke down fastest and the departure was statistically significant (**Figure 2B**) suggesting subclonal clusters can retain the same centroids while increasing in variance but the VAFs overlap between clusters making it virtually impossible to segregate them. CRUST scaling maintained a concordance of 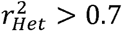 in presence of at least one representative sample (TCF > 0.7) given that the difference in TCF between samples was less than 30% of the maximum; with a larger than 30% departure, the concordance decreased drastically.

**Figure 2.**
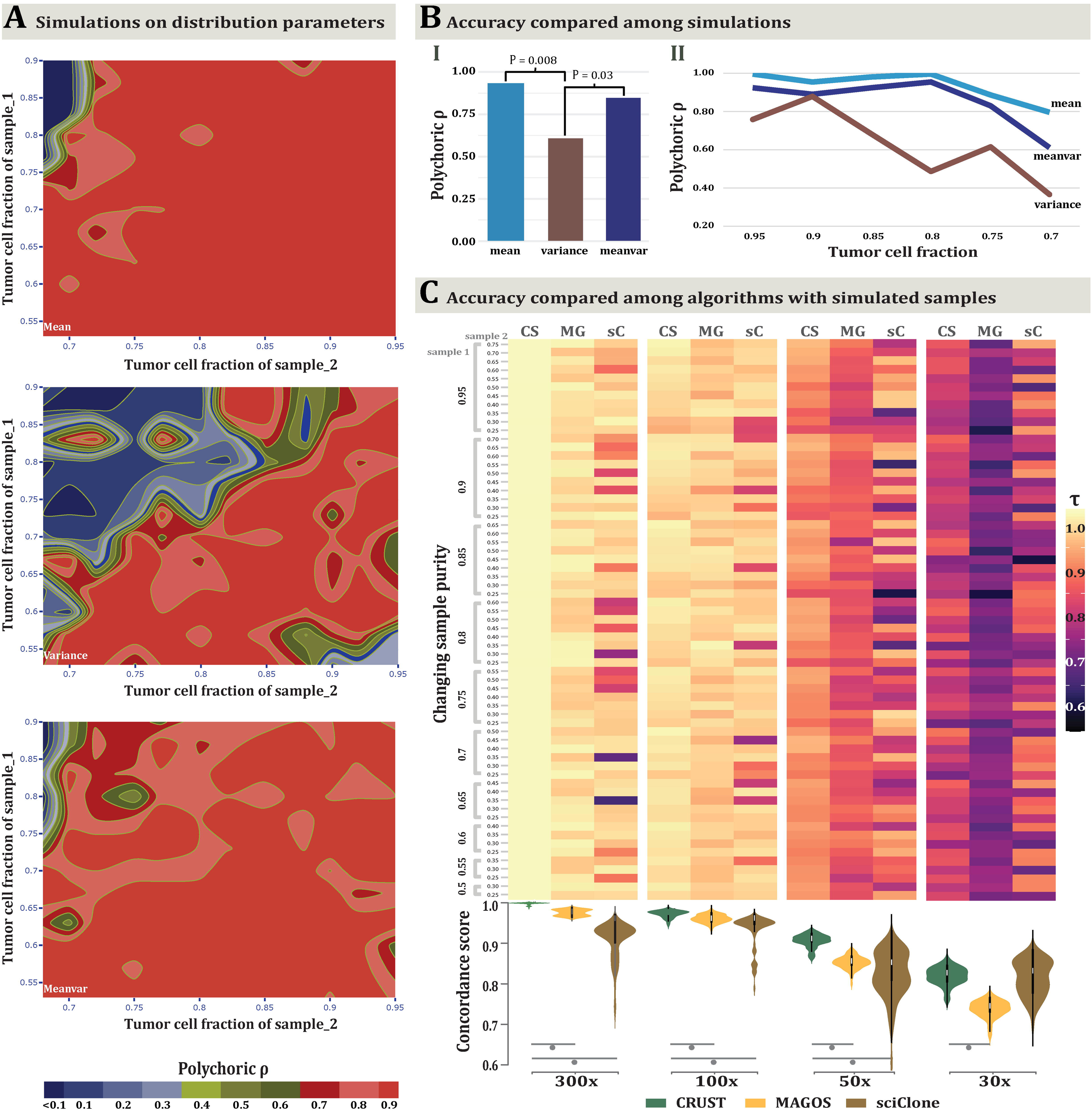
Evaluation of efficacy of CRUST with simulation. In (**A**) simulations of scaling with varying sample composition are shown. Each iteration generates two samples, say X and Y with tumor cell fractions (TCF) Tx and Ty, respectively. Assuming Tx > Ty, CRUST rescales the variants in Y based on those in X. Simulations are performed to see how well the scaling works when Tx and Ty are varied. Three parametric beta-log normal models are in effect to generate simulated samples. The top panel shows changes in TCF that only affects the mean of the VAF distribution. The middle shows changes in TCF affecting the variance (ergo spread) of the VAF distribution and the lower most panel shows when it dynamically affects both mean and variance (referred as *Meanvar*). The measured statistic is polychoric correlation among predictions and its scale for all three simulations is the same, as is indicated at the bottom. In (**B-I**), average marginal concordance is estimated with geometric mean for all three methods and tests are performed between pairs. Only significant deviations are marked with corresponding P values. In (**B-II**) the trend of change in average concordance with varying levels of TCF between the three algorithms is depicted. A comparison across deconvolution methods was done with simulation of varying sequencing coverage **(F)**. Samples are drawn with varying TCF for four sets of coverage at 300x, 100x, 50x and 30x. Ordinal cluster similarities were assessed between CRUST (CS), MAGOS (MG) and sciClone (sC) with Jaccard coefficient (τ). The four combined heatmap and violin plots correspond to four coverage settings denoted in the x axis. Each combination represents summary statistics obtained as median τ for paired TCFs. Each cell in the heatmap reflects that obtained from a paired simulated sample denoted in the joint y axis TCF. The leftmost y axis annotation denotes TCF for sample 1 (Tx) and the inner annotation denotes that of the second sample (Ty). The highest Tx was 0.95 and the lowest was set at 0.5. For Ty, the highest by default was chosen to be 0.2 lower than that of the highest Tx, hence 0.75 and the lowest was set at 0.25. The violin plots are drawn correspondingly under the heatmaps on the lower panel denoting the dispersion and central tendency of the estimates with significant p values of the paired association tests marked by grey points.

Next, we compared CRUST against some of the contemporary and frequently used clustering algorithms (sciClone, MAGOS)[11, 12]. All tumor biopsies were simulated as containing two samples with varied TCFs drawn from a lognormal-binomial mixture. At sequencing coverage of 300x all three algorithms were able to maintain a τ_median_ (median Jaccard index that varies between 0 and 1) of 0.95 with CRUST leading (τ_median_ = 0.99, **Figure 2C**). This pattern remained consistent throughout 100x and 50x simulations and all comparisons with CRUST were statistically significant at the 0.001 level. At 30x, CRUST had a lowered τ_median_ of 0.83. The interquartile distances between the τ_median_ estimates also markedly increased at 30x (range: 0.037 – 0.11). At lower coverages (≤ 50x) sciClone failed to predict the correct number of clusters often predicting as many as 6 clusters instead of 4 although CRUST and MAGOS predicted the simulated clonality status more accurately (statistically significant with two-tailed P value < 0.05, Mann-Whitney test) with at least 50x coverage **(Figure 2C)**[11, 12].

### Illustration of the impact of scaling for correct deconvolution

As an example of the importance of scaling, we extracted from a published dataset on childhood cancer[4], three neuroblastoma tumor tissue samples from a patient with varied TCF (55%-90% tumor cells), two from the primary tumor (NB12, P1 and P2) and one from metastatic relapse (R) (**Figure 3A**). Available copy number data and whole exome sequencing summaries were filtered for variants at a 1+1 allelic composition and sequenced at a depth of at least 100x, resulting in 32 variants. Rescaling the VAFs of two samples (P2, against that with the highest purity (P1), had a major impact on the subclonality prediction of the relapsed sample R (**Figure 3B, C I-II**). If unscaled, almost all variants shared among the three samples were predicted to be subclonal in sample R, contradicting their status as clonal (present in all tumor cells) in the other two samples (**Supplementary Table 1; *Scaling***). For example, the unscaled data predicted that a *SIAH1* mutation, was clonal in the primary but subclonal in the relapse which was rectified post scaling resulting in a prediction of clonal mutation across all samples. Only one mutation, in *ST8SIA2,* exhibited changed clonality status between two samples, i.e., the two regions of the primary tumor (**Figure 3C III-V**). This was indicative of a regional clonal sweep at geographic transition between these regions, an event corroborated by copy number profiling, which showed a subclonal copy-number neutral imbalance of chromosome 4 in P1, which transited to clonality in P2 (**Supplementary Figure 3**).

**Figure 3.**
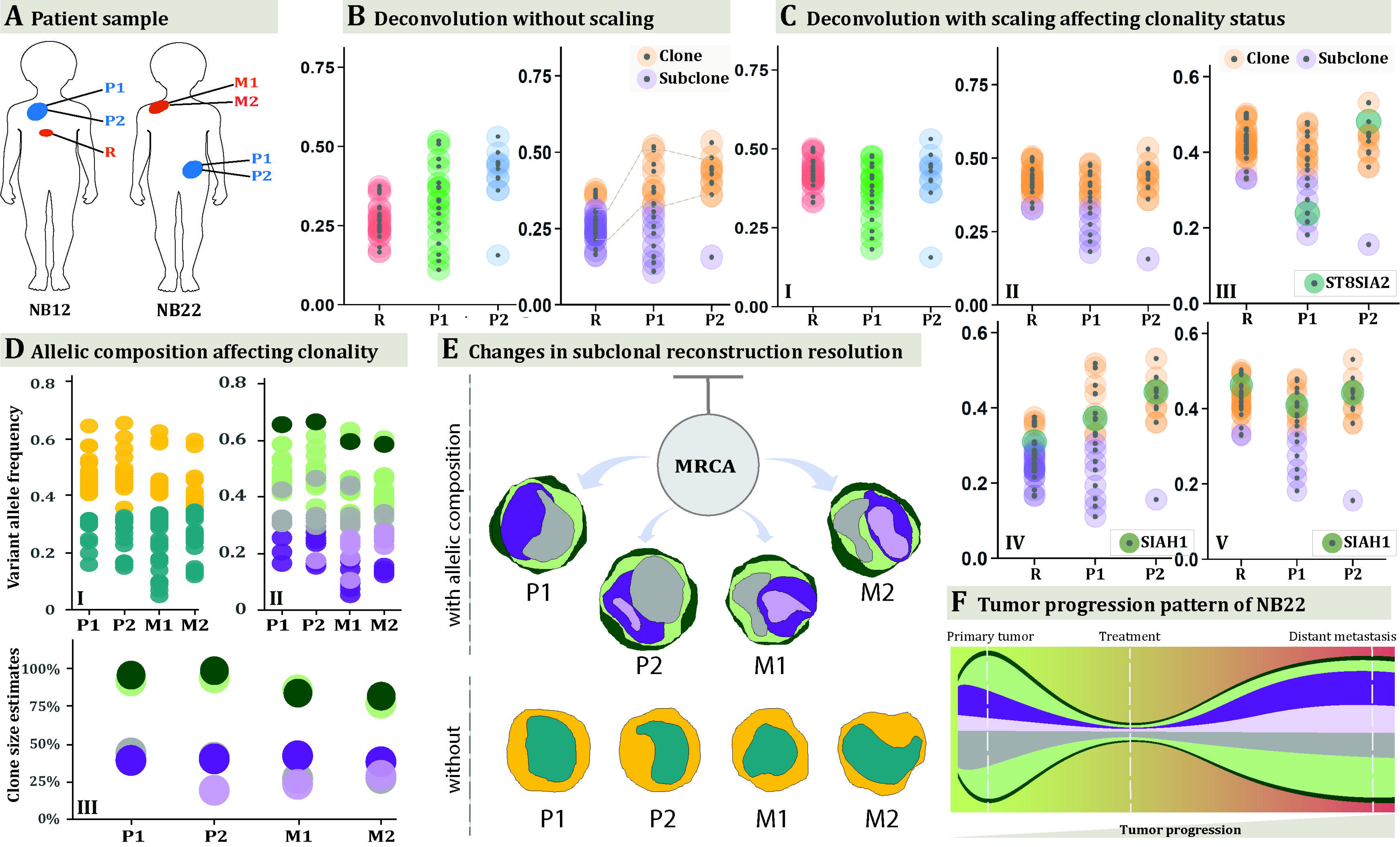
Tracing clonality trajectories across samples. Multiple samples with varied purity from each of two neuroblastomas (NB) were used as examples (**A**). NB12 is represented with three samples, two primary tumor biopsies (P1, P2) and one relapse (R). The primary sample was ∼90% pure whereas the relapse sample contains only 55% tumor cells as estimated by previous studies[4]. Hence, a deconvolution without rescaling the variant allele frequencies (VAFs) results in all shared variants (linked by grey lines between R and P1) being classified as subclonal in R (original sample specific VAFs are on the left, clonality predictions are on the right, **B**). Post-scaling (**C**), the relapsed variants re-adjust (**C-I**), and the predictions reflect a reasonable nature of the clonality (**C-II**). It is worth noting that in both analyses, the optimum cluster number is unchanged. This indicates that a traditional subclonality reconstruction algorithm would fail to account for the noise in the relapsed sample if analyzed in conjunction with the primary samples. The next three panels demonstrate how scaling impacts the predictions. In panel (**C-III**), an *ST8SIA2* mutation changes clonality status between P1 and P2, in concordance with a clonal sweep between these regions (see **Supplementary Figure 3**)[4]. Panels C-IV and V show a *SIAH1* exonic variant that is present in all three samples. In R, it is classified as subclonal while unscaled, but the prediction is overturned to be clonal post scaling. Deconvolution of the copy number aberrant neuroblastoma NB22 **(D)**, based on samples from the primary tumor (P1, P2) and a metastatic lesion (M1, M2). This tumor contained several copy number changes that required consideration for accurate deconvolution. CRUST was used to detect the segmental copy number alterations of all variants which were classified in two allelic composition make-ups, balanced 1+1 segments, and unbalanced 1+2 segments. These were deconvolved separately. Predicting clonality status without consideration of the copy number aberrations results in two predicted clusters **(D I)**, whereas considering allelic composition results in five clone/subclone clusters across all four samples **(D II).** This deconvolution would not have been possible without copy number data taken into account. Estimated clone sizes are depicted below with tumor cell fractions of each cluster (**D III)**. Inferring tumor evolution from deconvolution **(E-F)** shows how starting from an unknown MRCA (most recent common ancestor) one of the primary clones (in grey) shrinks whilst another subclone (in light purple) expanded at metastatic sites. Clone sizes estimated from the set of variants with two different allelic compositions indicated a major clone size (1+1 in dark green and 1+2 in light green) of about 92% (mean) indicating the aberrations carried forward from a most recent common ancestor. The bottom panel in (**E)** devoid of copy number data lacks resolution to detect any such change.

### Illustration of the importance of accounting for allelic copy numbers

As an example of how CRUST improves deconvolution by accounting for copy number variations, we then analyzed four different patient samples (NB22; **Figure 3A**) comprising of two primary (P1, P2) and two metastatic tumor samples (M1, M2)[4]. CRUST is inherently dependent upon variant specific allelic composition information. Although best practice is to assimilate sequencing summaries with separately obtained SNP array for a precise allelic copy number estimate, CRUST contains a function to approximate copy numbers from sequencing data on variants called from the constitutional genome. To identify distinct aneuploidies across samples it graphically presents segmental allelic imbalance and average log-relative coverage as done elsewhere[15]. We compared the segmental plots generated from SNP array data and that estimated by CRUST (**Figure 4**). The allelic imbalances estimated from exome sequences closely resembled those obtained from the SNP array, but with slightly less fidelity (**Supplementary Table 1; *phs000159_seq*)**. The 1q gain, 6p gain, whole chromosome 7 gain and, 17q11 loss and distal 17q gain were clearly identified with the estimates (**Figure 4, Supplementary Table 1; *phs000159_iontorrent***). Overall, the CRUST-based copy number estimation resulted in only a small number of discrepancies (2.7%) in the estimated allelic compositions compared to the available array-based estimates.[4] These were removed prior to analysis (**Supplementary Table 1; *NB22_copynumber***).

**Figure 4.**
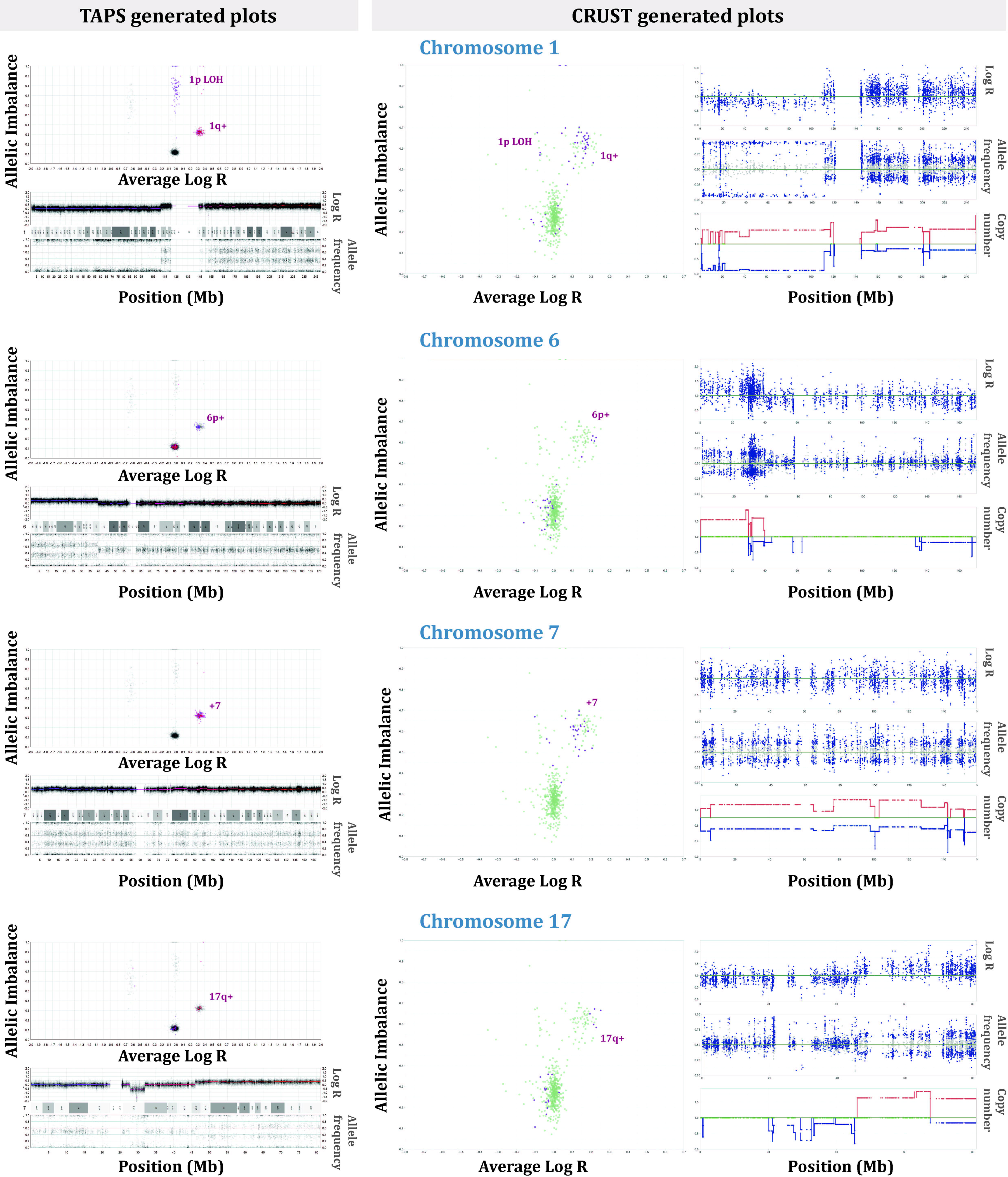
Estimation of copy number with CRUST. The segmental copy number estimates generated with CRUST are compared with SNP array profiles from the same tumor (NB22). We generated chromosome-wise plots of estimated segmental allelic imbalances against corresponding average log transformed relative coverages. In the left panels the plots are generated with TAPS from SNP array data[15]. In the middle panel the same plot is generated with data estimated with CRUST (highlighted chromosome in red). The rightmost panel includes three figures per chromosome, from top to bottom representing the estimated relative coverage, allelic frequencies and of segmental copy number. As demonstrated by TAPS, clones with different allelic compositions would show up at unit-separated distinct clusters along a fixed tangent in the allelic imbalance plots and the corresponding subclones would appear with a slight departure in the Y and X-axes. The allelic imbalance plots generated with CRUST retained the copy number specific cluster structures albeit with compromised resolution. The estimated copy numbers are close to that seen with SNP array. Larger events such as 1p gain, 6p gain, chr7 doubling and 17q loss are very clear. We recommend the users to consult these plots to verify the CRUST estimated allelic compositions.

To elucidate geographical makeup and temporal evolution of NB22 biopsies VAFs were then scaled against a diploid background and tumor cell fractions were calculated with allelic copy numbers considered. This revealed a varied tumor architecture across samples with evidence of polyclonal seeding of the metastatic sites, well in accordance with previous analysis of this case based on copy number alone (**Figure 3D**)[4]. Disregarding the copy number information, we reanalyzed the data assuming a balanced copy number state (1+1) for all chromosomes. The resulting deconvolution failed to pick up between-sample variations in clonality with considerable loss of resolution at backtracking of clones into geographic domains (**Figure 3E**). By integrating copy number and sequencing data, CRUST thus revealed details in the evolution of this tumor that would have had passed unnoticed if copy number data were not considered **(Figure 3E-F).**

### CRUST resolves clone topographies in published datasets

We extracted publicly available whole exome sequencing (WES) data on 20 multi-regionally sampled local primary non-small cell lung cancer (NSCLC, adenocarcinoma) from the initial release of the TRACERx project[16]. Deconvolution with CRUST, including copy numbers of mutated alleles in clone size estimates, made it possible to infer clonal topographies for all included samples at a level of detail not provided in the original publication (**Supplementary Figure 4**). Subclones could be distinguished from clones in all tumors and about 15% (median) of all variants in each tumor were predicted to be subclonal. All 20 tumors presented evidence of topological genetic diversity, i.e., variants were predicted to have changed clonality status (clonal crossover, clonal to subclonal and vice versa) between different samples of a tumor given the allelic composition remained unchanged, indicative of clonal sweeps across the primary tumor space. Some tumors were found to have an exceptionally high number of subclonal and crossover mutations, in particular CRUK003 (71%), CRUK004 (73%) and CRUK0018 (55%). This finding was well in accordance with the high proportion of branch mutations previously reported in these tumors (71%, 76% and 45% respectively)[16].

The relative proportion of genes predicted to be globally clonal or subclonal were sometimes different between CRUST and the original analysis (**Figure 5A**)[16]. For each patient, approximately 3% (median) of the quality-controlled variants (i.e., 1% of all variants) were responsible for clonal crossover. These belonged to about 5.6% of all annotated genes retained post quality control. Most of these crossover genes detected by CRUST (91%) were found to be absent from the published phylogenetic analyses[16]. A majority (18 of 31) of those retained in the original analysis were in fact predicted to concur between CRUST and the original analysis as subclonal driver mutations (ex. *TP53*, *KRAS*, *PIK3CA*, *CDKN2A*, *ATM* etc.) and the proportion of variants discarded from the original study resembled the proportion of crossover variants detected by CRUST (**Supplementary Table 2**). This indicates that CRUST adds value by resolving shifts in clonality status between samples.

**Figure 5.**
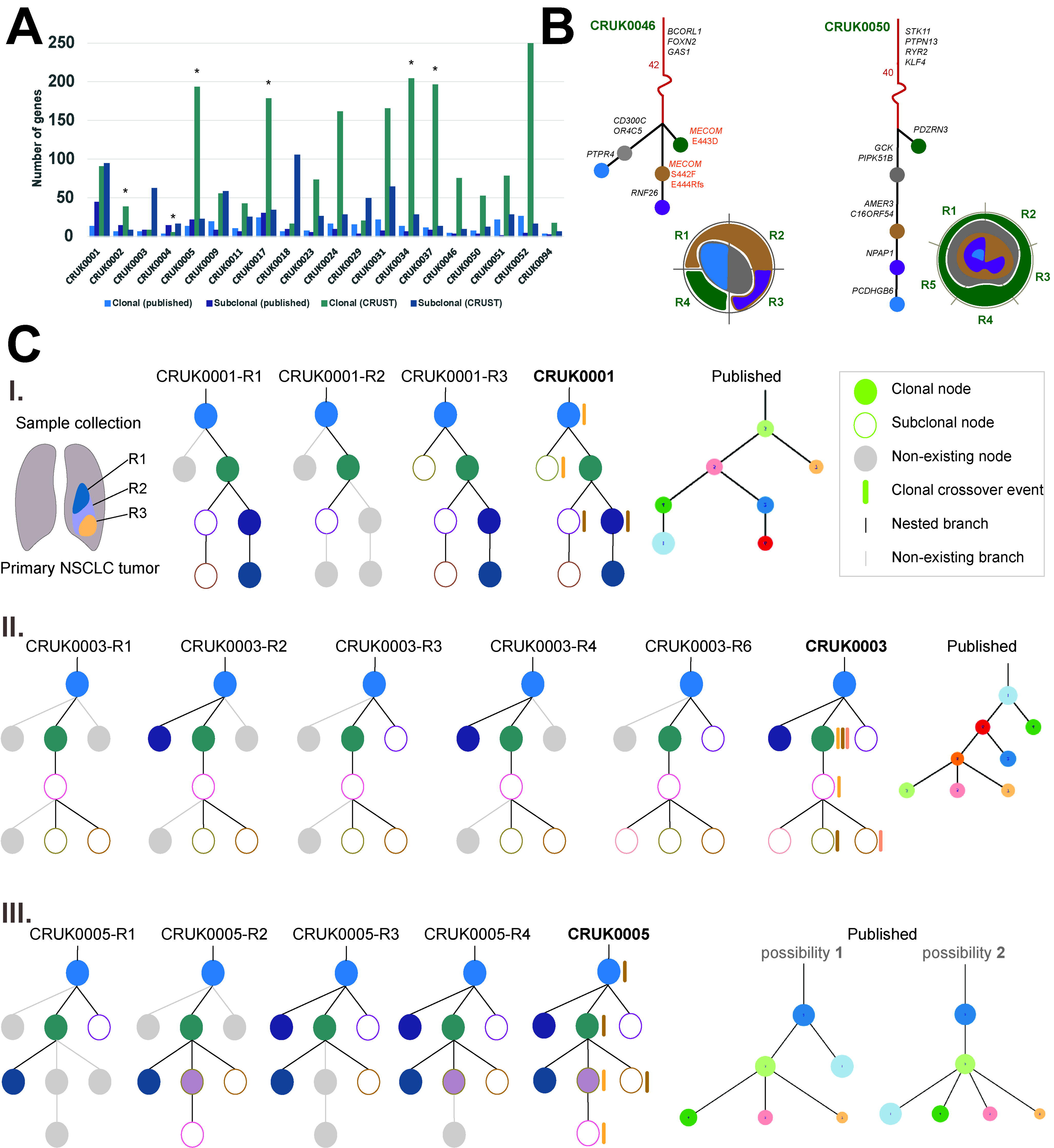
TRACERx Clonal geography. A comparison is made between clonality prediction from CRUST and that previously published (**A**). The bar chart shows the number of globally clonal and subclonal genes identified across all 20 samples in the two respective analyses. A non-parametric two tailed z-test is performed to denote significance difference (*P*<0.0025) between them, highlighted by star. In **B** phylogenetic trees of the two tumors, CRUK0046 and CRUK0050 are depicted along with the tentative sample specific clonal geography in the corresponding radial diagrams. The stems are in red with some of the noteworthy genes accompanying in black. Each subsequent branch are in black and the corresponding branch length is in proportion to the number of genes in each branch. The three variants corresponding the gene *MECOM* is highlighted for CRUK0046 as it depicts convergent evolution. The clonal / subclonal population nodes in the tree carry forward their colors in the respective clonal geography diagram. **C** depicts comparison of nested phylogenies between CRUST and the published analysis for three tumors, CRUK0001 (**C-I**), CRUK0003 (**C-II**), and CRUK0005 (**C-III**). Each sample has its own phylogenetic tree and all sample from a tumor is combined to build a consensus tree. If nodes and branches that are present in the consensus tree are absent from a sample-specific tree, they are greyed-out in the sample tree. Accompanied by the CRUST-predicted consensus trees, the corresponding published tree (s) is provided alongside. The clonal and subclonal populations are represented by filled and hollow circles respectively; clonal crossover events are denoted only in the consensus trees with colored bars alongside the clone (s) and the subclone (s) where the variants appeared. The largest clonal population in each sample are assumed to represent the stem of the phylogeny. Please refer to the original publication for color codes of published trees.

We then compared the proportion of driver mutations predicted by CRUST to be truncal with that suggested previously.^16^ The original analysis included 100 patients whereupon inferences were drawn with the constructed phylogenies. The CRUST analysis with 20 individuals retained approximately a third of the original set of annotated genes. When only this subset of 20 patients were considered, 63 genes in the published study were annotated as driver of which 31 (49%) were annotated as ‘subclonal driver’ in at least one patient. Post quality control, CRUST retained 56 of the said 63 driver genes among which 21 (38%) were subclonal in all samples along with 18 (32%) having undergone clonal crossover bringing the totality of subclonal drivers to 70%. Of these 18 crossover genes 12 were present subclonally at least in one of the other 80 patients analyzed in the previous study. This was in line with the original finding, as 75% of the tumors were observed to have acquired driver mutations late indicative of temporal fluctuations in evolution. The lacking spatial resolution of the previous study meant only 9 driver genes were inferred to have both clonal and subclonal status in different patients. One such gene was *PIK3CA*, for which crossover was confirmed by CRUST. This led us to speculate that crossover variants if seen in genes of lesser biological impetus had been misclassified as ambiguous observation previously. Omitting these might deprive the analyses of spatial heterogeneity as shown in several of the tumors (CRUK0001, CRUK0005, CRUK0051), and miss the presence of regional clonal sweeps (CRUK0003, CRUK0029, CRUK0050, CRUK0094).

To understand the impact of copy number consideration in clonal deconvolution, tentative clonal geographies were constructed for CRUK0046 and CRUK0050 (**Figure 5, B**). These tumors were selected as they had adequate number of samples (4 and 5) coupled with relatively simple ploidy profiles (up to tetraploid). In CRUK0046, we found several genes (*FOXN2*, *GAS1* etc.) in the stem of the phylogeny that are putative for lung cancer initiation, resistance, and metastasis[17, 18]. Even more interestingly, we noticed convergent evolution in different sites on the gene *MECOM*, commonly shown to be harboring structural aberrations in NSCLC[19, 20]. CRUK0050 on the other hand had mostly undergone linear evolution except for one monogenic branching. Here the tree stem consisted of several well-known mutations in lung cancer such as *STK11* (prognostic marker, aids drug resistance with fusion partner *LKB1*)[21, 22], *PTPN13* (tumor suppressor in lung adenocarcinoma)[23], *NOVA1* (promotes telomerase activity in NSCLC)[24], *ANXA2* (influential on lung cancer cell survival, apoptosis, as well as a construct of EGFR-fusion gene)[25–27], *RYR2* (associated with high NSCLC mutation burden in conjunction with exposure to high air pollution), and *KLF4* (regulates lung cancer initiation)[28]. In both tumors, the stem harboring most influential aberrations allude to these being early events in initiation.

To further understand the prevalence of crossover variants and their subsequent effect on the tumor clonal landscape, we constructed cross-sample phylogenies of the tumors and compared them with the published phylogeny. Three interesting cases (CRUK0001, 0003 and 0005) are highlighted in **Figure 5C**. For CRUK0001, the published and CRUST-inferred phylogenies were essentially the same, despite crossover variants being present across two different clone – subclone pairs. In CRUK0003 the crossover events resulted in subclonal diversification. The published results identified the distinct clones and subclones correctly, but the nesting structure was quite possibly different lacking the reinforcement from the three pairs of crossover events. CRUK0005 demonstrated how crossover events may aid in untangling ambiguity in phylogeny reconstruction. The published results included two possible tree structures for this tumor as the nested clones could be placed in different hierarchies. There were three distinct sets of crossover events detected by CRUST, which again pointed towards a greater degree of subclonal diversification than previously detected. This was consistent with one of the two phylogenies originally suggested. We conclude that CRUST adds significant detail by adding a component of cross-sample spatial to solid tumor evolution, but that sample specific purity estimates were unavailable in this (as in most) public datasets, possibly hampering the detection of false positive crossovers.

### CRUST deconvolution across platforms is stable but coverage dependent

We finally turned to a case of acute myeloid leukemia (AML), with samples available from presentation and relapse[29]. We selected a case (AML31) that had two biopsies sampled (primary and relapse) that were sequenced on several different platforms. The VAFs needed to be normalized as the samples varied greatly in purity (90.7% and 36.2% respectively). Post scaling, CRUST was able to identify a single clonal population existing in both the primary and the relapse samples while analyzing whole genome, whole exome, and a custom-made mutation panel (**Supplementary Figure 5A-C**). The predictions concurred with that obtained from sciClone. However, a custom ion torrent assay resulted in a clonal/subclonal separation in disagreement with others (**Supplementary Figure 5D**). While investigating the respective total coverage provided by all four technologies, we noted for the whole ‘*platinum list*’ of SNVs declared by the original authors, that the ion torrent assay had a median coverage of < 50x for both samples (**Supplementary Figure 5E**). To increase robustness, we therefore extracted only SNVs called at a minimum depth of 15x in ion torrent for both samples resulting in 33 SNVs (**Supplementary Table 1; *phs000159_iontorrent***), with increased median coverage of 79x and 88x respectively for the primary and the relapsed sample. Thereafter, the remaining variants in the primary sample attained a similar distribution as that observed by the other three technologies and scaling resulted in variants in the relapsed sample undergoing a similar magnitude of displacement, whereupon the centroid of the relapse sample VAFs realigns itself with that of the primary sample. This inferred a single clonal population, consistent with assessment by the other techniques (**Supplementary Figure 5F**). In all, this cross-platform analysis confirmed the result from simulations, indicating that sequencing depth is a limiting factor for deconvolution with CRUST.

## Discussion

Parameters based on clonal deconvolution and tumor cell phylogeny have been shown to carry prognostically essential information in a range of cancer types [30–32]. Such phylogenetic reconstruction of tumors has mostly relied on bulk sequencing data, although this is about to change with the advent of single cell analysis. However, because single cell DNA sequencing data are still limited by a high cost and a relatively low resolution, we anticipate more studies will take place where bulk sequencing is used to investigate cancer cell evolution based on multiple samples from the same tumor. Distinguishing clonal from subclonal mutations is a critical step in all studies where tumor phylogenies are deduced from such data.

Here we presented a parametric semi-supervised method of clonal deconvolution developed to interrogate variant clonality in multi-regional/temporal samples of a tumor. In comparison to most available tools for clonal deconvolution, CRUST has several major features including a robust normalization for purity, an inbuilt assessment and integration of copy number alterations, and a possibility to take à priori biological knowledge into account through user supervision. While it determines clonality with stochastic algorithms, depending on sequence quality variation between samples or technical artifacts, sometimes no mathematical model can adequately harmonize spurious signals. As the variance of each clonal subcluster inflates with compromised quality of sampling/sequencing, CRUST expands on the prediction with a non-parametric test indicating the probability of a variant belonging to a certain cluster that compensates for hard thresholding. Because copy number profiles are not always available by a dedicated method such as SNP array for sequenced tumors, CRUST can estimate copy numbers from sequencing datasets. However, there remain risks of detecting spurious signals if copy numbers are solely estimated by this approach. Hence, a dedicated estimation should always take priority and we would recommend strict monitoring of sample quality, purity, sequencing technology variation, variable coverage across chromosomes, unstable GC content scaling and other factors. Another quality issue arises from low sequencing depth, leading to allele frequencies unsuitable for scaling. Nevertheless, even at 50x coverage with at least one sample with 70% purity, the clonality determination was accurate in simulations free of artifacts.

Through CRUST, we have demonstrated how a semi-supervised model can yield biological insight with a purely mathematical framework to determine clonality of mutations. CRUST not only delivers a high-resoluton spatio-temporal clonal deconvolution of multi-sampled tumors, but also provides users a much-needed means for manual curation. The sequence of somatic mutation can thus be traced more accurately across multiple samples from a tumor especially in cases with a large burden of chromosomal gains and losses.

## Methods

### Quick user guide

Clonal deconvolution can be performed with CRUST following a minimal number of steps. The argument data needs to contain at least two columns specifying sample ID and the VAFs. Annotation for the variants are optional but is required to be able to create variant specific plots of the deconvolution. If sample(s) need to be scaled, an additional column should be provided with tumor cell fractions for each sample. The allelic composition of each variant corresponding to each sample should be available to the user. Here we will assume that SNP array data is not available to the user. Hence it needs to be estimated and we will provide a protocol for a sample analysis:

- First make sure the .vcf file contains calls from the constitutional DNA that enables CRUST to estimate copy number profiles. Each .vcf file should only have summaries for one tumor sample and one corresponding normal tissue sample. For each tumor sample this analysis needs to be performed separately.

a. Sample specific allelic composition can be estimated with the function *AlleleComp*. Input for this function requires the allelic depth (*AD*) identifier usually present in the *FORMAT* argument of the .vcf file. There are two compatible methods for this estimation that can be selected at user’s discretion.
b. A summary of the estimates can be visualized with allelic imbalance and copy number summary statistics if the estimation was performed with the ‘naive’ method. The *View_summary* function can generate this.
- The data thus obtained can be merged with the original data file containing sequencing summaries. As each distinct allelic composition have a distinct VAF distribution, the data needs to be analyzed separately for each allelic make up.

a. Before deconvolving, the VAFs need to be scaled according to the corresponding TCFs with the *seqn.scale* function.
b. scaled VAFs now can be used for deconvolution with *cluster.doc*

I. User is here asked to provide a postulated clonal/subclonal composition of the variants from visual inspection of the dot plot
II. A measure of clustering accuracy in terms of Bayesian information criteria or Smin statistics is provided to the user along with the best (statistically) suggested clonal deconvolution, which the user is free to retain or override (**Supplementary Figure 1)**.
- Depending on biological agreement and objective plausibility, the user can choose to fit different deconvolution structures with *cluster.doubt*
- Variant specific plots of the deconvolution can be obtained with the function *variant.plot*. It is also possible to automate this process and obtain plots for all variants with *variant.auto.plot*
- The clonality estimates thus obtained can be used to estimate clone sizes and to formulate phylogenies. We estimated clone sizes for the example dataset named *Neuroblastoma*[33] provided along with CRUST.

### Study design and data preparation

Clinical samples included here were fresh frozen tumor biopsies analyzed as a part of a larger study[4]. The DNA was extracted with the AllPrep DNA/RNA/Protein Mini kit (Qiagen); segmental aberrations were analyzed using the Cytoscan HD platform (Thermo Fisher Scientific/Affymetrix). Whole exome sequencing of neuroblastoma samples were performed by SciLife lab (Stockholm, Sweden; Illumina) on NB12 (two primary and one metastatic relapse sample) and NB22 (two primary and two metastatic relapse samples). Subsequently a targeted resequencing on all NB22 samples and the two NB12 primary samples were performed based on the variants identified by exome sequencing at SciLife lab (Uppsala, Sweden) using the AmpliSeq technology for design and Ion Torrent for sequencing (both from Thermo Fisher Scientific). Further details on human sample collection, preparation, relevant technologies, quality control measures and basic bioinformatic analyses are described elsewhere[4].

### Baseline normalization of VAFs

Human tumor samples are often contaminated with non-neoplastic cells, e.g. local epithelium, fibroblasts, endothelium, pericytes and, immune cells. Even though these cells are expected not to carry the clonal somatic mutations present in the tumor cells, they can affect the VAFs of such variants by a dilution effect. A quantitative measure of sample purity is given by the tumor cell fraction (TCF). It can be estimated from either SNP arrays, sequencing or methylation data[34–36]. Given that TCFs may vary among samples of a tumor procured at different sites or time points, variants may be present with altered VAFs even though the relative abundance of them among neoplastic cells are identical in the respective samples. On the contrary, they can also appear to be identical despite being influenced by selection or genetic drift that significantly changed their relative frequency among tumor cells. A normalization strategy can realign the VAFs of a variant from several different samples, given a common baseline such as the TCF.

It is customary to avail an integral normalization in presence of a k-dimensional reference space to transform the n-k dimensional scalable space, assuming the total integral uniformly influences dilution of all signals in the space. In its stead, we incorporated probabilistic quotient normalization of VAFs based on sample specific TCFs, as quotient normalization assumes that changes in unique elemental signal dilution affects only the signal of that element in the complete spectrum and dilution in global signal of a spectrum influences that of the overall spectrum[37]. This normalization algorithm is carried out between spectra where,

a. Integral normalization is performed on the scalable spectra.
b. Median spectra of the reference sample are calculated; lacking a reference spectrum, a control sample can be obtained from the scalable spectra but is advised against.
c. Quotients are calculated for the scalable spectra scaled against that of the reference.
d. Scalable spectra are normalized with the median of the quotients.

As the reference spectrum is same as the TCF and VAF combination for the highest quality sample, it is assumed to be devoid of (detectable) signal dilution. Hence, we realign VAFs of all other samples against their corresponding purity scaled with the departure in TCF between that sample and the purest one.

### Cluster detection and assignment

We use VAFs as the metric for clustering as is popularly used in the clonal deconvolution literature. VAF is defined as the relative frequency of read count pertaining to a single variant and is calculated by

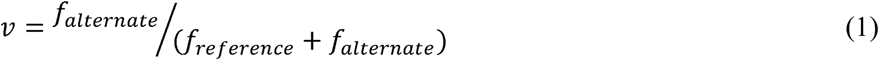

where *f* denotes read depth of an allele, alternate allele is the mutate allele of a presumed biallelic variant, and the reference allele is the wild type allele.

We take a set of *N* variants that may be observed in *S* samples procured from a tumor. Given the set of *S* samples, VAF of the *i^th^* variant can be represented by *v_i_* for a set of *k* VAF values where 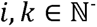 and 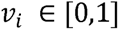 obtained from (1). Hence, we find a *K* dimensional matrix 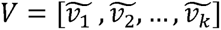. To determine clonality of the variant space *V*, we employ two different techniques: first, parametric stochastic modeling with Gaussian finite mixture modeling [38] and second, bootstrapping estimated cluster stability modeling [39]. Here *V* can be interpreted to be a random sample of *K* independent identically distributed random variables. A joint probability function defined with a finite mixture model of *C* components takes the form

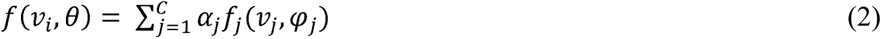

Where the parameter space of (2) is defined by 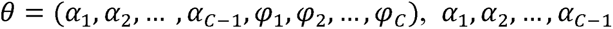 are the weights of the *C* mixture components given 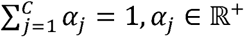; and marginal density function of *v_j_* is 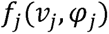 with parameter *φ_j_* Applying the assumptions of a gaussian mixture model, the marginal density of each mixture component follows a multivariate normal distribution N(*μ_j,_Σ_j_*). As each of the mixture component represents an ellipsoidal mutually exclusive cluster, these are centered at *μ_j_* and the shapes are defined by the variance-covariance matrices *Σ_j_* respectively. Henceforth, Bayesian information criteria is computed with penalized log likelihood so that with increasing likelihood in proportion to increasing number of mixture components, information loss is incorporated with the logarithm of the number of estimates. Thus, CRUST determines the optimum number of mixtures. Decided the ideal clustering parameter, the algorithm performs hierarchical multifactor analysis to obtain cluster centroids[40]. With the centroids determined, sample points are assigned to clusters with supervised *k*-means clustering[41]. To elaborate on the features of the data we include a provision for multivariate mixture component. This method can be utilized on such data with dimensionality reduced sample space, although for univariate samples of VAFs the first principal component and the original vector is identical.

In addition, the stability of the clusters can be assessed with strength of the clustering method[39]. This technique relies on bootstrap resampling and iterative clustering. As the resampling is performed *B* times, the minimum mean Jaccard index is observed and scaled over the *B* resamplings. With an increase in the number of predicted clusters over the number of true clusters, the stability (defined by *S_min_*) decreases affecting the Jaccard index as a given independent cluster is split in random sub-clusters. We leverage *S_min_* to obtain the number of clusters devising partitions closest to true separation in VAFs. The optimum number of clusters thus observed is used to fit a *k*-means clustering algorithm with the centroids obtained from the *B* resamplings.

### Clonality determination

In absence of clone size estimates corresponding to a specific variant, a cluster-wise detection of subclonality by available algorithms often lacks prediction of clonality. If the longitudinal sequence in which a variant appears in the tumor is not apparent, then we cannot distinguish a clonal event from a subclonal one. We conceived a metaheuristic process to determine the clonality of identified clusters. First, an iterative unsupervised density-based neighbor joining algorithm is used to cluster the VAFs by varying epsilon boundaries from 0.1 to 0.3. The epsilon boundaries define the minimum margin around the cluster centroid which passively characterizes the distance between the centroids at initiation. In a diploid, balanced copy number state, the separation between centroids of a clonal and a subclonal cloud is expected to lie in this range[13]. An emergent feature of this unsupervised clustering with the prediction obtained from the algorithm above is that a pairwise contingency comparison of the two produces a matrix ( ) spanned with either mutually orthogonal vectors or that belonging to same subspace i.e.,

If 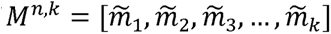

where *n* is the optimal number of predicted (or user defined) clusters and *k* is that obtained from the density-based algorithm; then,

either 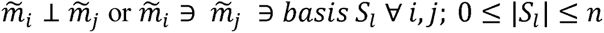

This property ensures that a set of predicted clusters is reduced to a set of vectors belonging to the basis of a *d* dimensional space that spans the vector space defined by the *n* clusters. The orthogonal vectors and corresponding clusters are linearly independent as the linearly dependent clusters are merged to reduce the matrix to have *d* rows. Hence, we obtain a set of clusters with cardinality *d* ≤ *n*. The clonality is assigned sequentially thereafter with a reductionist assumption that the subclones in a sample are represented in the left heavy tail of the VAF distribution if noise corrected[13]. This is justified by largely late-arriving subclonal events and stronger signal dampening experienced by rarer variants[42].

### Estimation of allelic composition

Somatic copy number alterations are notoriously capable of modifying the relative abundance of a variant from sample to sample. Since subclonal copy number events can often occur during cancer progression, stratification of VAFs by allelic composition produces a better clonality prediction. CRUST stratifies variants according to baseline copy number at start and performs separate analyses for each stratum. In the absence of SNP array or other specific copy number data, the allelic composition is estimated from two separate standalone processes that enquires somatic tumor variants along with polymorphisms in the constitutional genome [43, 44]. These predict variant specific copy number by modeling mean relative depth ratio and segment specific B allele frequency for each estimated segmental copy number alteration, adjusting for cellularity, and overall tumor ploidy. Estimation of allelic segmentation from bulk sequencing data are known to contain several sources of bias. GC content is one such major source of bias which induces an undulating pattern in allele frequencies [45]. As a measure of quality control, prior to copy number estimation, CRUST normalizes the B allele absolute frequencies against small segmental GC content with penalized lasso regression[46]. Additionally, it rescales the same against aggregate allele frequency to account for technical artifacts. Subsequently, segmental copy numbers are estimated with *copynumber* or *falcon*[44, 47].

CRUST visualizes segmental copy numbers by estimating allelic imbalance ratio and average log relative coverage ratio. To this end, both are estimated over genome-wide dynamically created chromosomal segments. A principal component analysis is then leveraged for distance-based pruning to discard outlying variants in each segment. Allelic imbalance ratios are calculated with mirrored B allele frequencies as described elsewhere[15]. As median log ratio is assumed to reflect segmental copy number and allelic imbalance ratio dictates the relative state of segmental zygosity, the unknown states of segmental copy number are reflected in the cluster positions of the segments. Presence of a subclonal copy number event is thus indicated by departure of a smaller segmental cluster from its parental cluster.

### Calibration on simulated data

We presume the apparent alteration in representation of a variant in samples from the same tumor, given the same copy number state, is due to variation in the TCFs in each sample. The following stochastic modeling of the variability in variant abundance generated synthetic sample data closely resembling that of a solid tumor. Leveraging the heavy tailedness of log-normal distribution, a random variable with varying parameters drew sample points representing departure in VAFs from the true distributions due to fluctuating TCFs. We varied the mean of the distribution according to the logarithm of the re-centered true mean and re-scaled empirical variance in three separate set ups. Assuming a 100% purity and copy-neutral (disomic) ploidy status, true mean of VAFs for the clonal variants was assumed to be at 0.5 and that of the subclonal variants at 0.25[48]. First only mean then only variance and subsequently both mean and variance dynamically were made to affect the simulations by changing the nature of variability in the data (**Supplementary Figure 2A**). The samples were drawn with these three different strategies to compare how restructuring the clonality of the variants may affect the deconvolution efficacy. For all iterations, the TCFs were sequentially varied from 0.95 to 0.5. We used polychoric correlation estimates computed between scaled and unscaled predictions to measure level of concordance[49].

As sequencing depth contributes immensely towards reduction of the baseline noise in signals, the following adaptation describes a probabilistic model conceived for the above-mentioned simulation. VAFs from a well-covered region can be reliably estimated with a beta distribution[12], thus a beta-binomial process with beta priors can be used for simulation[50]. To incorporate a log normal deviation based on TCF, we rendered a convolution instead. A joint distribution of beta-log normal was used for this[51], the probability density function of which is thus defined by,

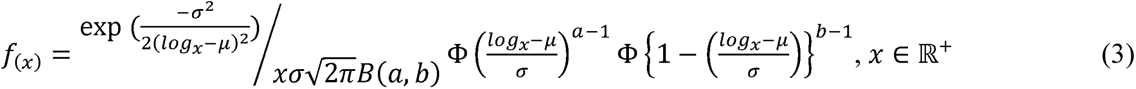

It jointly varies on the aggregates of the parameters of the two marginal distributions (a, b, µ and σ). The re-parameterized cumulative density function given by *F_(y)_* was inversed to obtain a canonical population of VAFs with subclonal mutations. A standardizing transformation on *x* in (3) gets us,

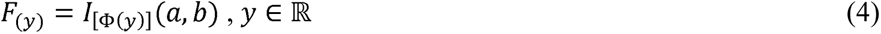

Where 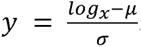. This method produces the prior distribution of a hypothetical sample. For a true clonal population, we center the marginal beta parameters with mean 0.5 (0.25 for subclonal) and variance 0.001. If these are denoted as a random variable 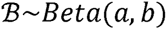 then, a closed form of the beta-log normal variable is given by *X* = *^eσ^*^Φ–1*(b)+*^ *^μ^*. Provided an expected sequencing coverage of λ, if *n_sm_* denotes the number of reads for a mutation *M* of sample *S*, then *N_sm_∼Poisson(λ)*. Presuming the variants are all biallelic, we can further estimate the read count *r* of an allele of the mutation following *r_sm_*∼*Bin*(*n_sm_*, *f*_(*x=sm*)_). Hence, the VAF for that allele is 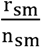. This process progressively aggregates noise as the coverage downsizes and the TCF degrades (**Supplementary Figure 2B**).

### Performance testing of scaling with simulation

We simulated an admixture of clonal and subclonal variants assuming a balanced background copy number. A parametric setup for the sample generation was favored under the assumption that sample purity indirectly affects the observed distribution of the relative abundance of variants. Three separate parametric assumptions were tested to obtain mutual concordance in prediction. All formulations are described in Methods and will here be referred to by the *mean*, *variance* and *meanvar*. For all three assumptions, 1500 simulations were performed respectively to generate two samples in each iteration. TCFs were sequentially modified for each iteration to have produced 100 variants for every single sample. As means vary, the clonal distribution and the subclonal tails are linearly translated on the x-axis which does not affect the clustering if the respective centroids are not too close. When variance was changed, the deconvolution and prediction accuracy broke down fastest as the two VAF distributions could retain the centroids while the range increased making them overlap. Hence the VAF distributions overlap between clusters making them virtually impossible to segregate.

Next, we compared CRUST against some of the contemporary and frequently used clustering algorithms, i.e. MAGOS and sciClone[11, 12]. We refrained from comparison against another popular method PyClone as both MAGOS and sciClone reliably outperforms it[11]. As these methods predominantly are clustering algorithms, there were certain assumptions we had to consider for the sake of contrast. Although CRUST is built to handle non-recurring variants that can appear in a (or disappear from) samples extracted in different stages of progression from a tumor, sciClone and MAGOS operate under the assumption that each ascertained mutation is present in all analyzed samples. Therefore, we used this as a basis for the subsequent comparisons. We also operated with the presumption of samples being copy neutral and clonally consistent subclonal populations do not undergo clonal sweep or fixation. Each simulation produced two admixture samples for every run of the procedure. Coverages were varied to generate 4 distinct sets of observations at 300x, 100x, 50x and 30x depth. Among the two samples included in each observation, the first was consistently drawn with a TCF higher than that of the second, sequentially reducing both with every iteration. The TCFs were thus progressively lowered from 0.95 to 0.5 for the first sample whereas the other one was continually initiated with a departure of 0.2 in TCF from that of the former and was successively lowered until 0.25. Hence the theoretically lowest quality of sample was restricted with a paired TCF of 0.5 and 0.25 with 30x coverage. At each combination we drew 10 observations consisting of a pair of 10 samples with 500 variants each. A sample thus drawn consisted of one clonal and one subclonal population.

We measured the performances of each method with the Jaccard index (τ, also known as Tanimoto similarity index) to quantify cluster agreement by creating a contingency table between the original population assignments and the predictions[52]. This statistic varies between 0 and 1, 1 indicating the highest possible concordance and can be interpreted similar to a correlation coefficient. We discarded CRUST’s internal prediction on variant clonality justifiably compromising information in sake of ordinality and generalizability of the classifications. Here it is worth mentioning that sciClone has previously shown to perform reliably given at least a coverage of 300x[29]. Furthermore, MAGOS has been shown to perform almost as good if not better at 300x[11].

### Deconvolution on hematological tumor patient samples

We further assessed constraints due to sequencing depth using publicly available data from an acute myeloid leukemia patient (AML31; dbGaP: phs000159) consisting of two temporally separated samples corresponding to the primary (90.7% pure) and relapsed tumor tissue (36.2% pure). These samples were queried with deep whole genome sequencing (up to ∼312x), exome capture (up to ∼433x) and further ultradeep targeted sequencing with custom capture assays (>2000x). To ensure quality, we extracted summary statistics on a putative ‘platinum list’ of SNVs (consisting 1,343 high-quality validated sites) 8. Subsequent pruning with read count (>10) resulted in a total of 37 variants. Statistics from four assays were used for testing: whole genome sequencing, exome capture, custom capture, and custom ion torrent platform, each providing varied coverage (**Supplementary Figure 5**).

### Deconvolution and phylogenetics of TRACERx tumors

To observe the effect of normalization and inclusion of allelic composition on the predicted clonality status in a large-scale mutational landscape of a typical solid adult tumor, we turned to publicly available data. However, since procuring multiple tissue biopsies from several regions or at separate time points from human tumors are not part of standard care there are only a handful of dedicated studies set up for such investigations. The TRACERx[53] study is one such endeavor where initially upwards of a hundred non-small cell lung cancer (NSCLC) patients were biopsied for tumor tissue, some multi-regionally[16]. We extracted WES data on 20 of these tumors along with their copy number profiles for which at least three viable regional samples were sequenced with at least one mutant variant present in more than one sample. We extracted sequencing summaries on a total of 21,000 variants (median: 911) from the twenty patients averaging 260 single nucleotide variants (SNVs) per sample. Only 9,400 variants passed the quality control.

Copynumber summary data was procured directly from the published study and was linked to each variant to obtain allelic composition. All 20 tumors were subjected clonal deconvolution performed with CRUST where all multiregional samples were analyzed in tandem (**Supplementary Figure 6**). Each unique allelic composition warranted a distinct run. Due to unavailability of sample purity data, we were unable to normalize the VAFs. The clonality determination was supervised in accordance with the allelic compositions and in case of VAFs clustering too close to be clustered across samples, all variants were presumed clonal. Next, clone sizes corresponding to each variant’s VAF were estimated as follows[54]

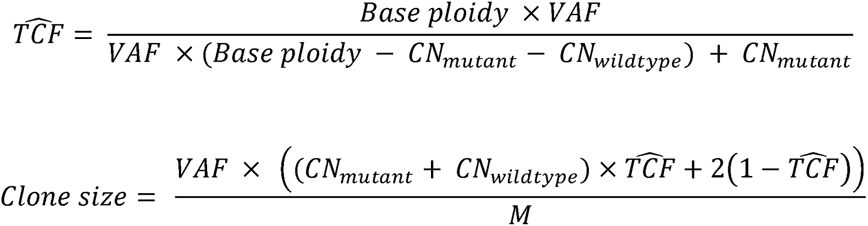

The base ploidy was determined as that of the clonal variant with the least total copy number; *CN_mutant_* is the copy number of the mutant allele and *CN_wildtype_* is that of the wildtype allele, *M* is alleles harboring the variant. We reconstructed the clonal or subclonal clusters based on these clone sizes. From top down, all clone sizes within 15% of each other were aggregated (separately for clones and subclones). The median clone size of each such aggregate were made to reflect the size of that population. The clone sizes needed to be scaled up as the estimates reflected the effect of purity. For each sample, a scaling constant was determined as follows:

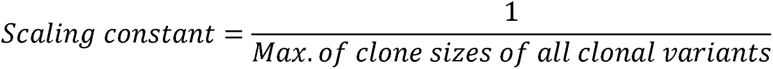

Clone sizes of all clonal as well as subclonal variants were multiplied with the scaling constant of the corresponding sample.

Additionally, we created spatially nested phylogeny of some of the tumors as done in the published study[16]. To this end clone sizes (for several clonal / subclonal clusters) corresponding to each allelic composition were aggregated under the assumption that variants with similar VAF and allelic composition belong to the same population. In case more than one such aggregate (with different copy number or from different samples) were seen to have similar clone size (i.e., within 10% of one another), they were inferred to belong to the same population. All such calculations were separately formed for clonal or subclonal clusters (see **Supplementary Table 2; *clonal nesting***). The population nesting pattern thus discovered were used to build phylogenies.

## Key points

- A new computational tool that predicts clonality of sequenced variants and help detect underlying evolutionary processes in a tumor
- Consideration of aneuploidy aids proper subclonal reconstruction which if absent jeopardizes phylogenetic analysis
- Multiregional sequencing can aid in uncovering spatial inclination of solid tumors increasing the genomic and etiological resolution

## Software availability

CRUST depends on R (>3.5.0) and is available for download from GitHub repository https://github.com/Subhayan18/CRUST

## Data availability

WES data on the twenty TRACERx tumors were extracted from cBioPortal (http://www.cbioportal.org/study/summary?id=nsclc_tracerx_2017). The SNP array summary of the TRACERx tumors were available in supplementary tables of the corresponding study[16]. AML samples are part of the *Whole-Genome Sequencing of Acute Myeloid Leukemia* study and is available via dbGaP (https://www.ncbi.nlm.nih.gov/projects/gap/cgi-bin/study.cgi?study_id=phs000159.v8.p4). The neuroblastoma samples are part of previous study[4]. Simulated data are generated with randomized seed using R and the test data sets are included in the package build.

## Authors’ contribution

**Conception and design:** S. Chattopadhyay, D. Gisselsson

**Development of methodology:** S. Chattopadhyay, D. Gisselsson

**Data acquisition:** S. Chattopadhyay, A. Valind, D. Gisselsson

**Analysis and interpretation of data:** S. Chattopadhyay, J. Karlsson, A. Valind, D. Gisselsson

**Technical support:** J. Karlsson, A. Valind

**Contribution towards manuscript:** S. Chattopadhyay, J. Karlsson, A. Valind, N. Andersson, D. Gisselsson

## Author description

S. Chattopadhyay, PhD is a post doctoral researcher in Lund University. His research focuses on mathematical modeling of evolutionary dynamics in cancer.

J. Karlsson is a research, PhD scientist in Lund University. She is researching genetic signatures and evolutionary trajectories in paedictric tumors

A. Valind, MD, PhD is a research scientist in Lund University focusing in tumor cell heterogeneity and evolution within individual tumors, both with a focus on biological issues but also on the clinical implications of these processes

N. Andersson, MS is a doctoral student in Lund University. Her research interest lies in modeling in silico processes to analyze tumor heterogeneity and reconstruction of phylogenies.

D. Gisselsson, MD, PhD is professor in Lund University heading the research group pathways in cancer cell evolution. His team combines high-resolution genomics on clinical samples with mathematical modelling and methods from species evolution to answer questions in cancer evolution.

## Acknowledgments

The authors acknowledge technical support from the Science for Life Laboratory, the Knut and Alice Wallenberg Foundation, the National Genomics Infrastructure founded by the Swedish Research Council, and Uppsala Multidisciplinary Center for Advanced Computational Science for assistance with massively parallel sequencing and access to the UPPMAX computational infrastructure. We would also like to thank the Swegene Centre for Integrative Biology at Lund University (SCIBLU) for assistance. This study was supported by grants to D. Gisselsson from the Swedish Research Foundation (2016-01022), the Swedish Cancer Society, the Swedish Childhood Cancer Foundation, the the Royal Physiographic Society, LMK Foundation and the Gunnar Nilsson Cancer Foundation. A special thanks to Chris Miller and Malachi Griffith for their assistance in procuring data from community resources (dbGaP: phs000159).

## Supplementary figure legends

**Supplementary Figure 1.**
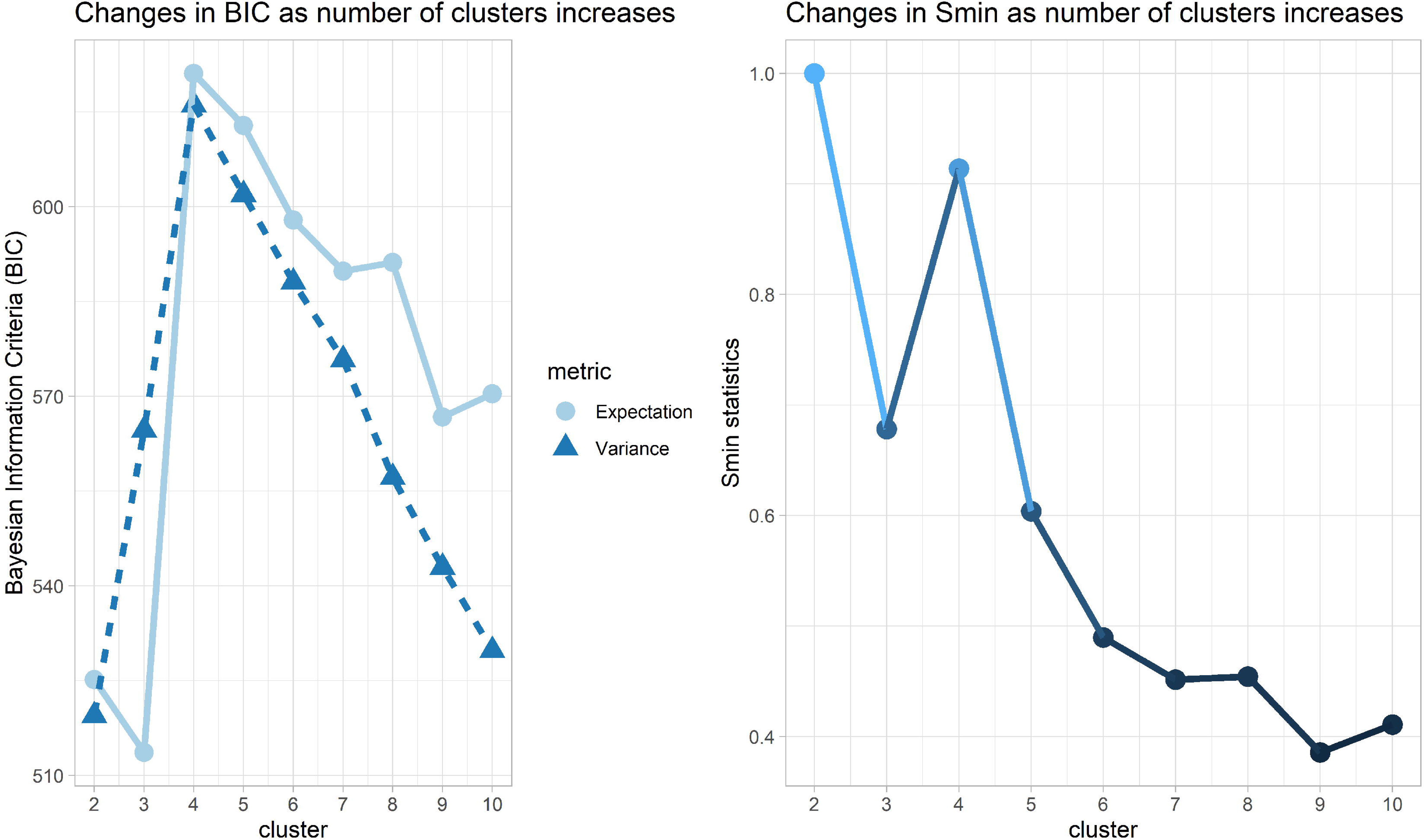
CRUST calculates the optimal number of clusters prior to deconvolution with two different methods (Gaussian mixture model on left, bootstrapping on right). The two methods correspondingly provide Bayesian information criteria or Smin statistics correspondingly which can be interpreted to be maximized with the least number of clusters to optimize the fit. In this example we see both statistics predict the optimal cluster to be four for the simulated data provided with CRUST (also used for deconvolution in Figure 1A-C). These predictions can be overridden by a user as shown in Figure 1.

**Supplementary Figure 2.**
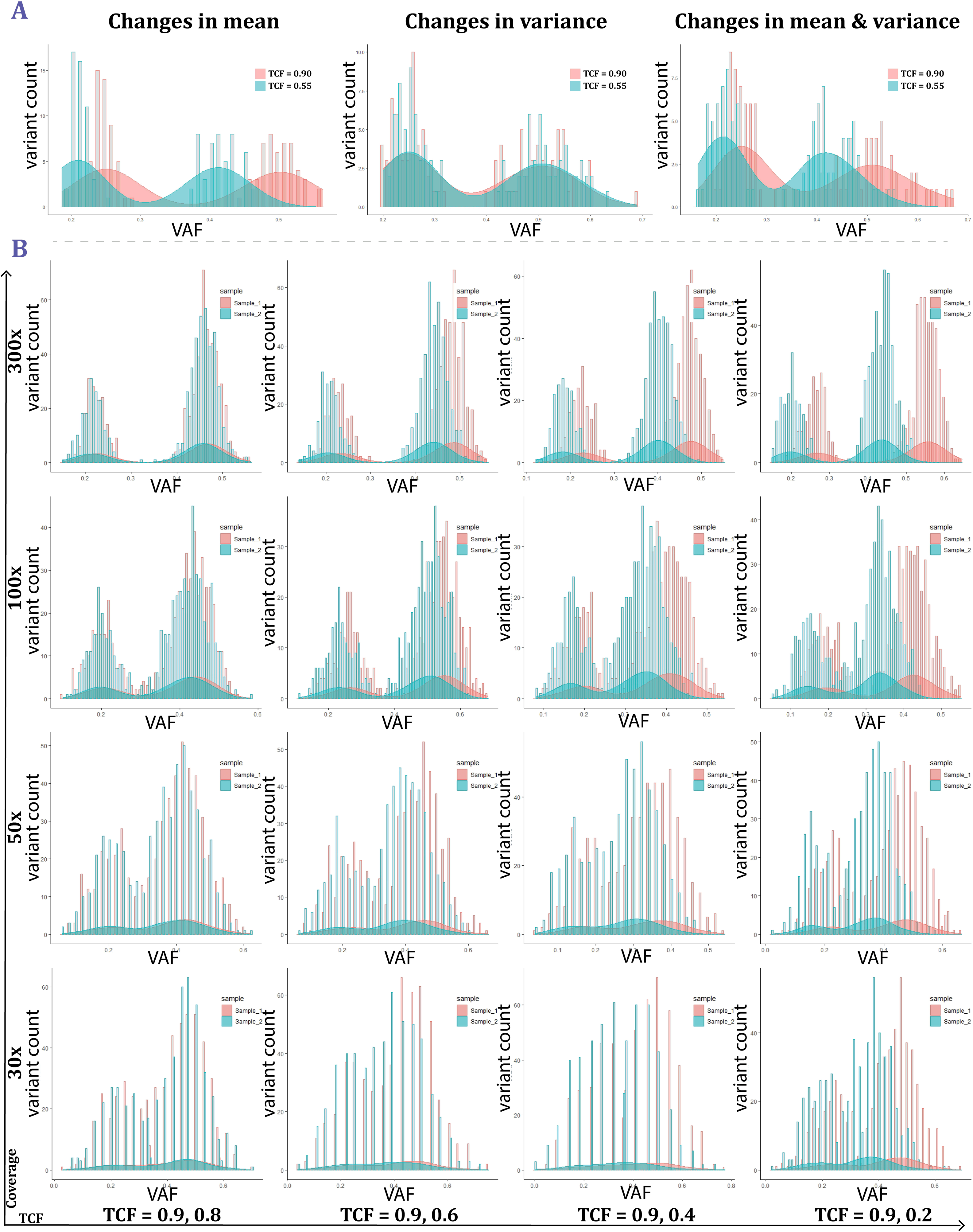
Effects of departure in location and scale parameters on the population distribution of variant allele frequencies (VAFs) is in (**A**). VAFs are beta-binomially drawn with mean and variance defined à priori. The population parameters are made to be influenced log-normally by the purity of the sample denoted here with tumor cell fraction (TCF). All three panels show population distributions of VAFs when two samples are drawn with the respective TCFs under different assumptions. The leftmost panel shows when the compromised purity is assumed to only affect the mean indicating that a low TCF result in a left tailed shifted VAF distribution, but the intra-sample heterogeneity remains unchanged. The middle panel demonstrates when TCF only affects variance keeping the mean unchanged i.e. low purity only increases intra-sample heterogeneity keeping the population mean unchanged. The right panel shows the changes when mean and variance are both impacted with change in sample purity. In (**B**) stochastic sampling with varying mutated clone fraction (TCF) is demonstrated. This is the result of a beta-log normal sampling procedure. Each panel corresponds to a coverage setting in the y axis and four corresponding panels on its right that are results of one single run of sampling for four different TCF combinations (T1, T2) denoted in the x axis where T1 is the assumed TCF of the first sample and T2 is that for the second. Each panel has histograms of VAFs plotted for two samples. Sample_1 in red is generated with T1 whereas Sample_2 was generated with T2 and is depicted in blue. As coverage declines (top to bottom along Y-axis), the ideal structure of the VAF distribution is compromised. A similar pattern also emerges for diminishing T2 (left to right along x-axis).

**Supplementary Figure 3.**
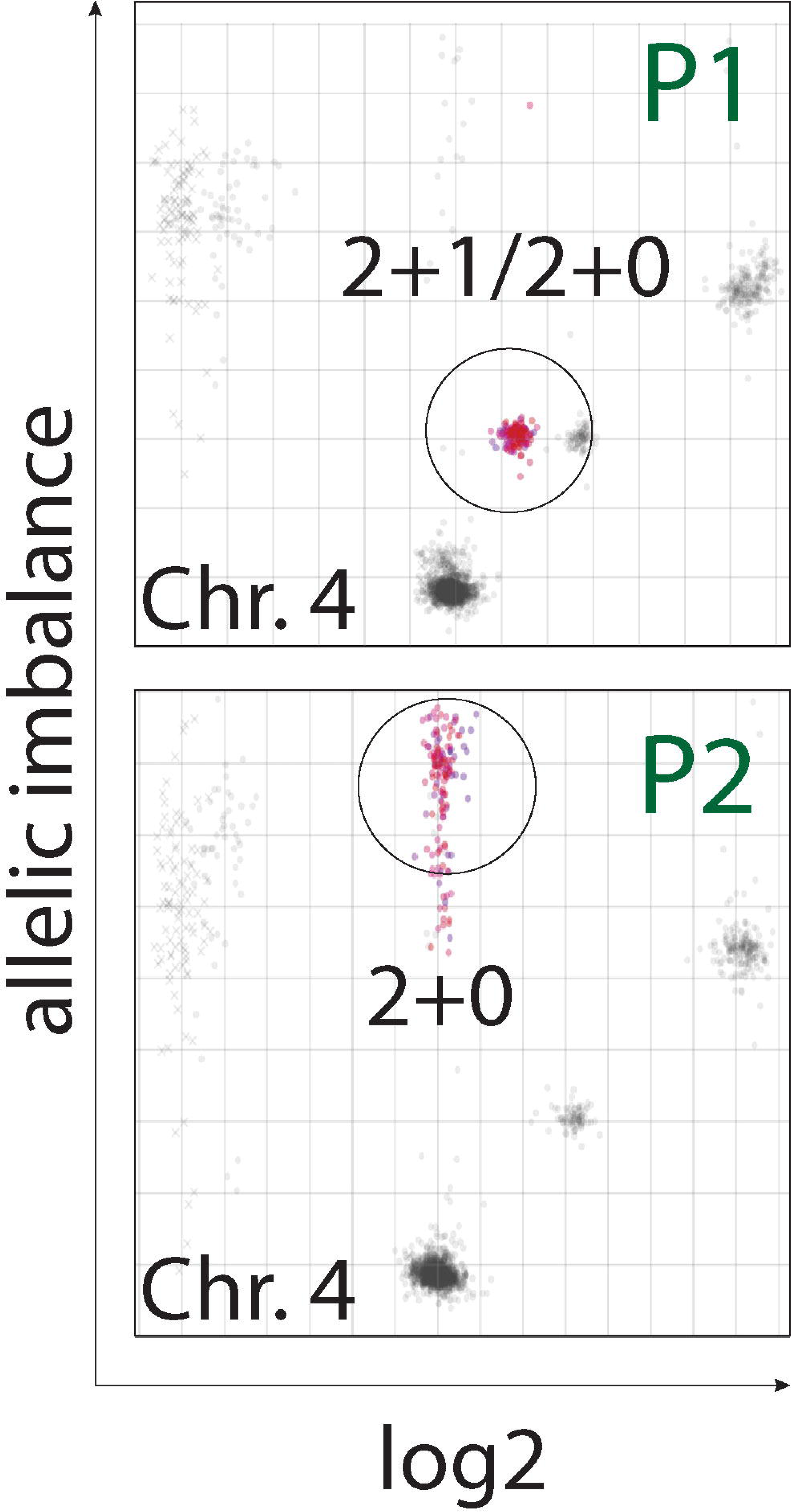
Allelic imbalance plots for two primary samples from neuroblastoma patient NB12[4]. Representation of chromosomal segments along the axes of allelic imbalance and log transformed relative copy number ratio creates clustering corresponding to allelic compositions[15]. All chromosomal segments are plotted for both samples in the two panels with those belonging to chromosome 4 highlighted in red. The top panel shows the red chromosome 4 cluster corresponding to a mix of cells with the allelic states 2+1 and 2+0 in P1, that in P2 alters into an almost pure population of cells with 2+0 (uniparental isodisomy). The results indicate a topologically restricted clonal sweep.

**Supplementary Figure 4.**
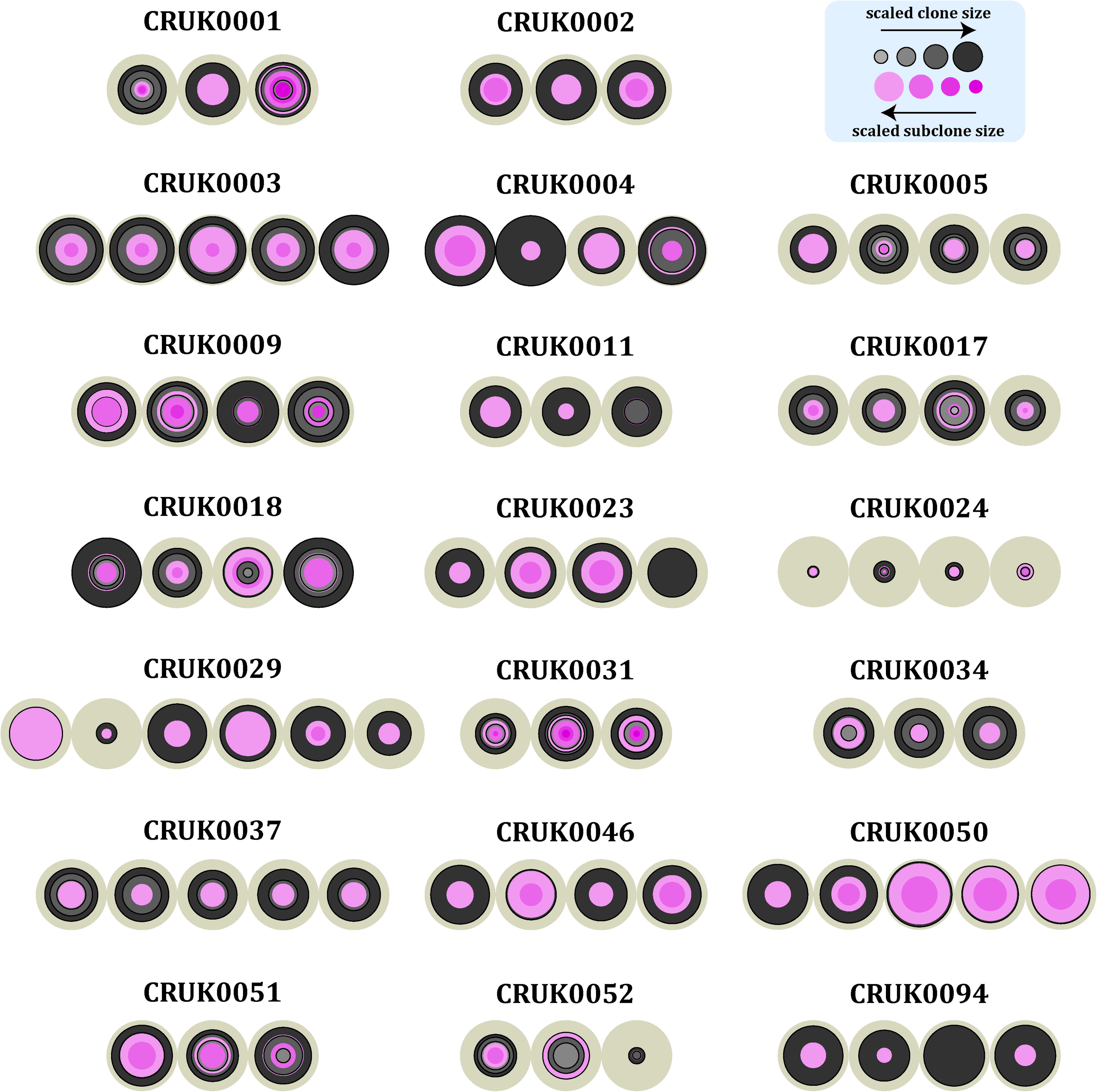
CRUST determines clonal compositions of the 20 NSCLC tumors from the TRACERx study. Each of the tumors is represented by three or more multiregional biopsies taken from the resected primary tumor. CRUST was used to determine clone sizes corresponding to each variant in a sample. The summaries can be found in **Supplementary Table 2**. Given the same chromosome, allelic composition and predicted clonality status separate clonal and subclonal populations were reconstructed according to the proximity of the estimated clone size. Clones (represented in grey) and subclones (in pink) are represented nested in each other as their size increases (not according to cellular phylogeny). Different contrasts are in use to indicate exclusivity of each population.

**Supplementary Figure 5.**
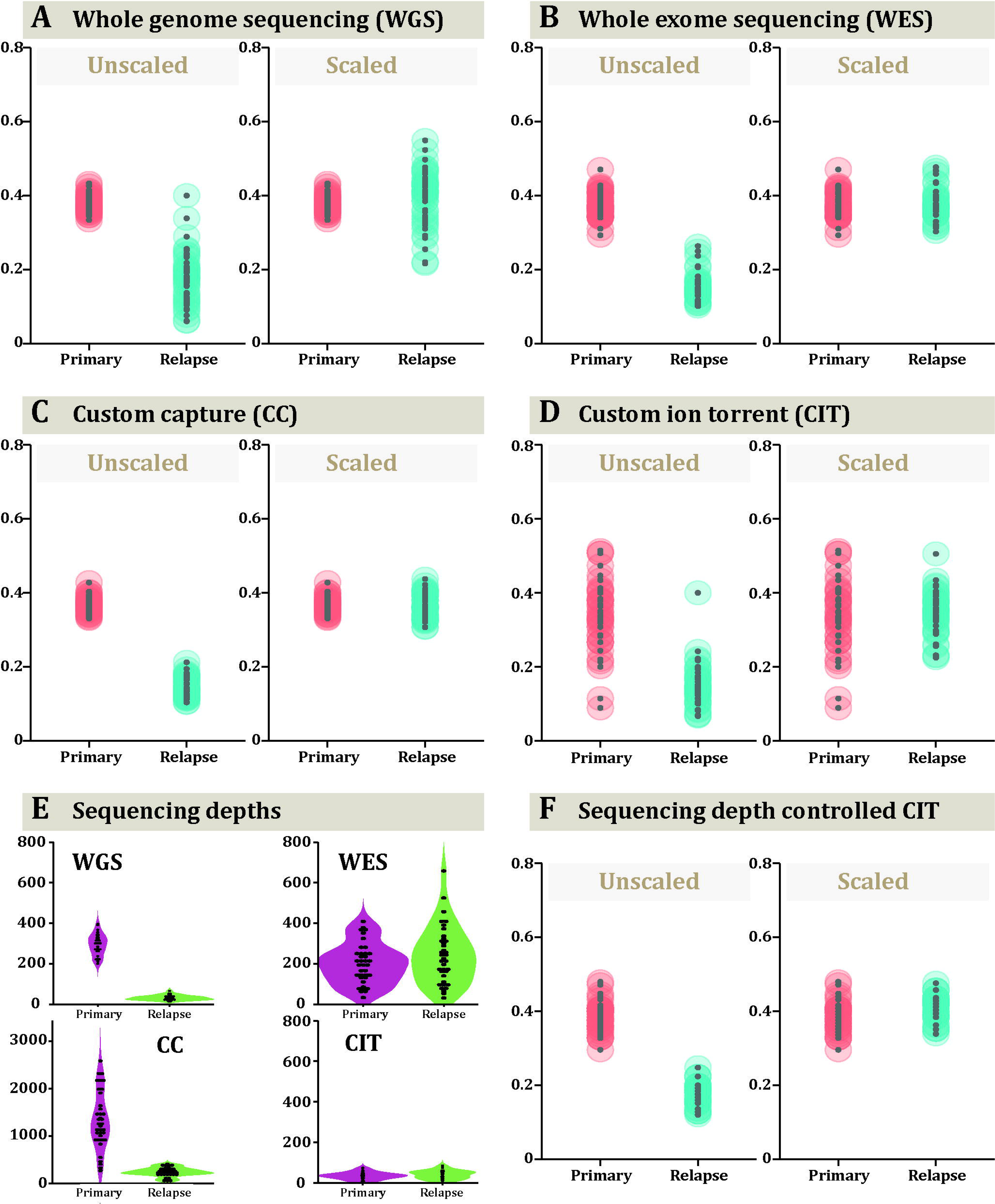
Clonal deconvolution of the patient sample AML31 extracted from the community resources (dbGaP: phs000159)[29]. The data includes sequencing summaries on two samples (a primary and a metastatic relapse). Data are shown on the effect of quotient normalization on the distribution variant allele frequency (VAF) data. On left are the original unscaled VAF distributions and on right are the same after normalization on whole genome sequencing (**A**), whole exome sequencing (**B**), targeted custom capture (**C**) and custom ion torrent sequencing (**D**). Violin plots of the sequencing read depths for the four technologies are shown in (**E**) (WGS: whole genome sequencing, WES: whole exome sequencing, CC: custom capture, and CIT: custom ion torrent). Variants with very low coverage were removed to create a pruned data from custom ion torrent (**F**), thereby identifying a similar VAF distribution as seen by other methods with normalization having similar effect.

**Supplementary Figure 6.**
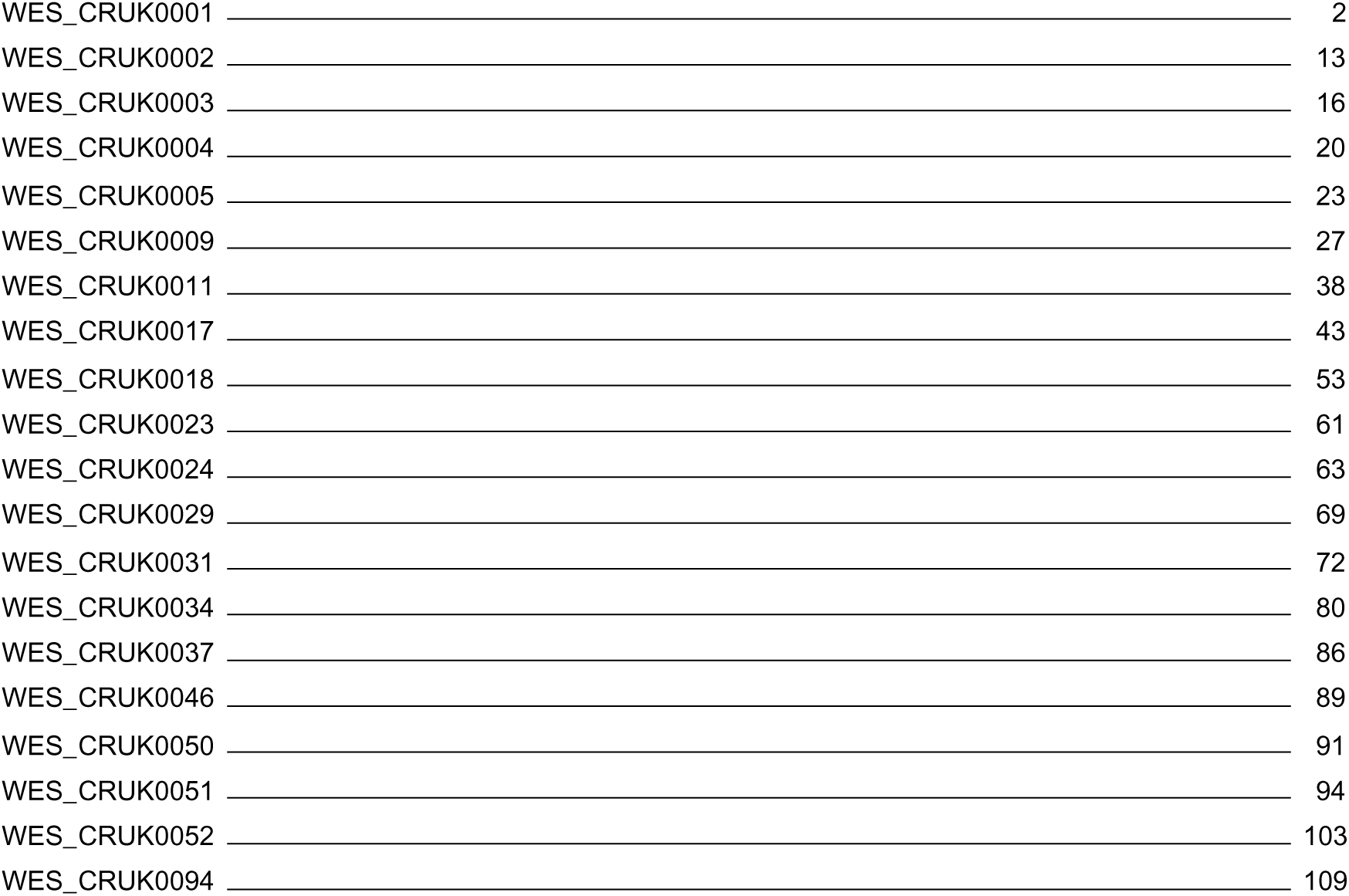
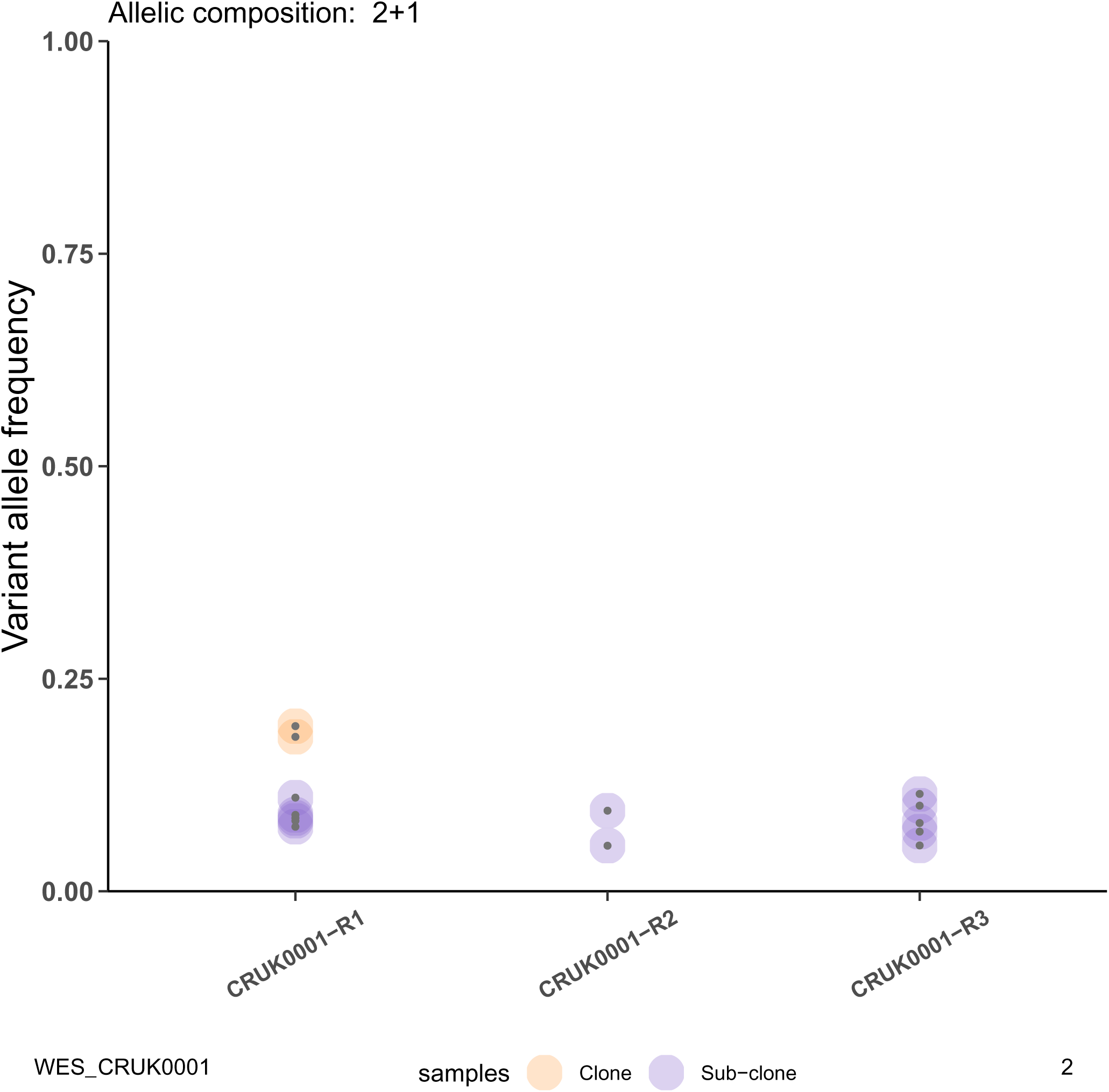
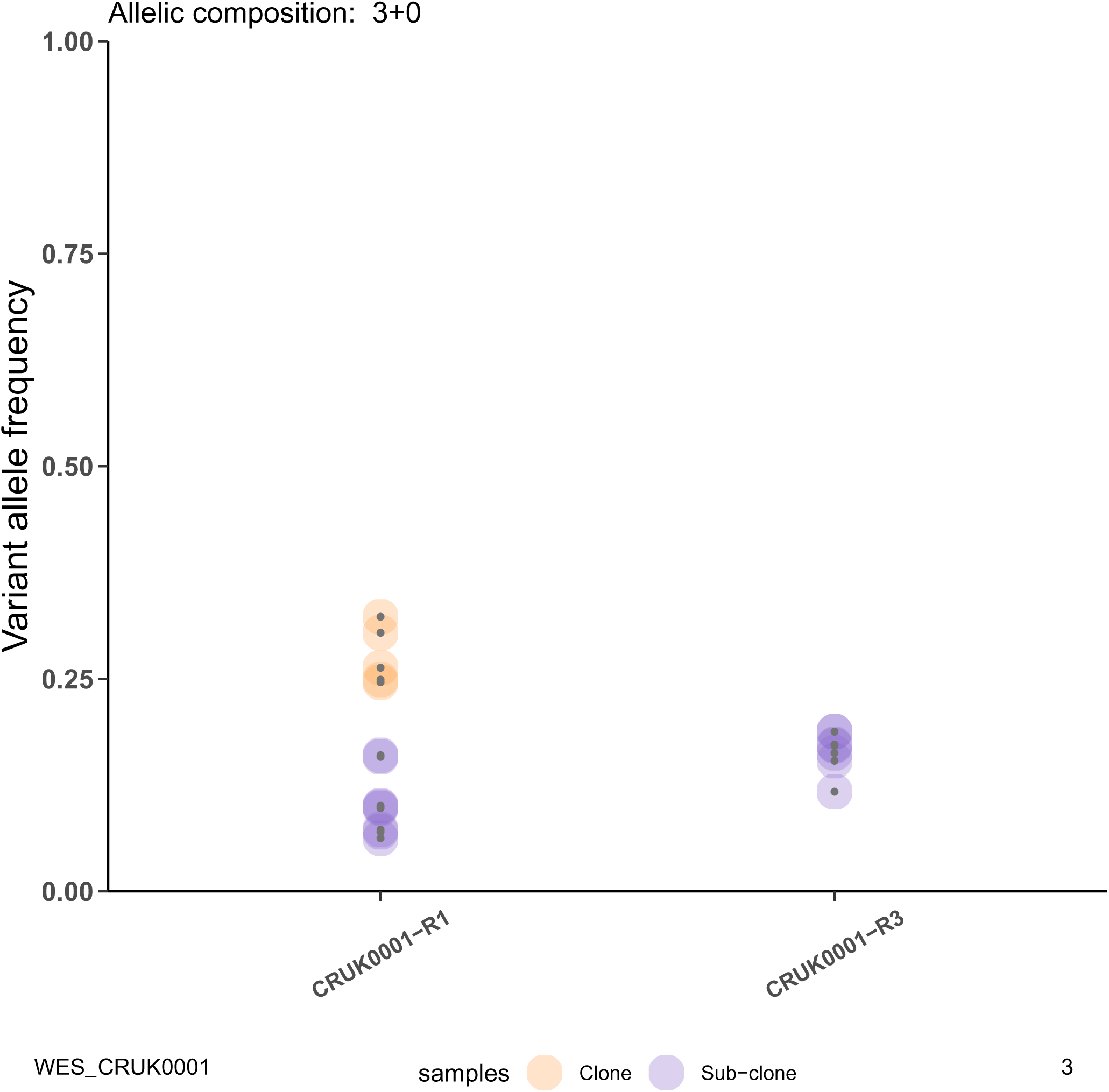
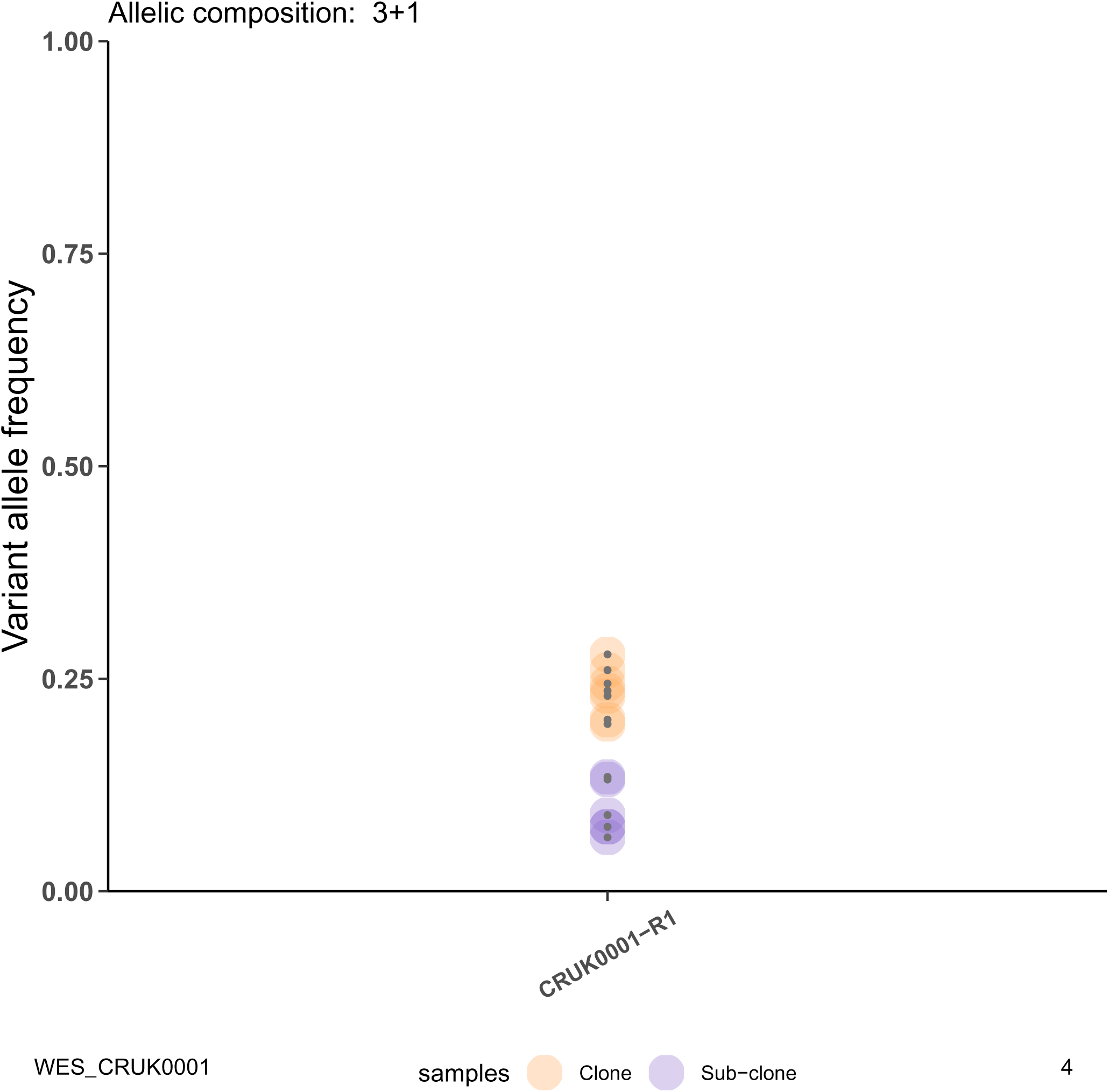
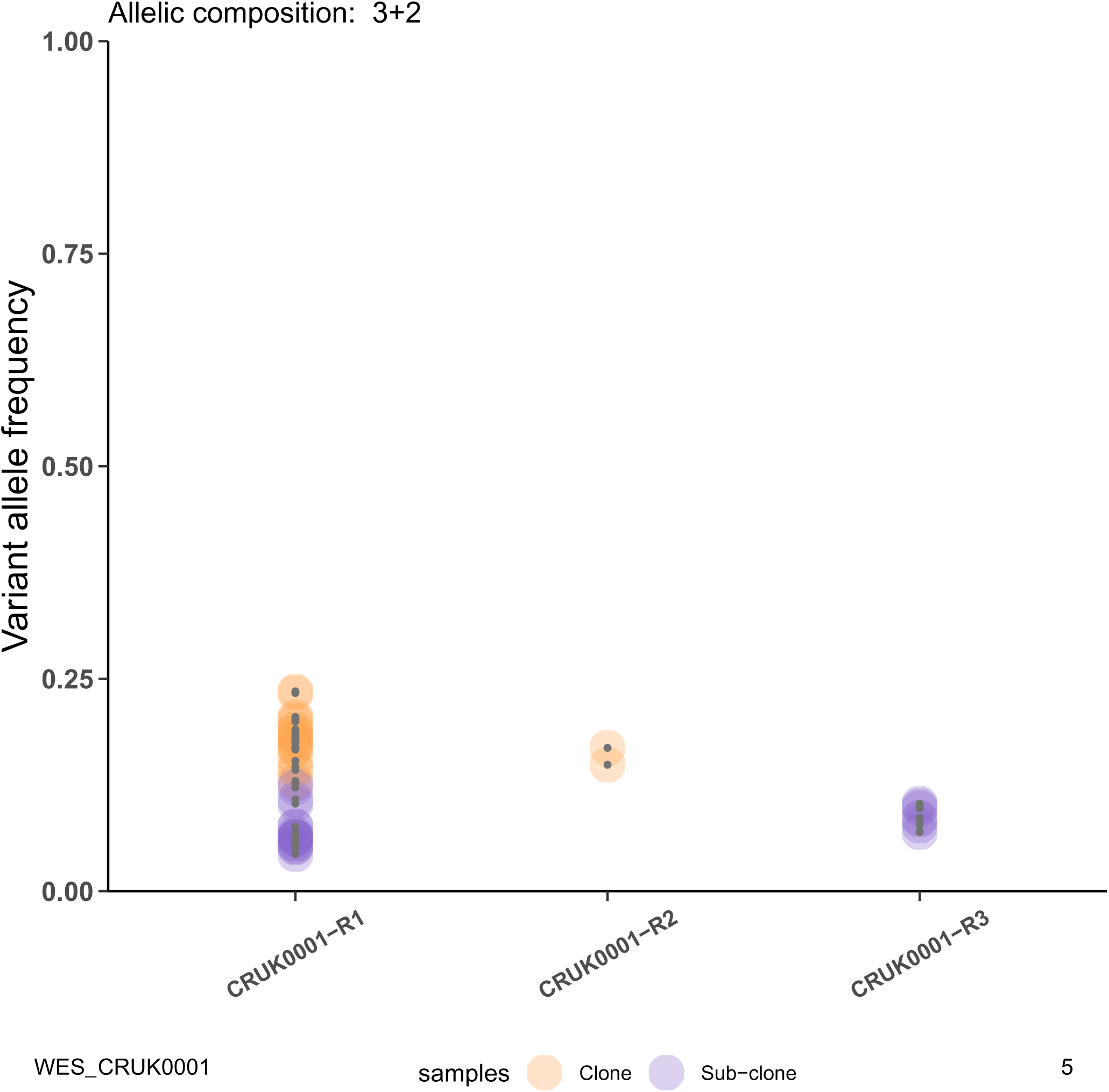
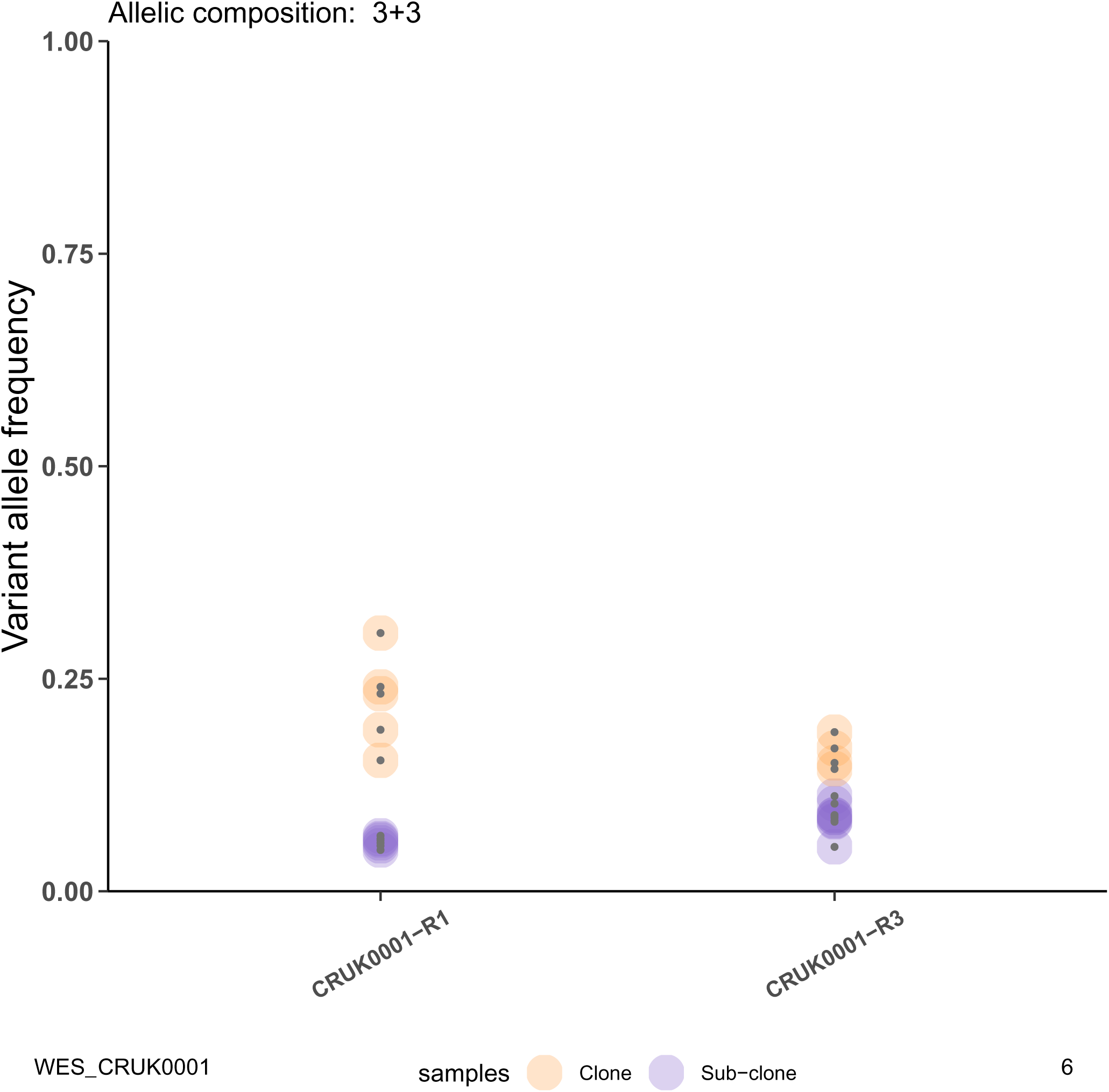
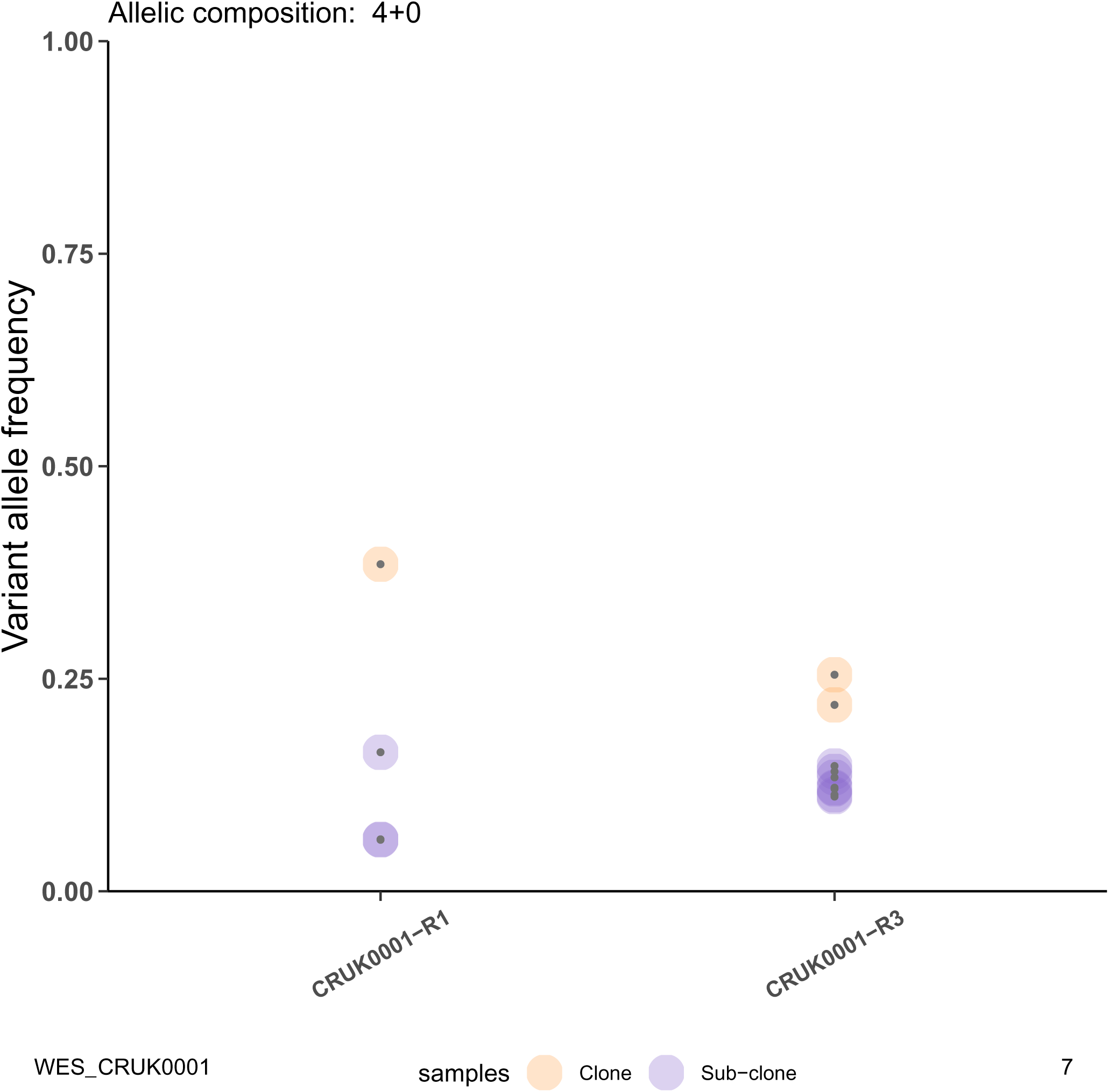
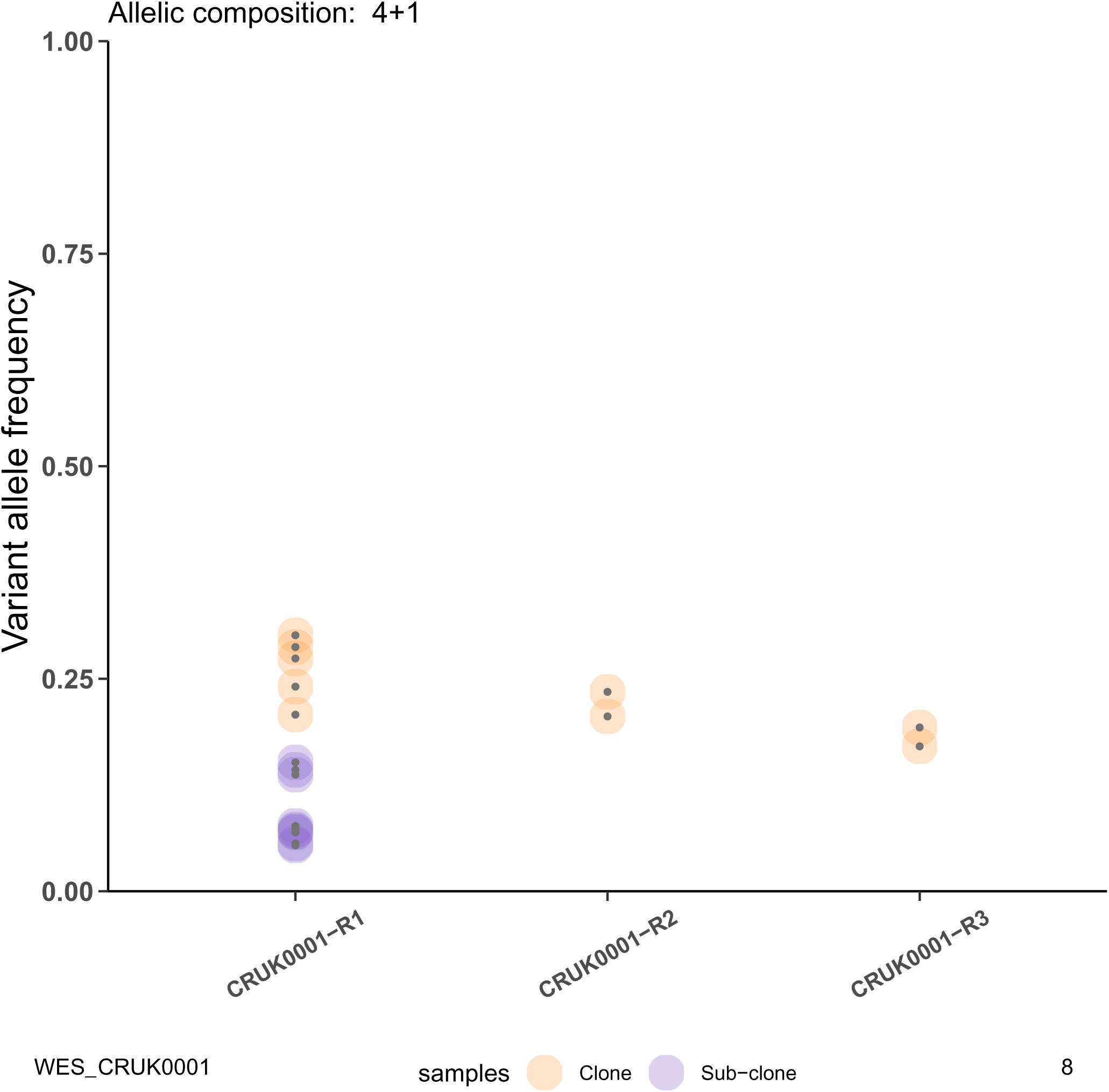
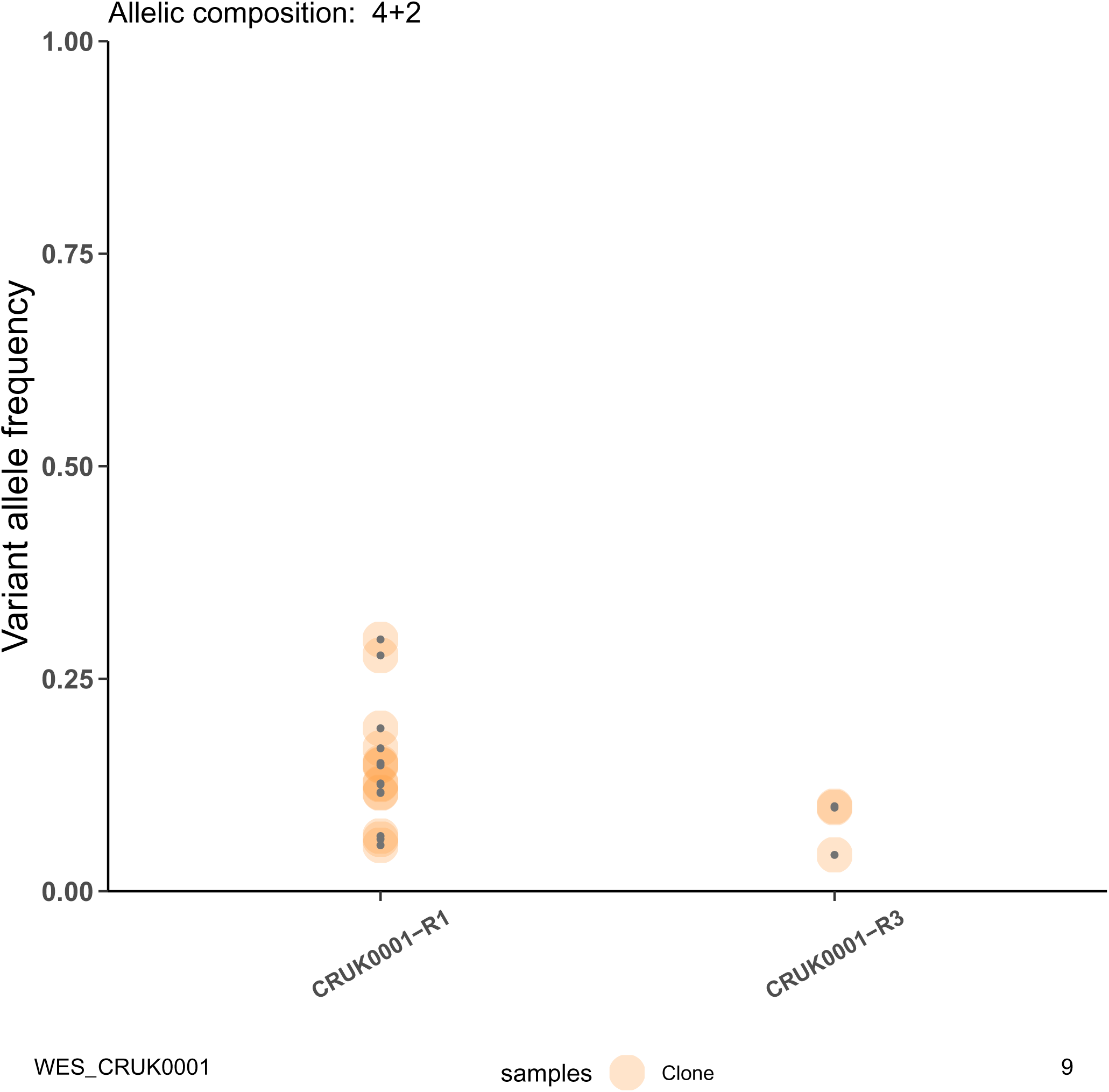
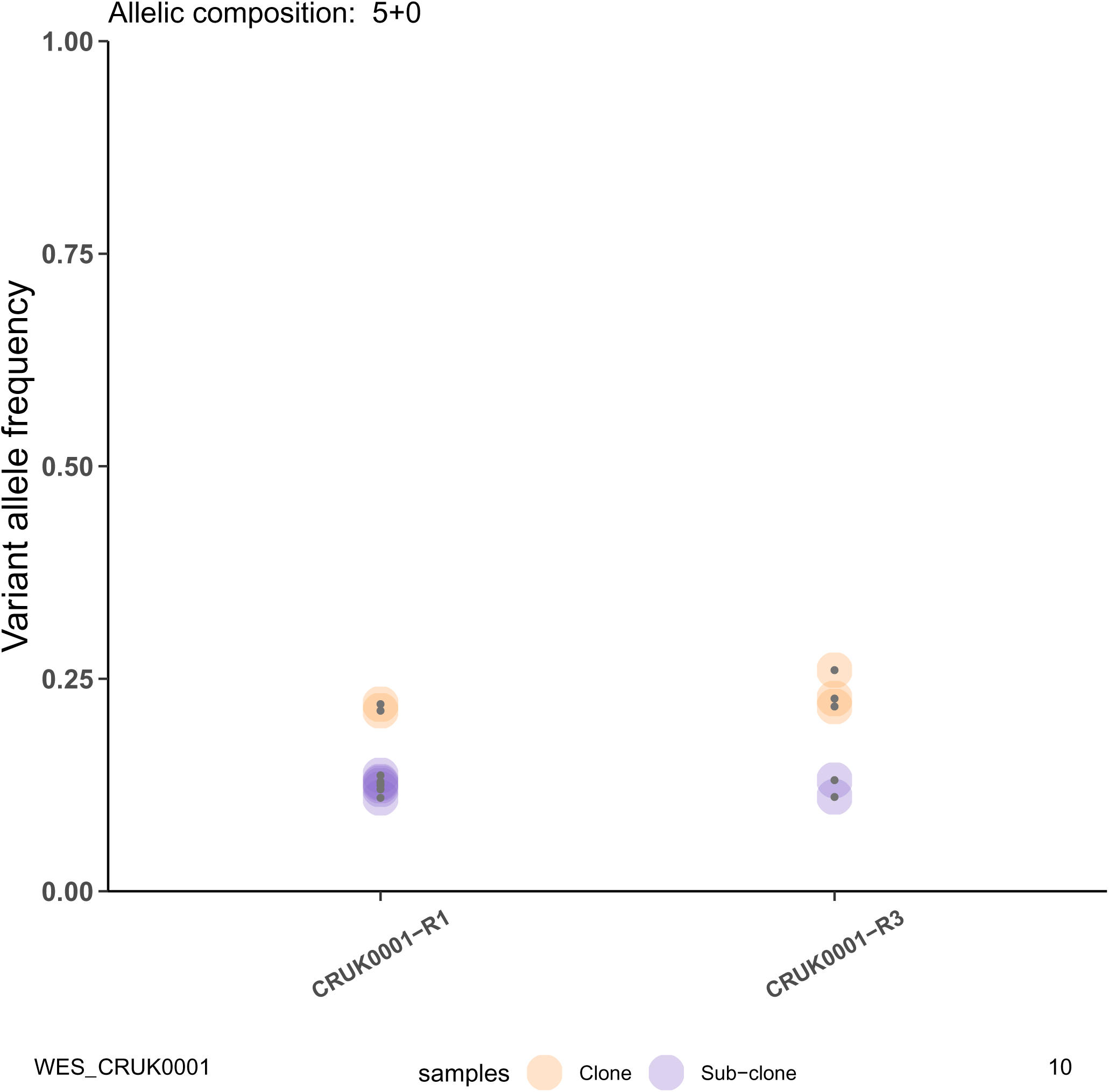
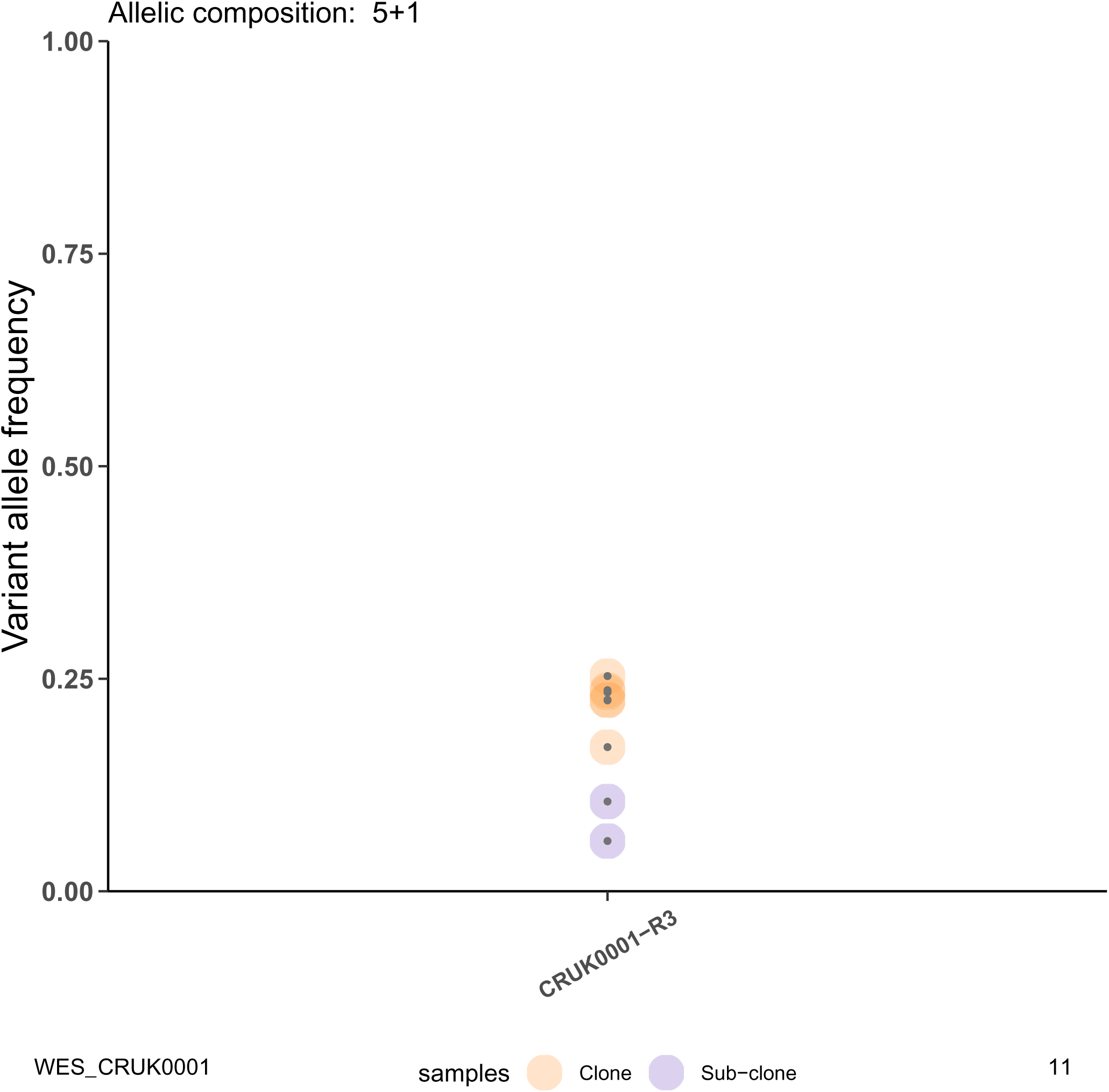
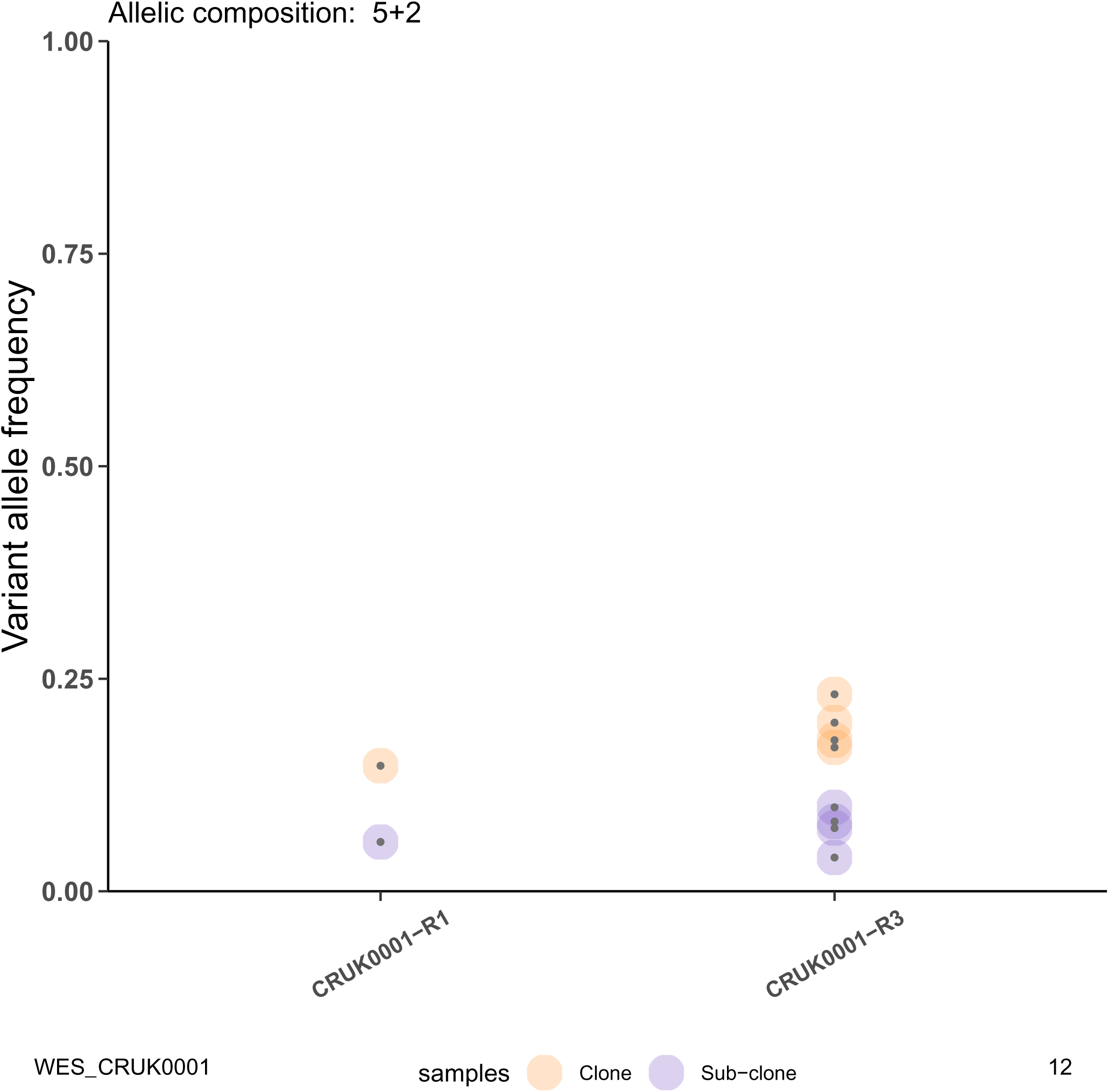
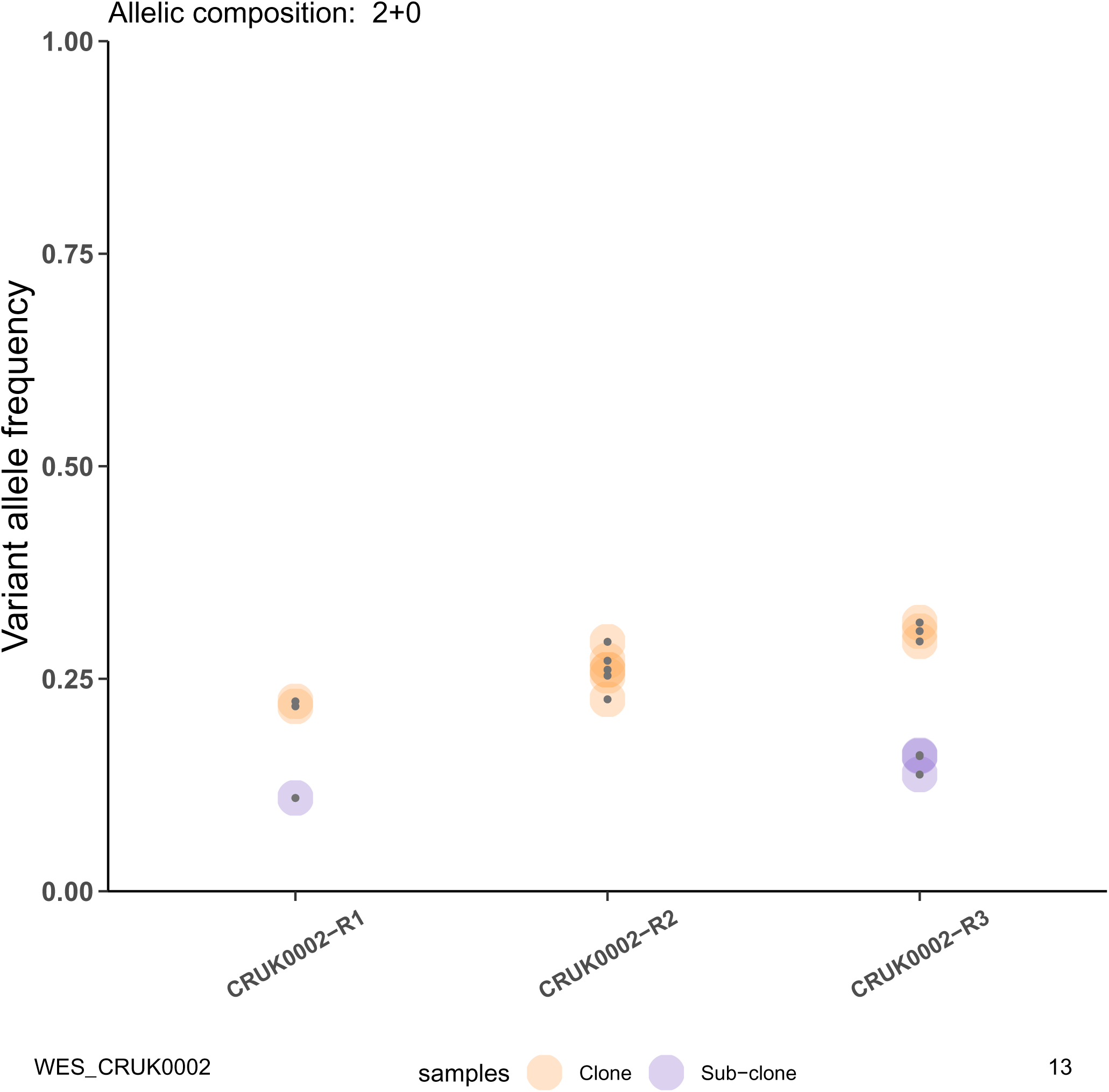
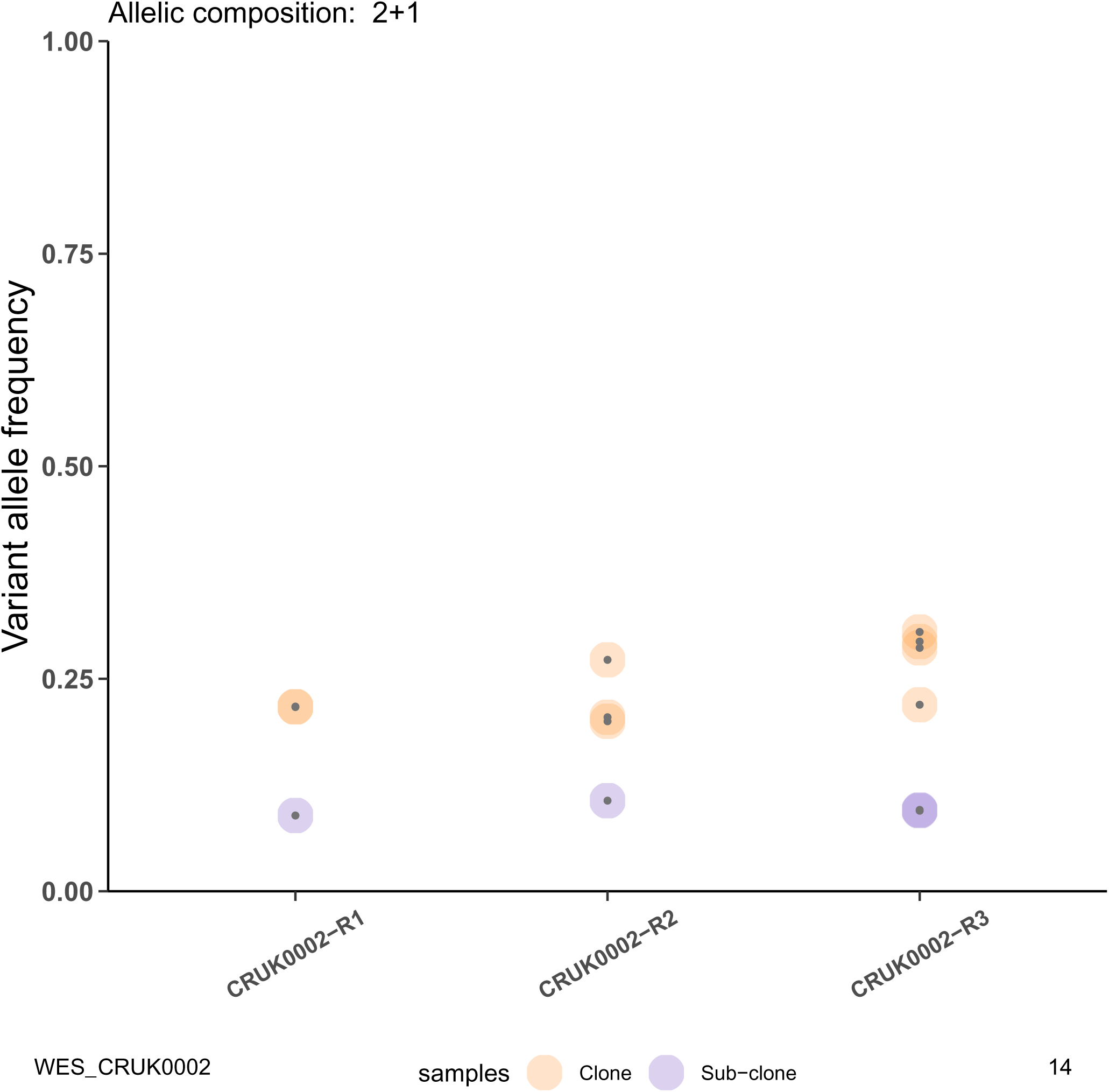
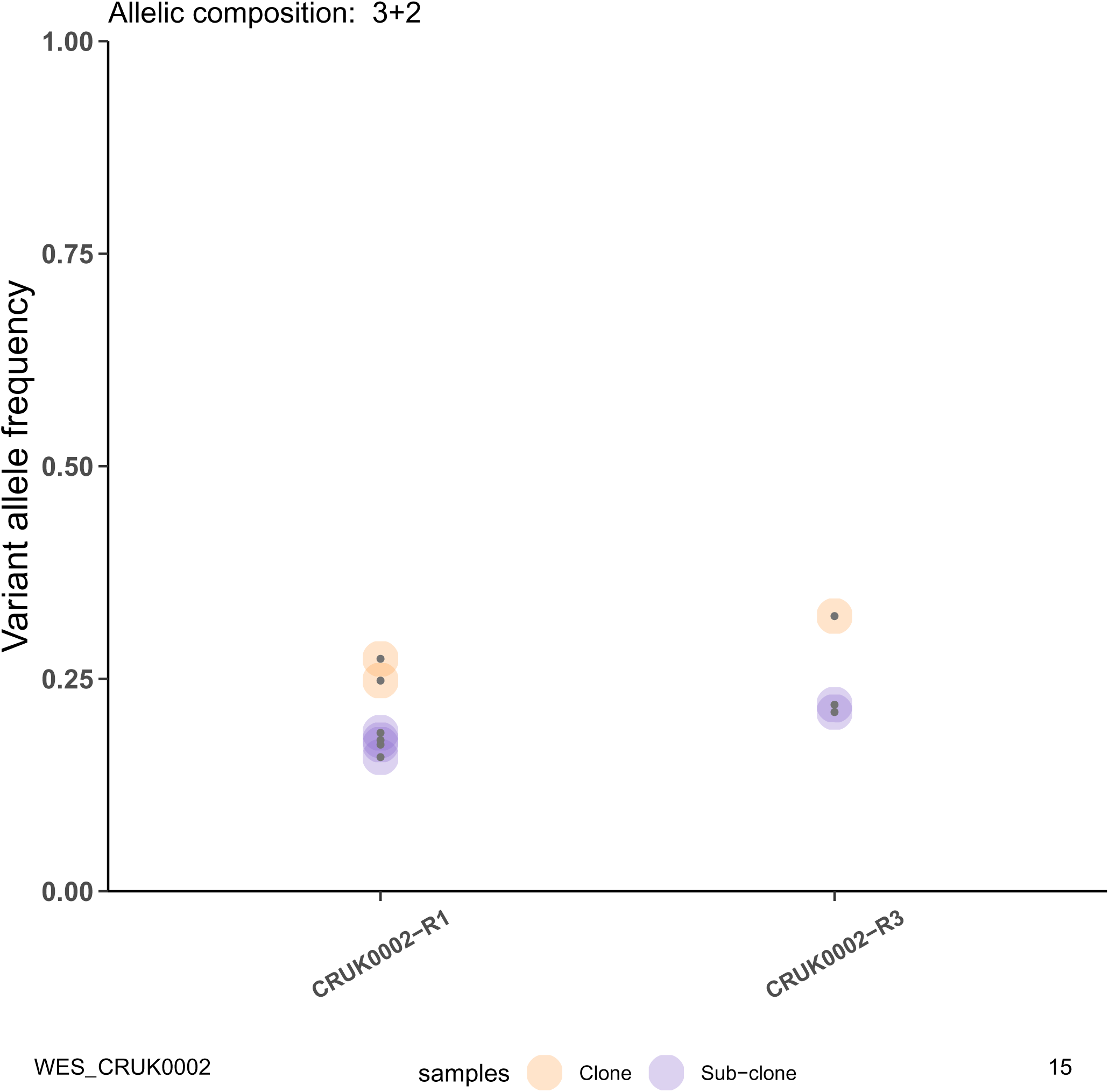
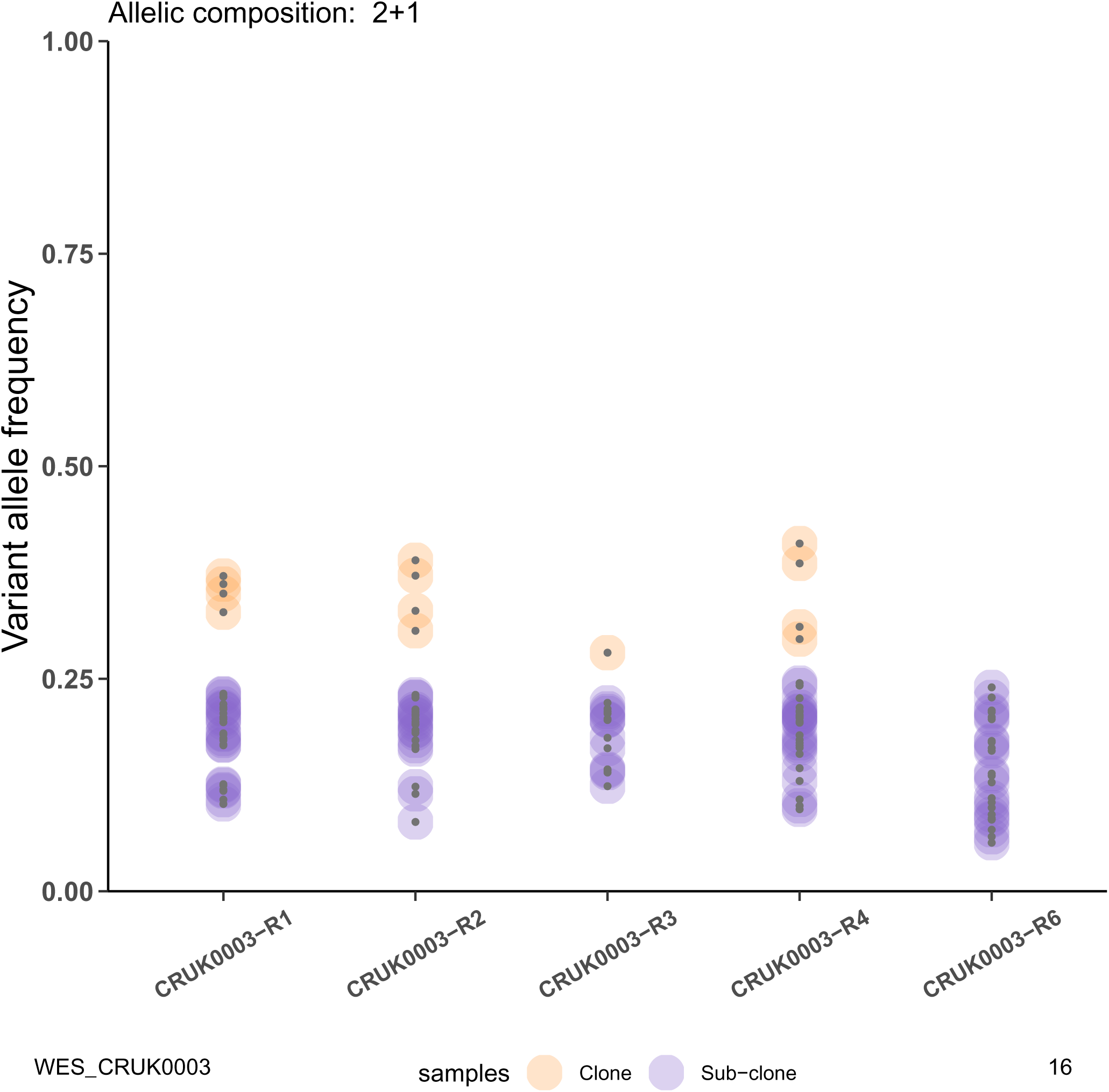
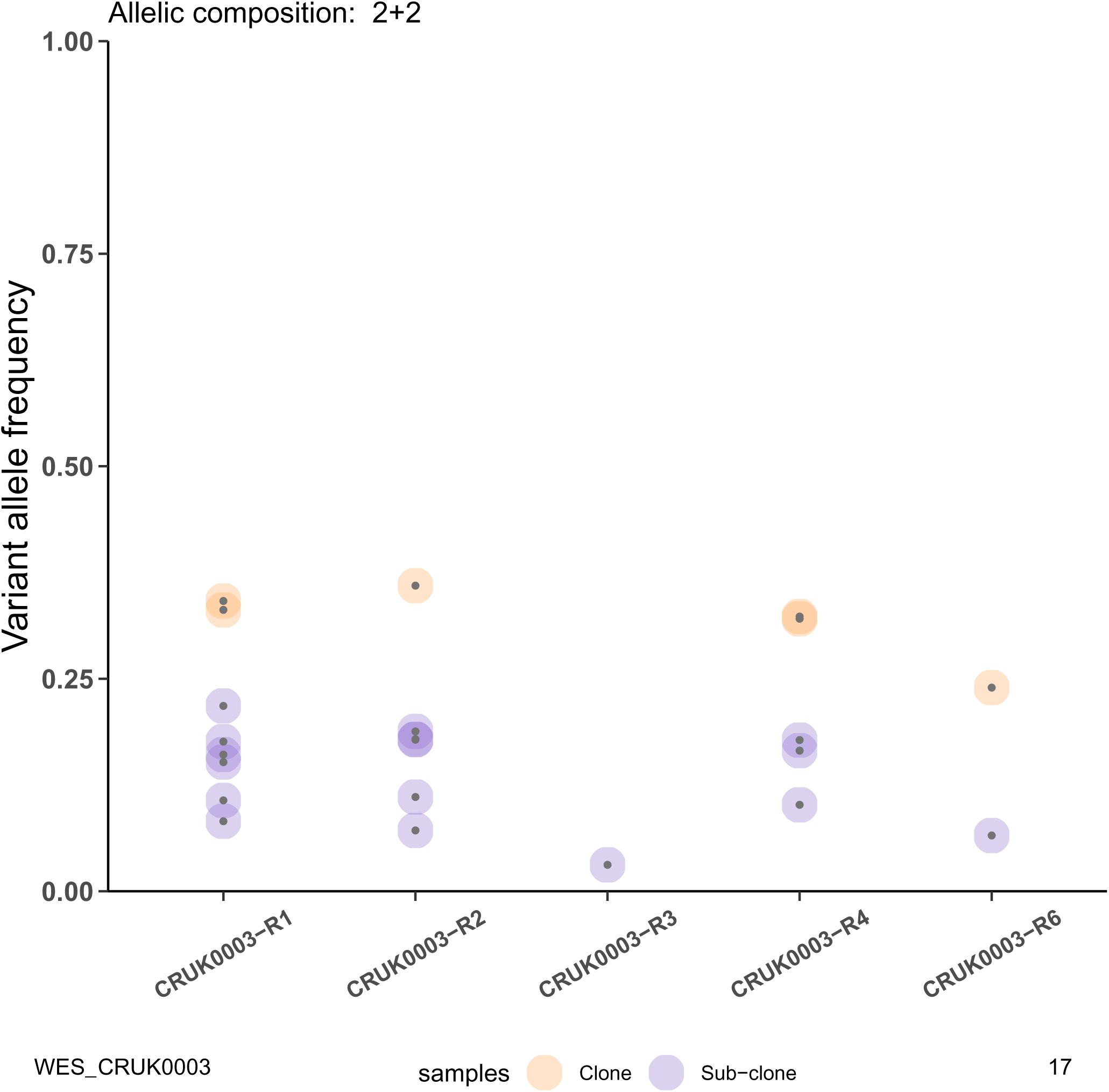
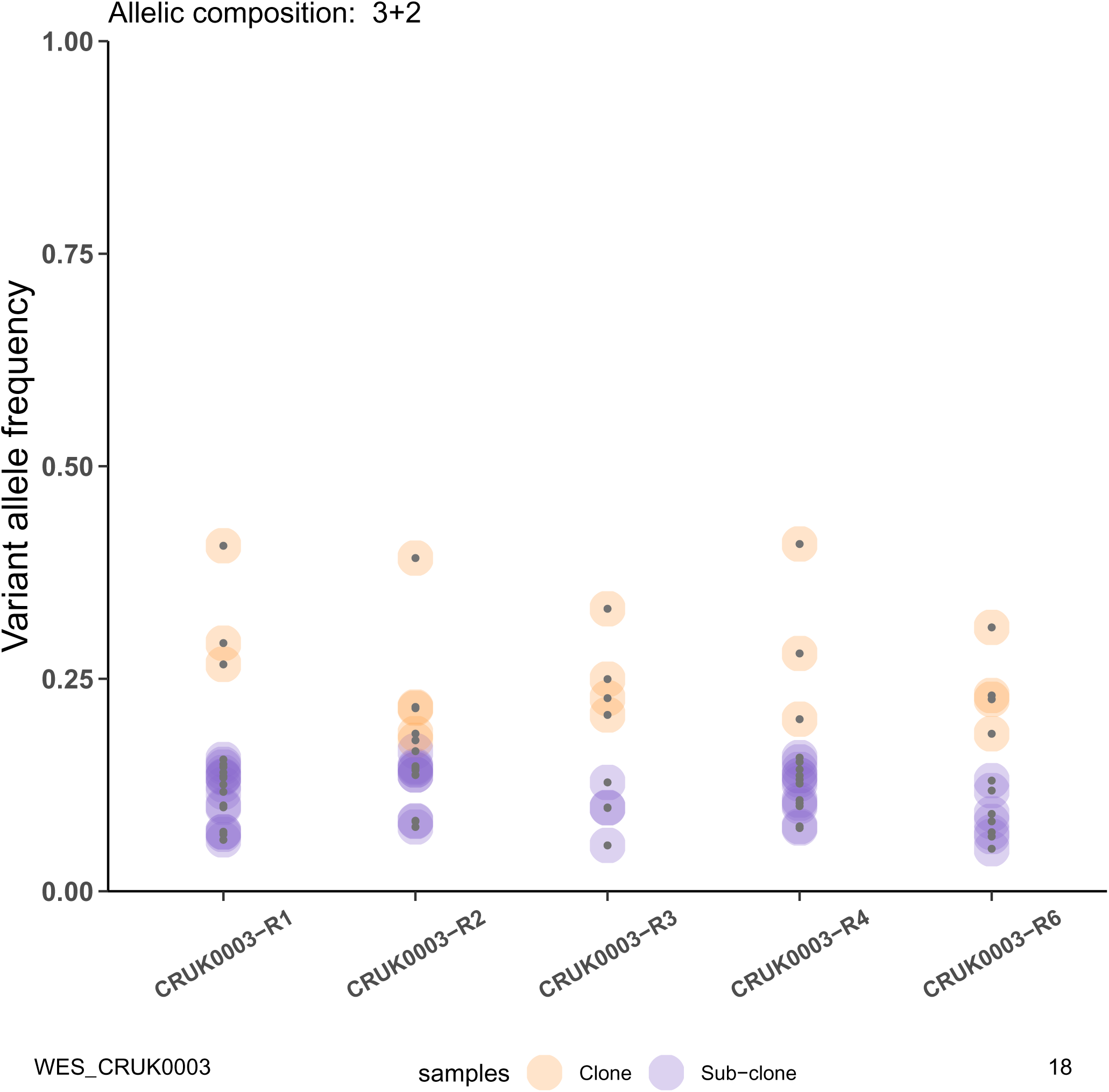
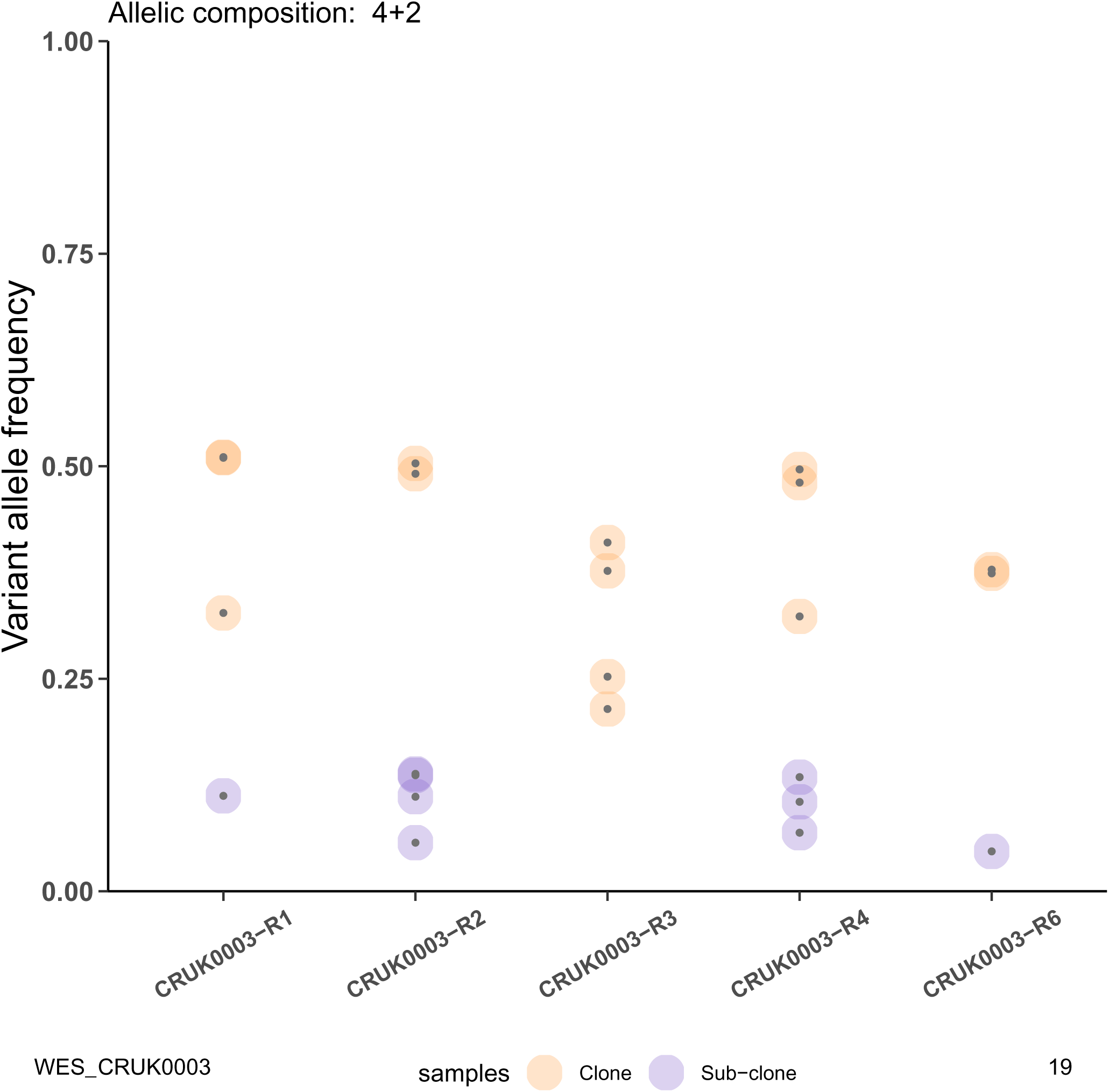
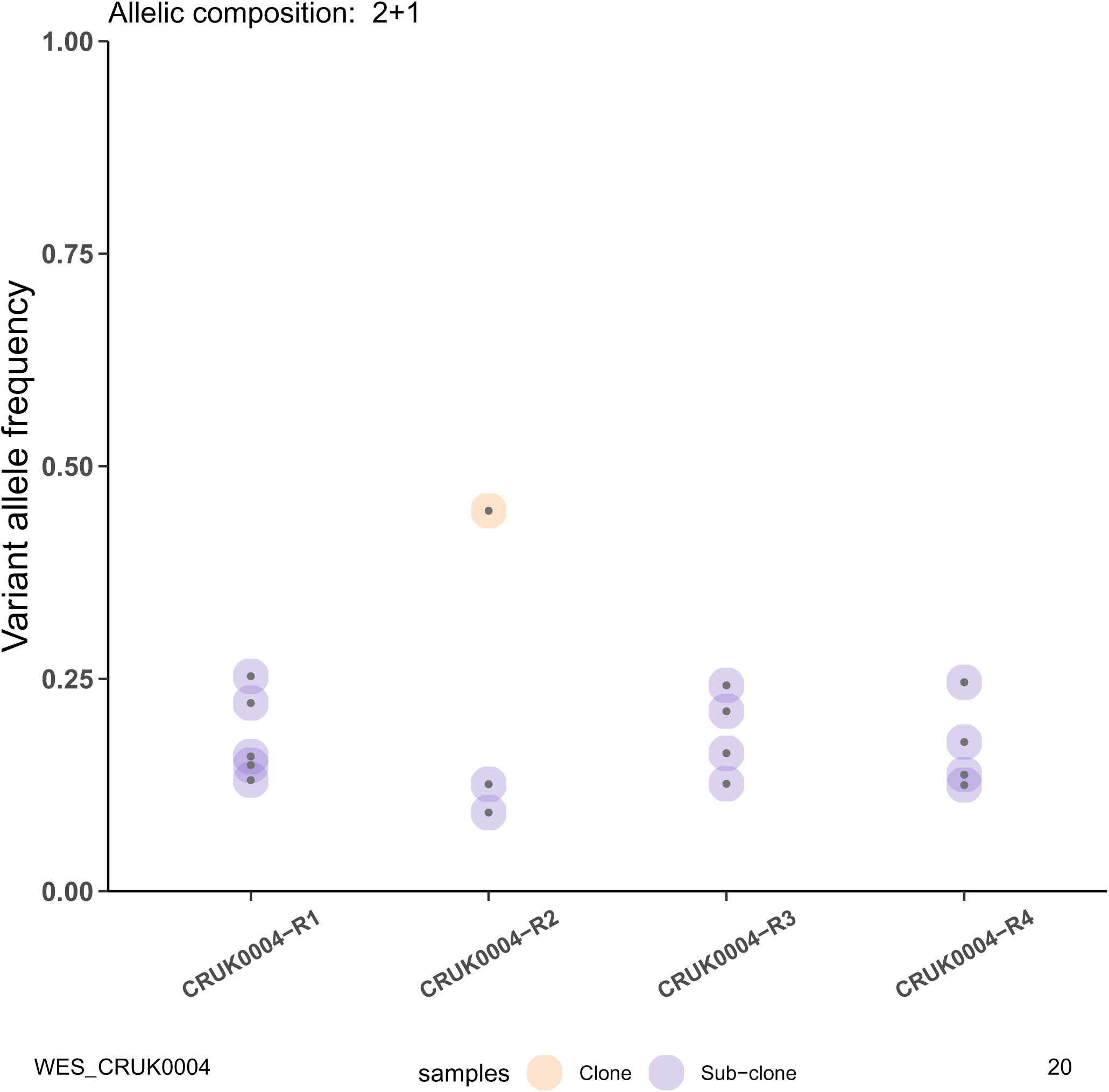
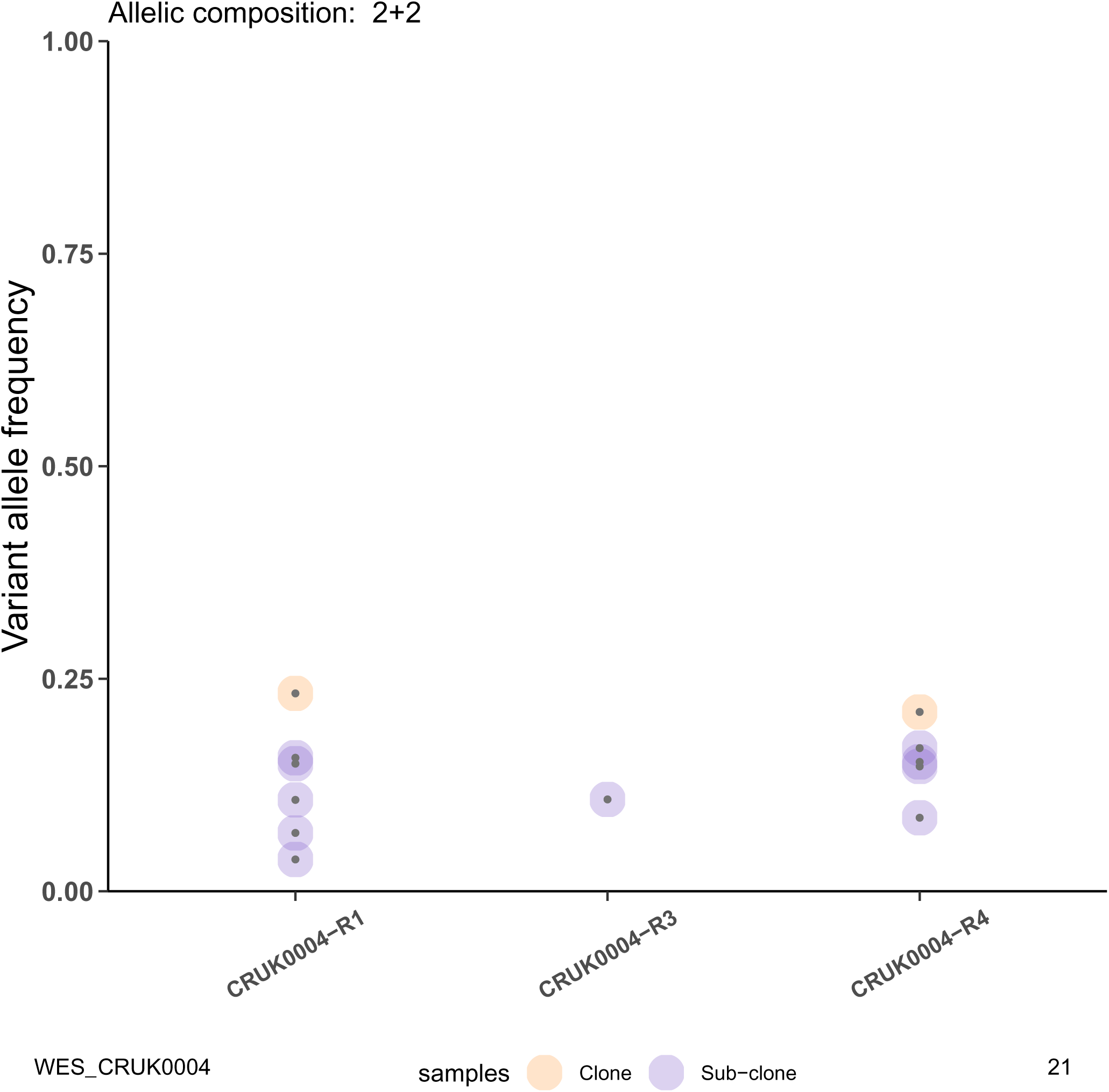
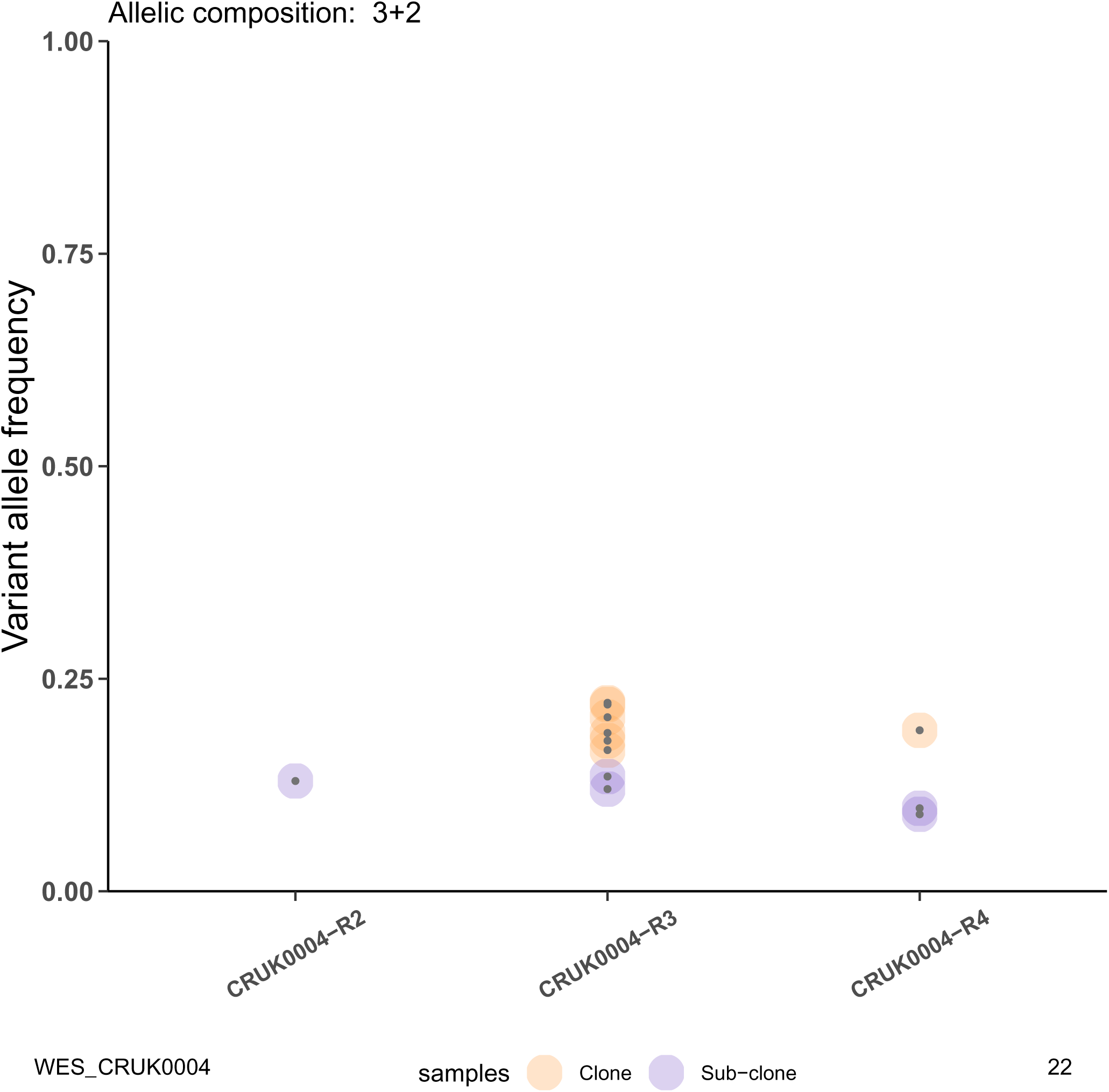
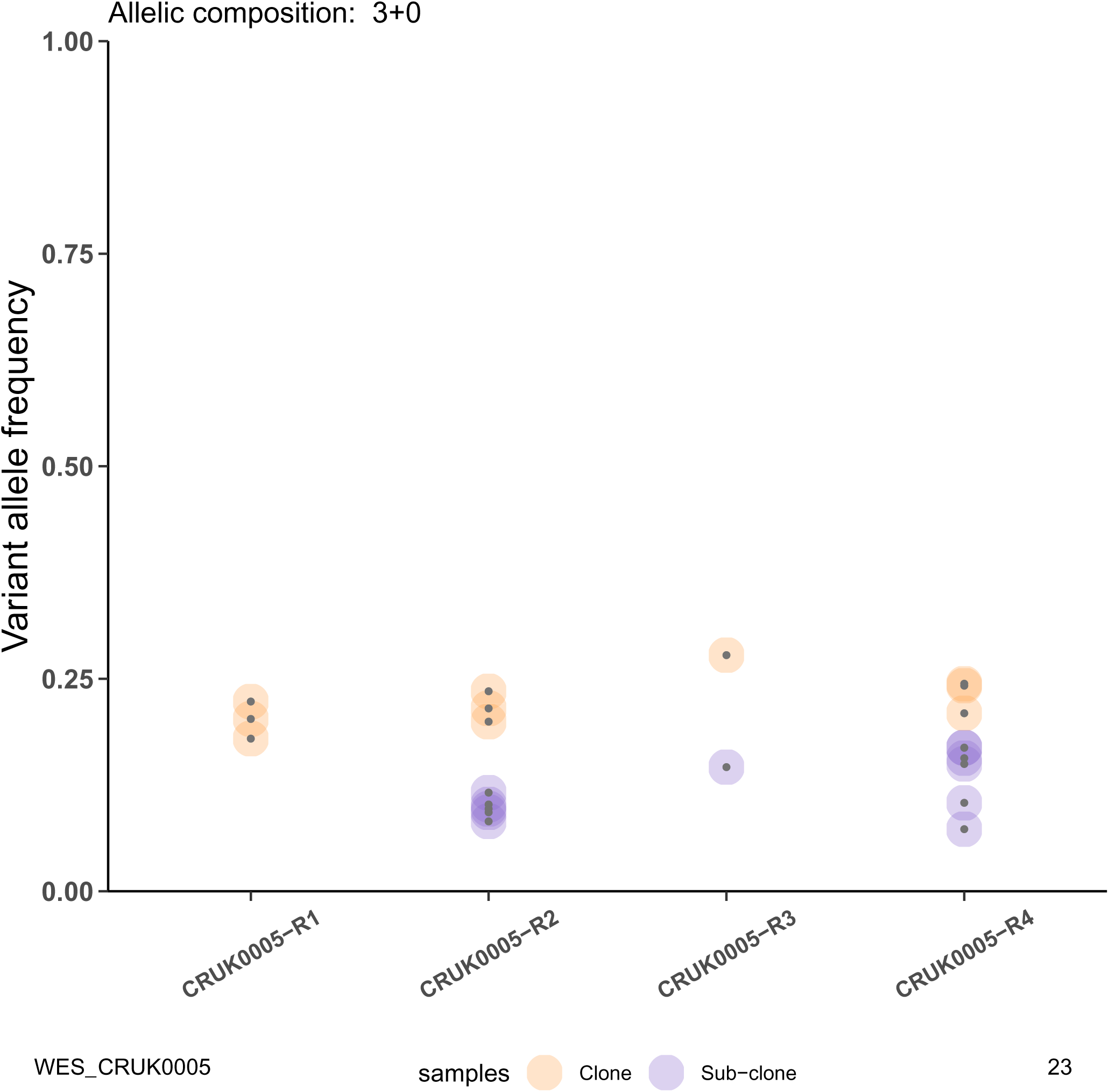
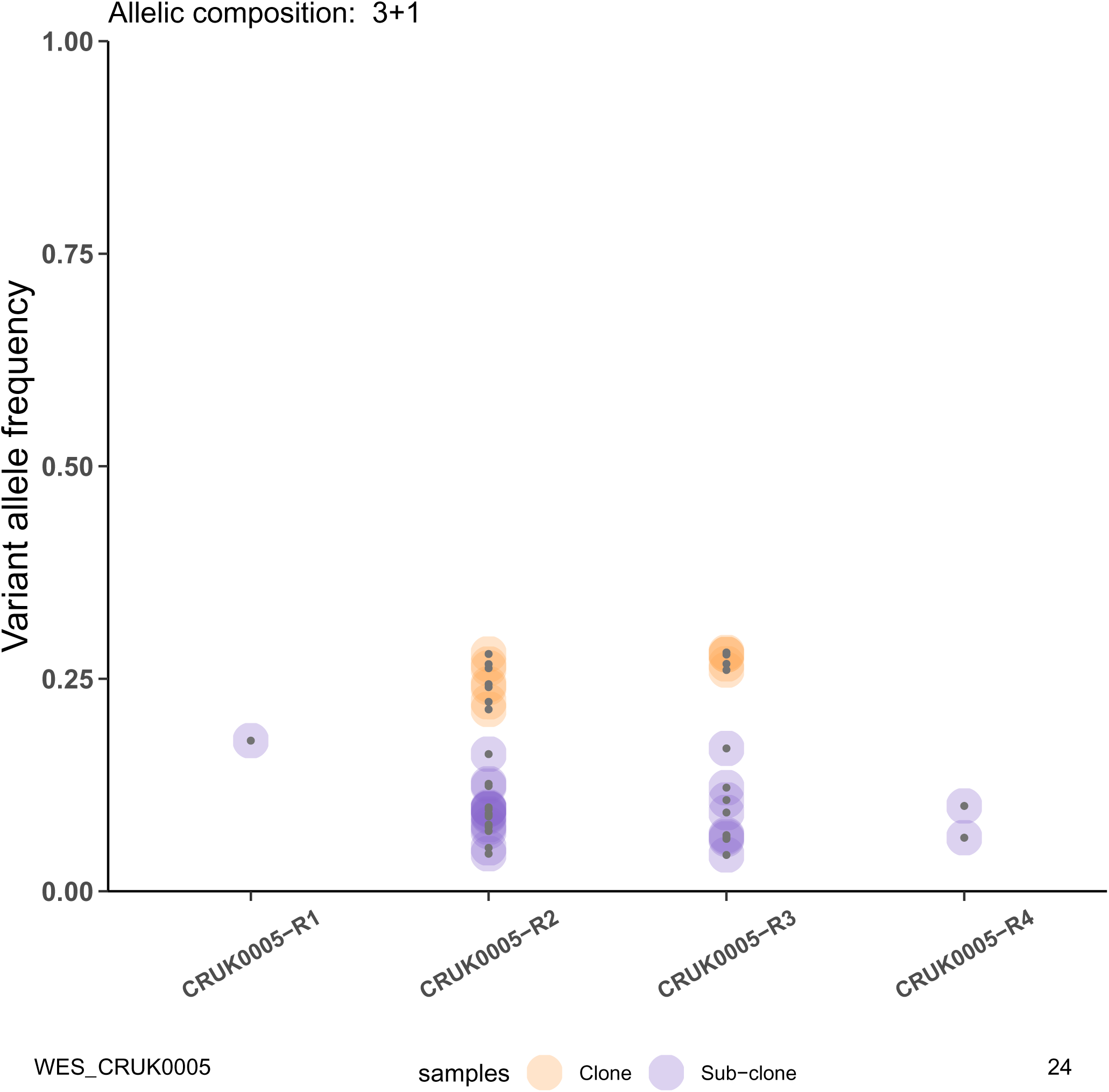
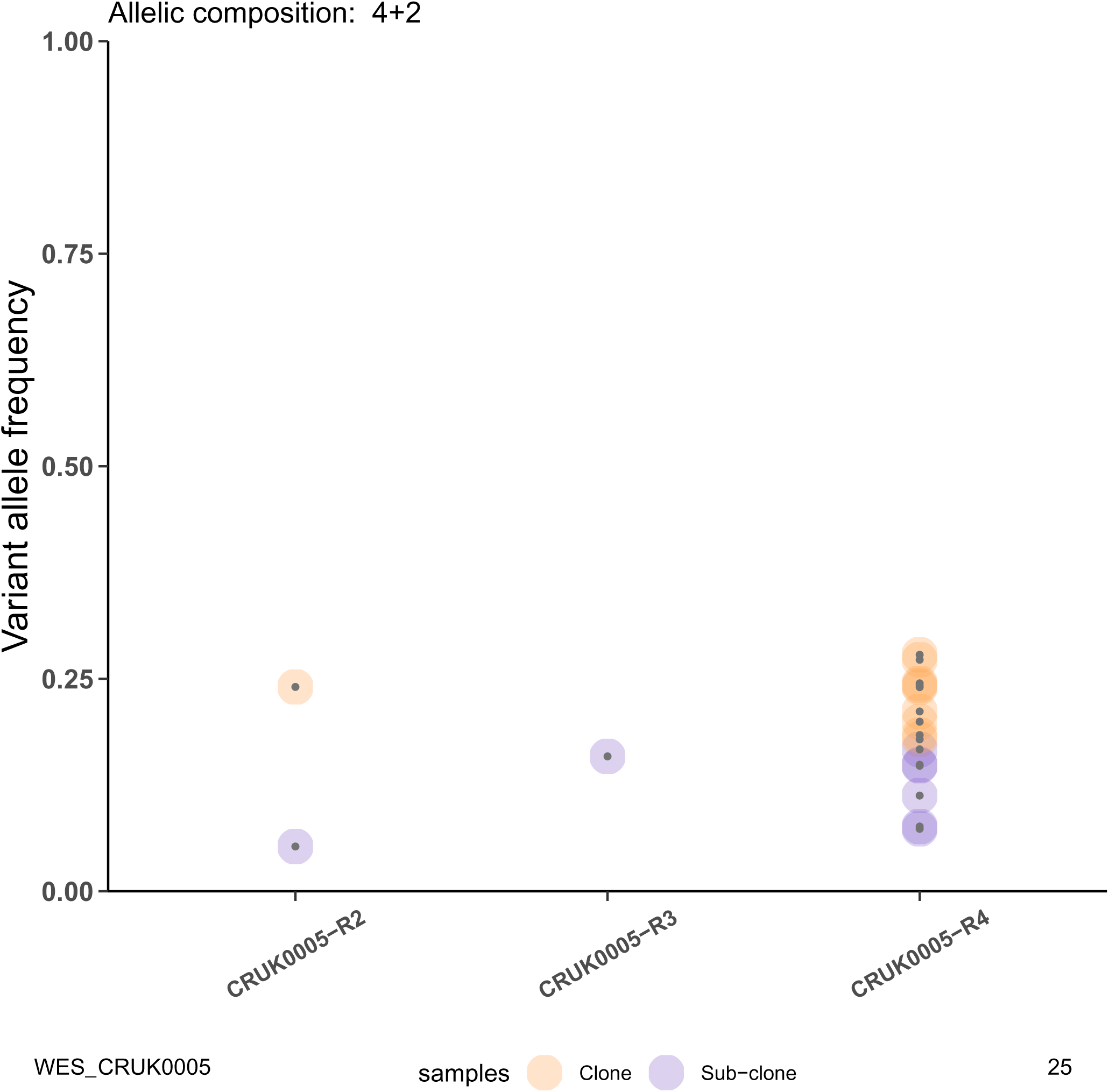
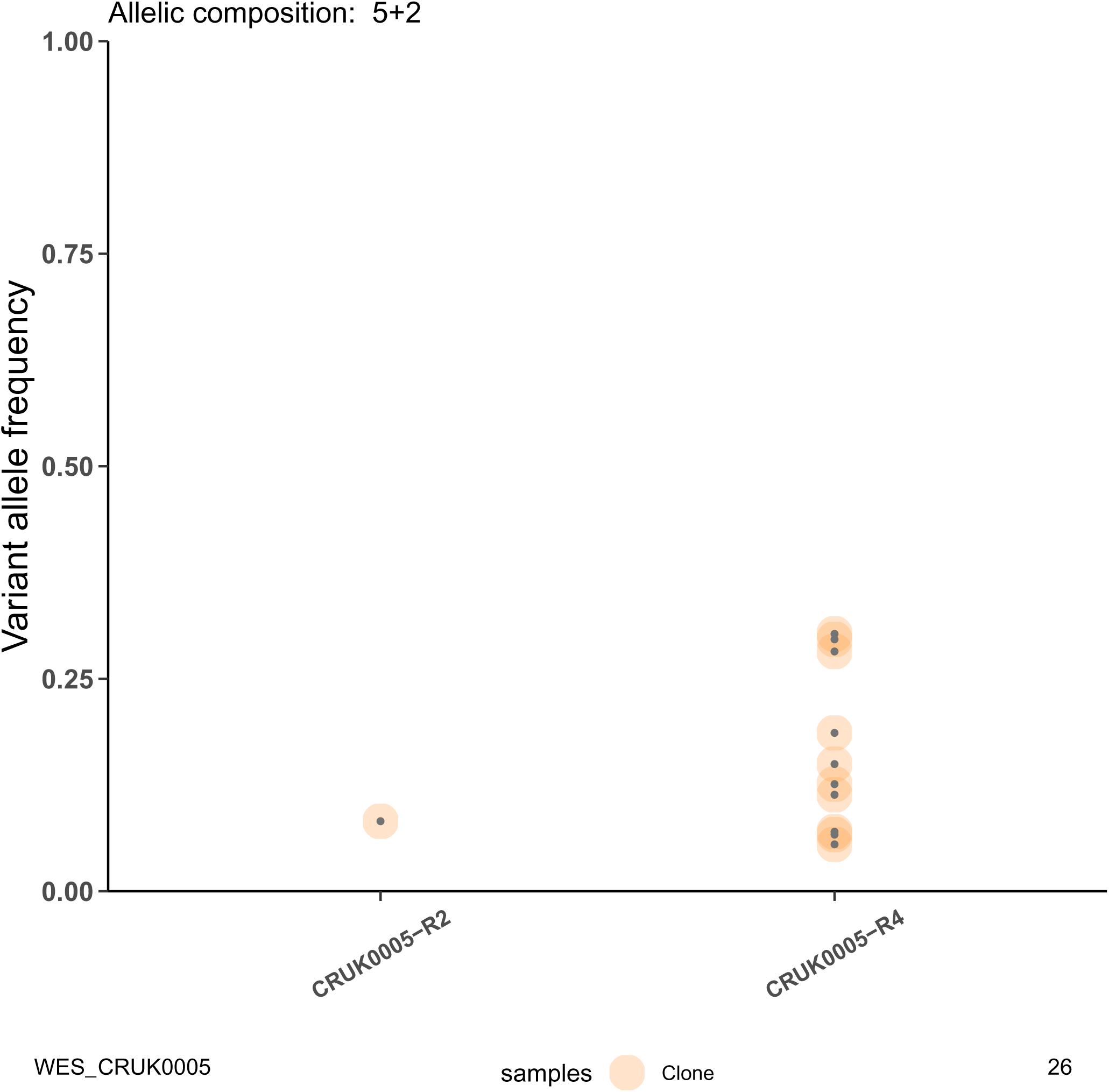
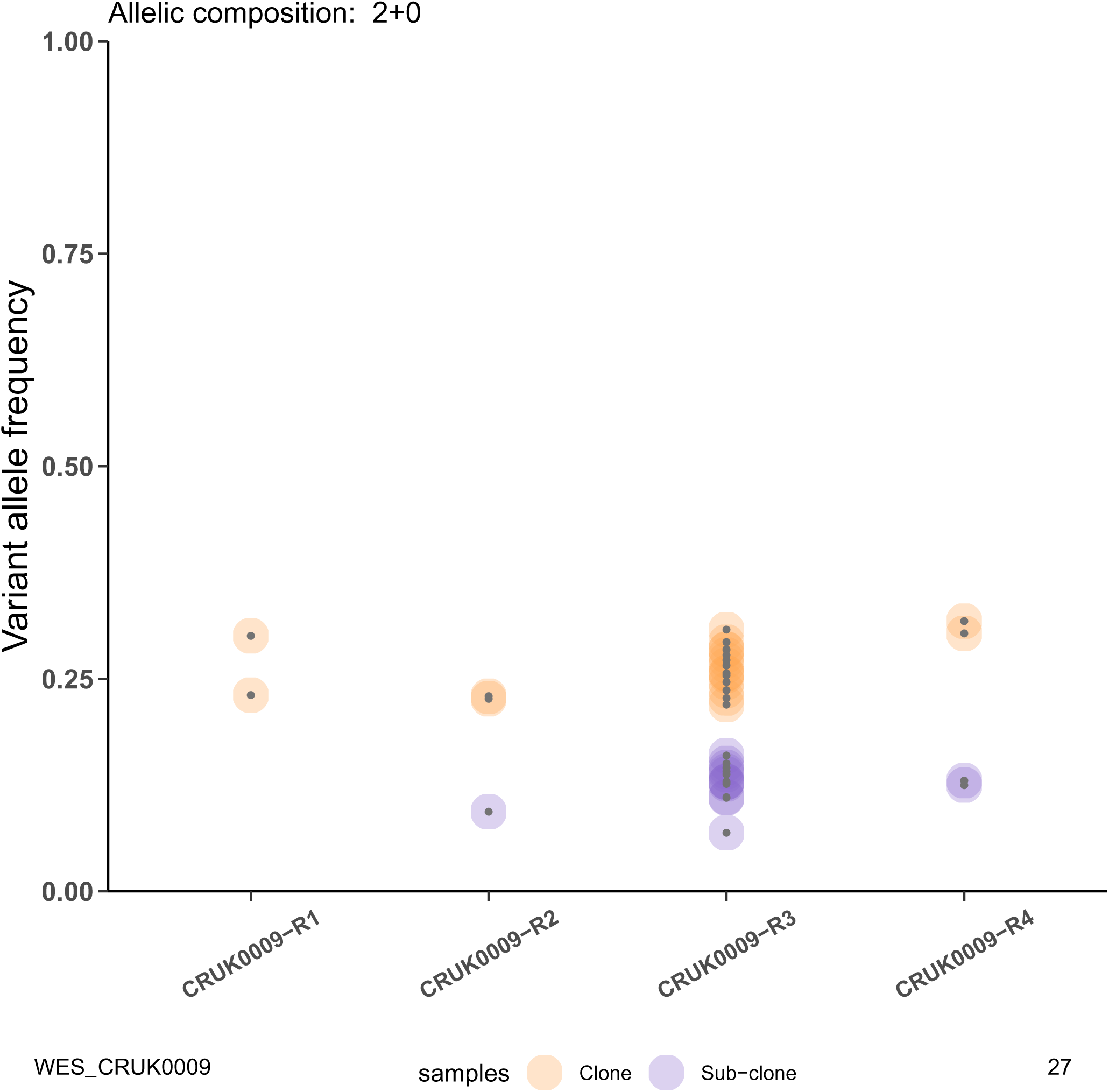
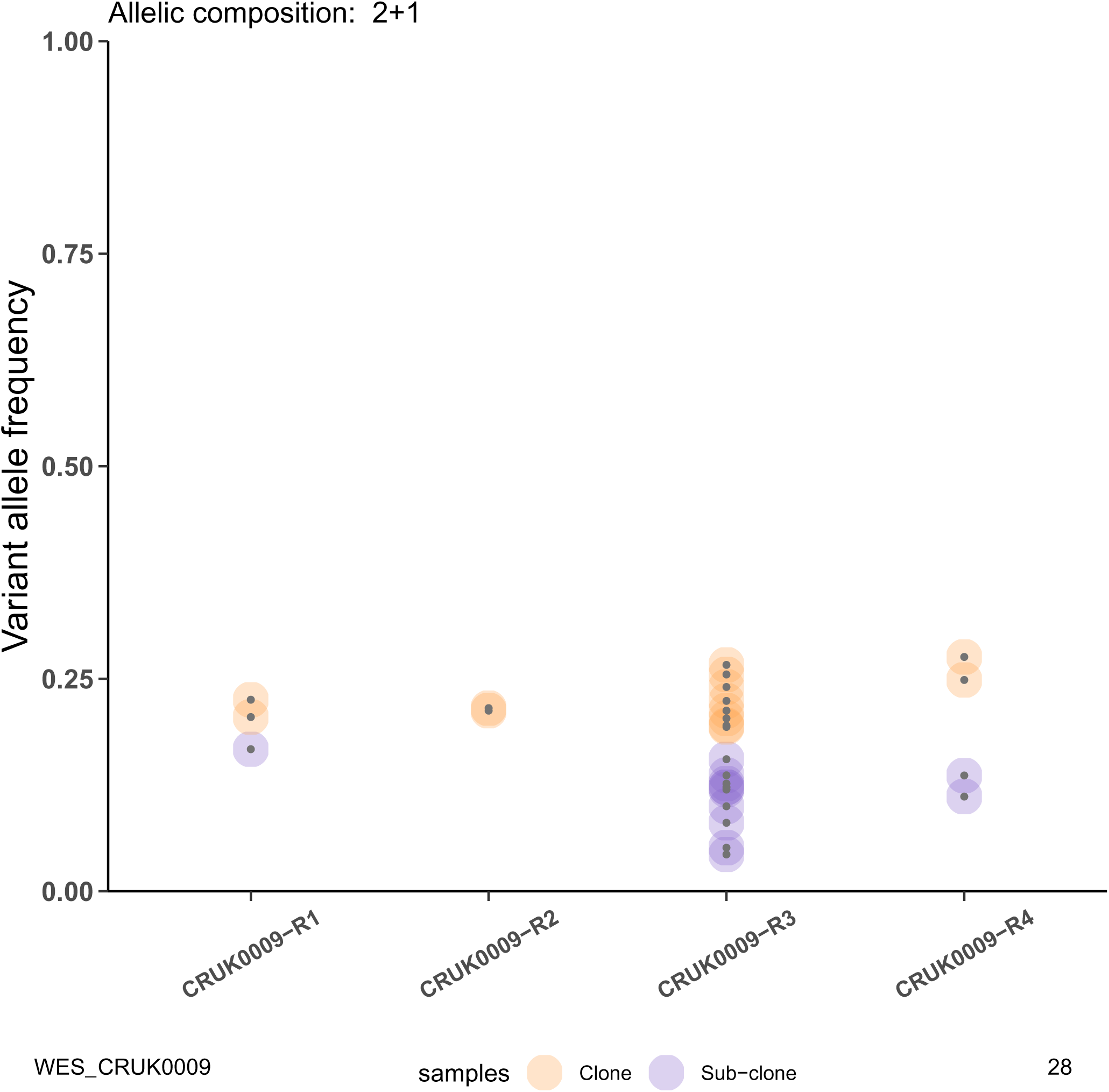
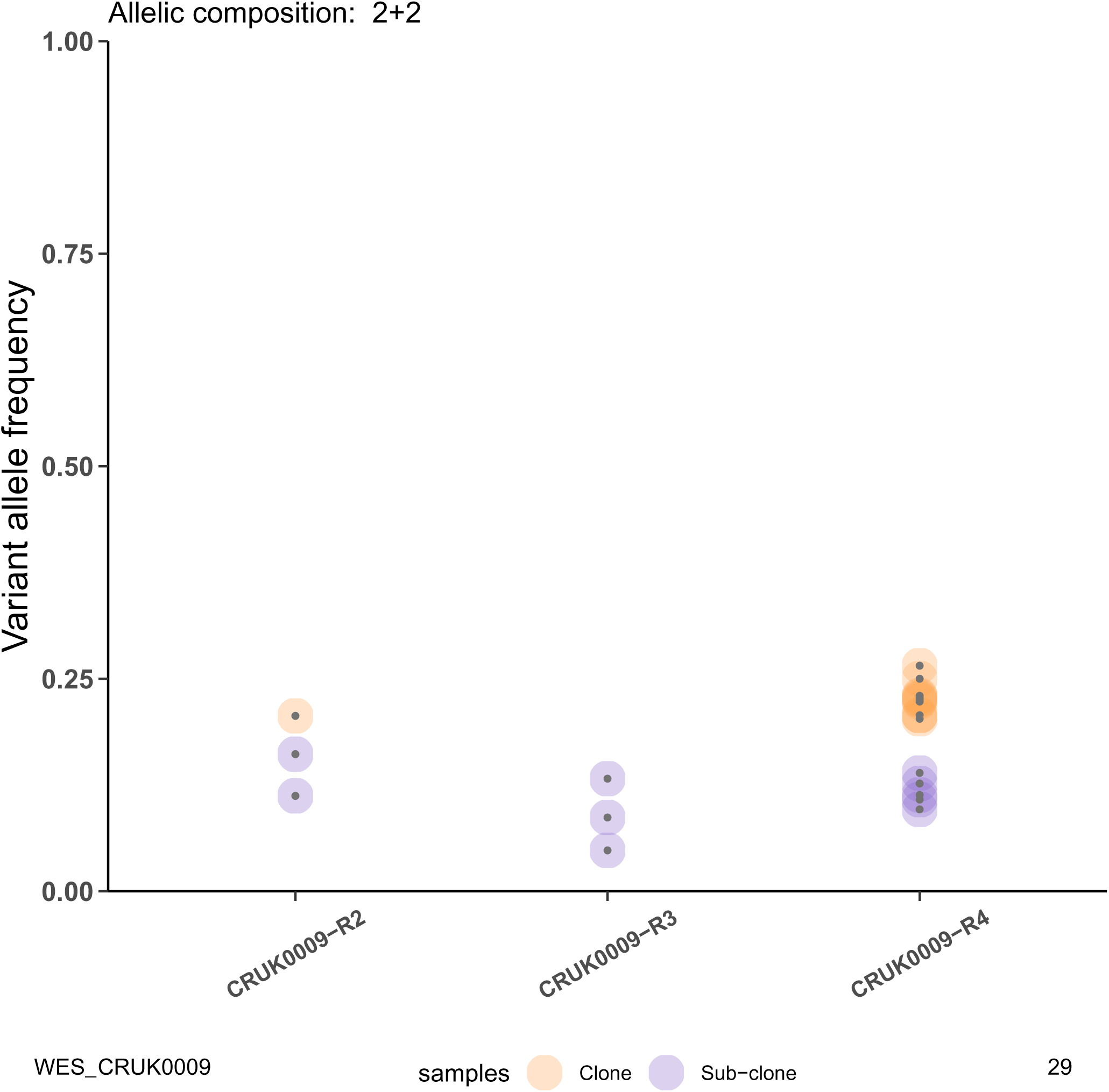
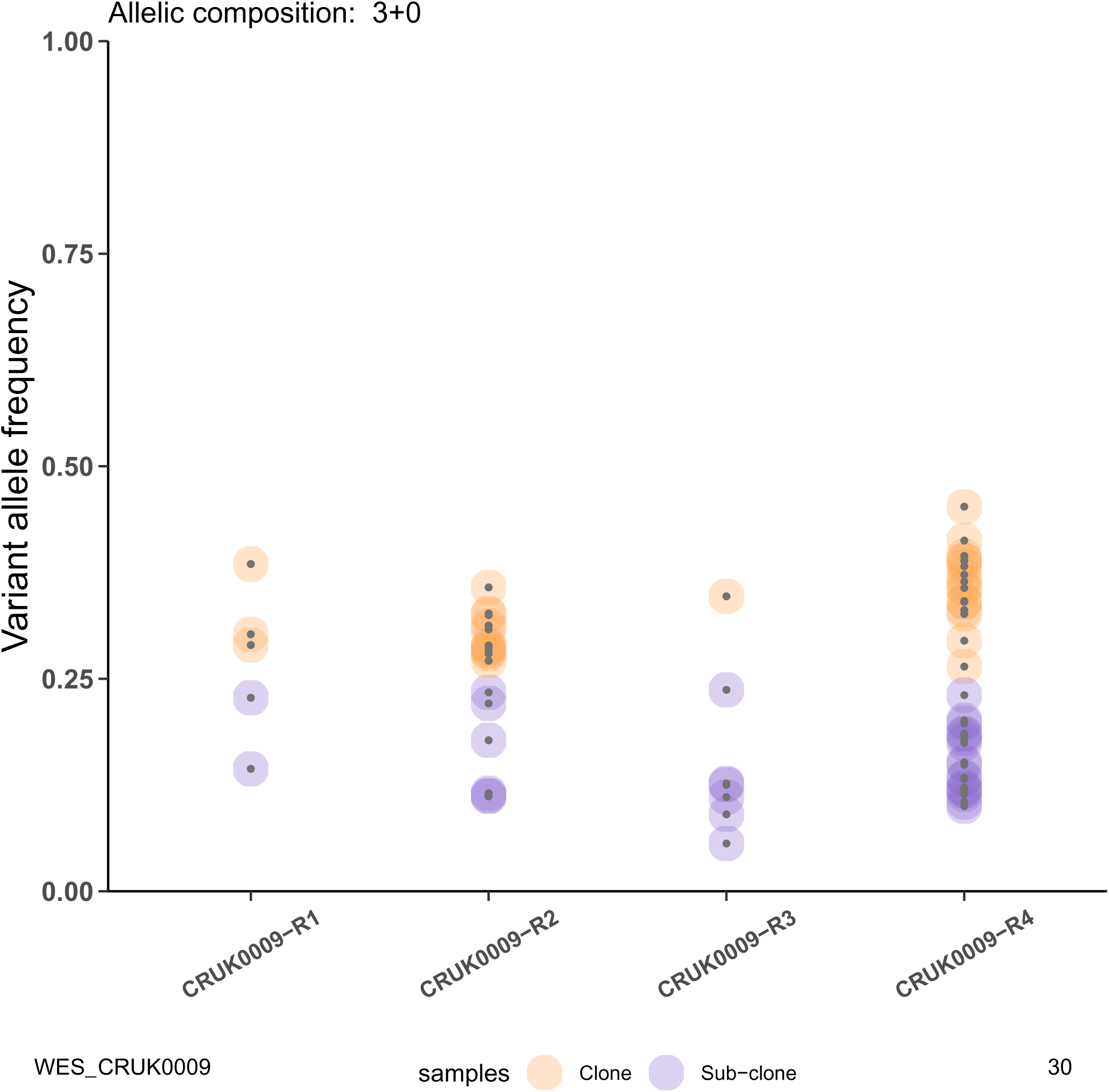
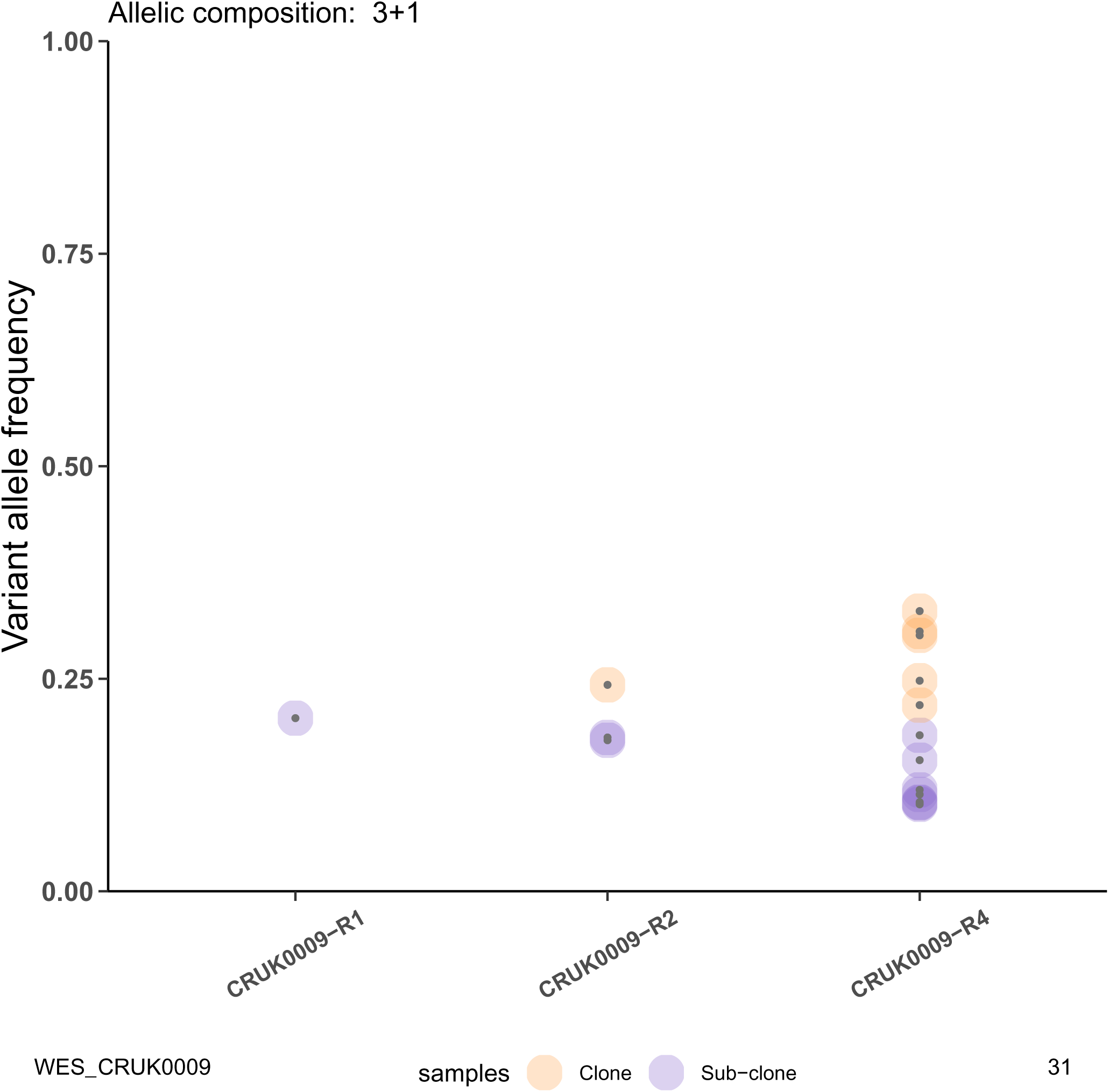
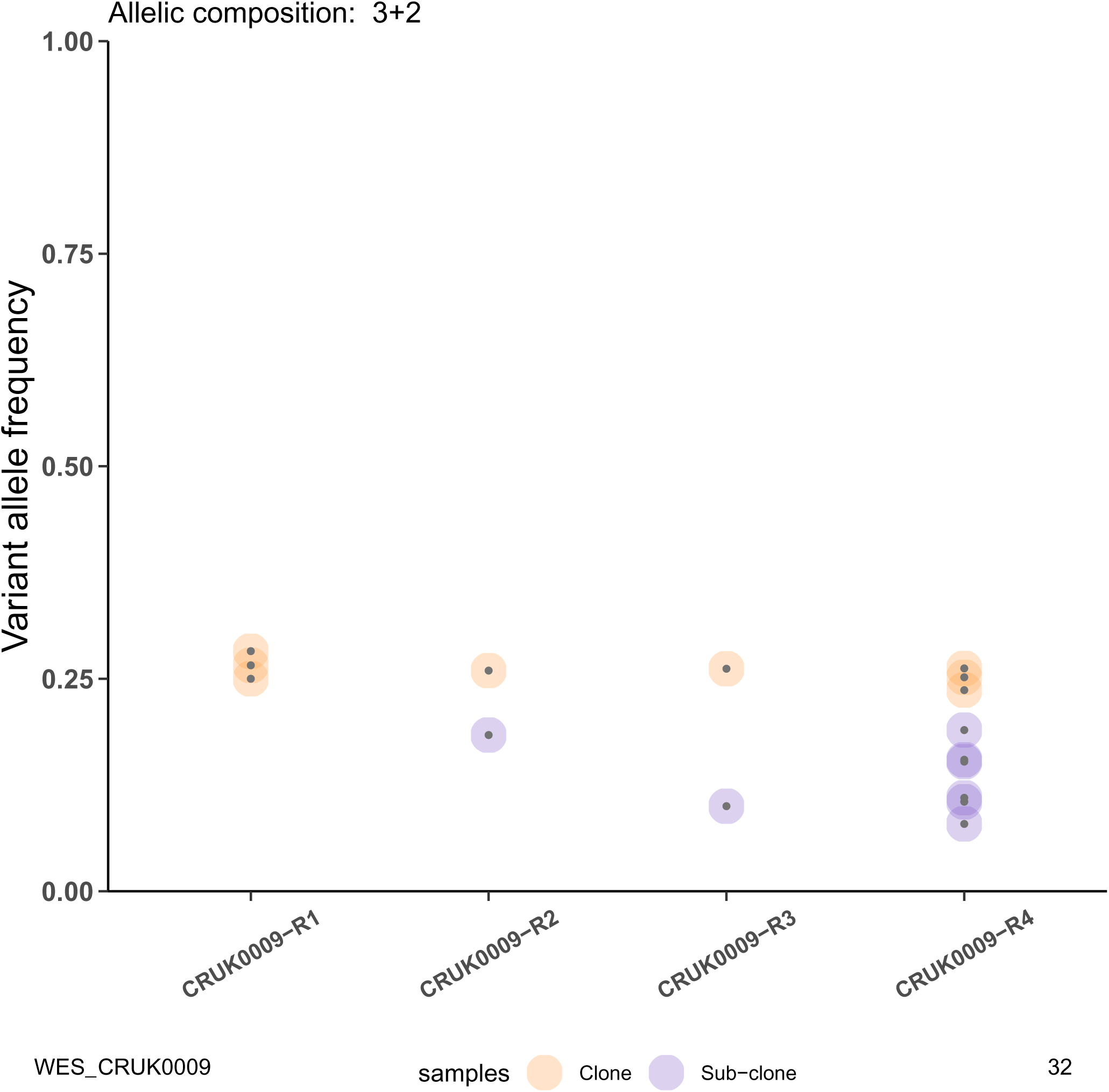
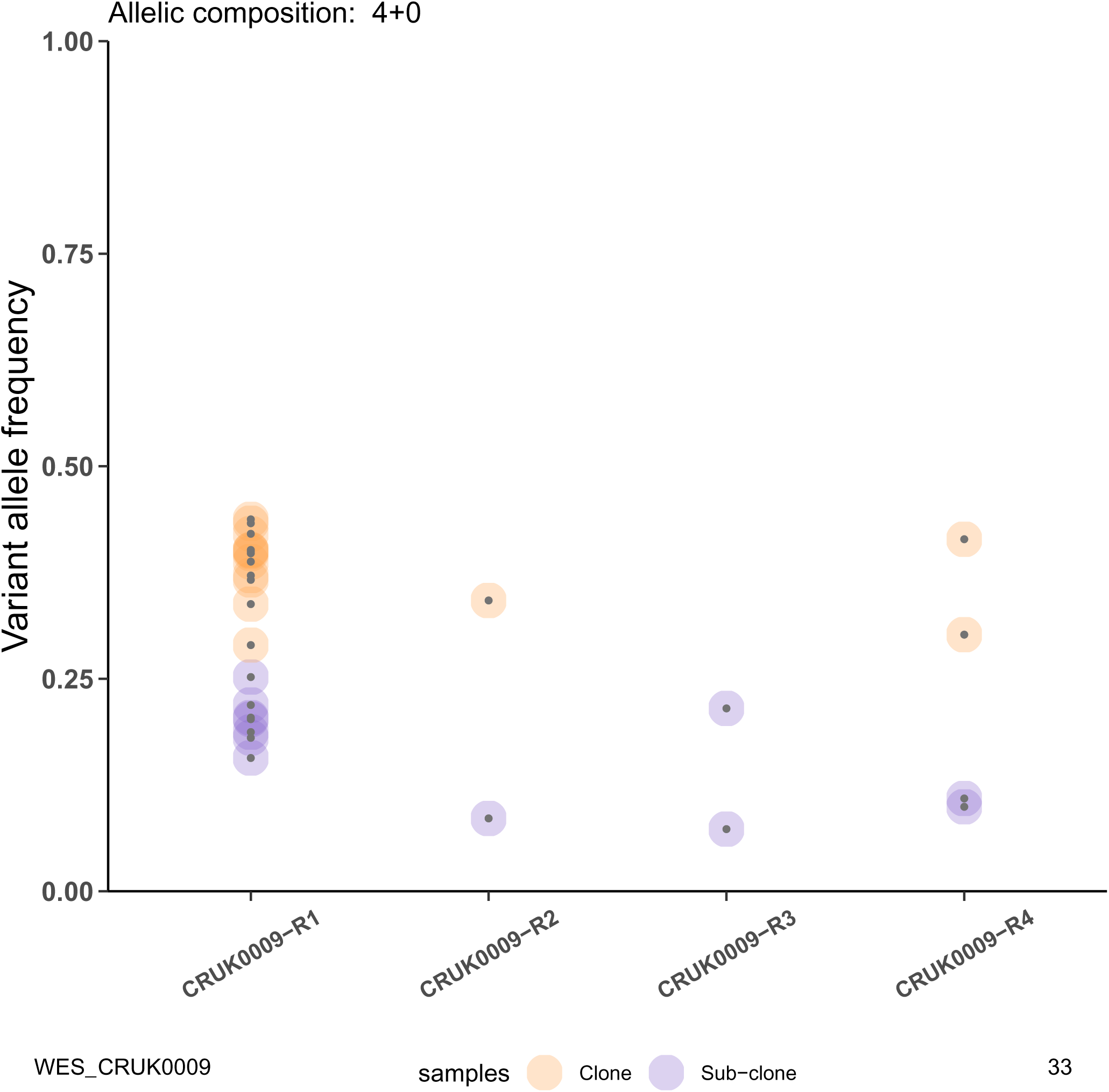
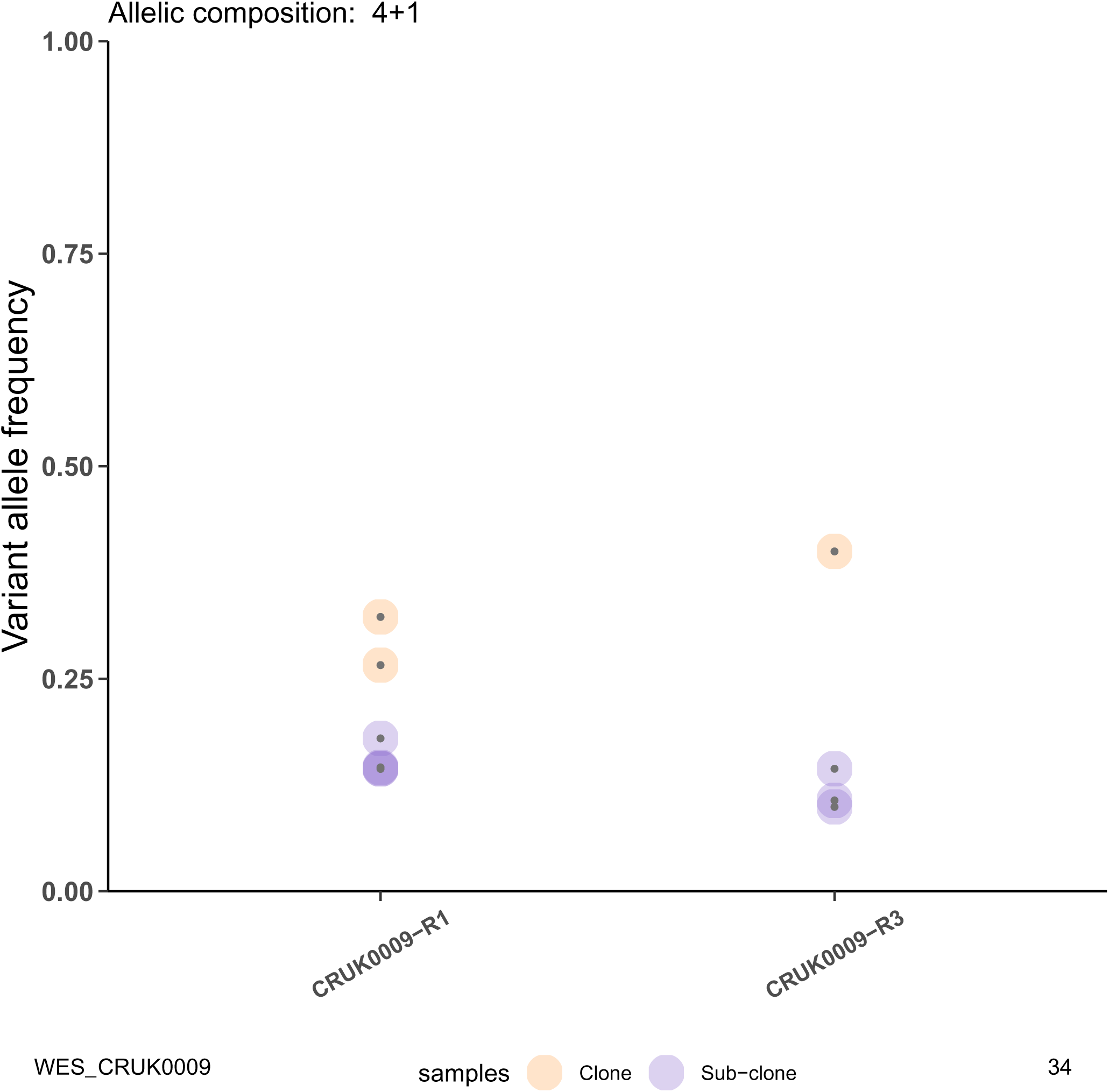
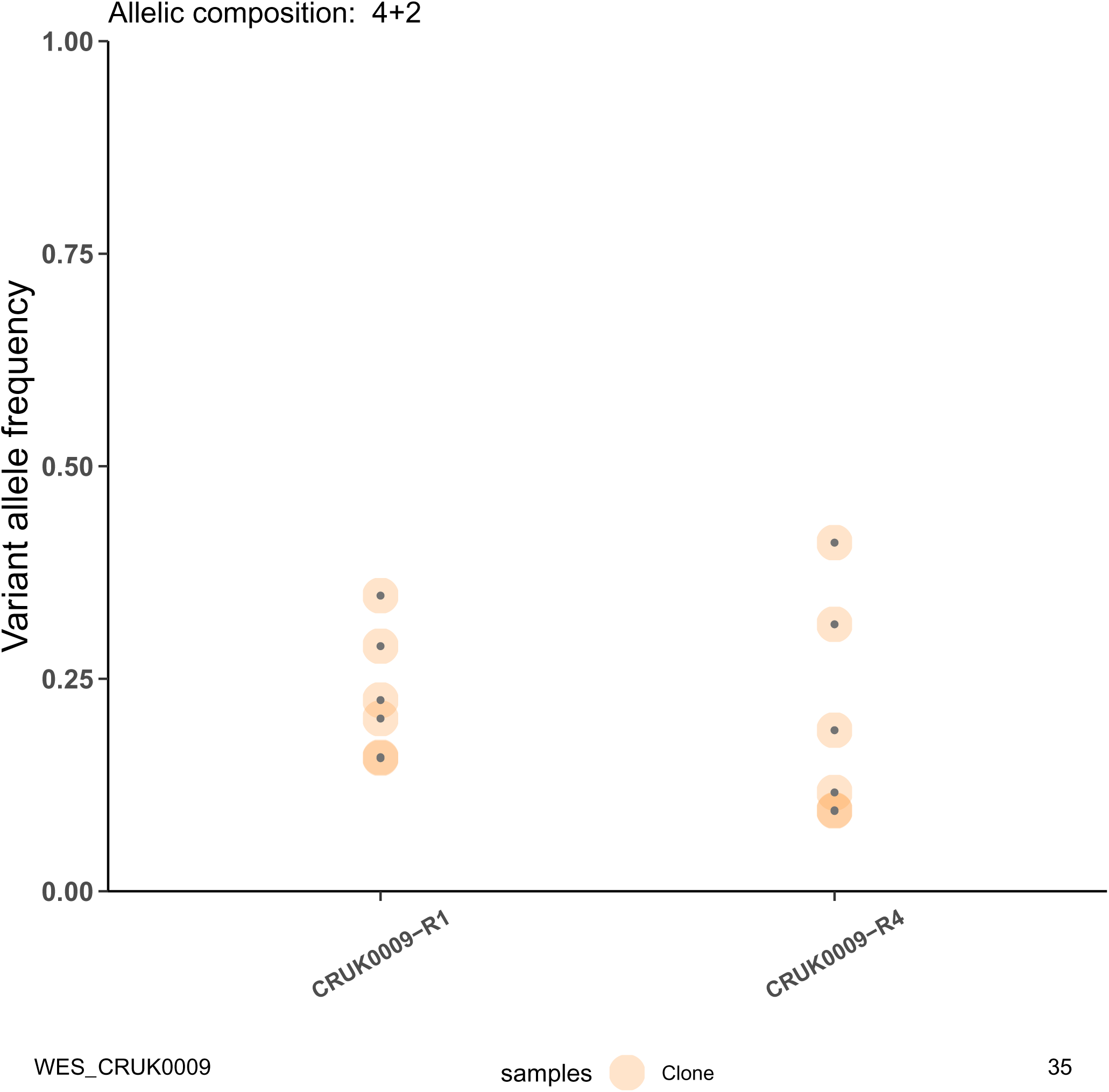
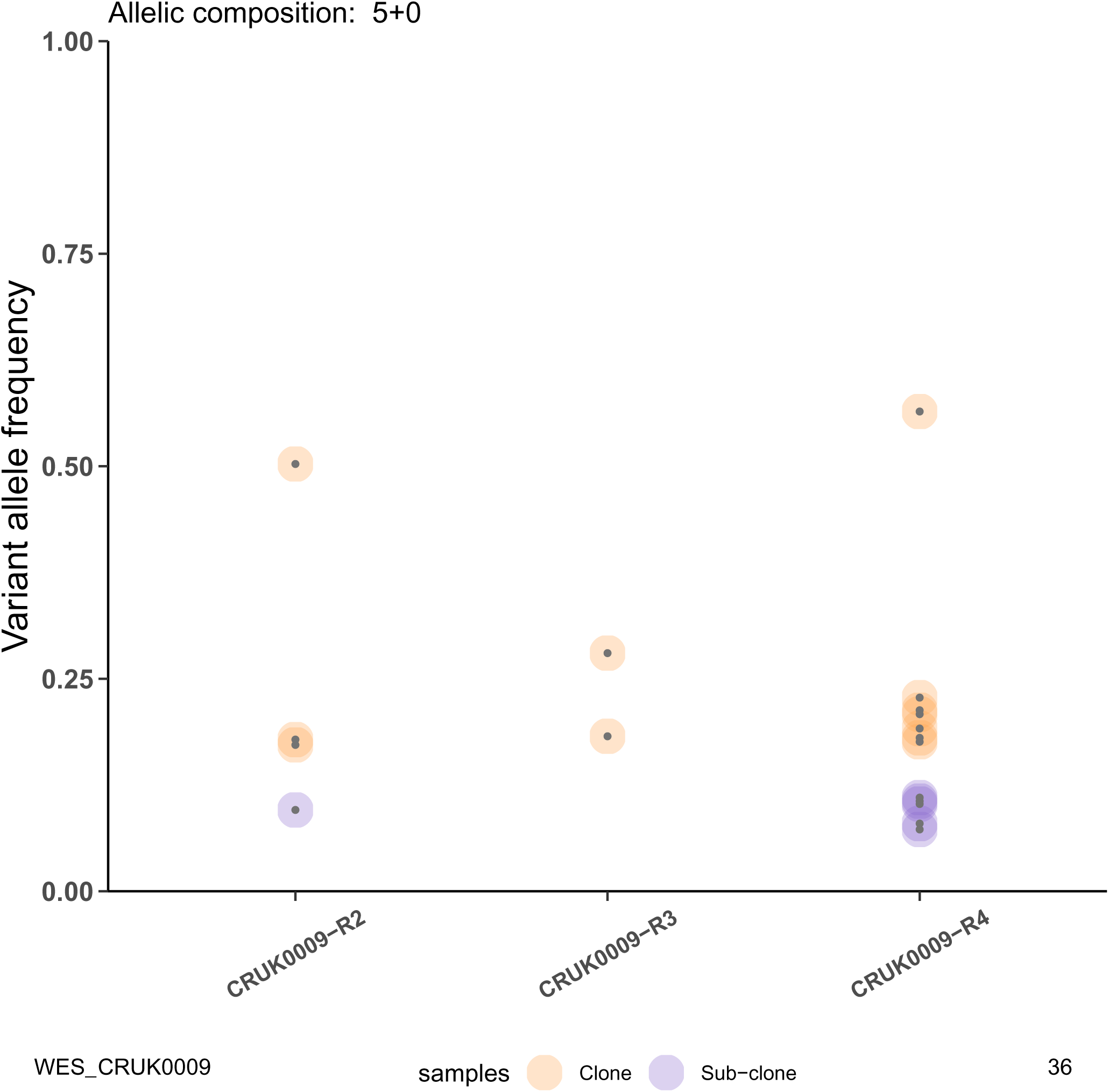
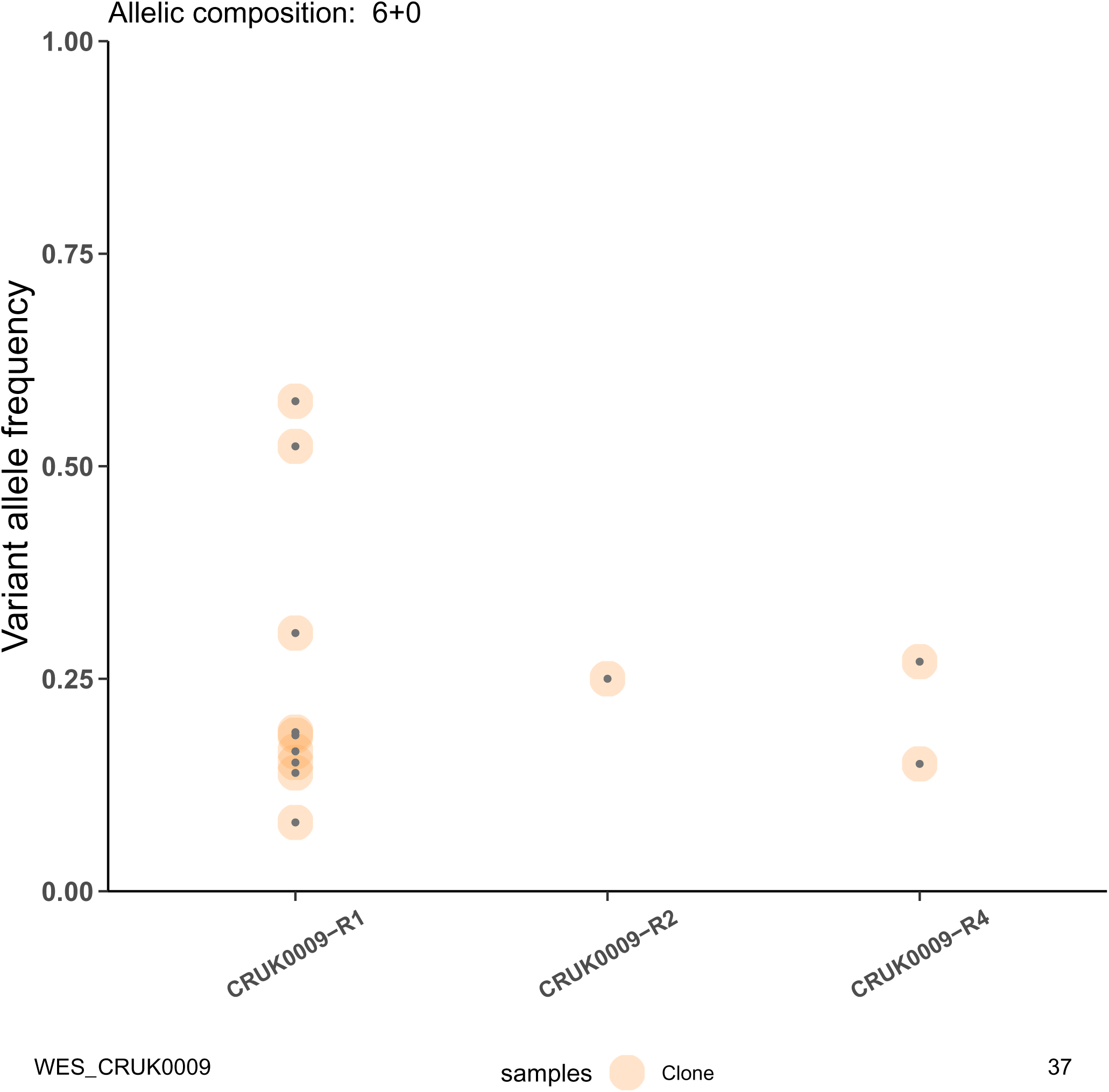
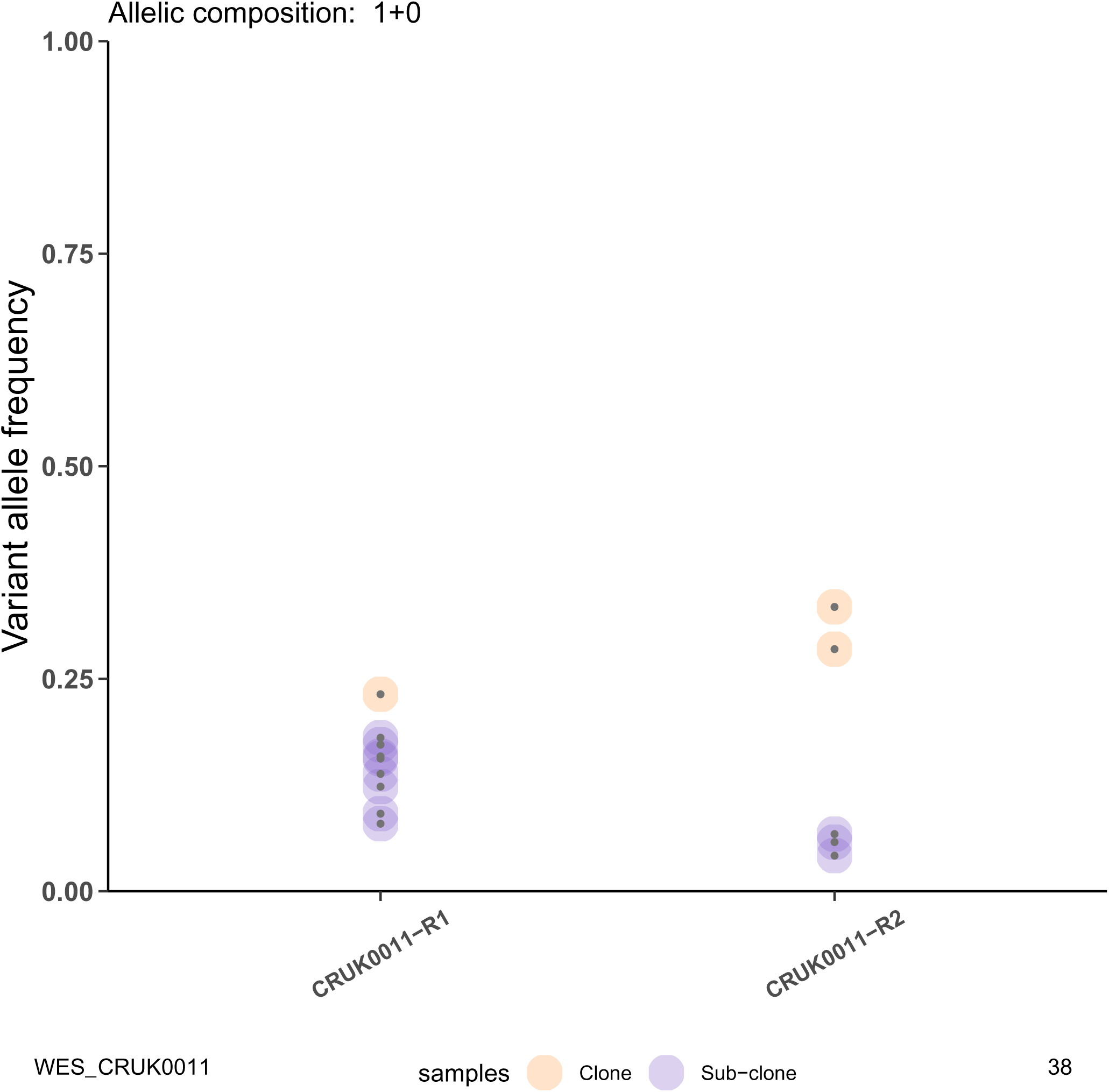
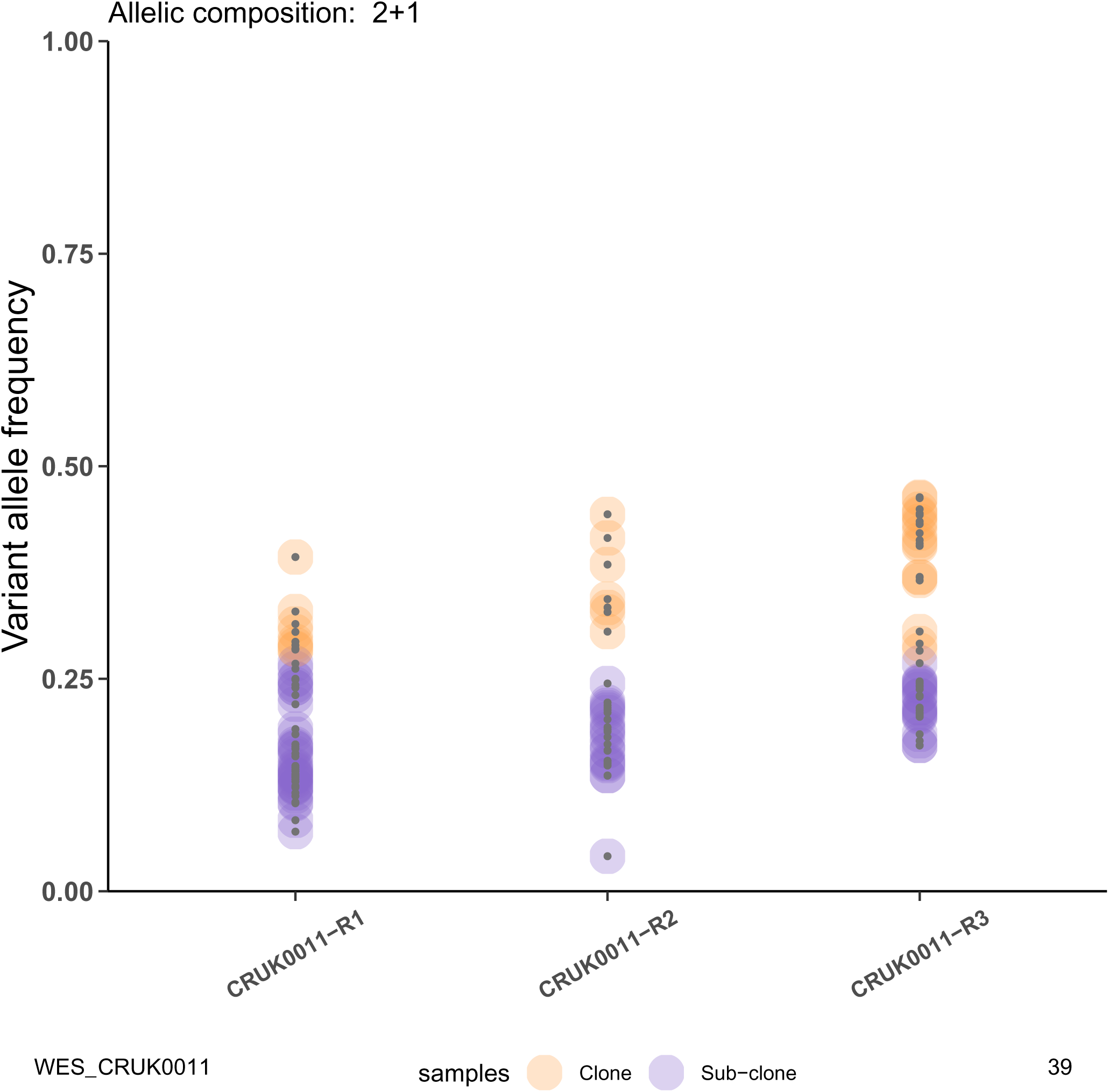
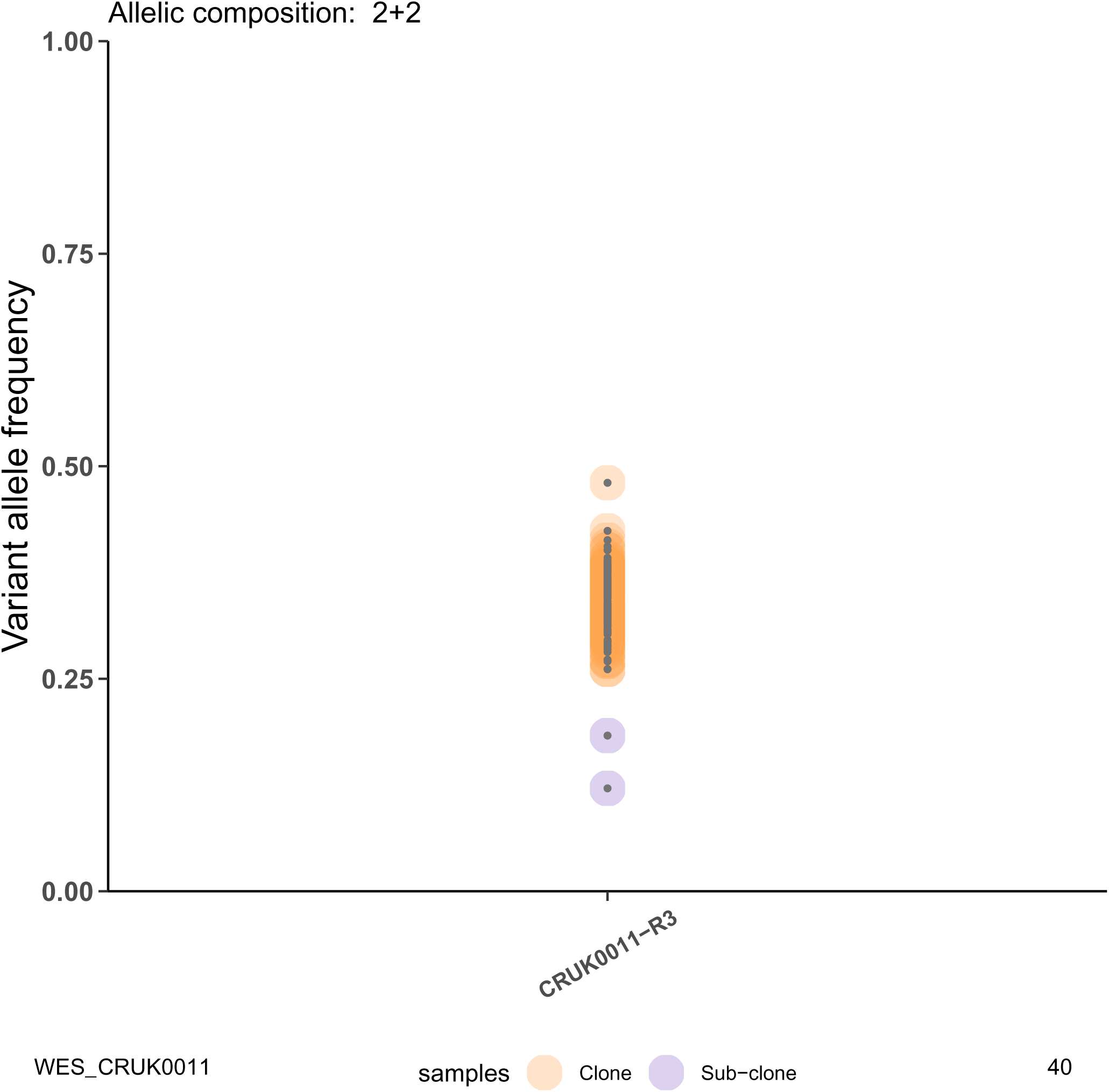
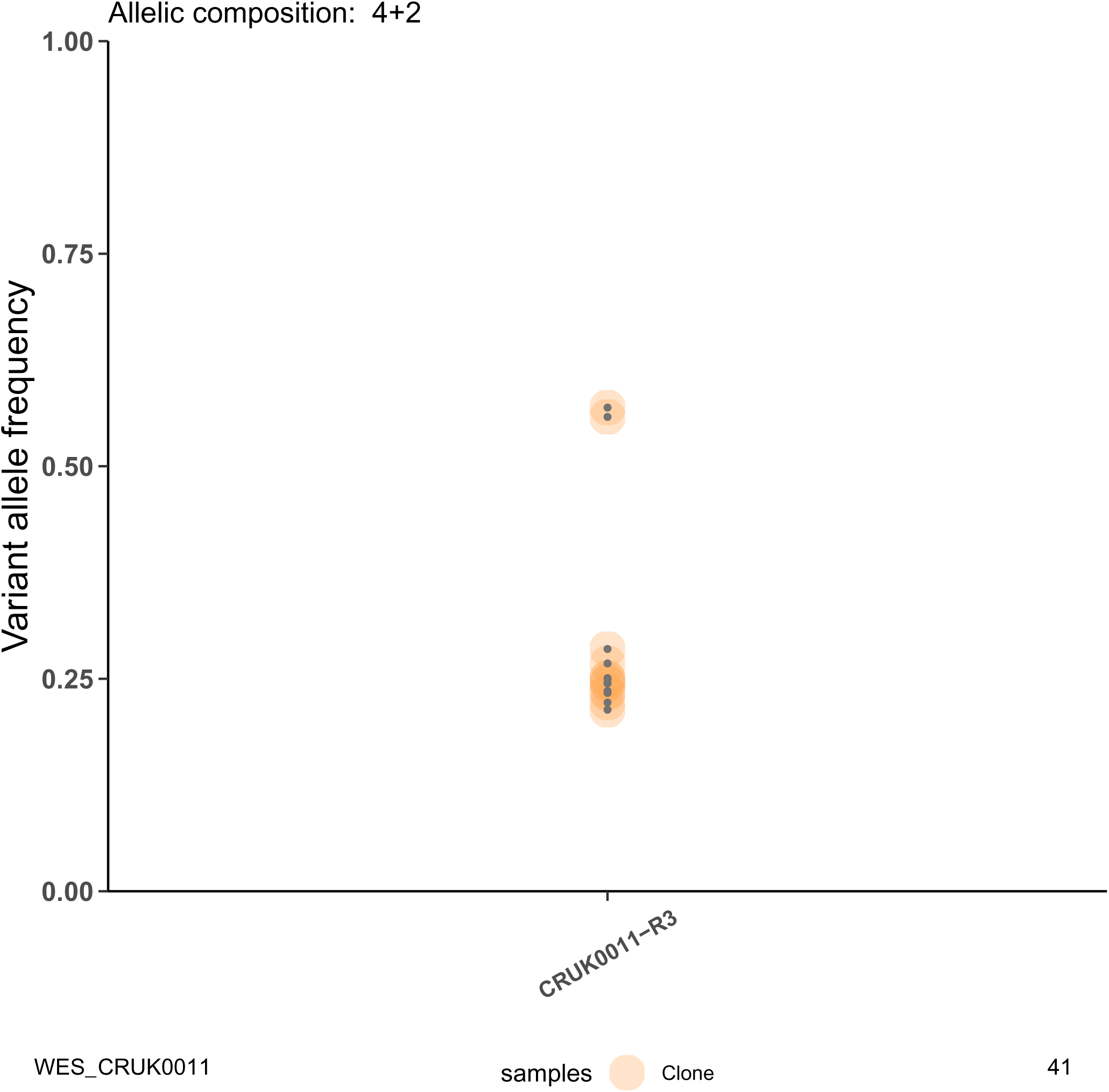
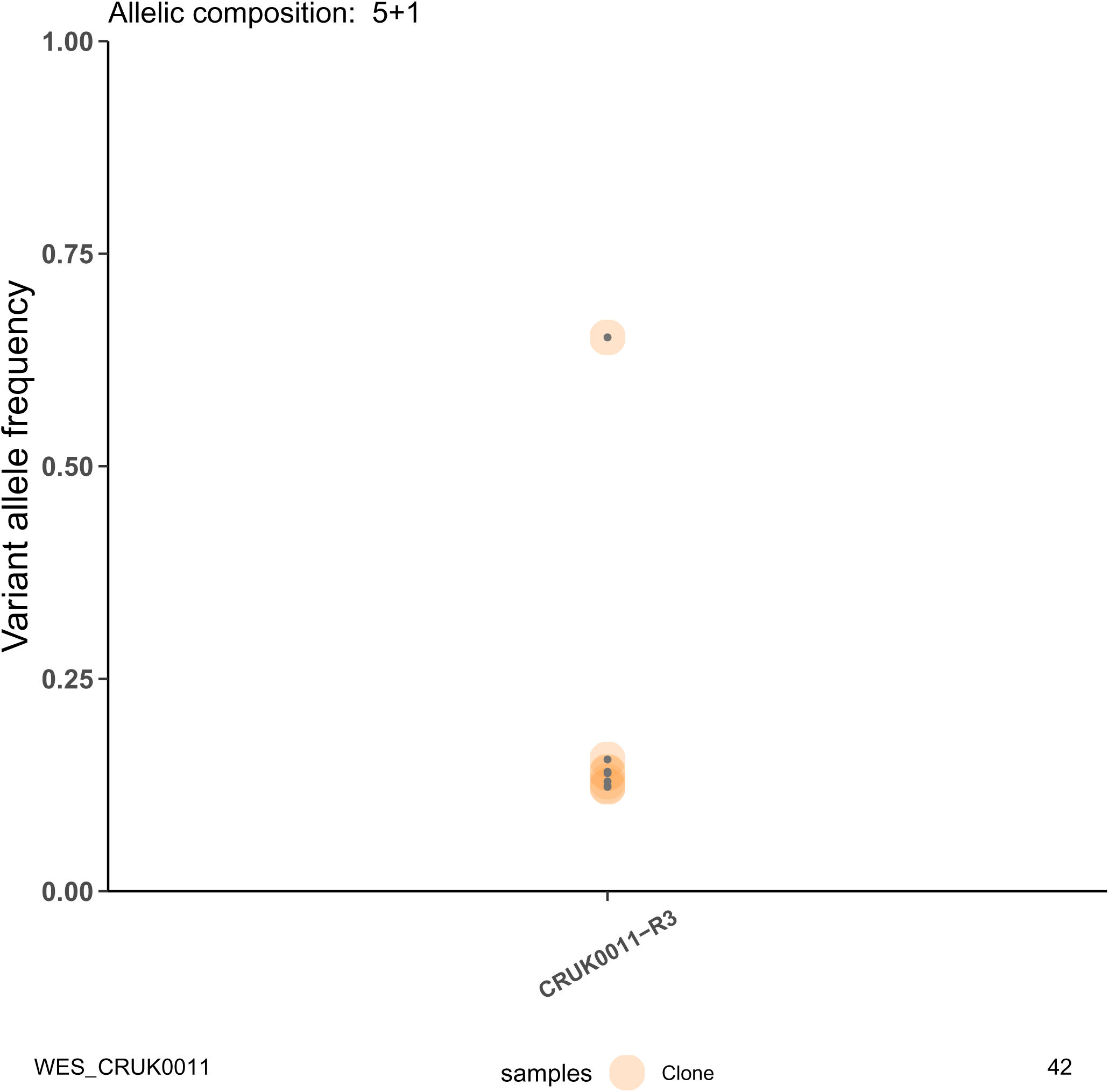
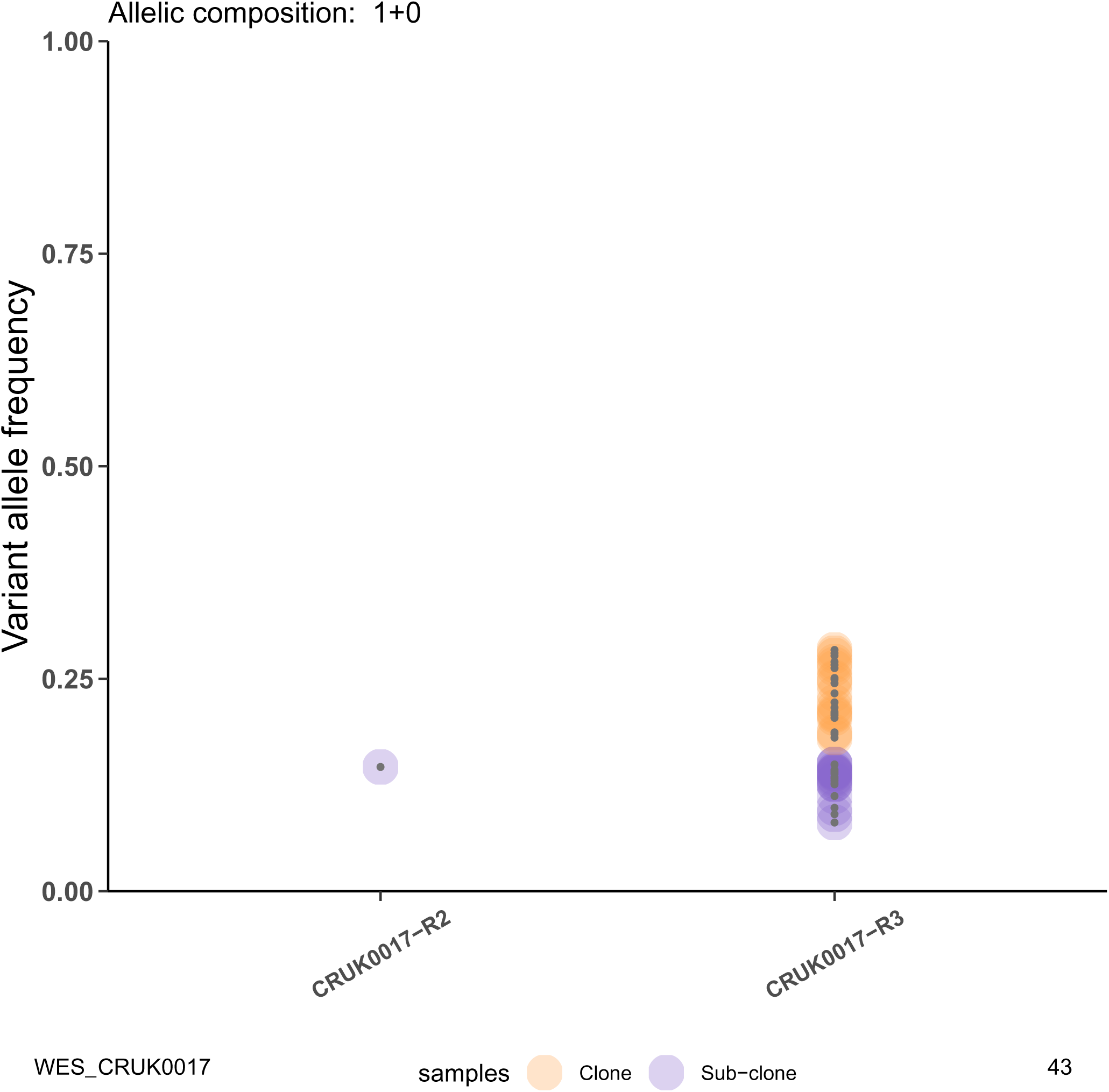
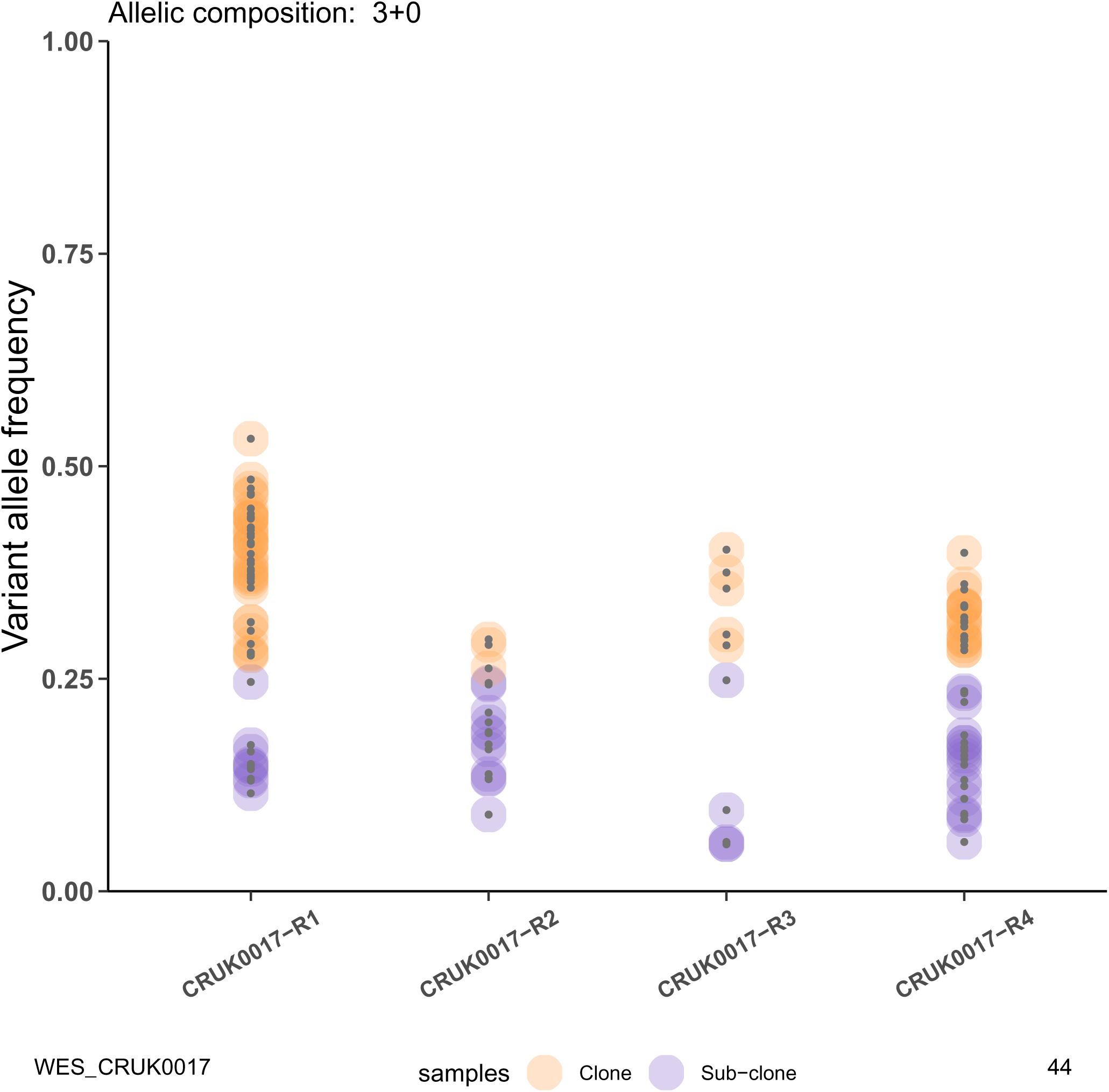
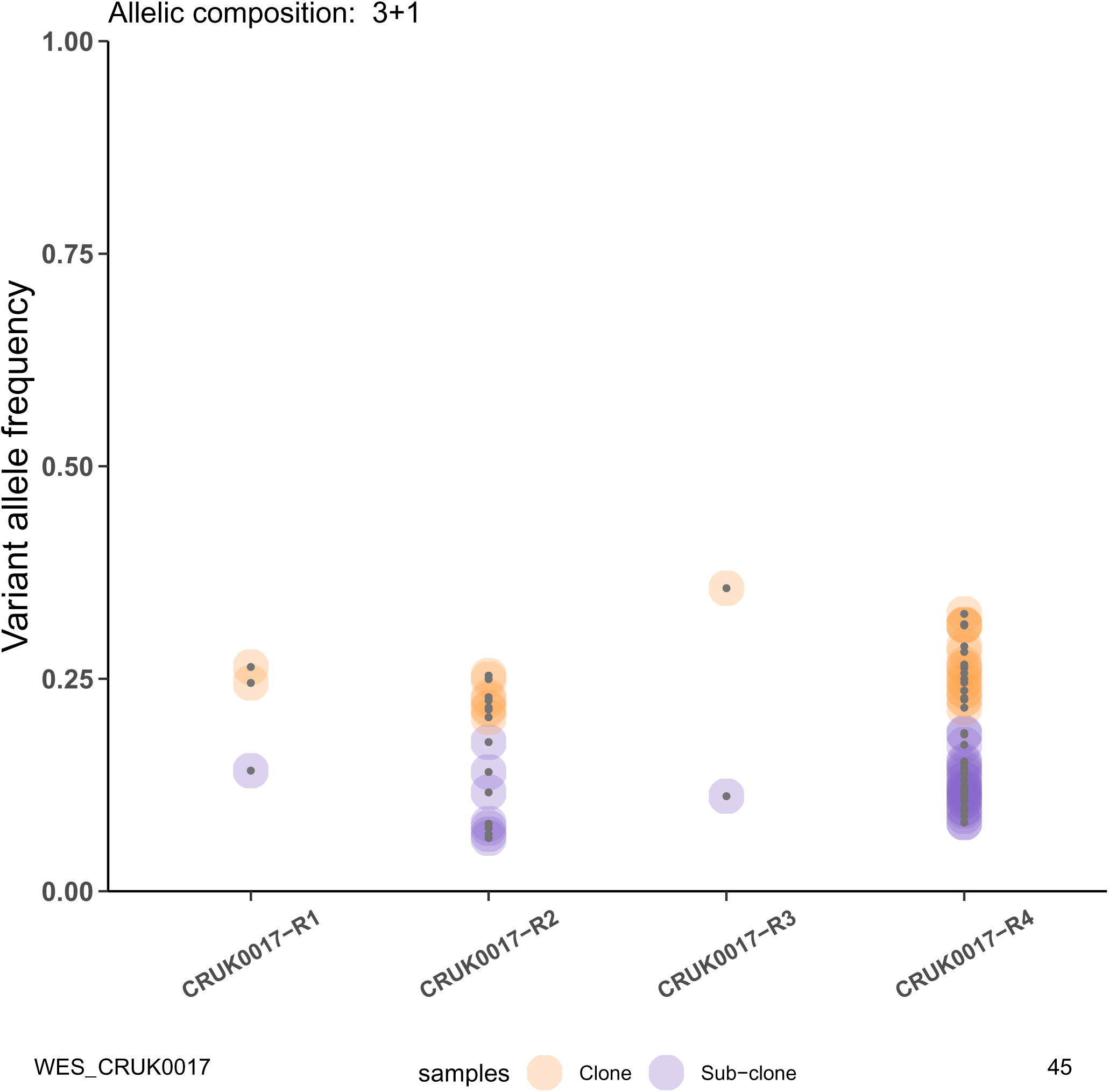
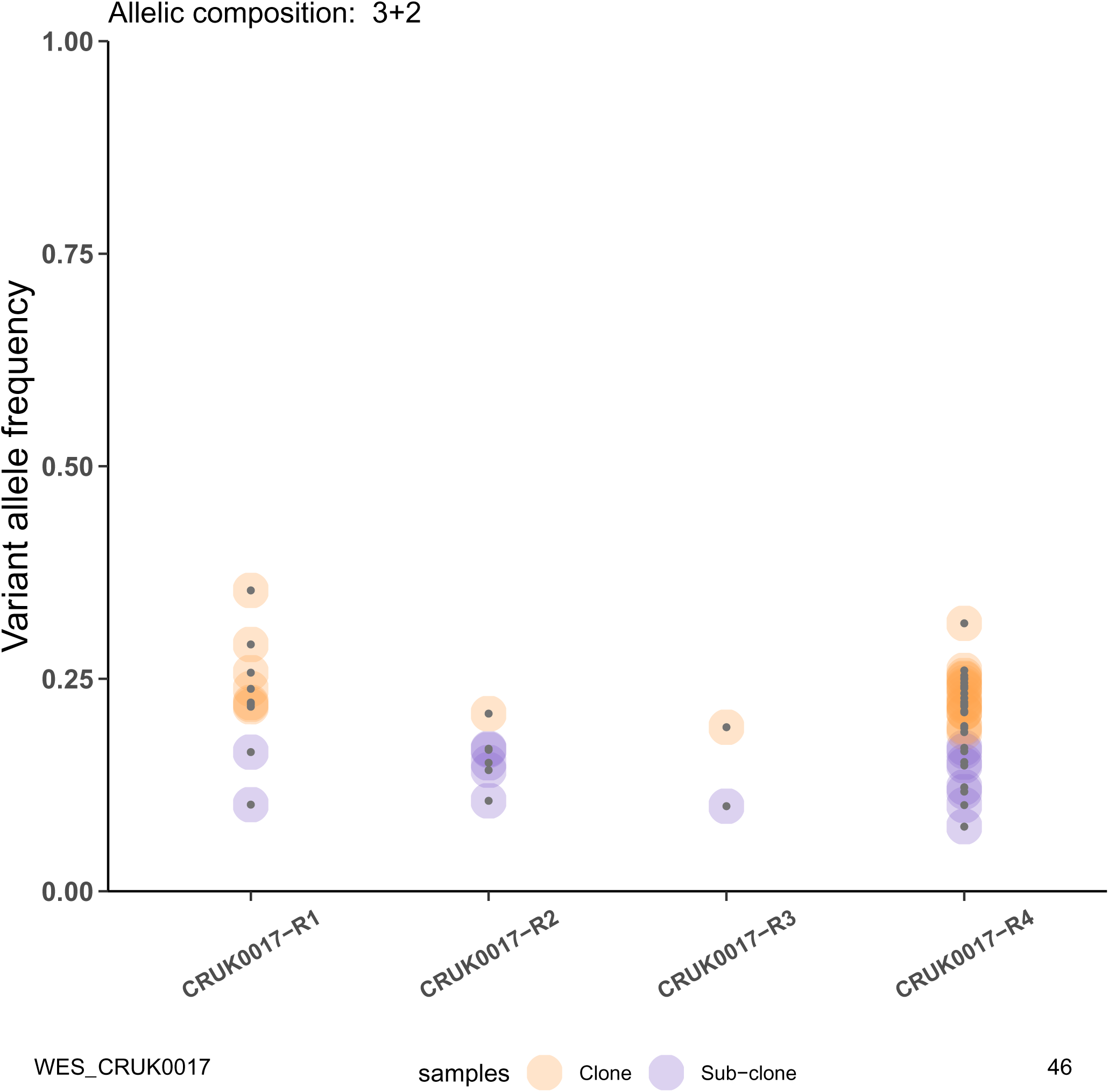
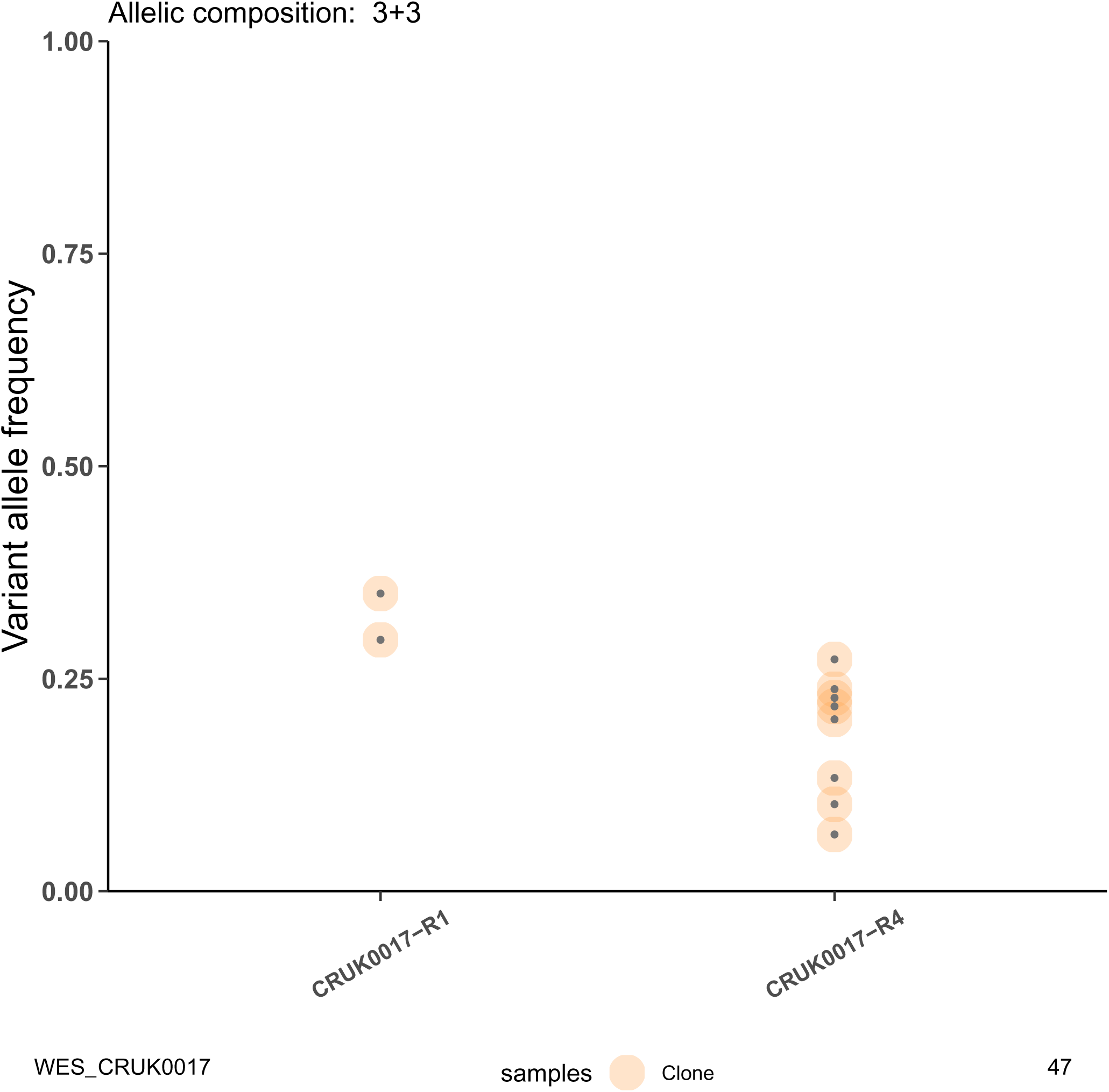
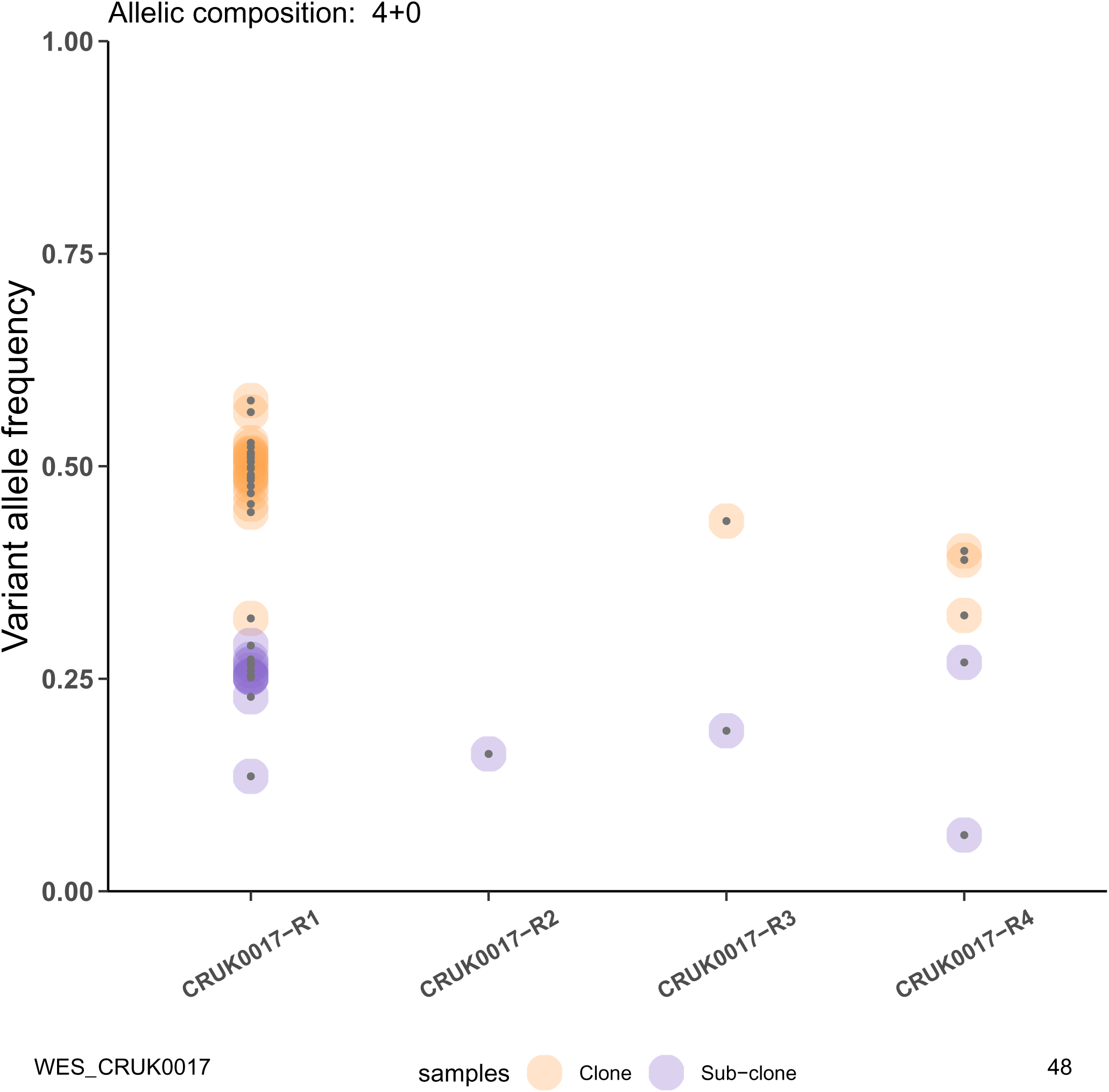
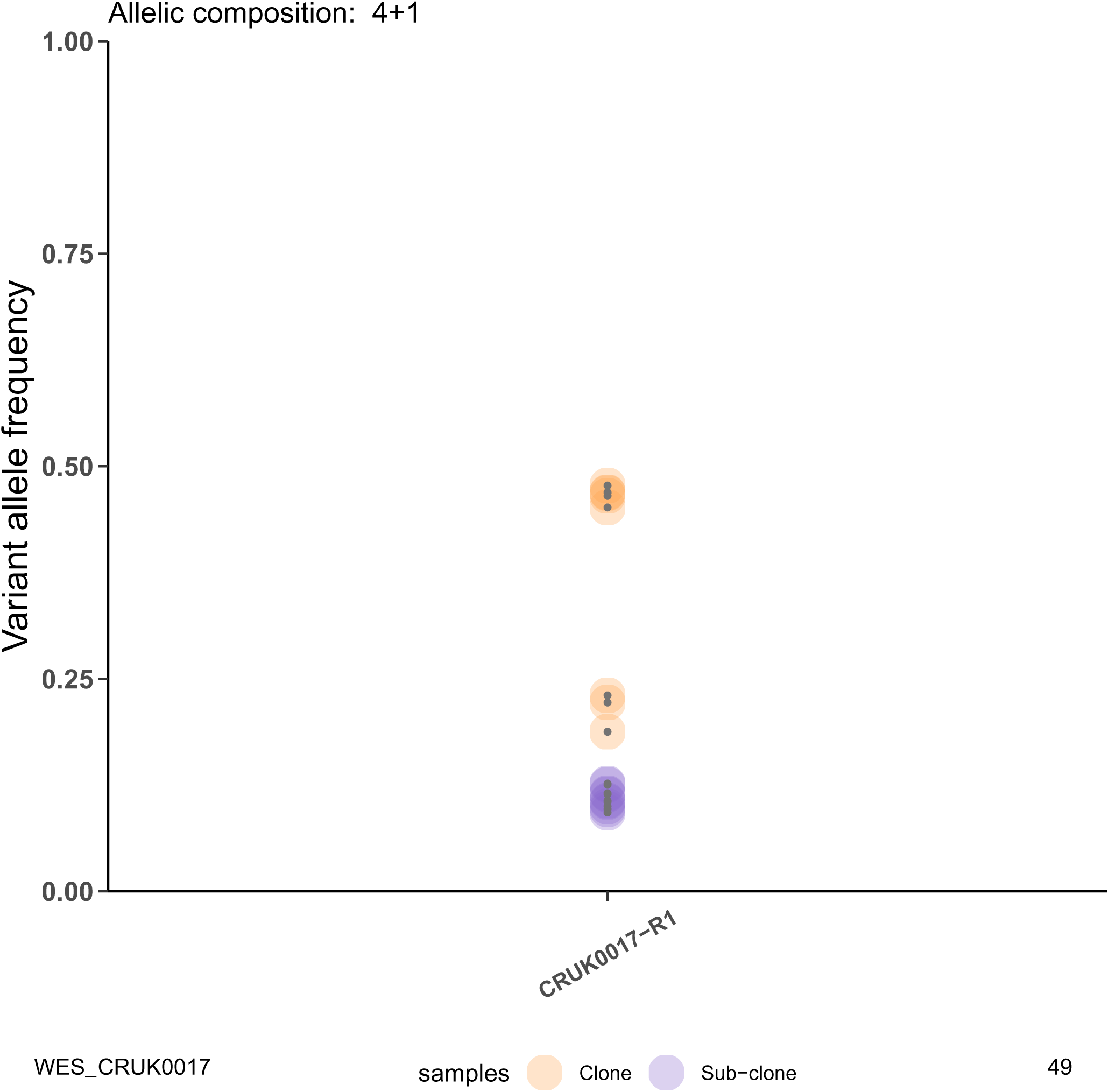
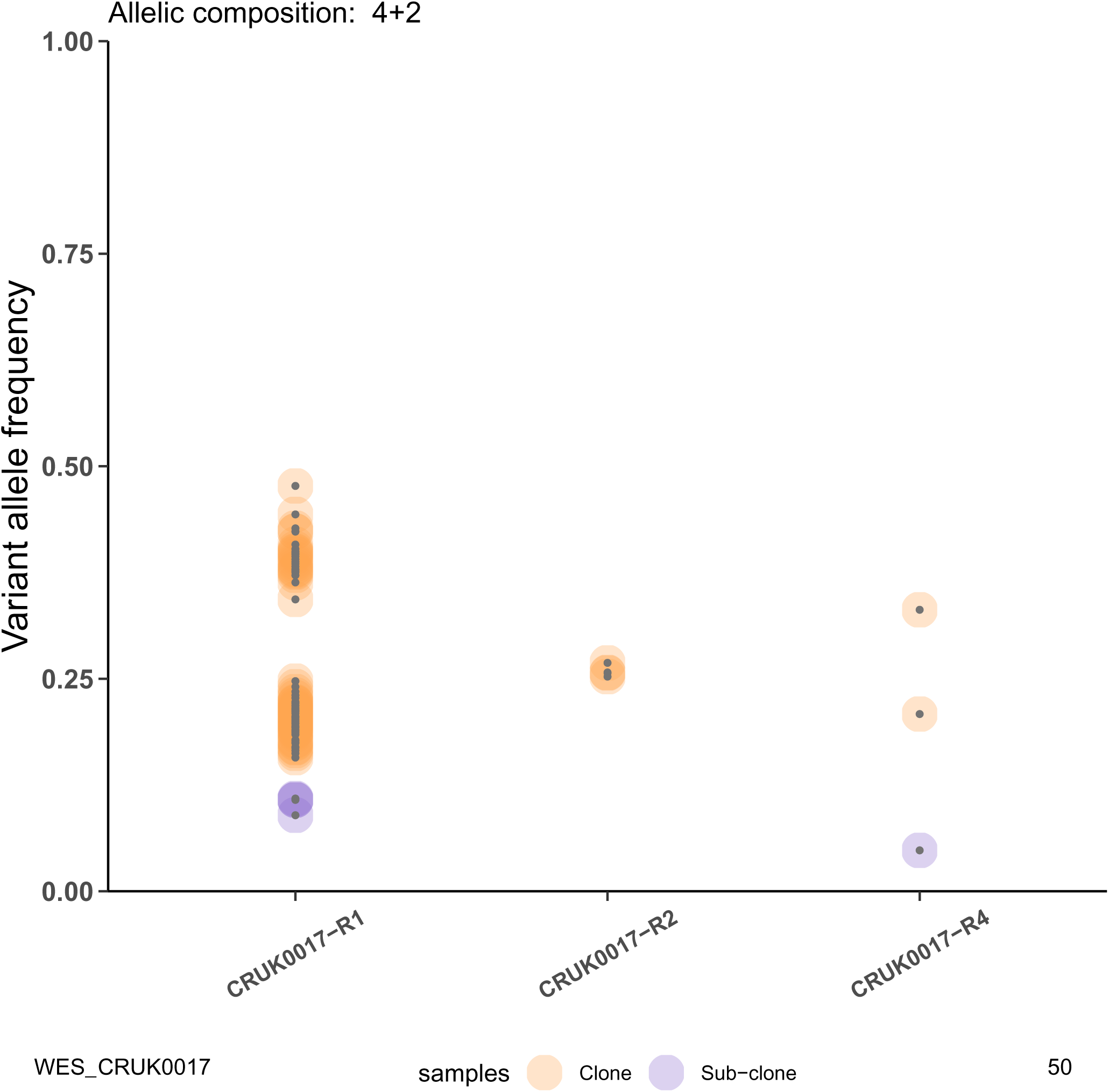
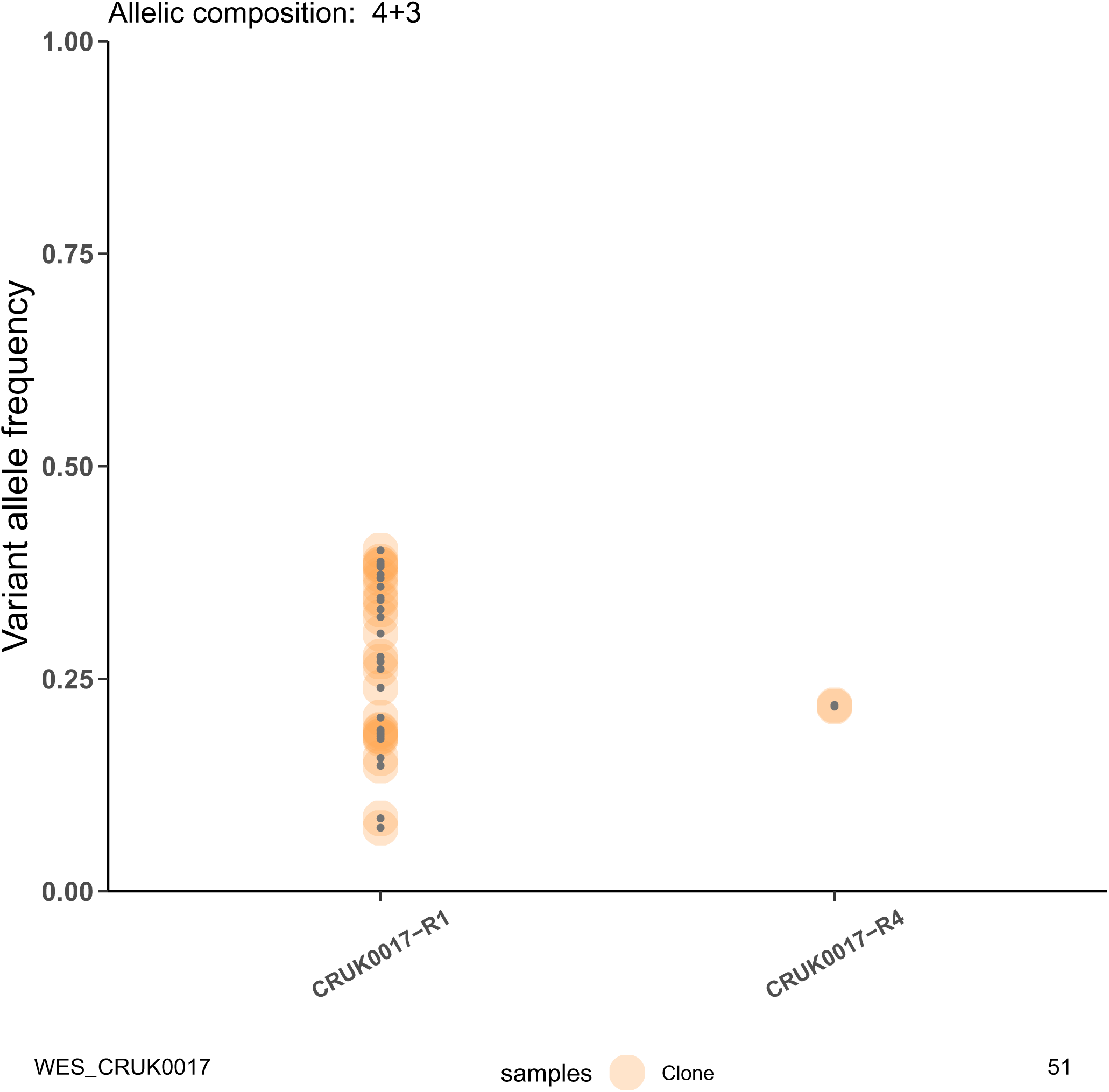
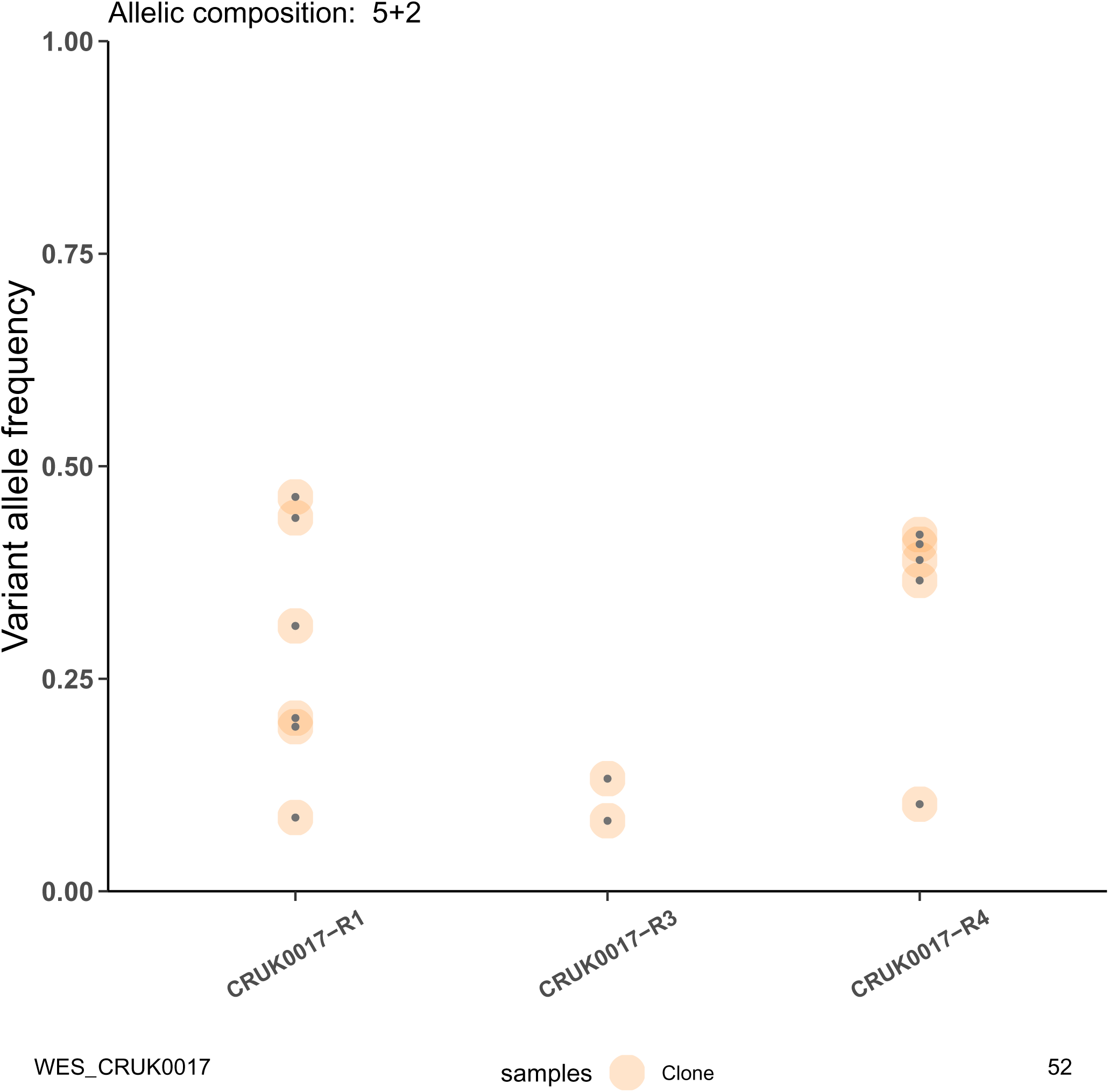
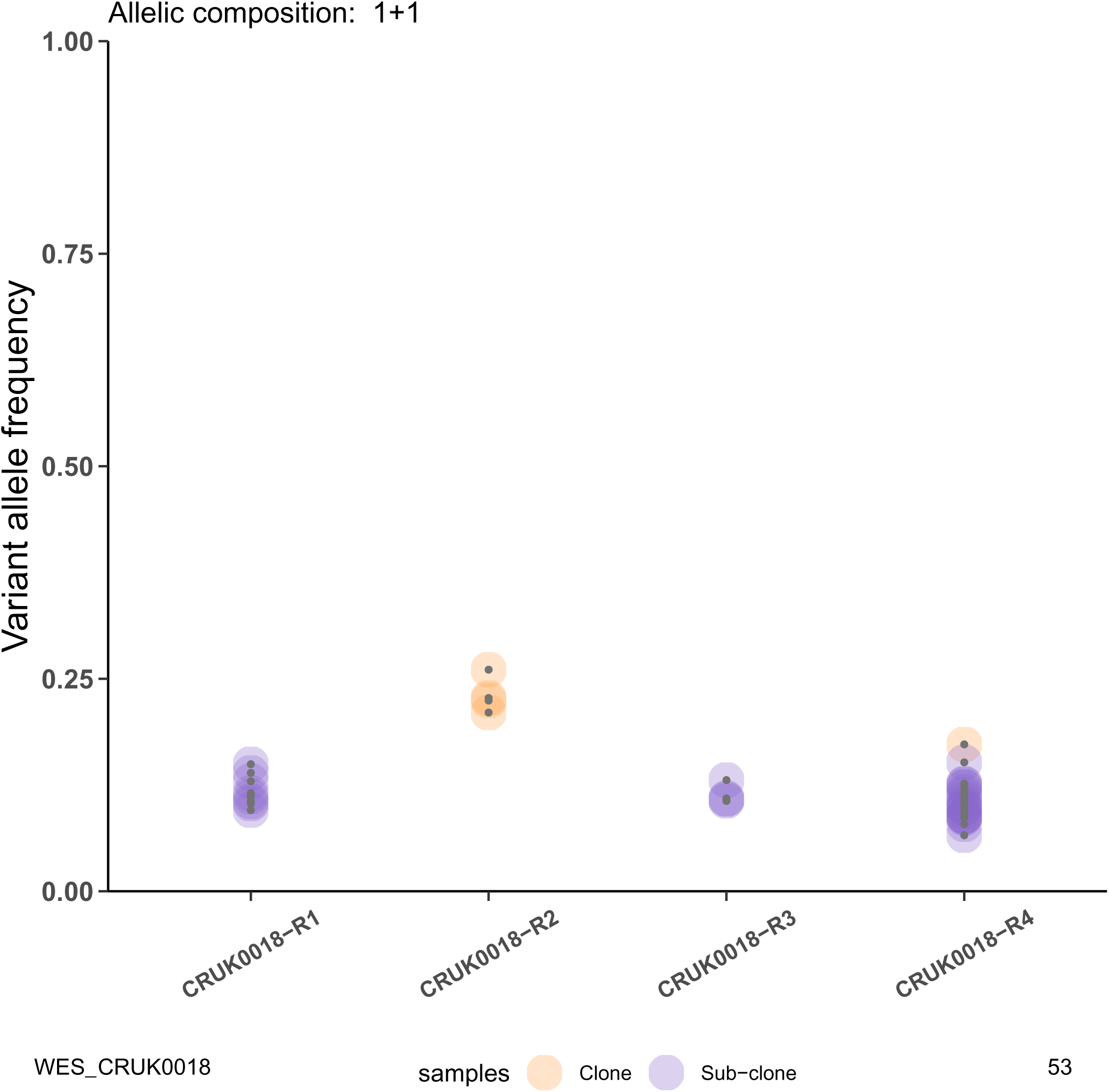
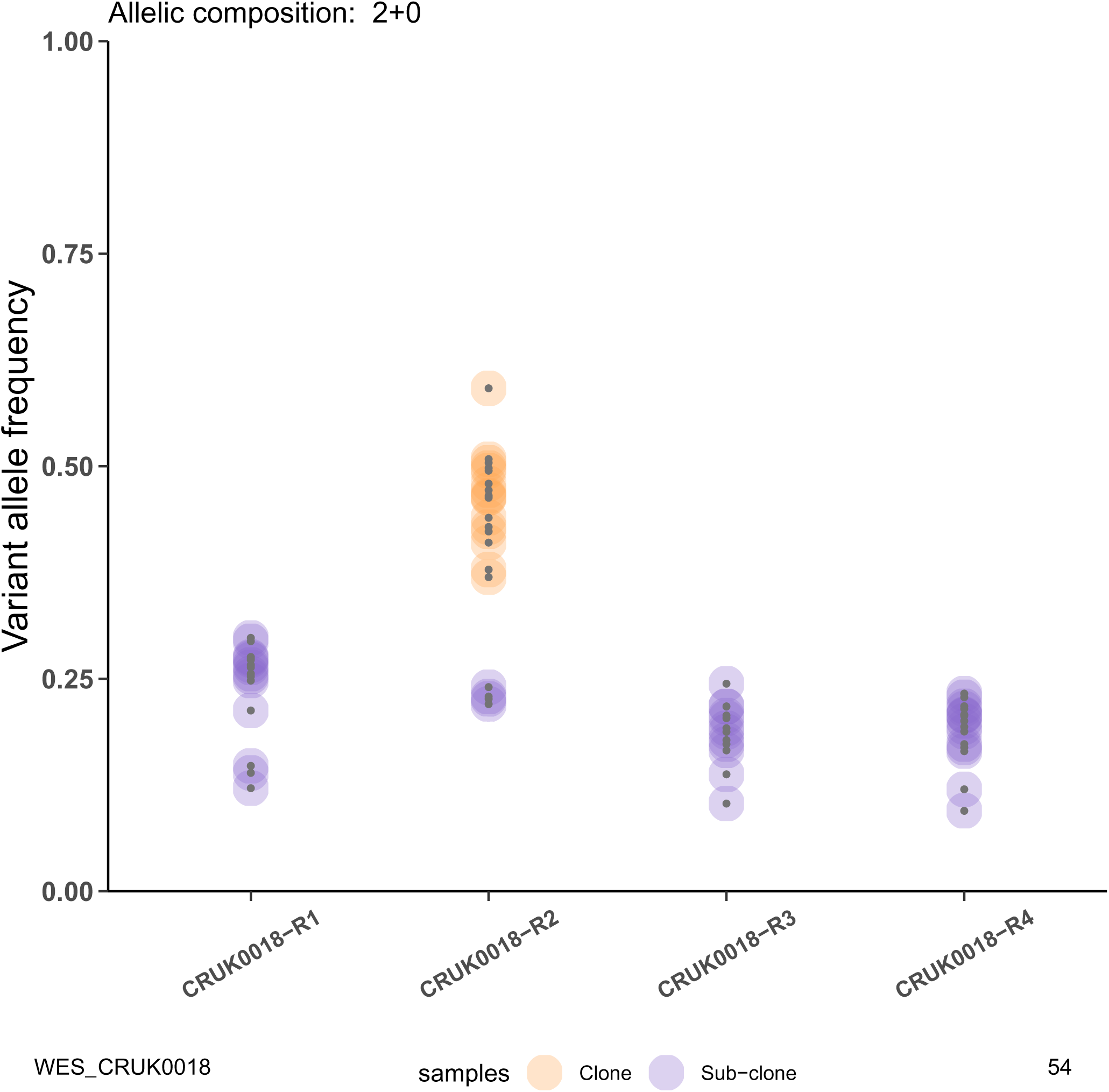
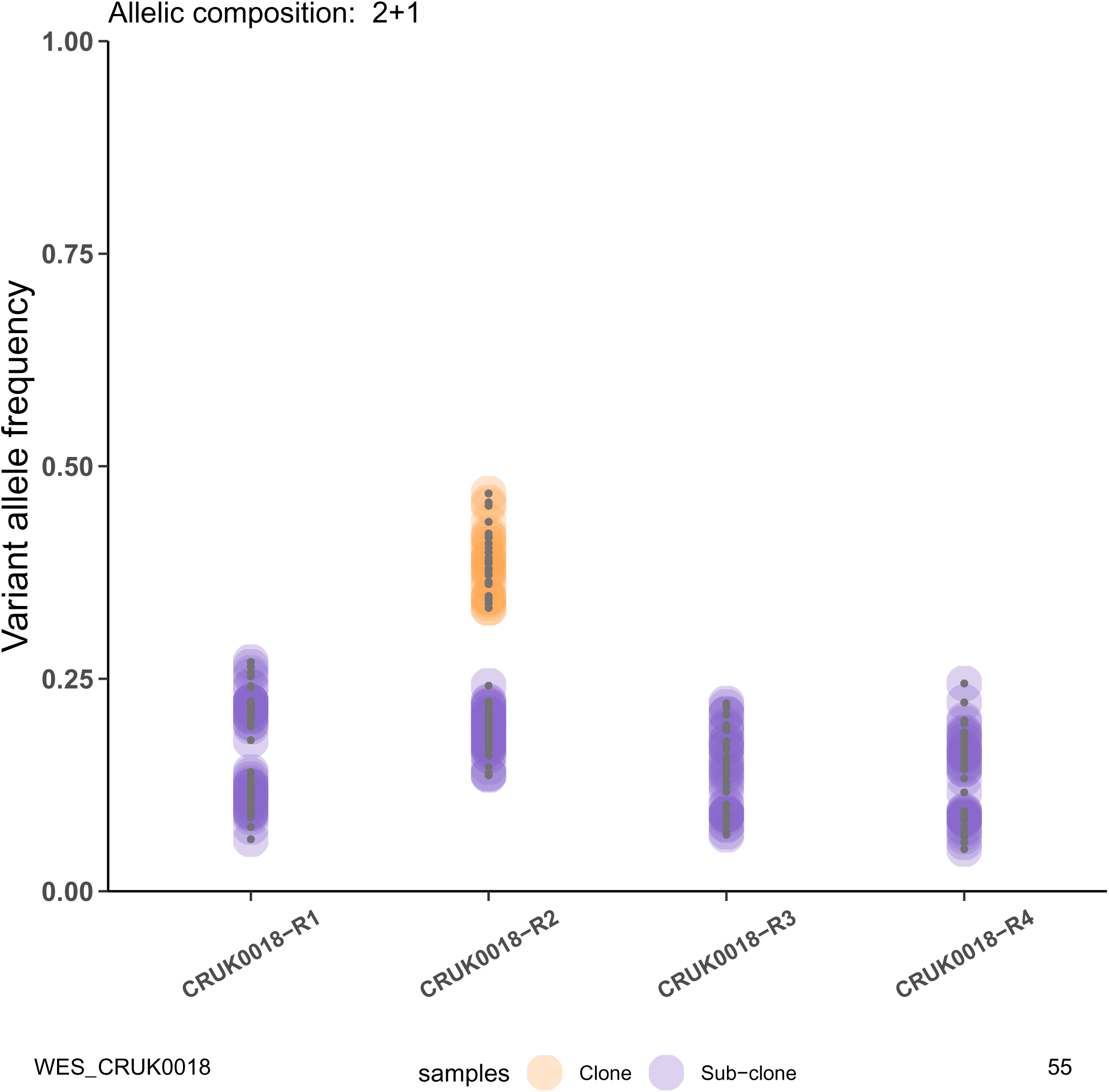
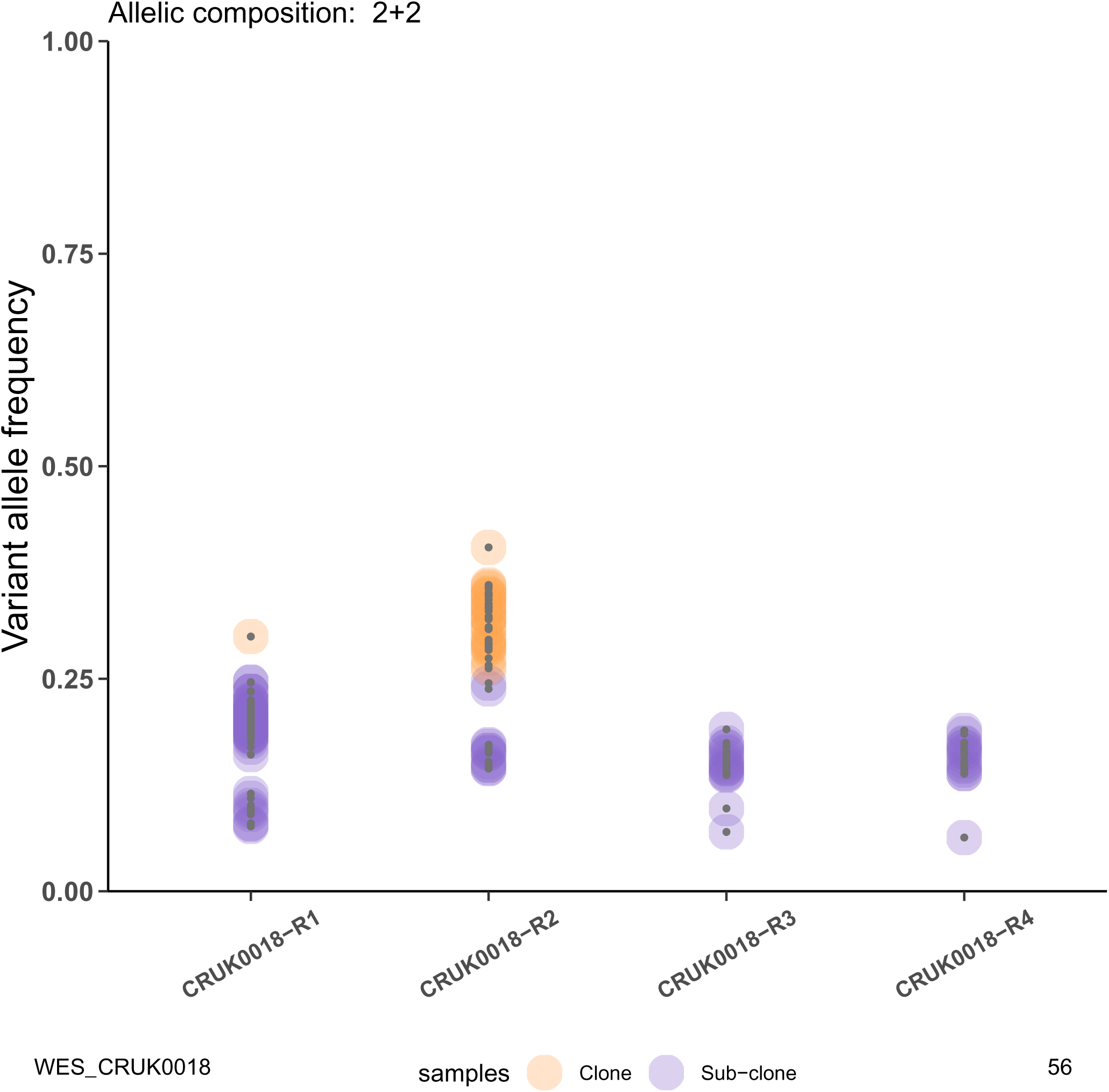
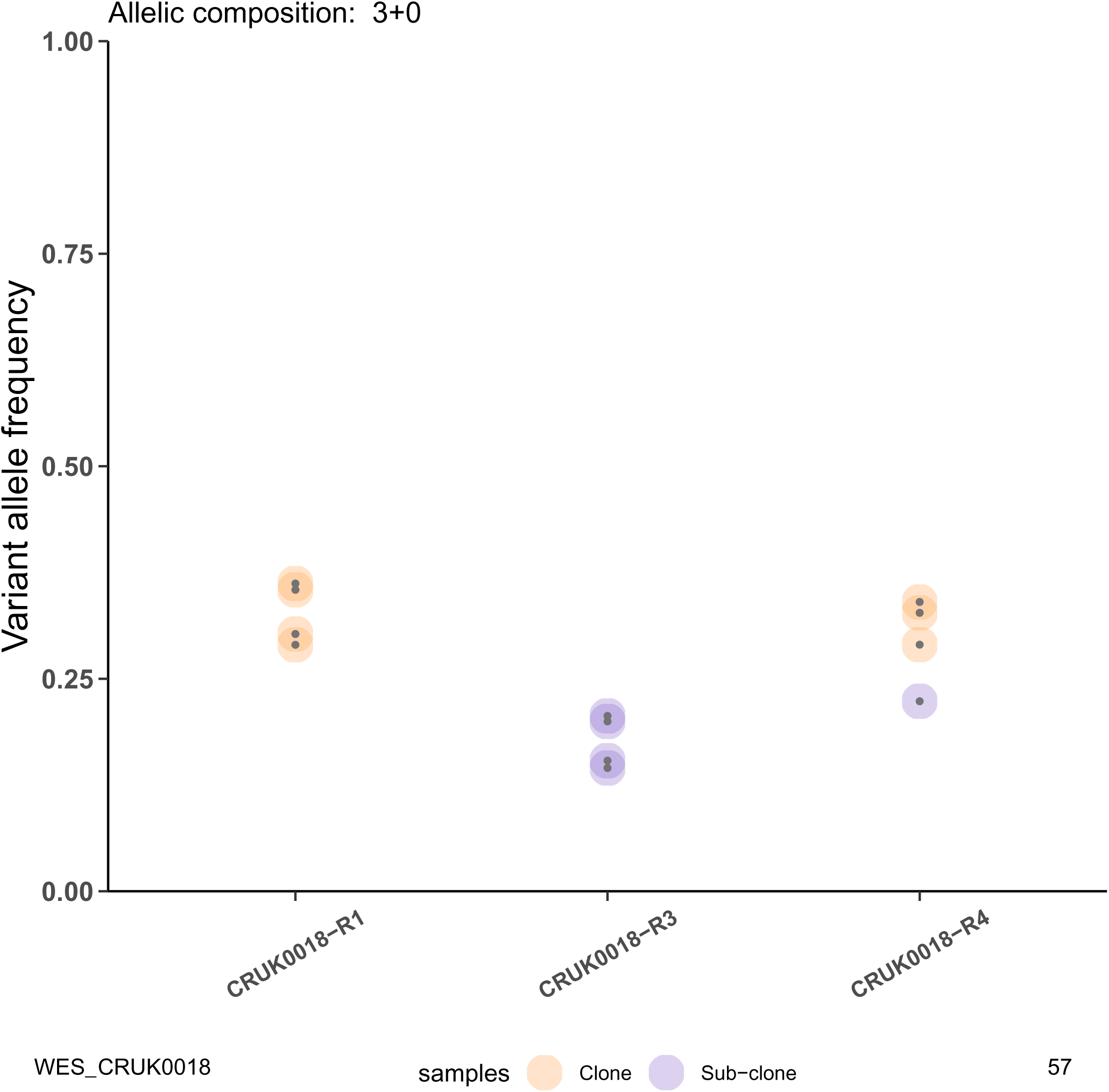
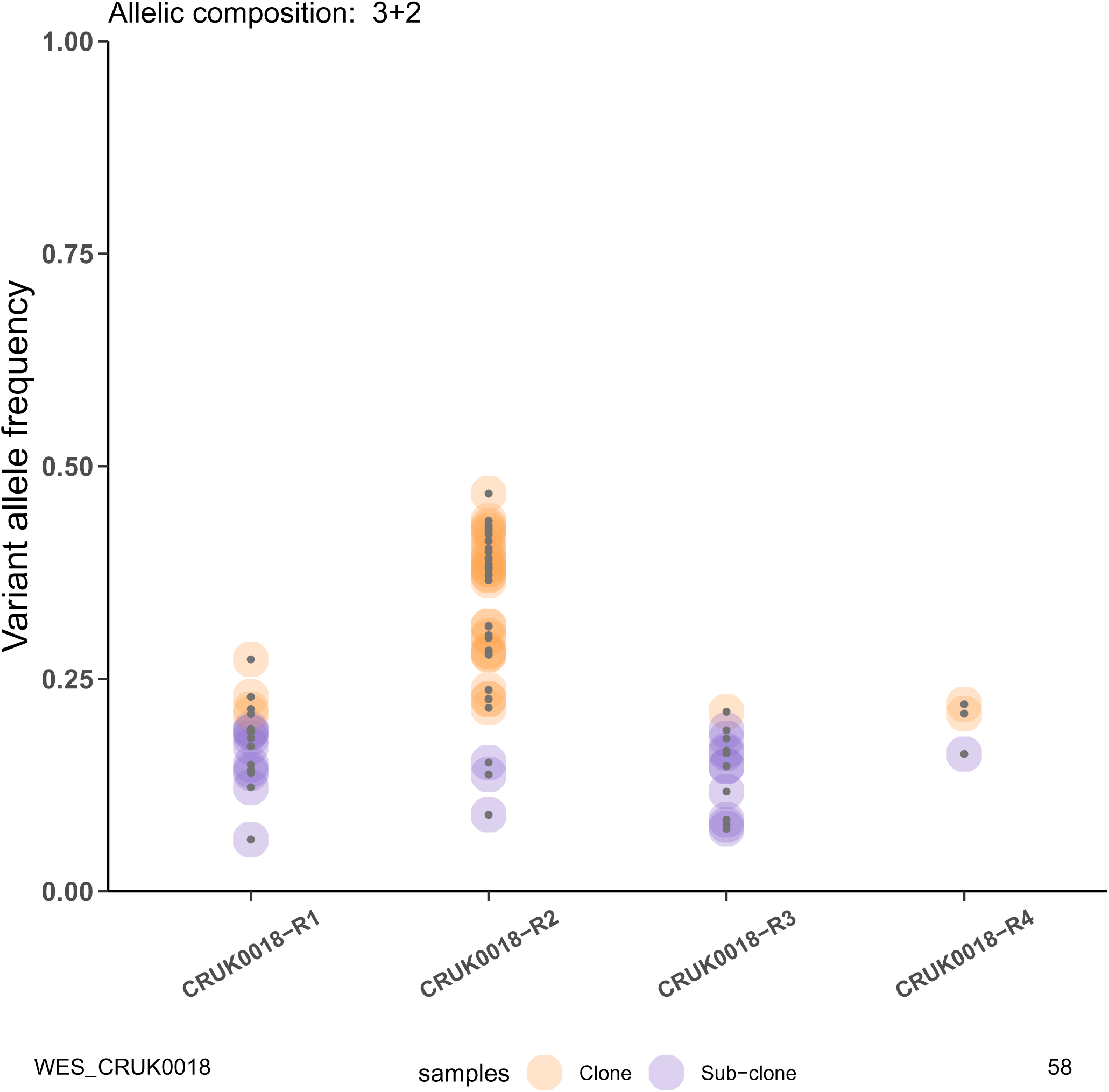
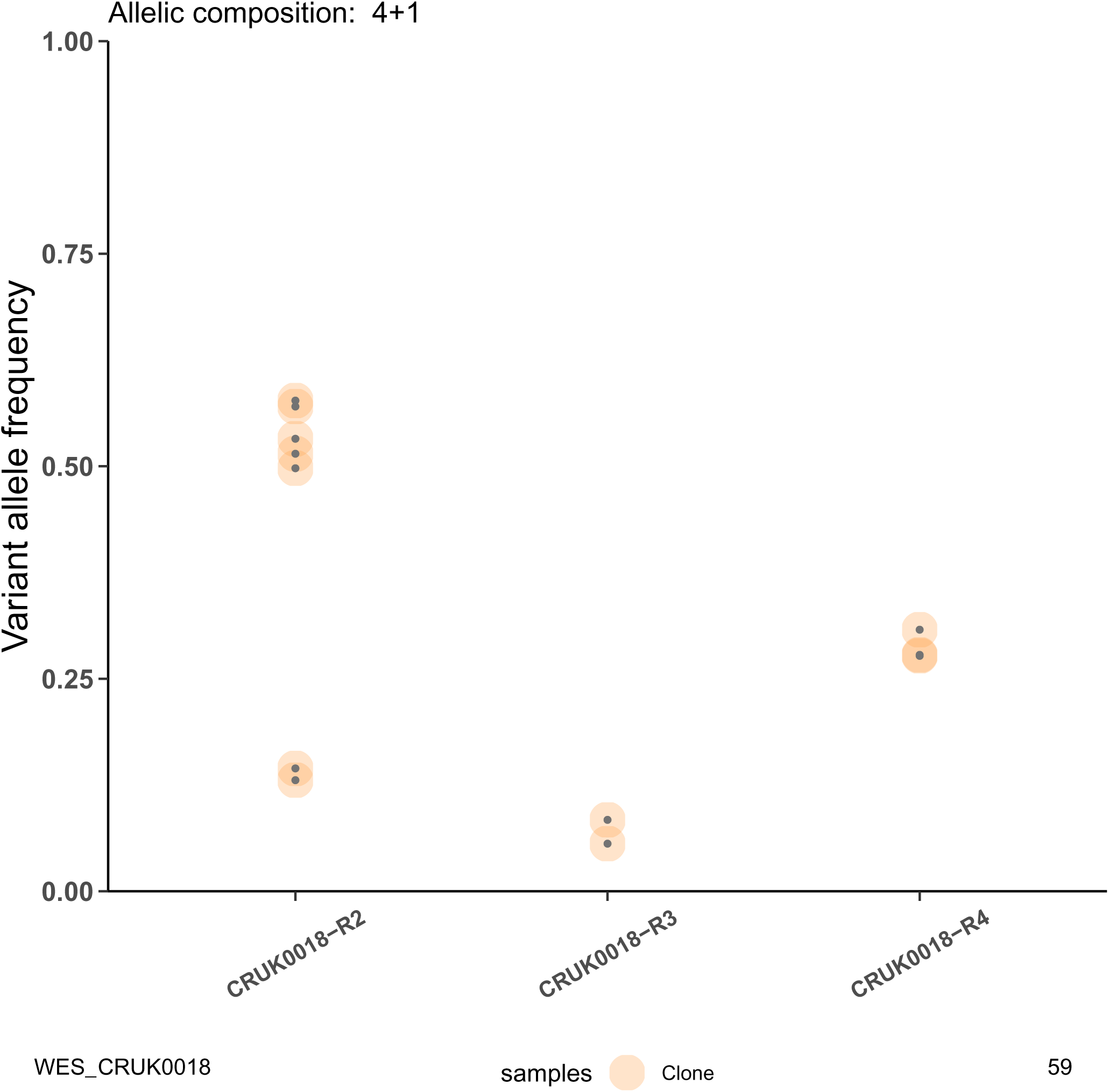
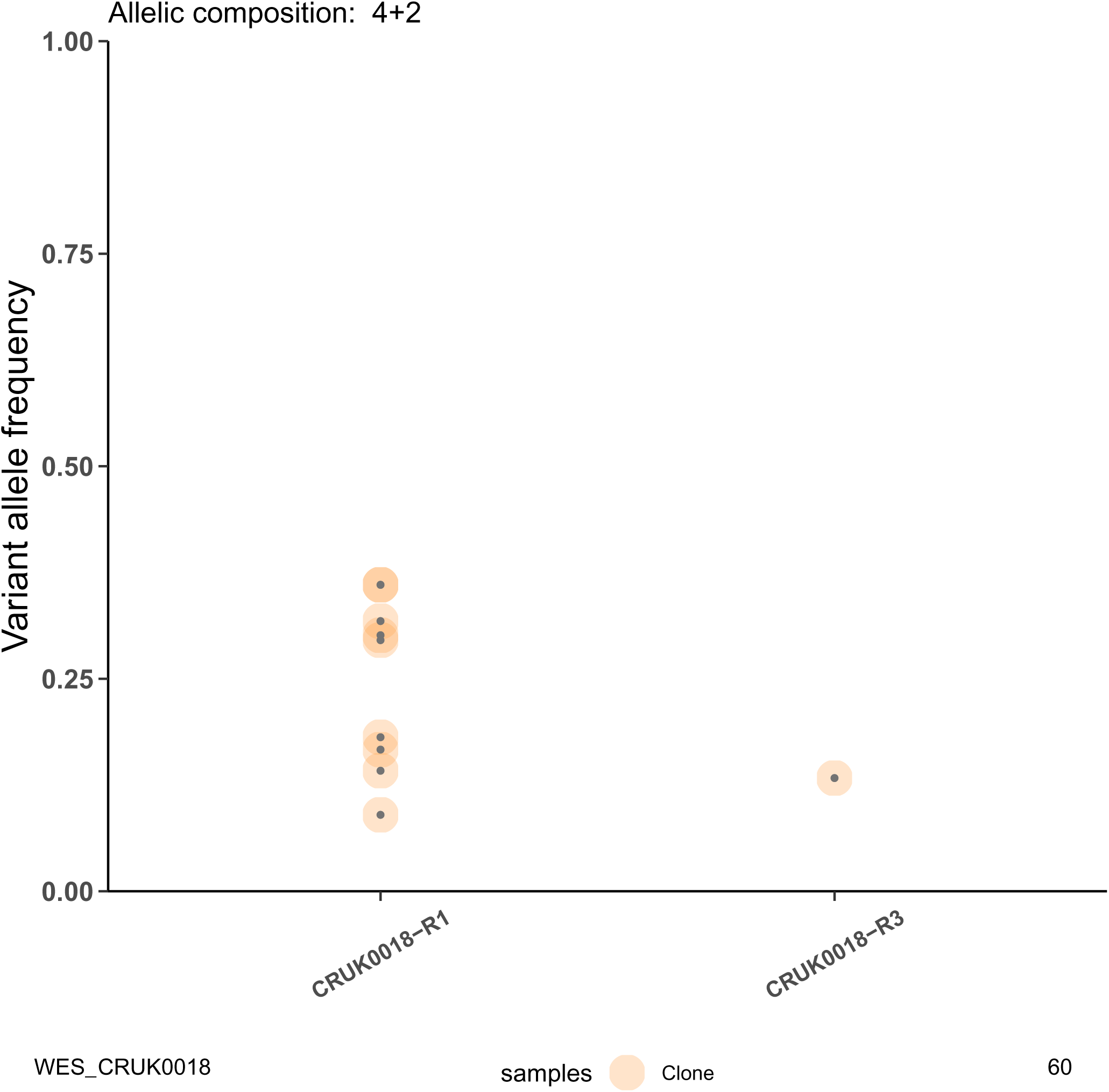
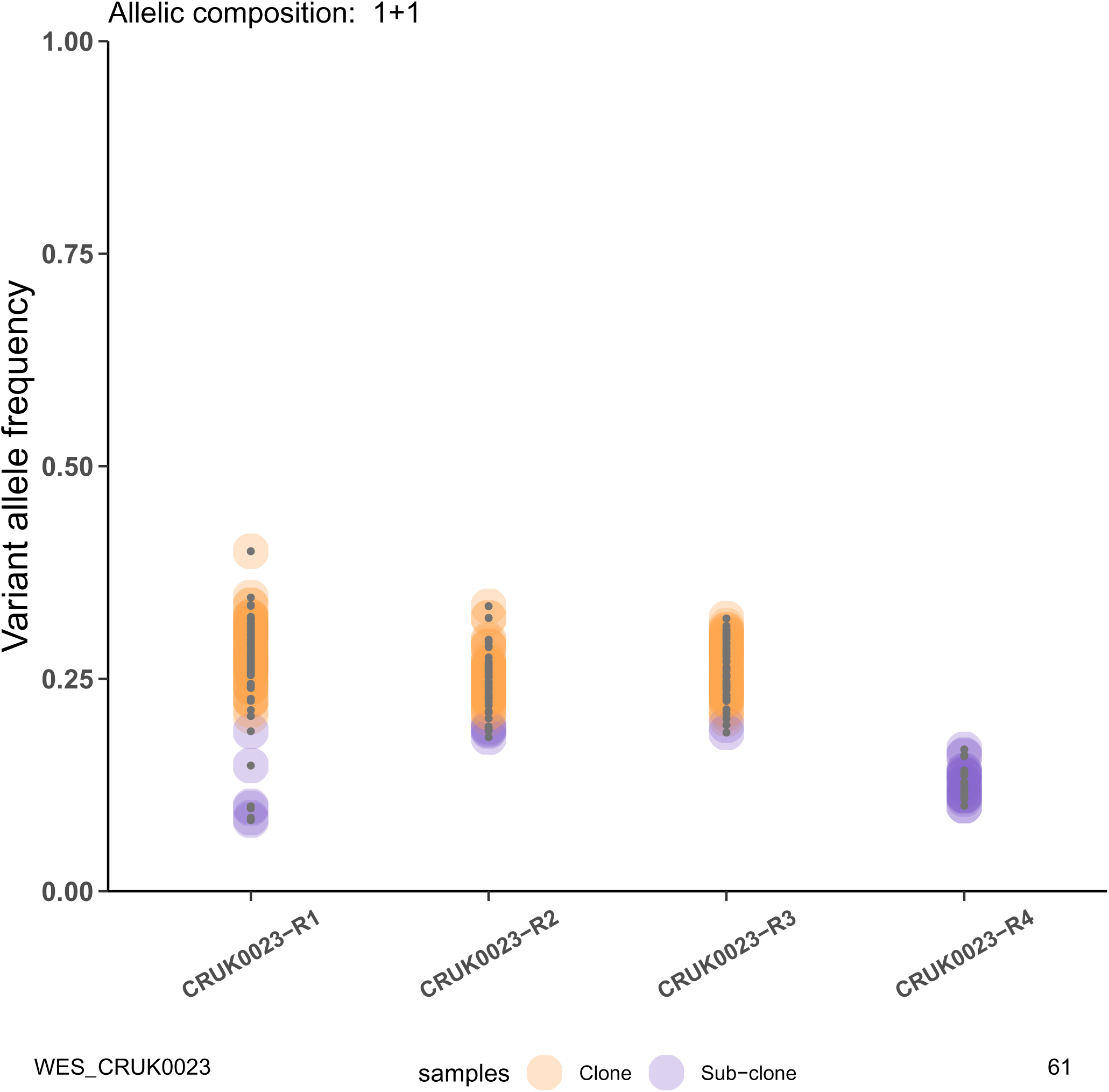
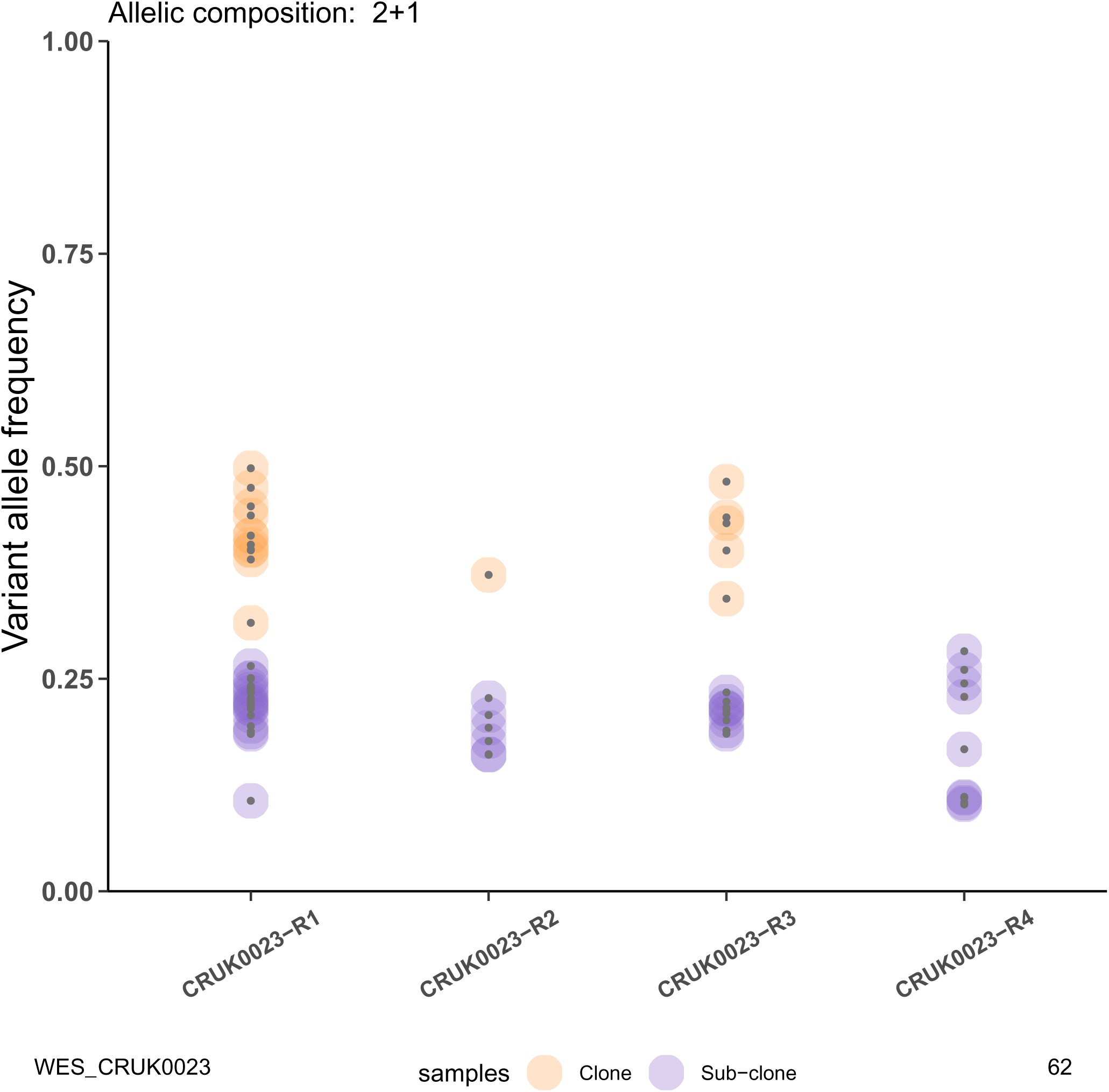
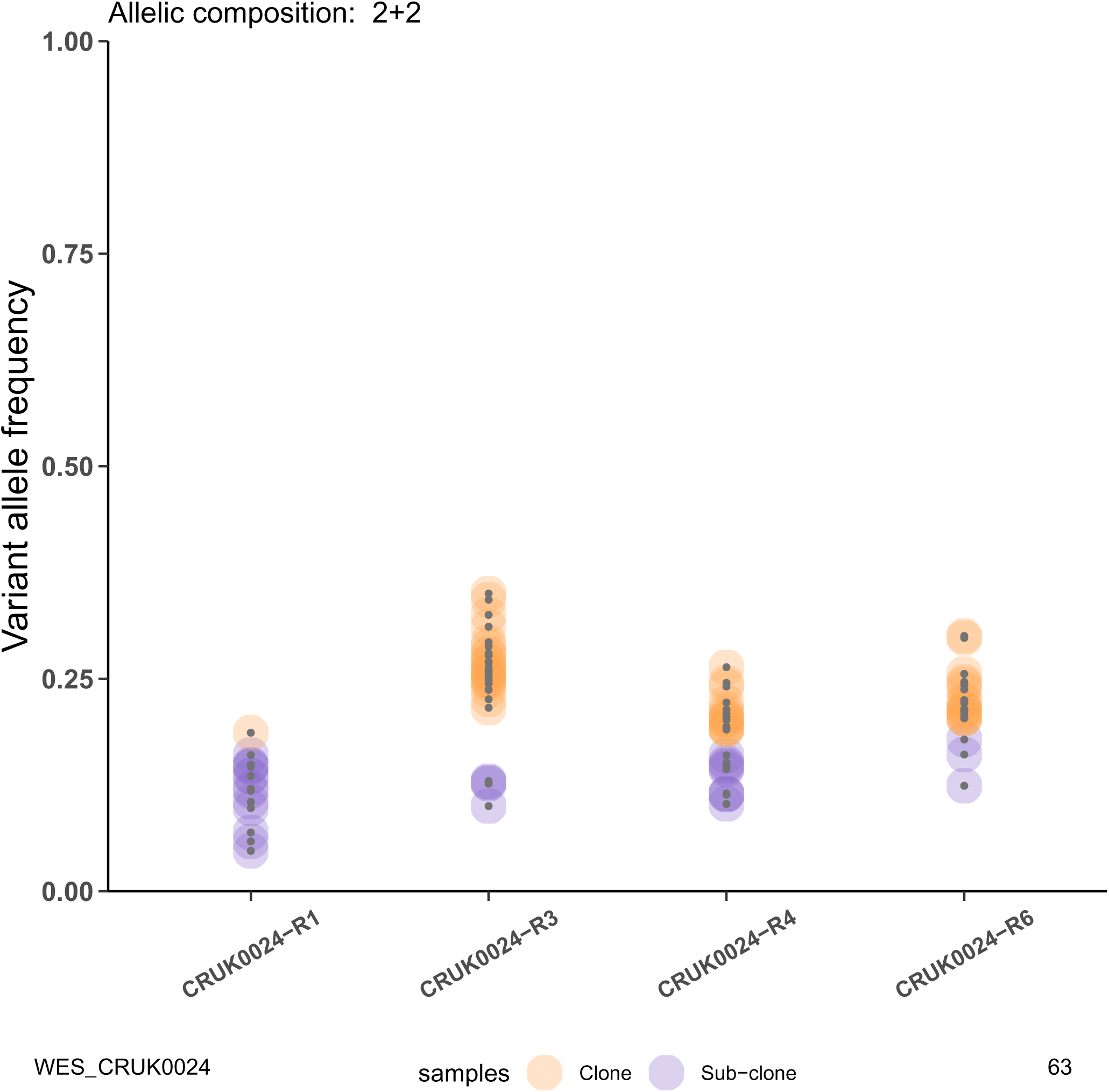
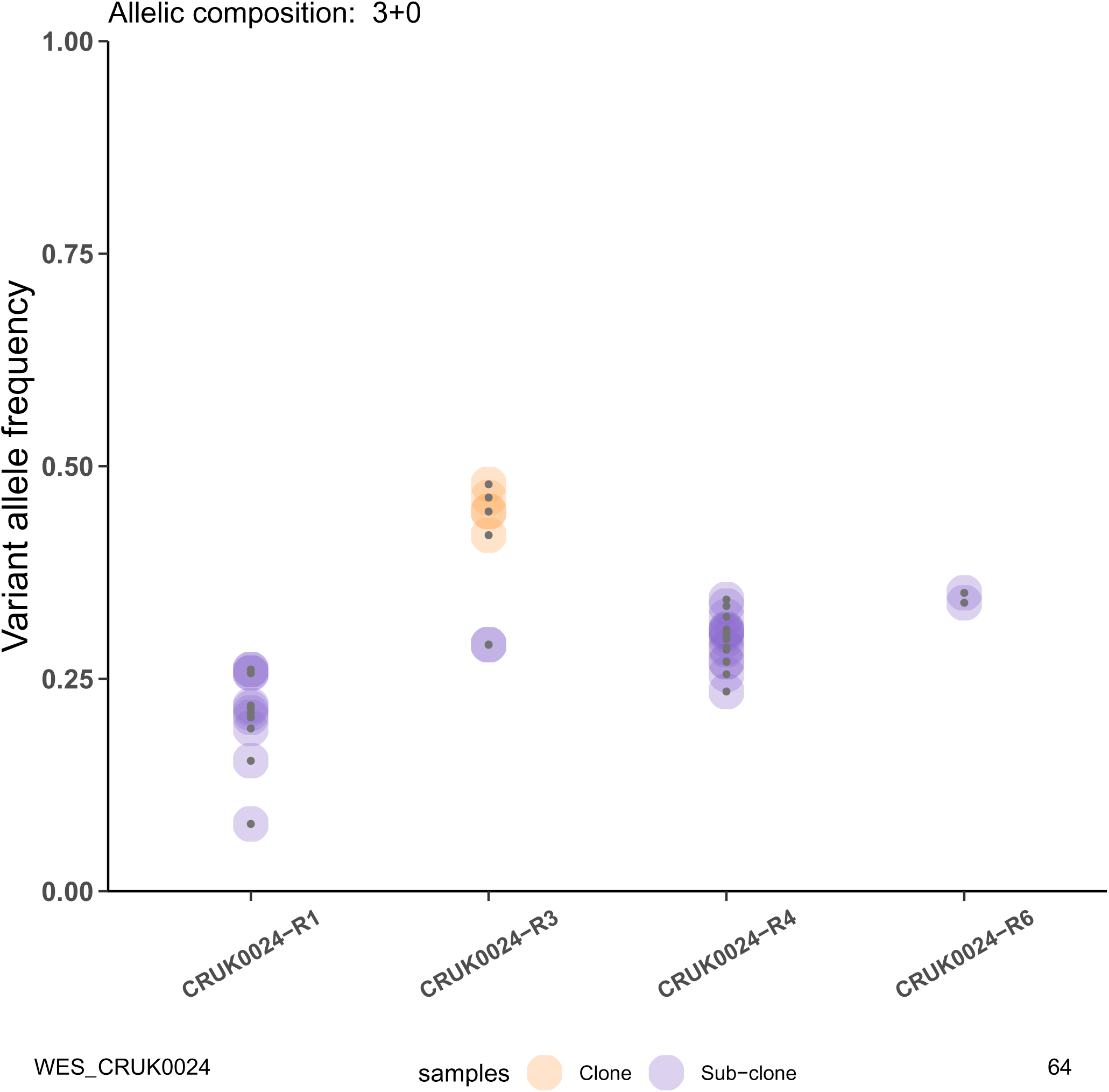
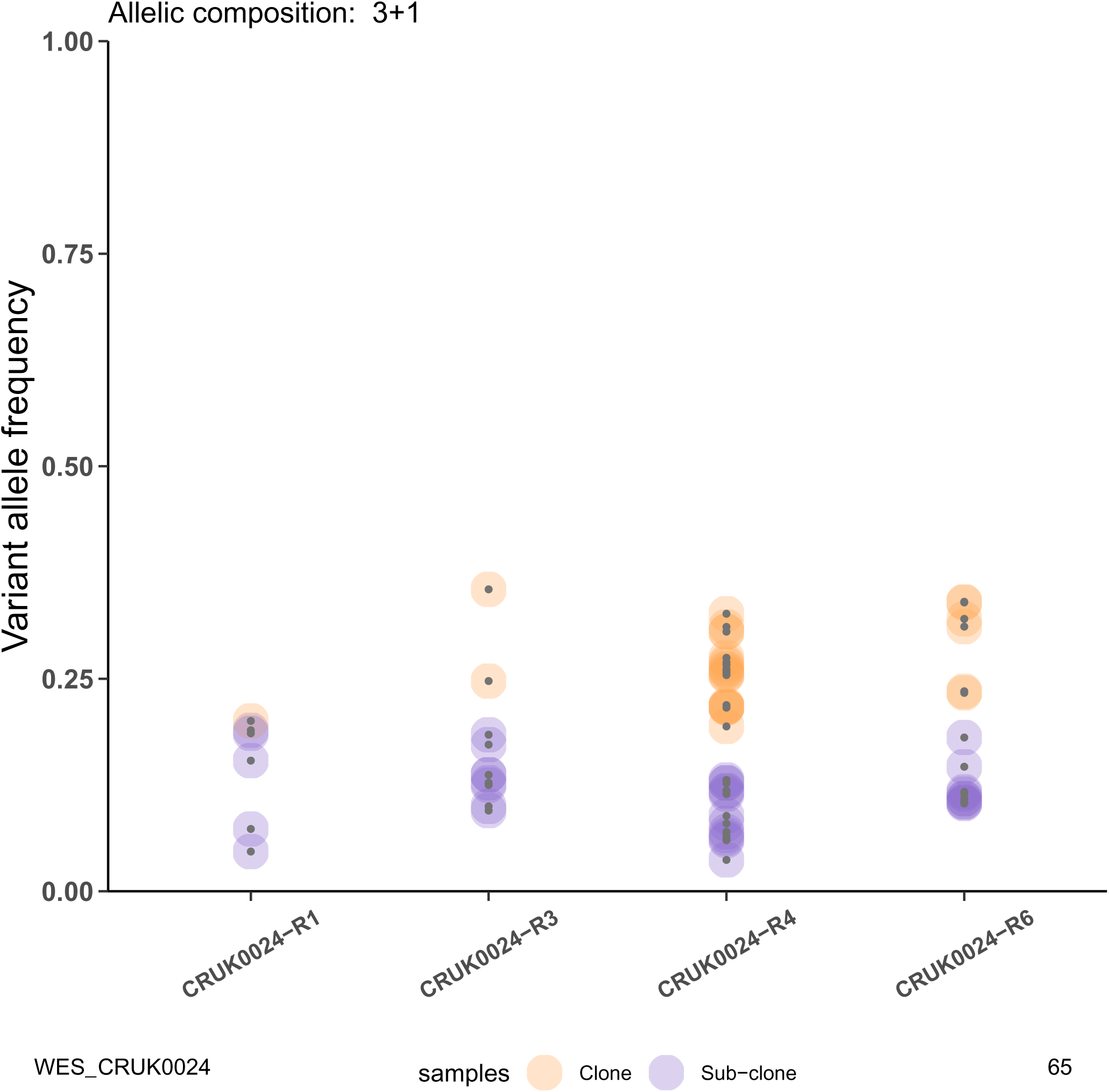
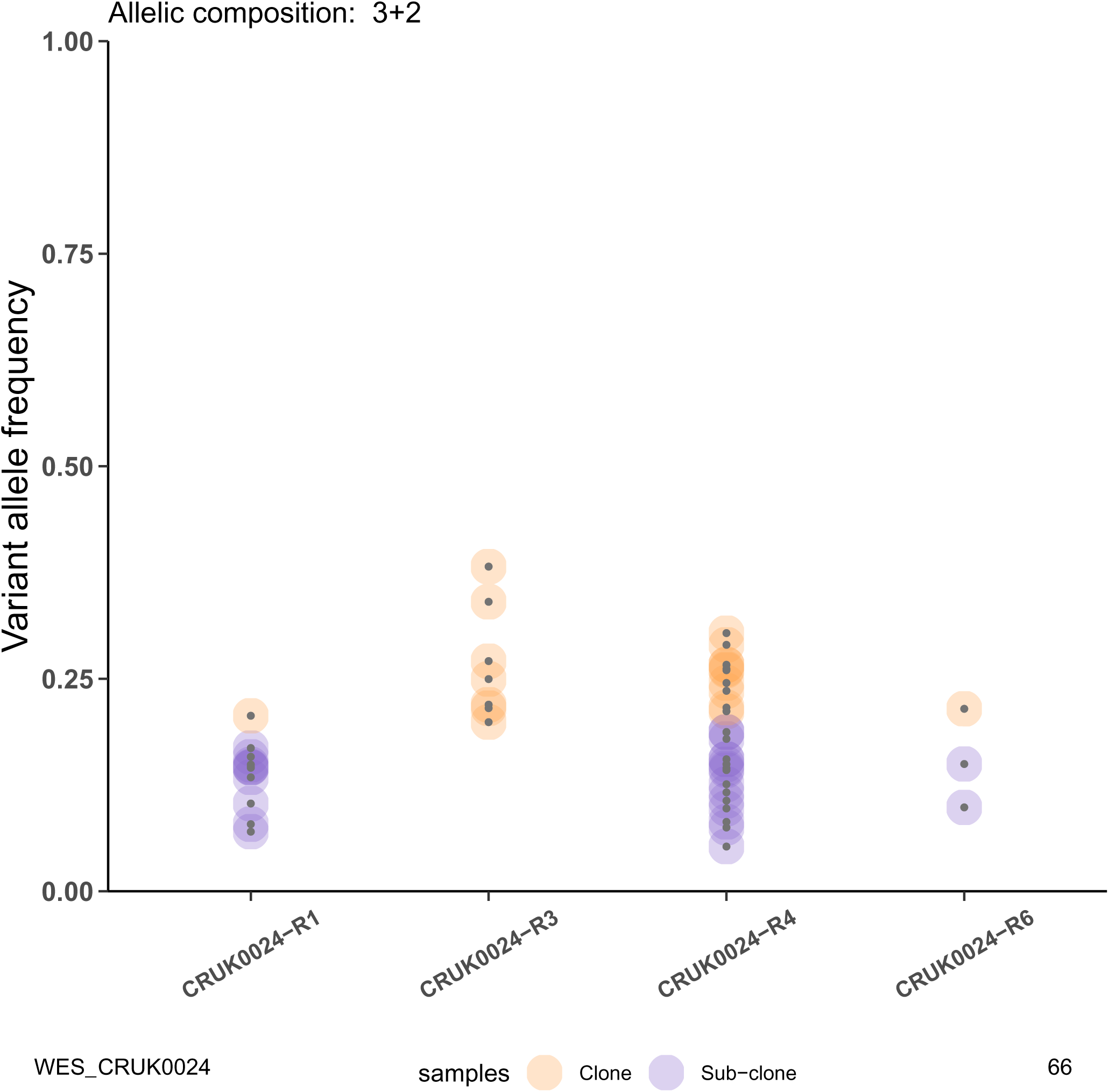
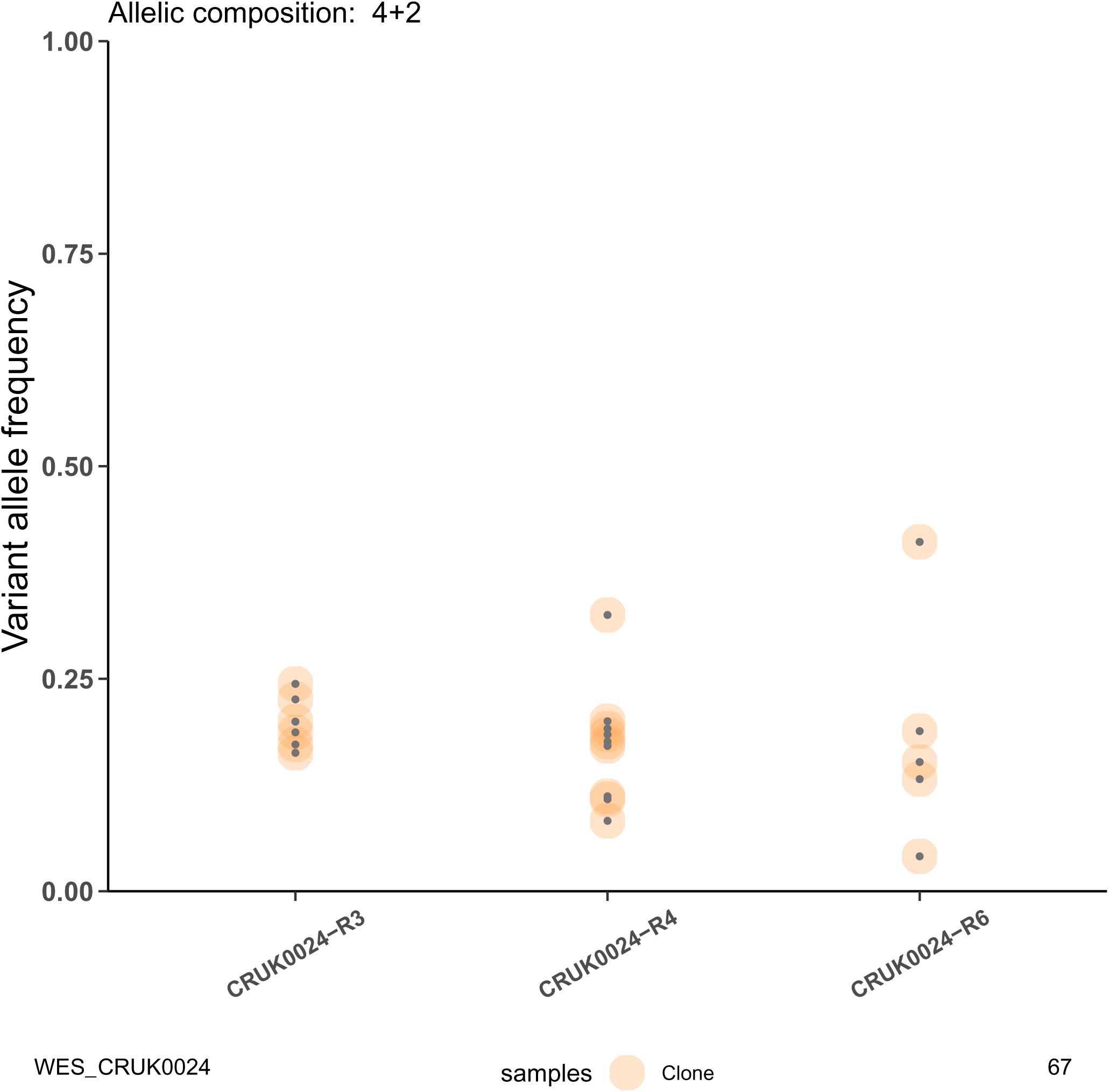
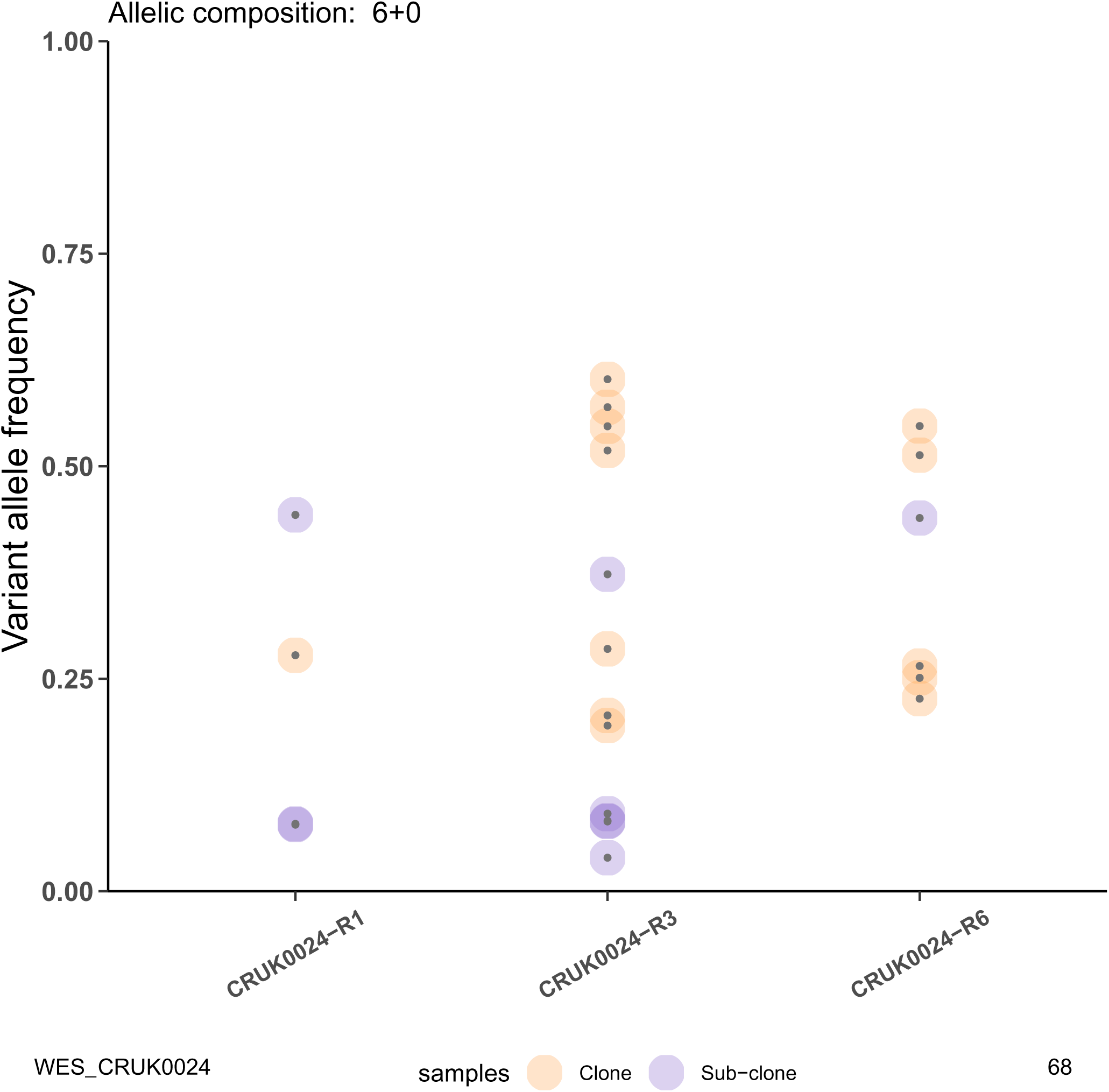
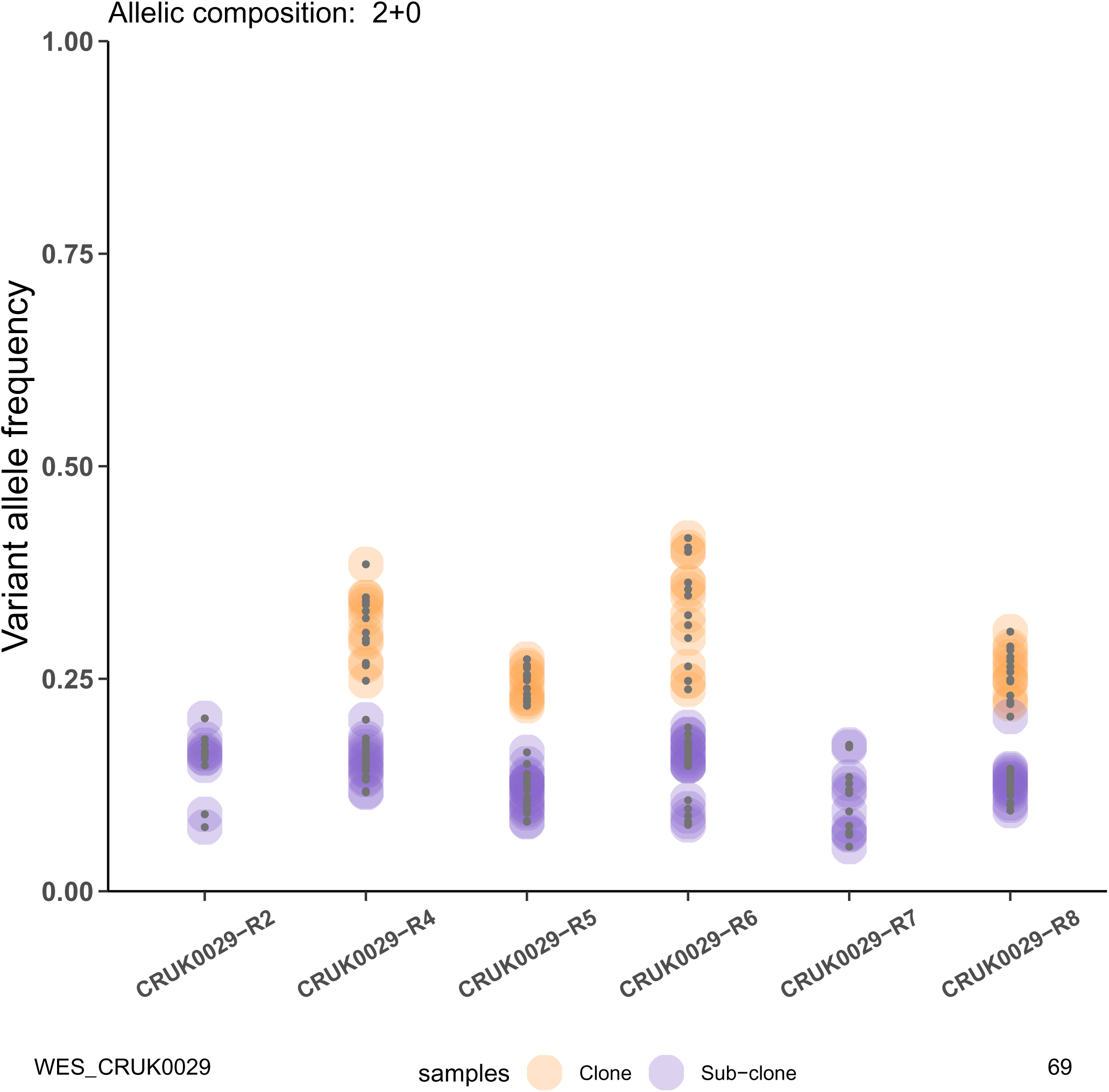
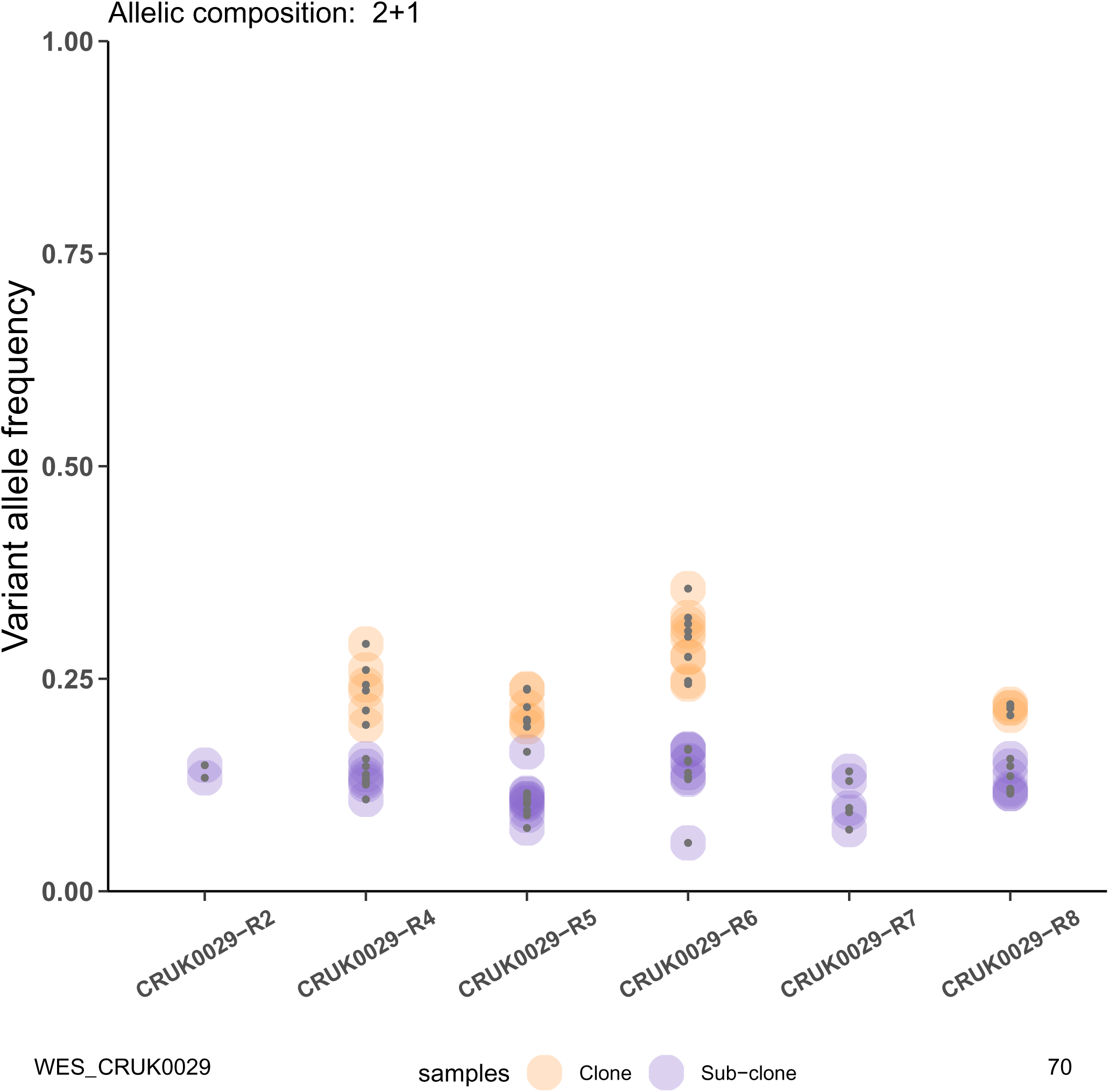
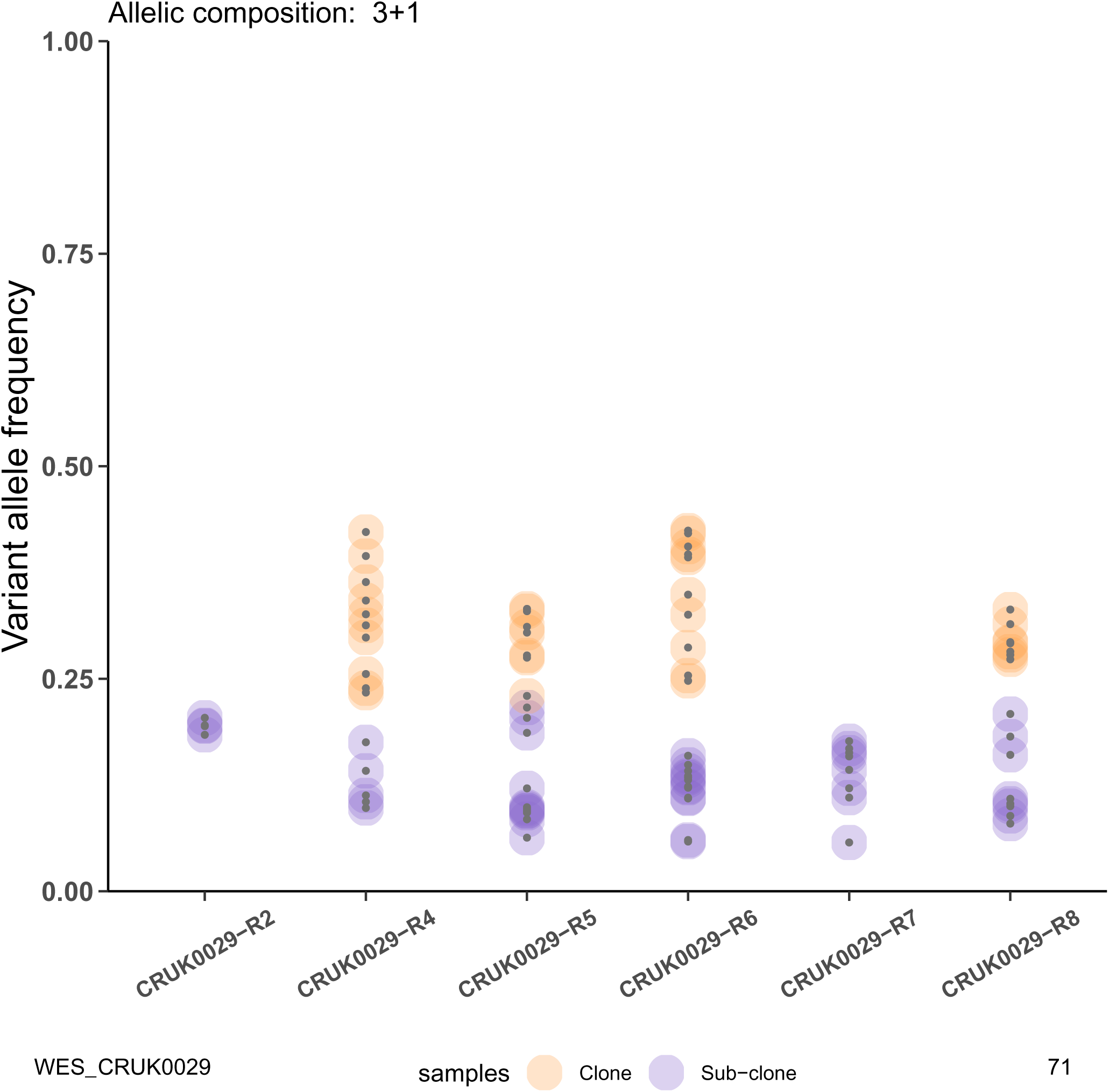
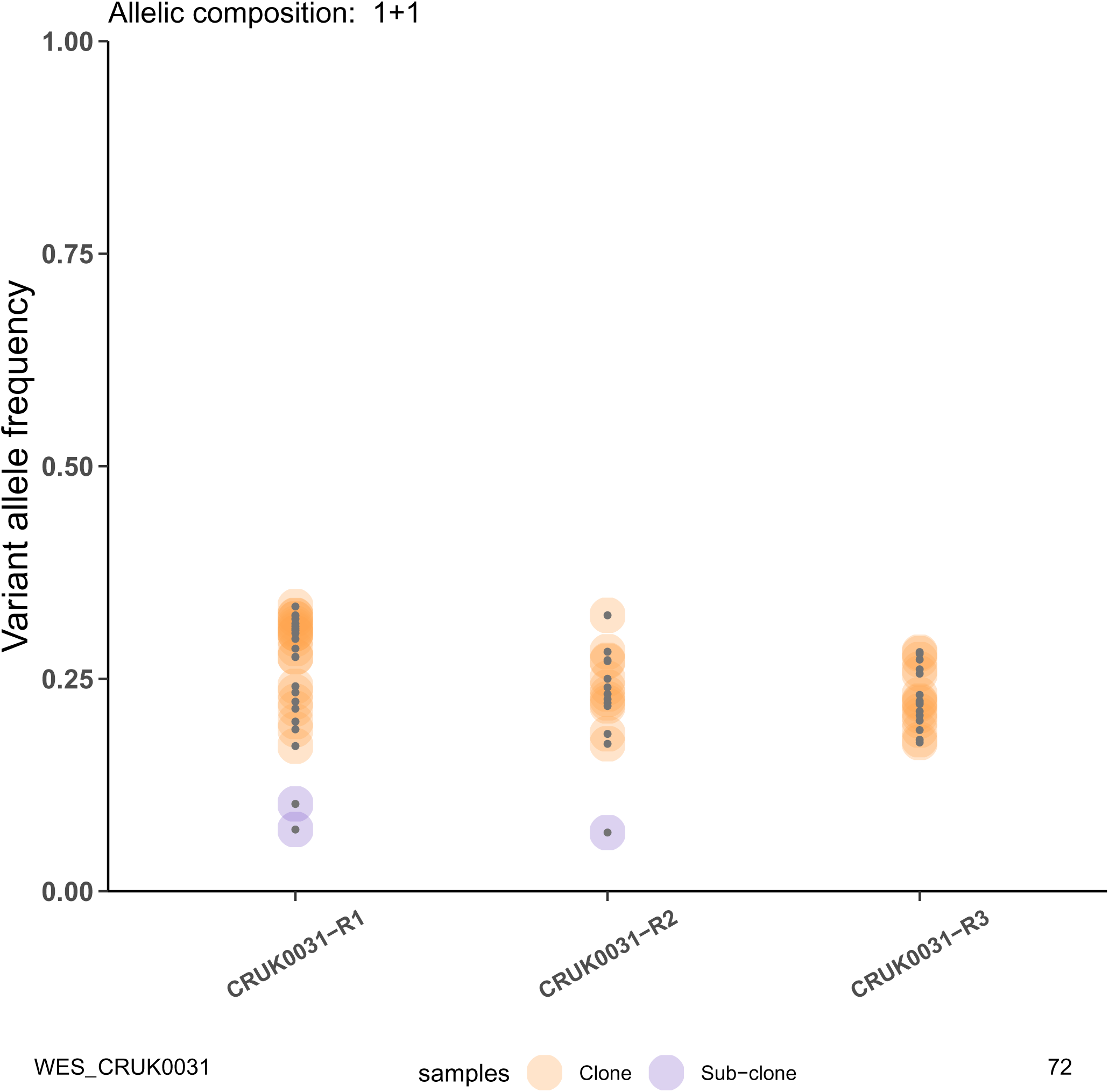
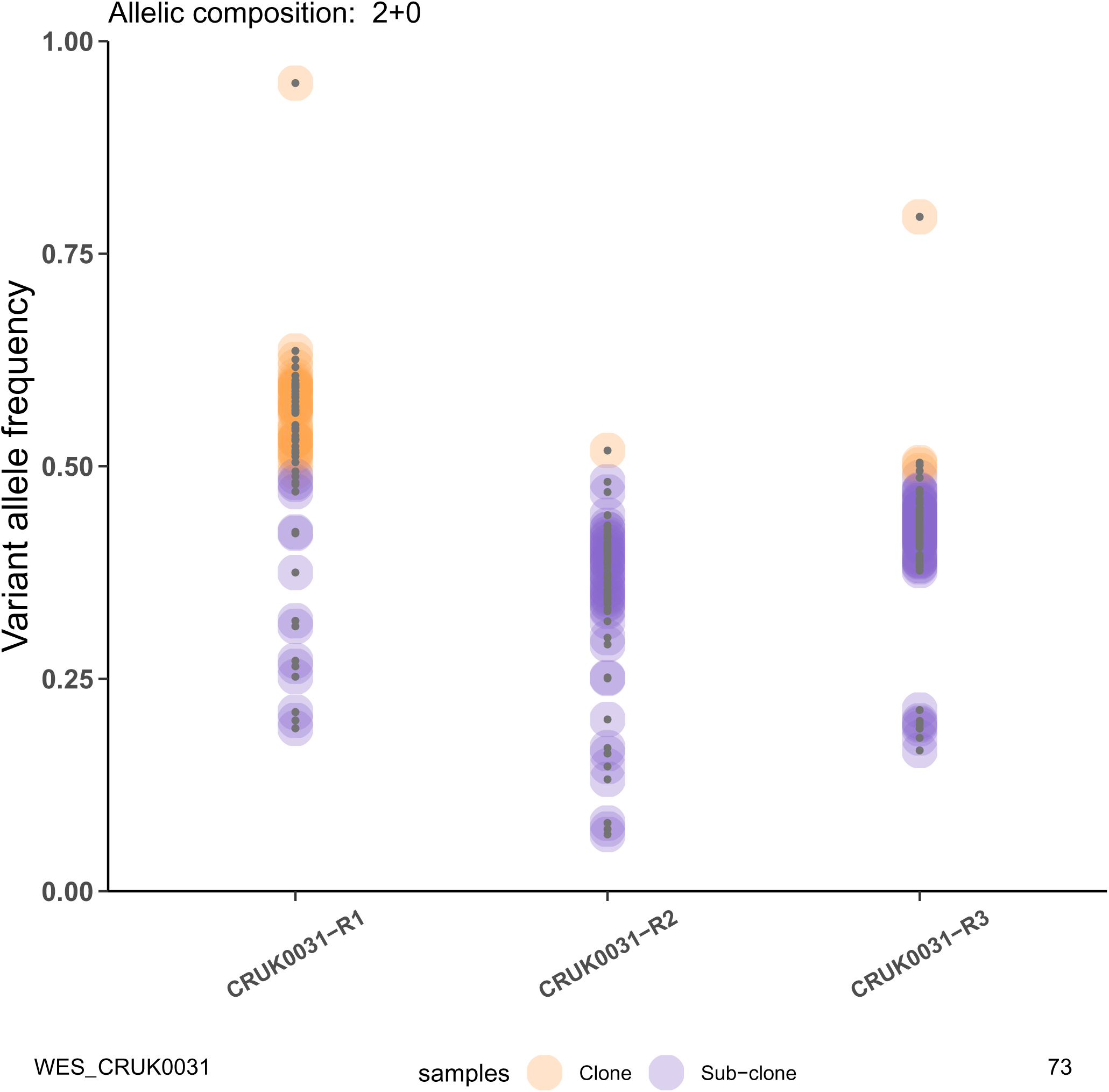
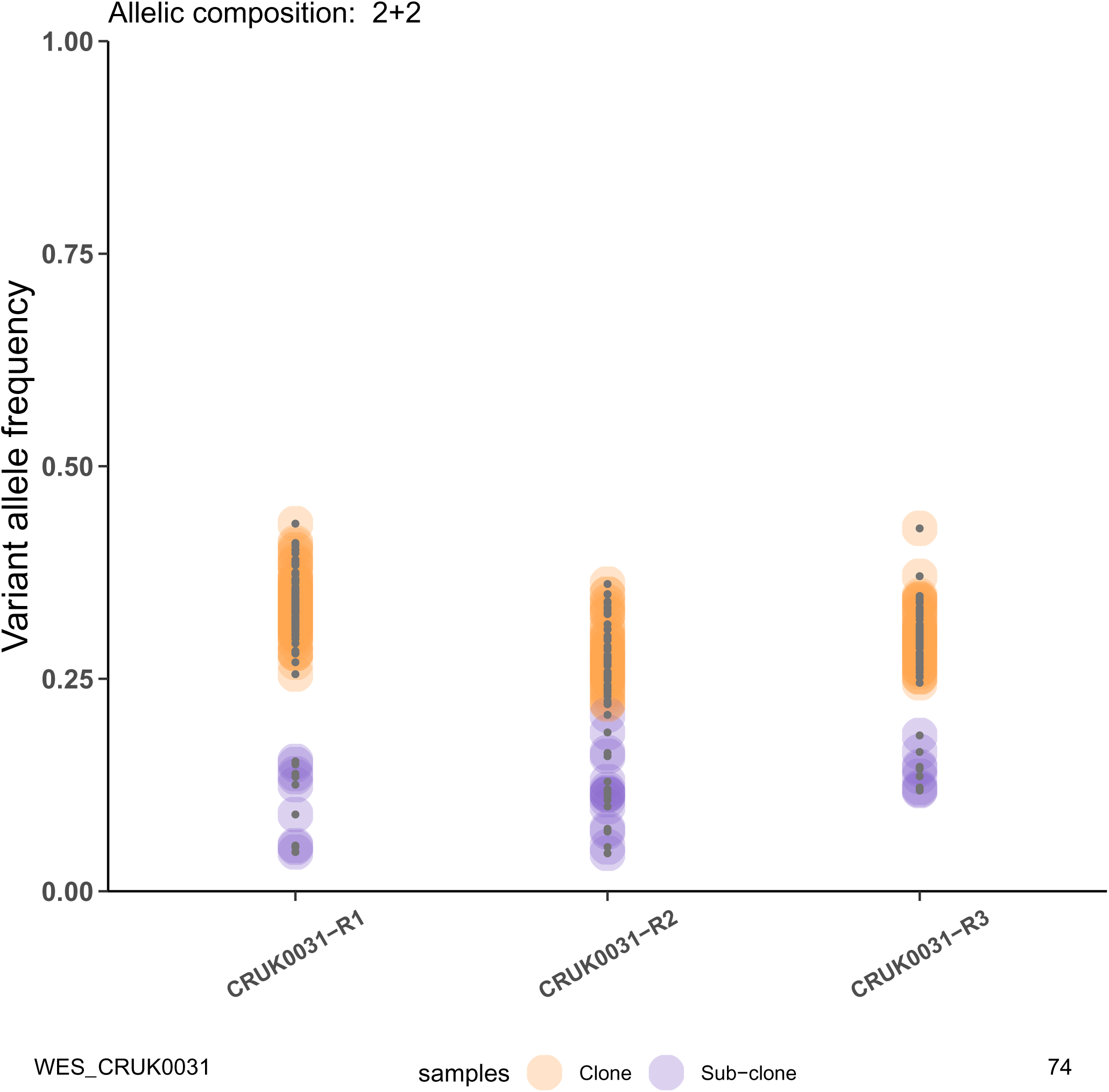
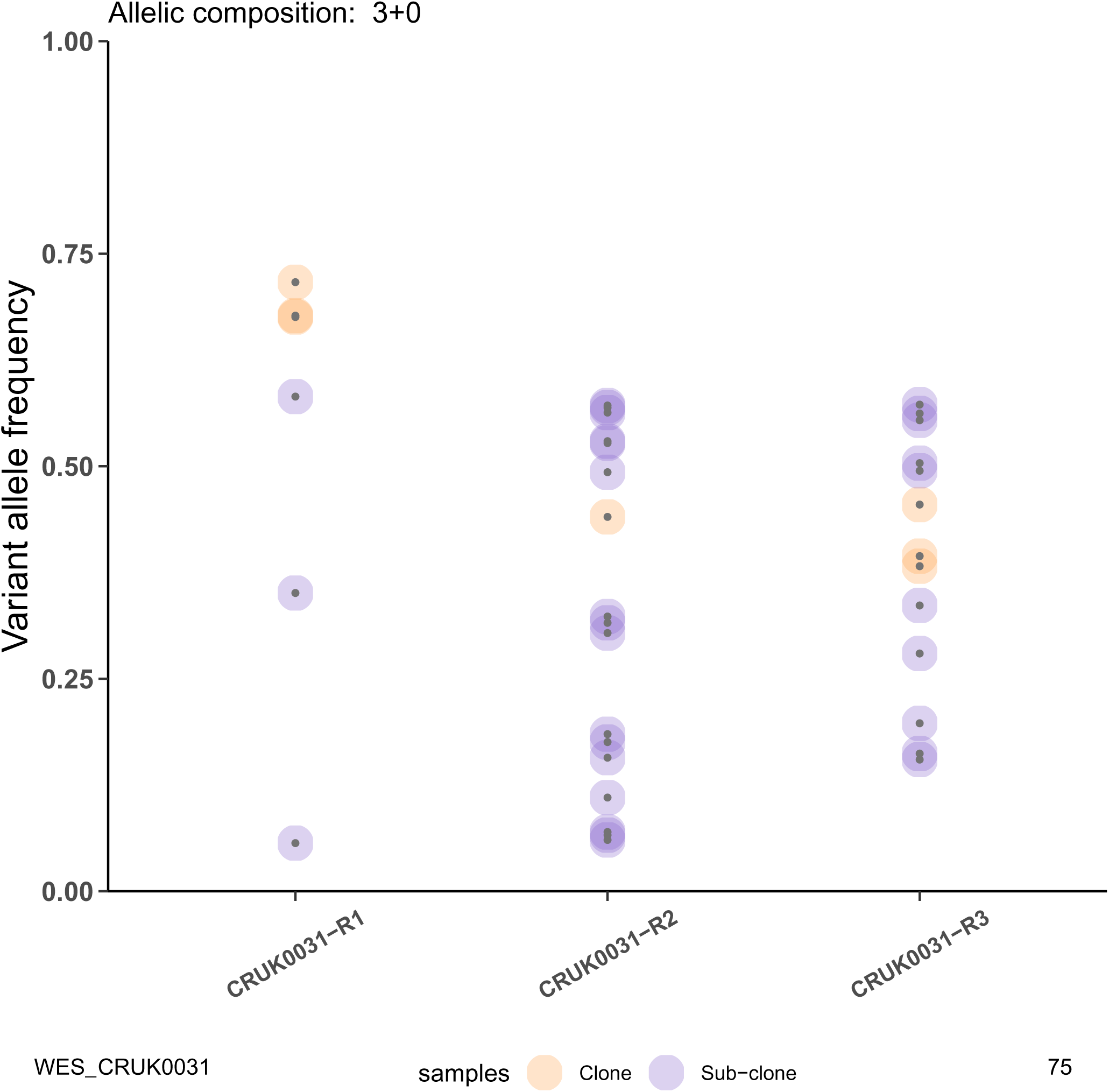
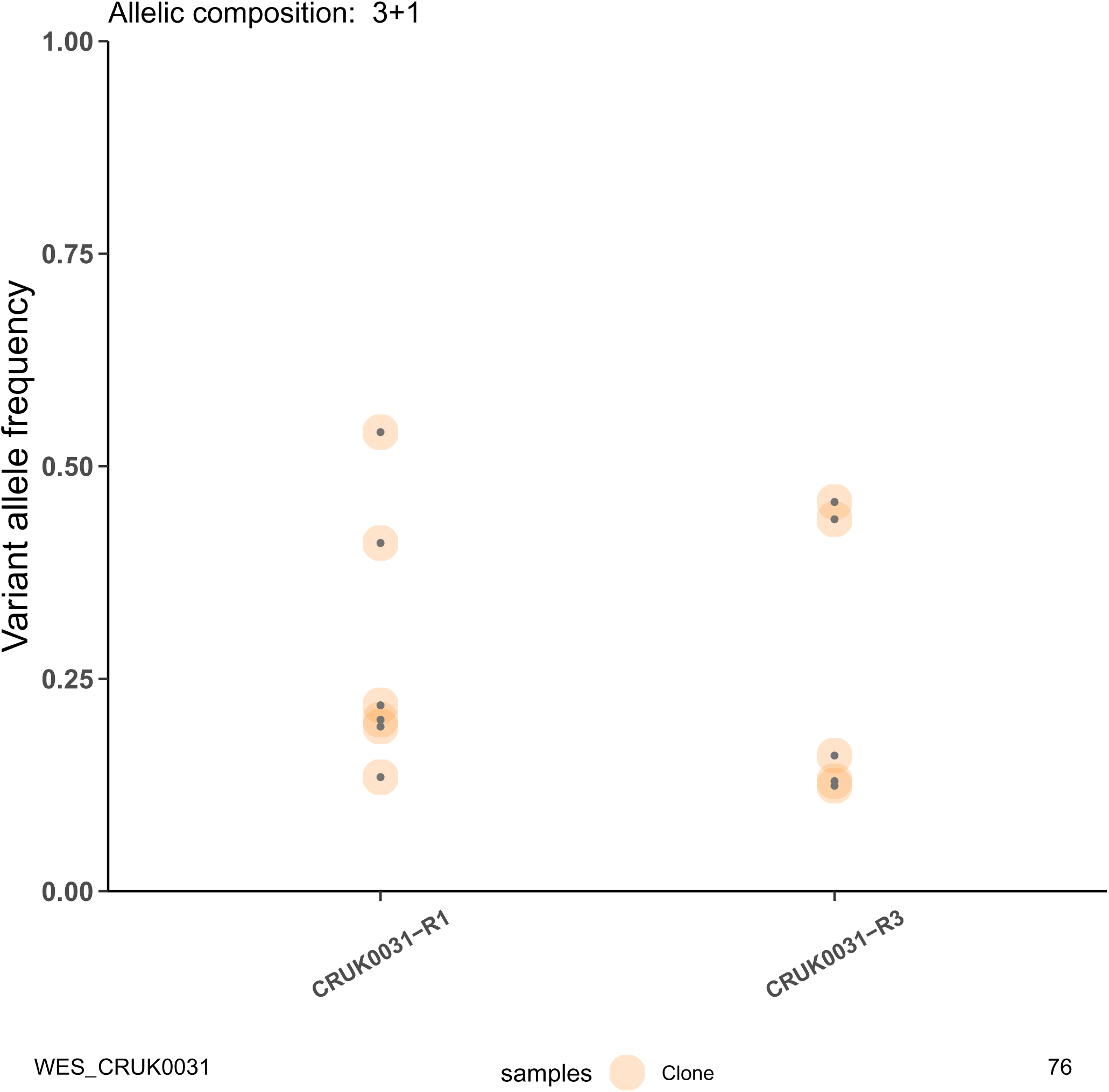
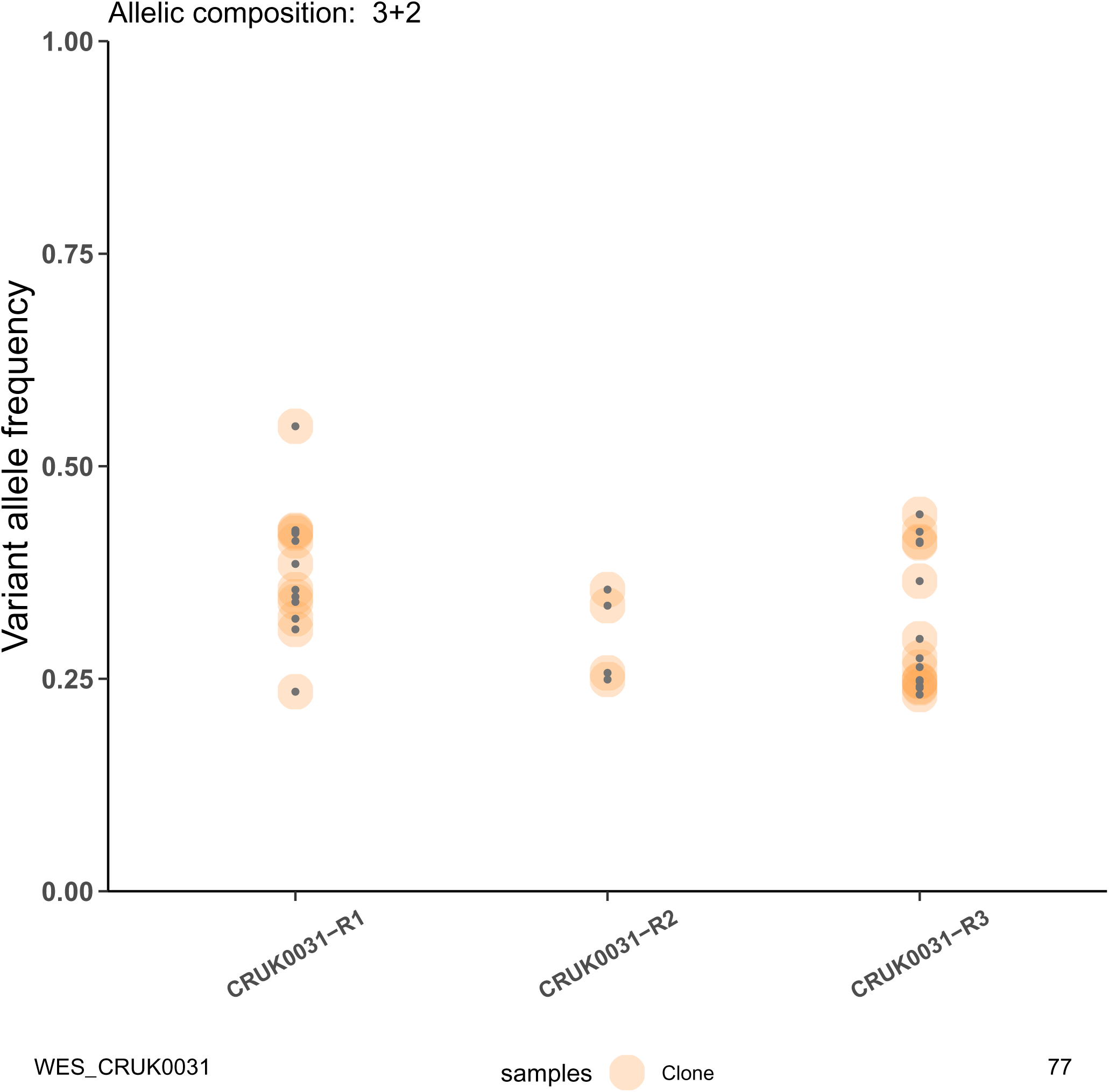
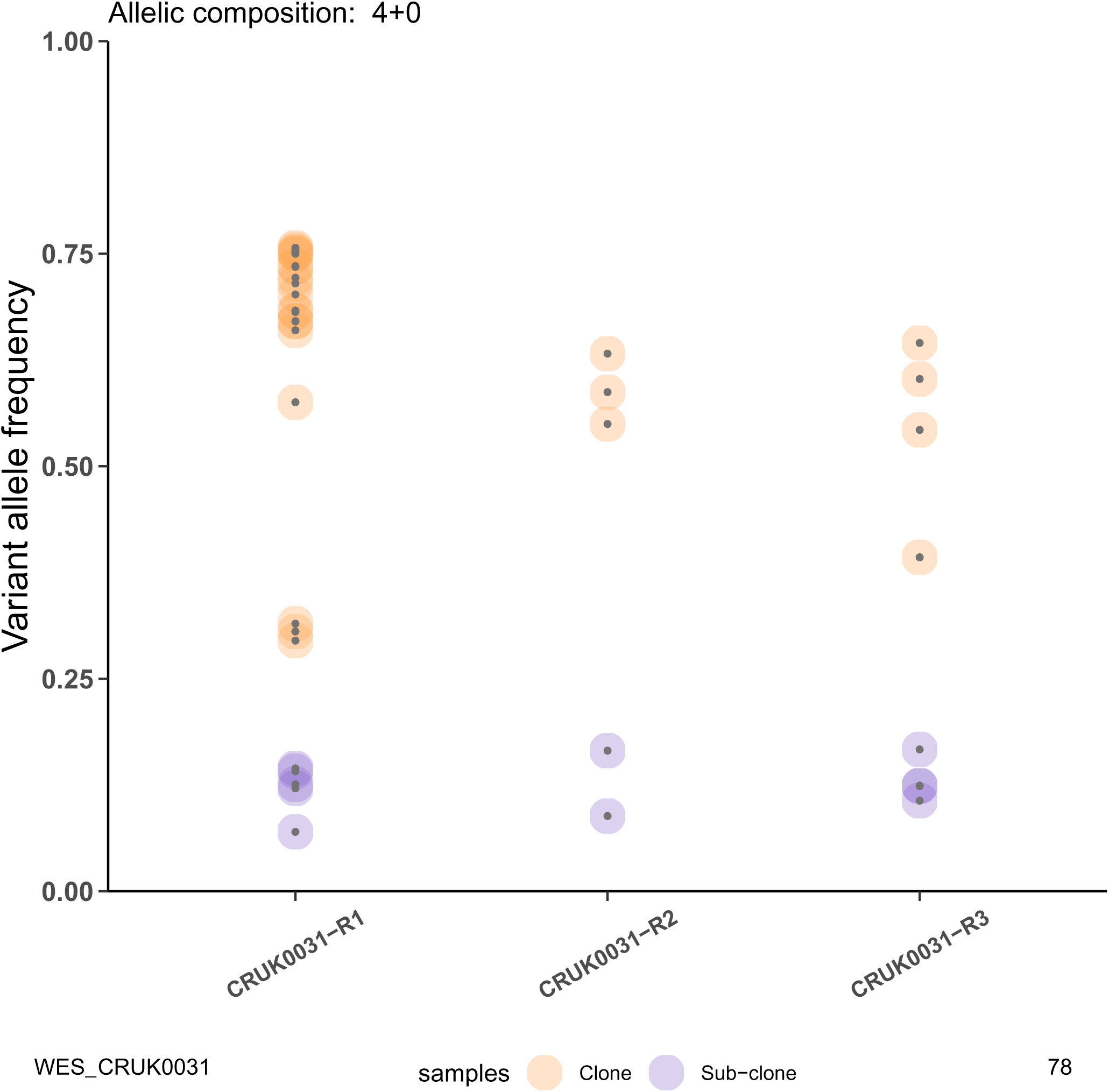
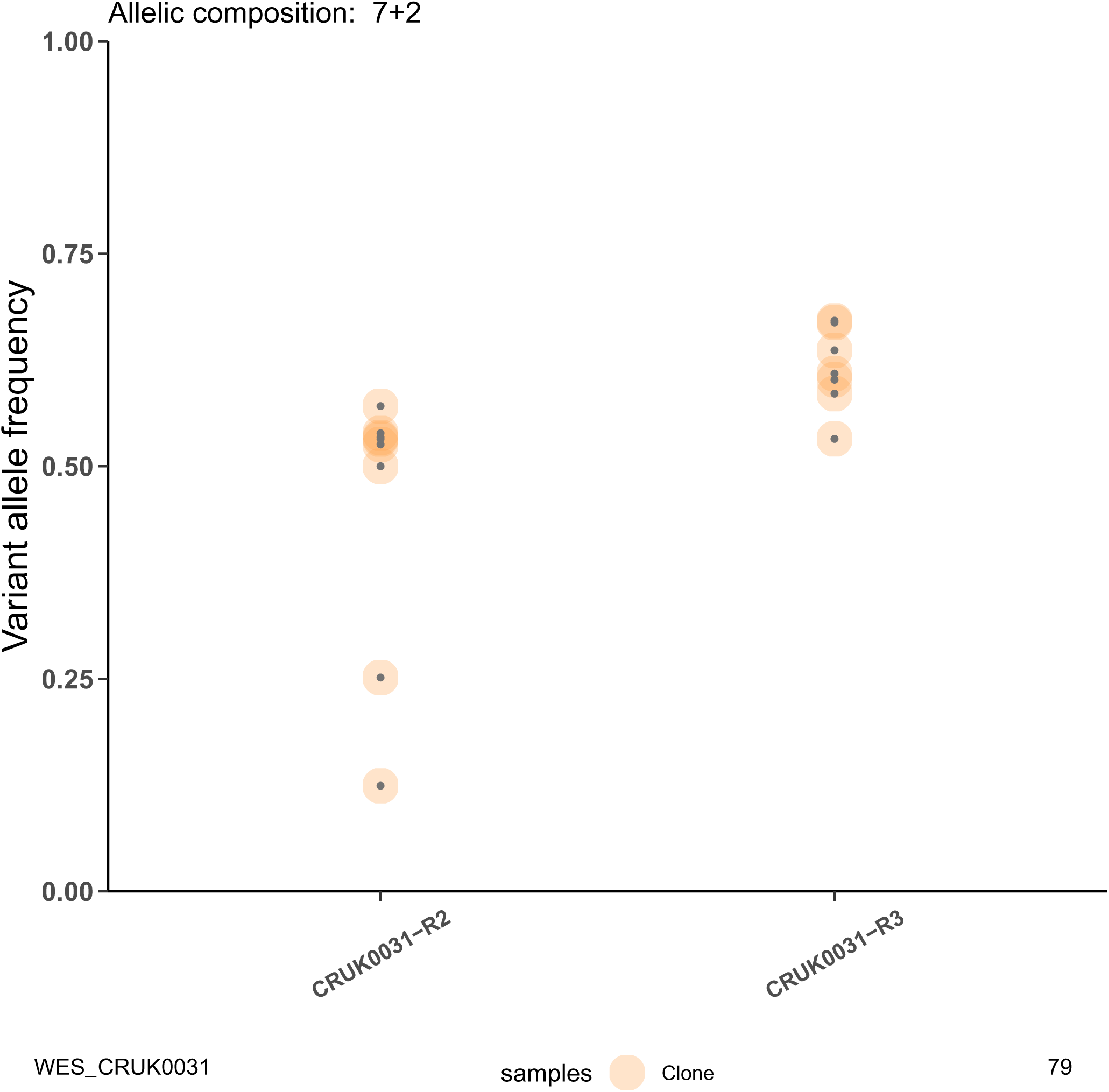
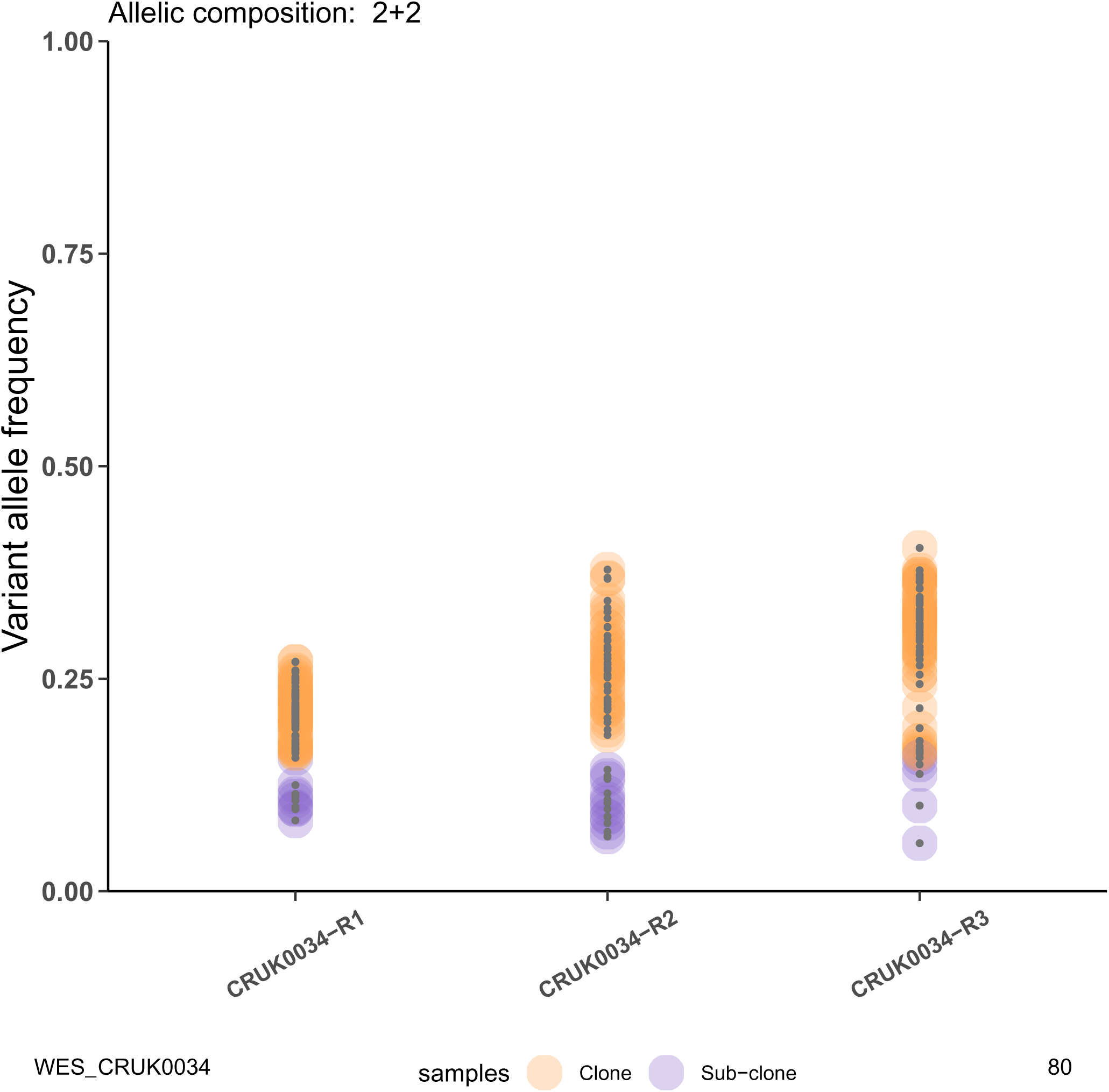
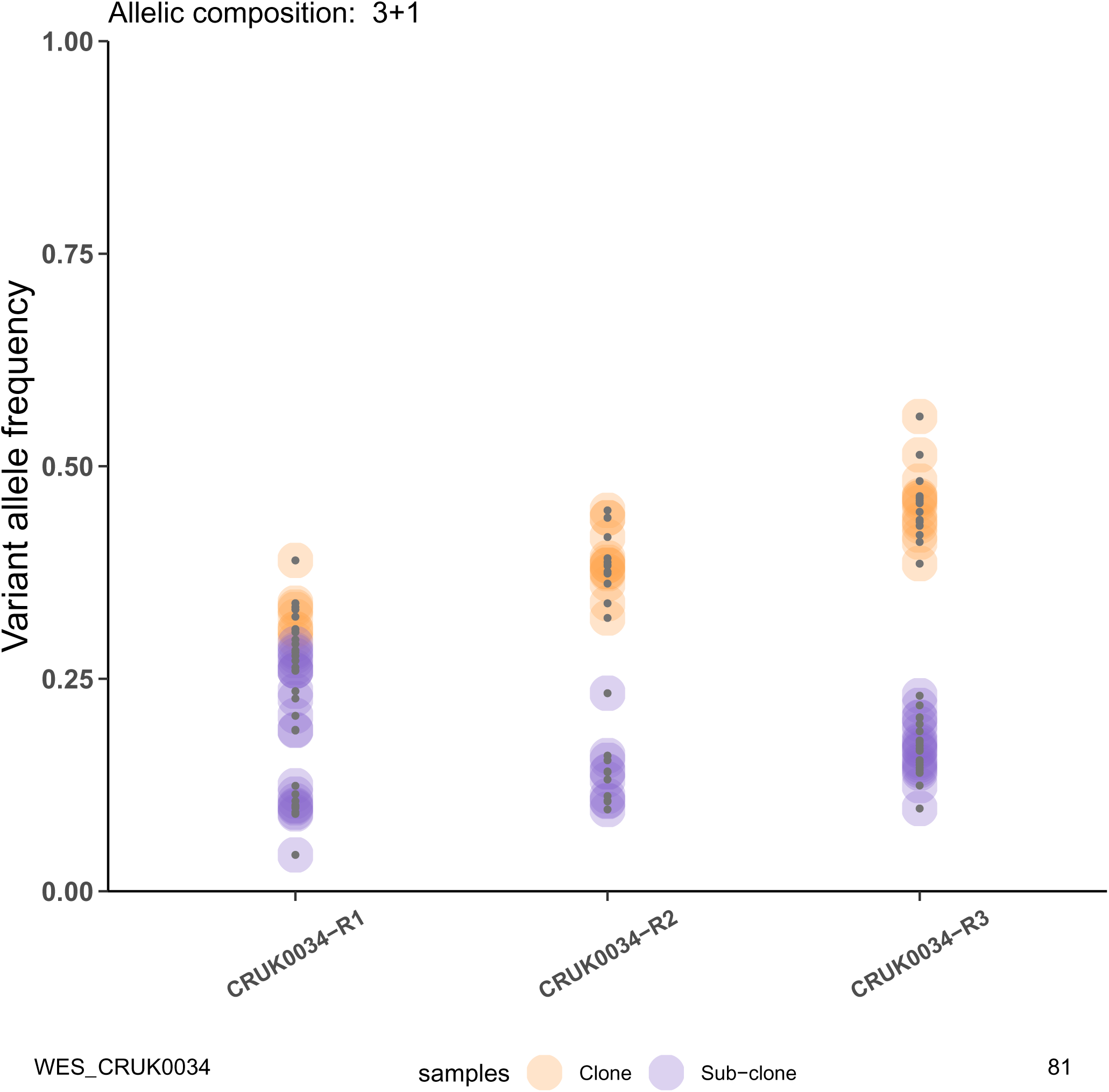
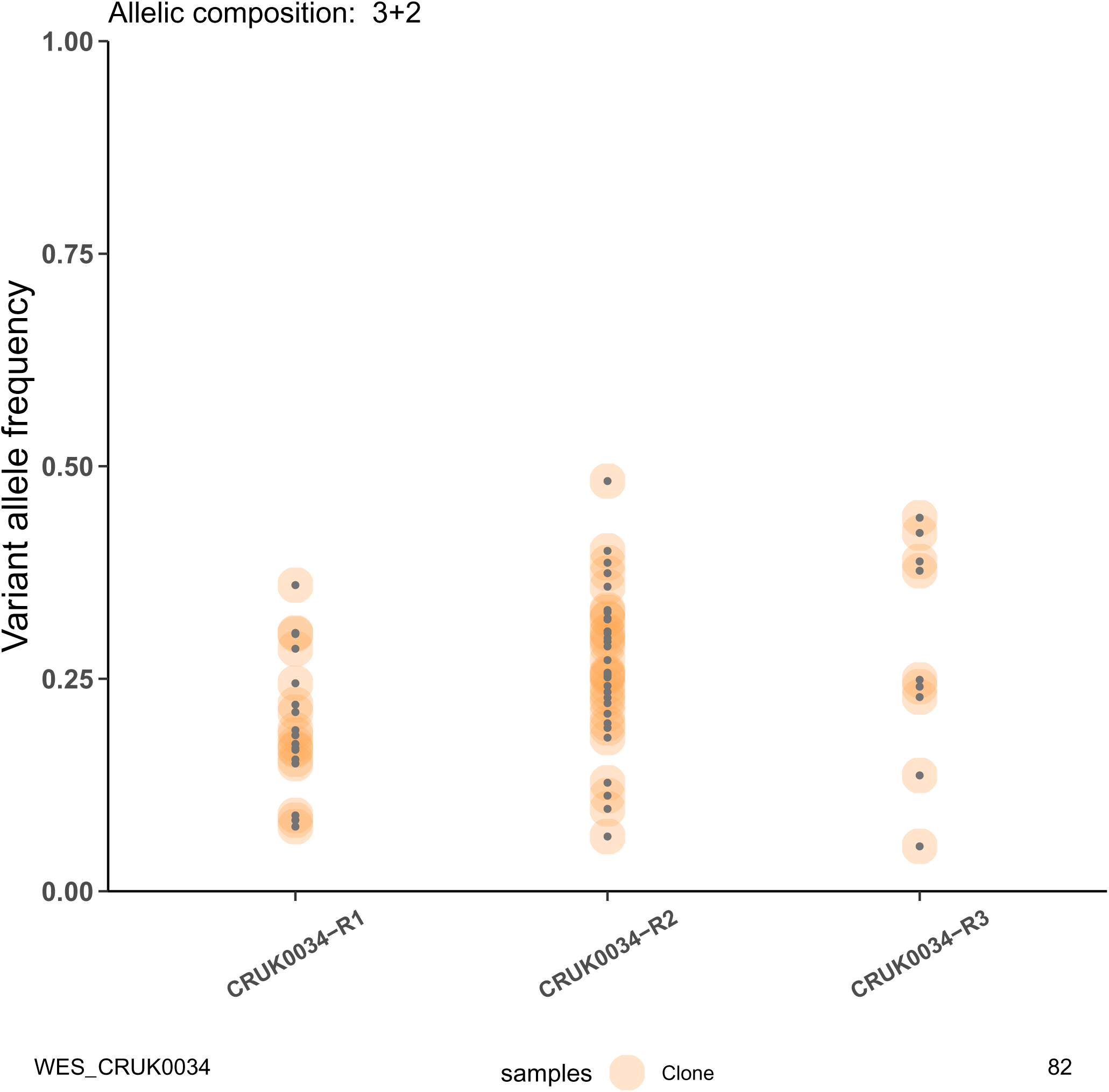
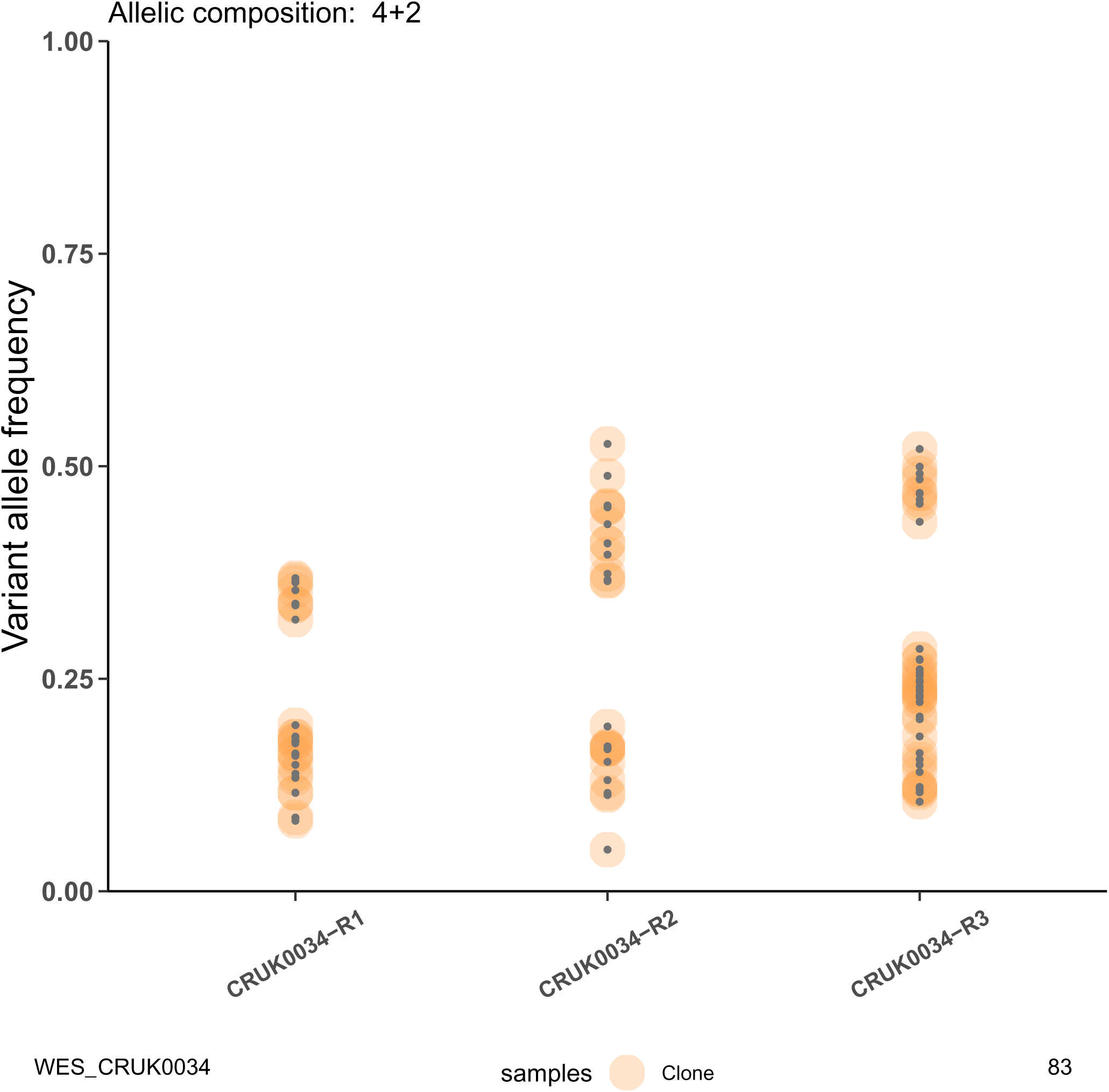
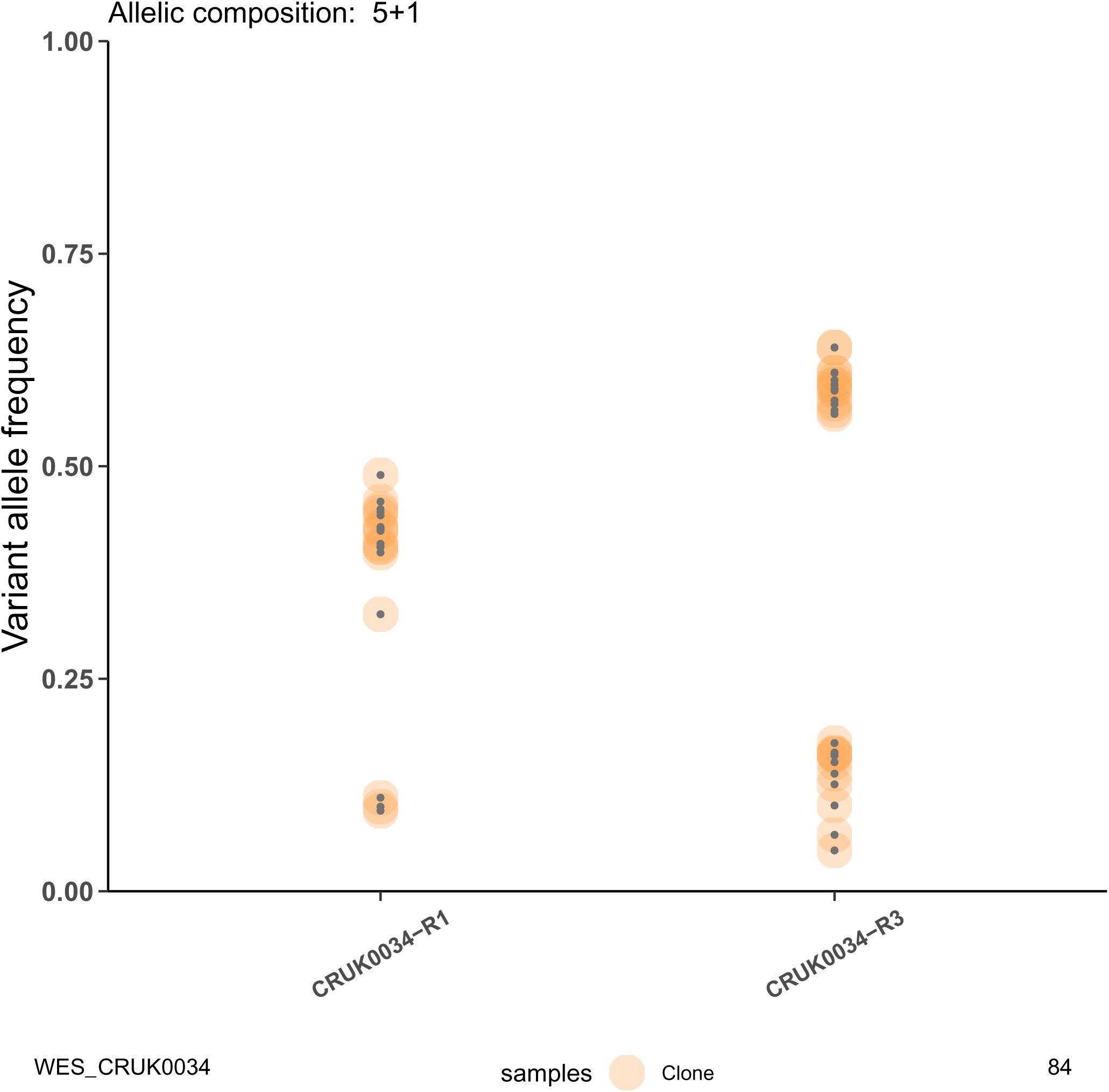
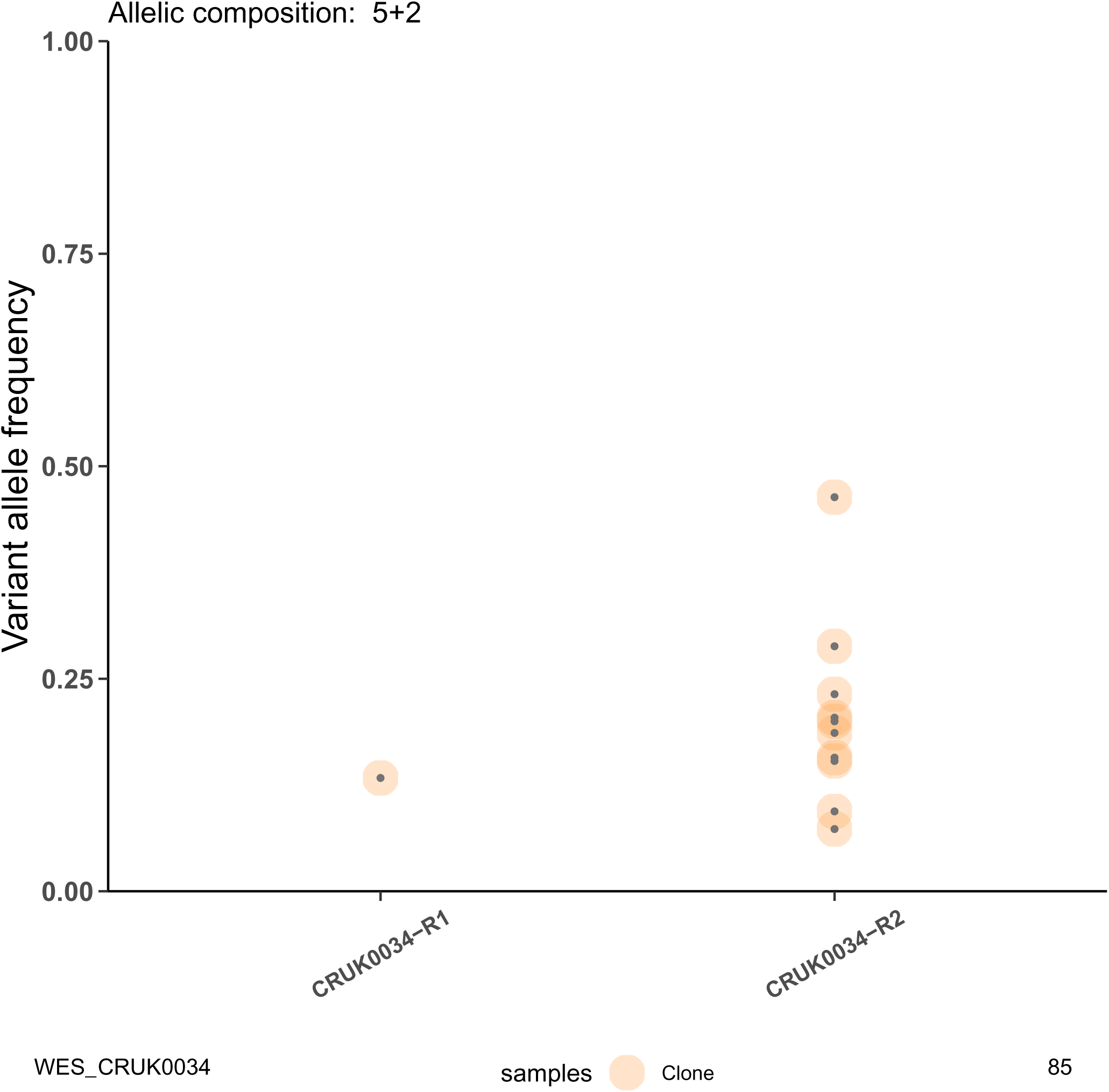
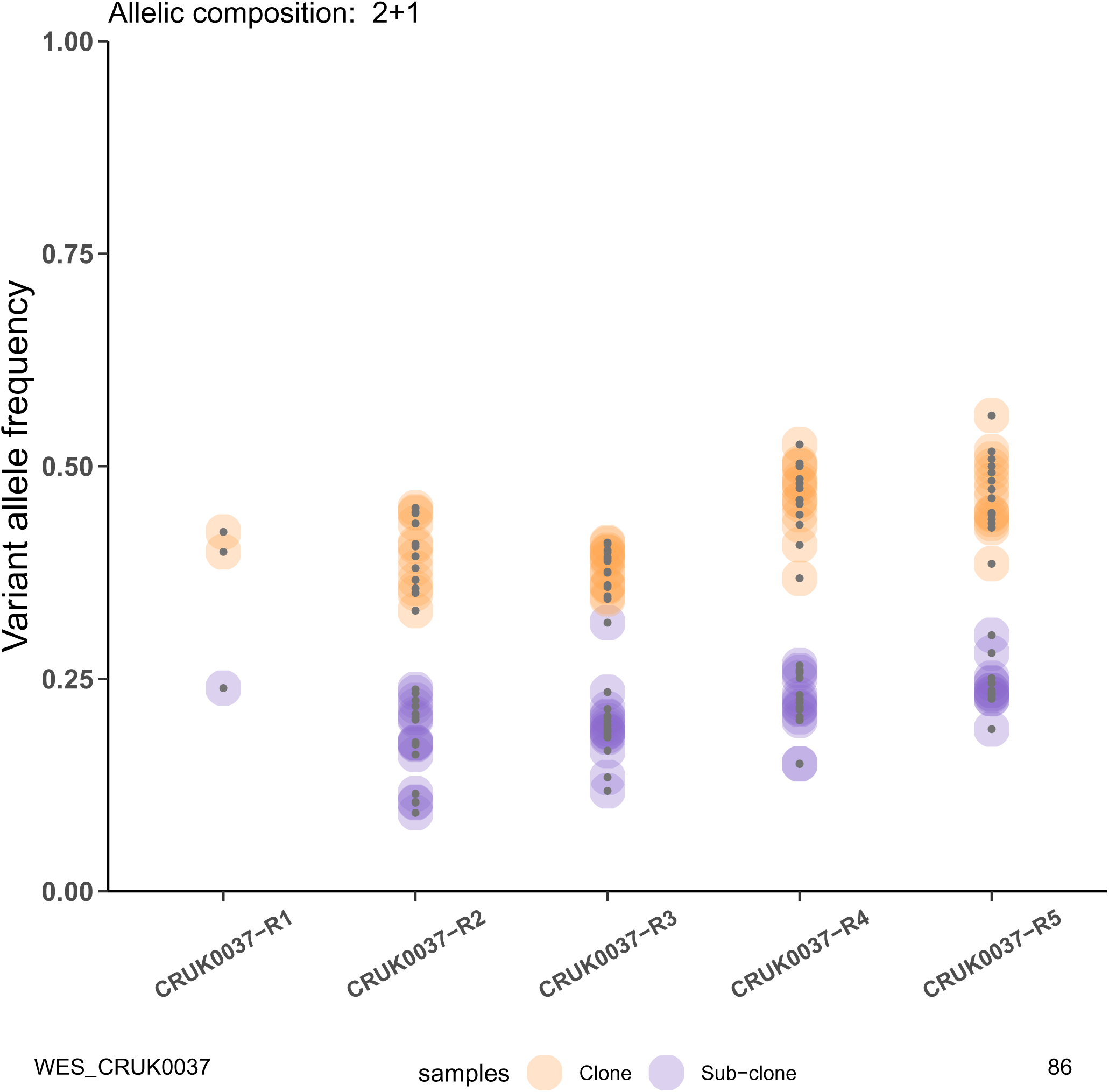
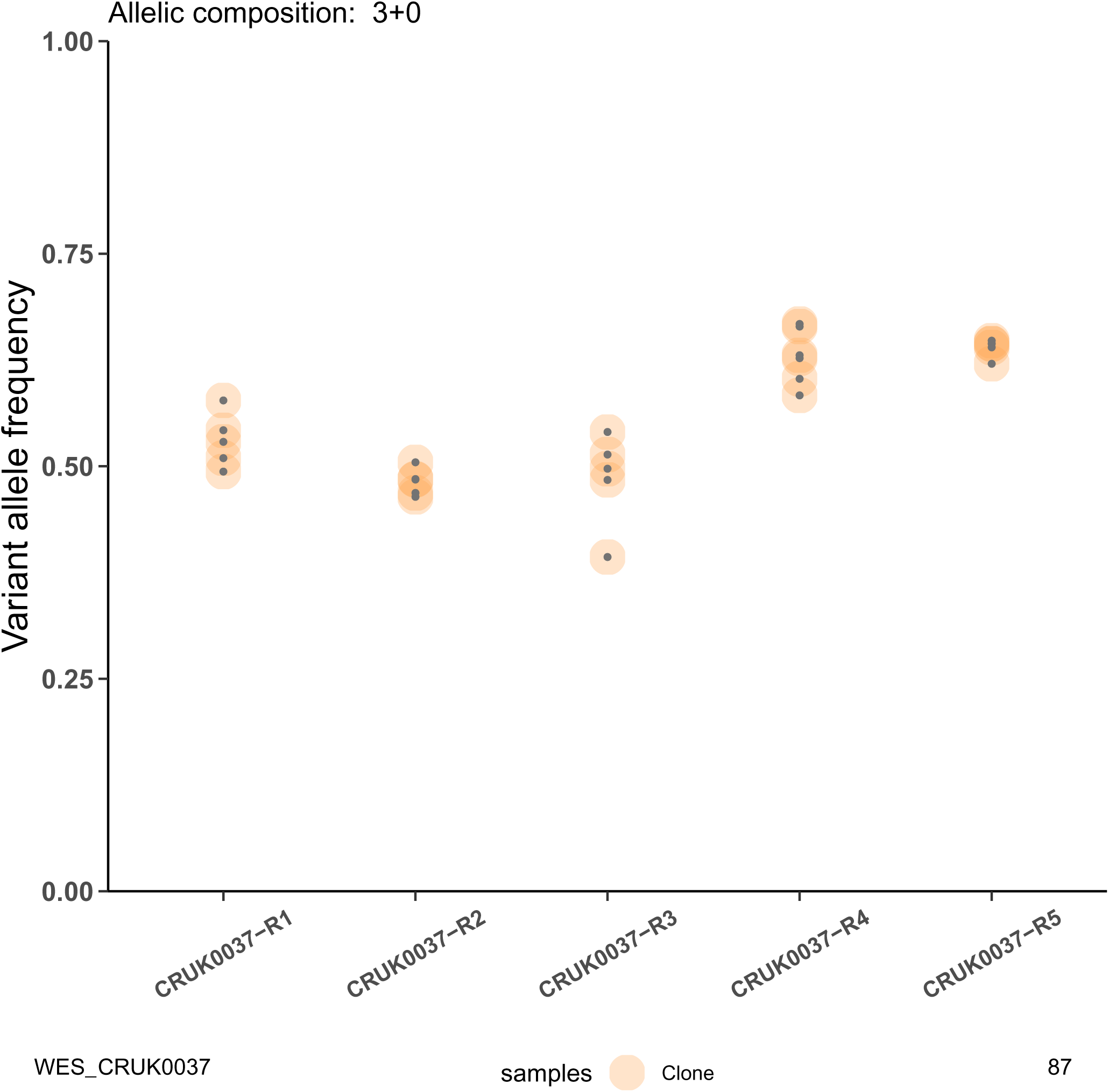
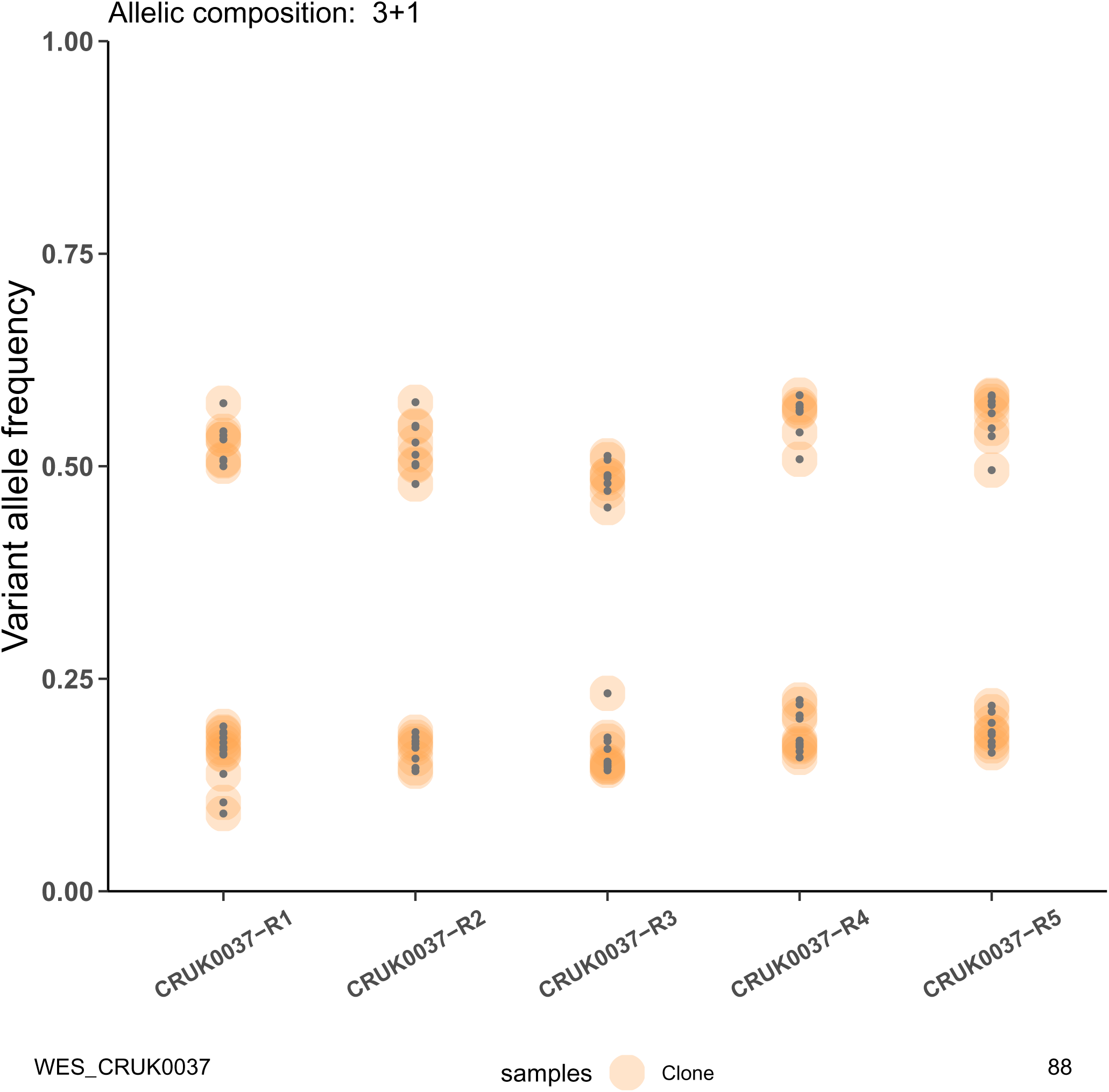
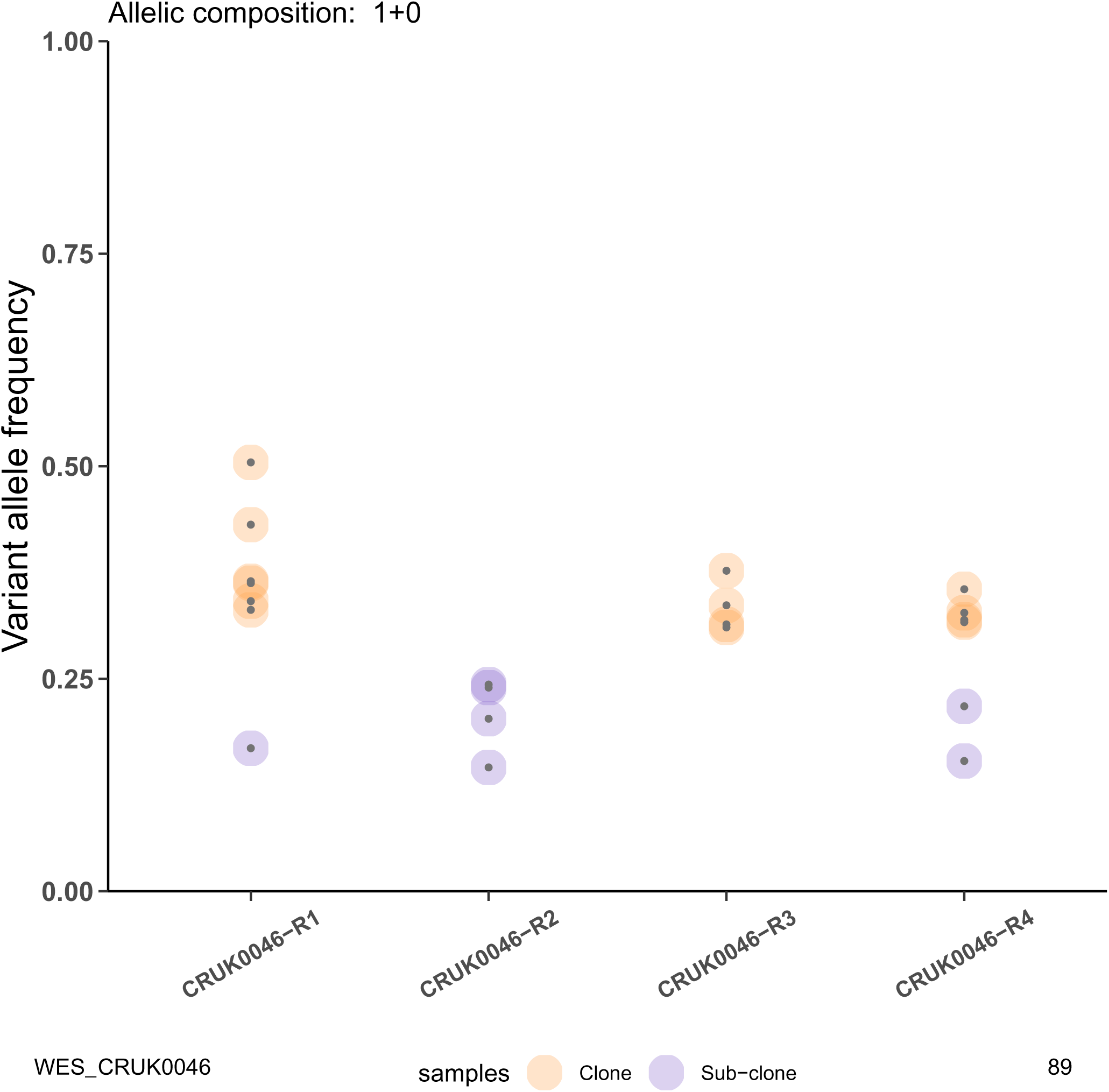
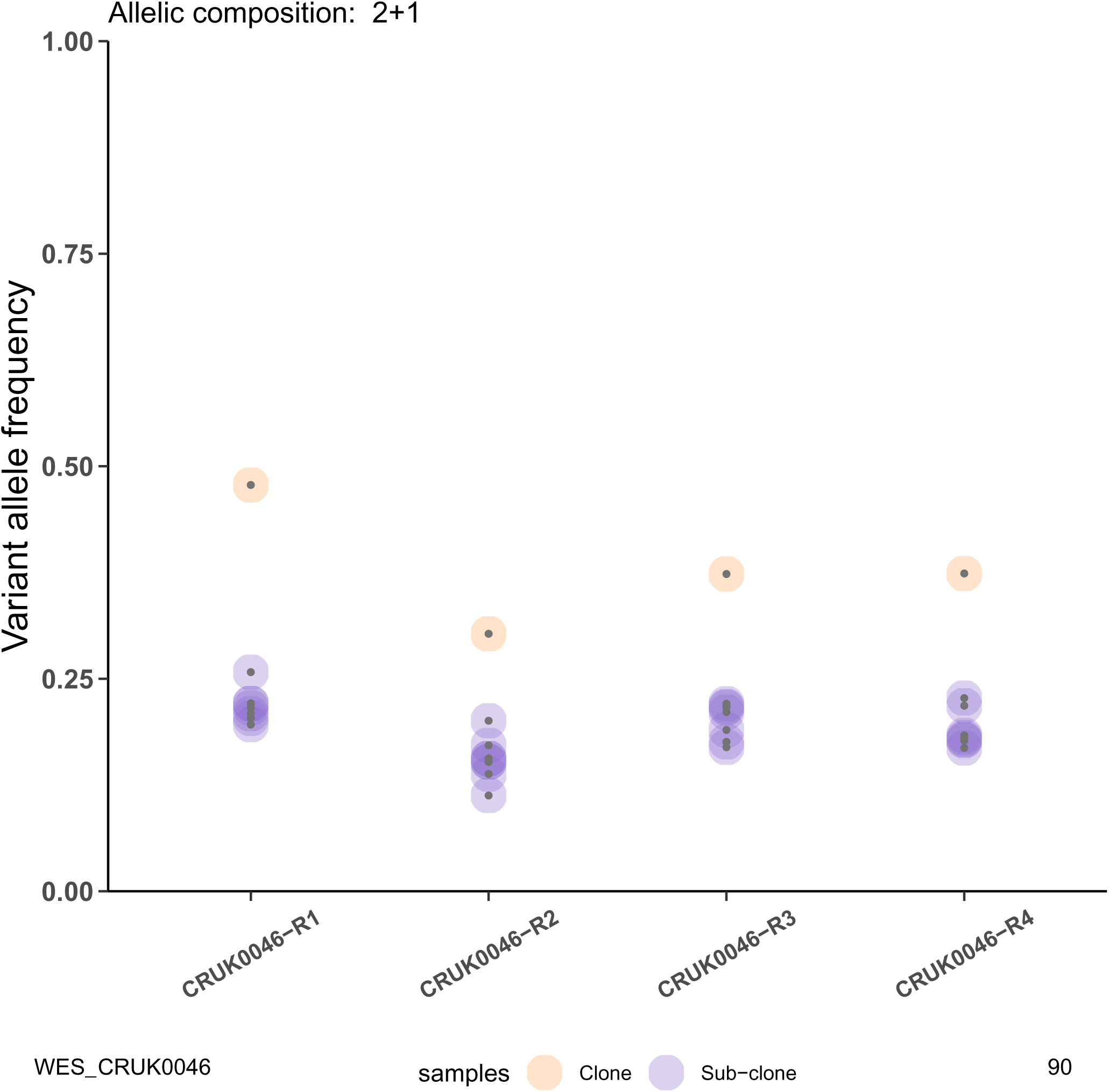
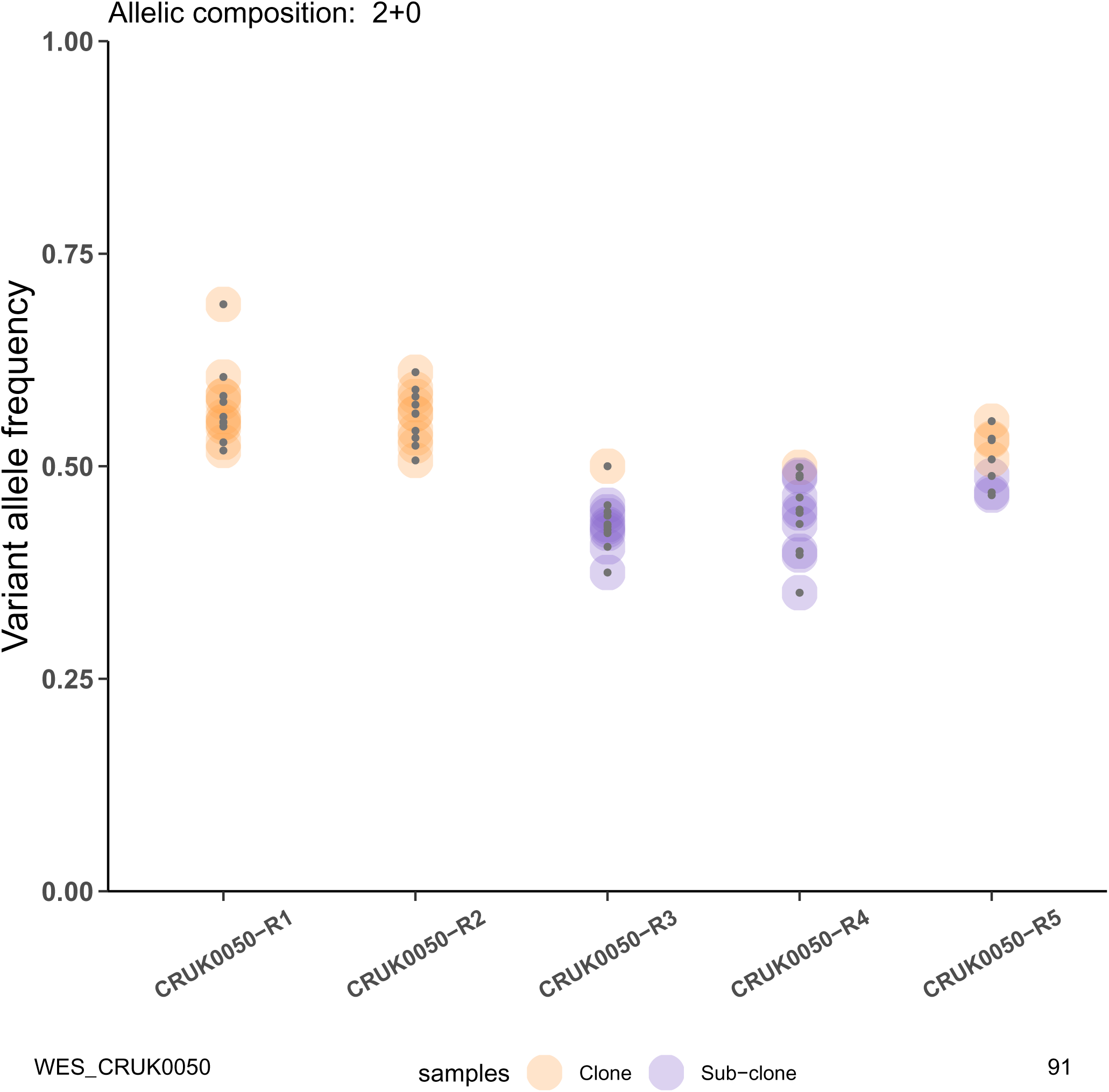
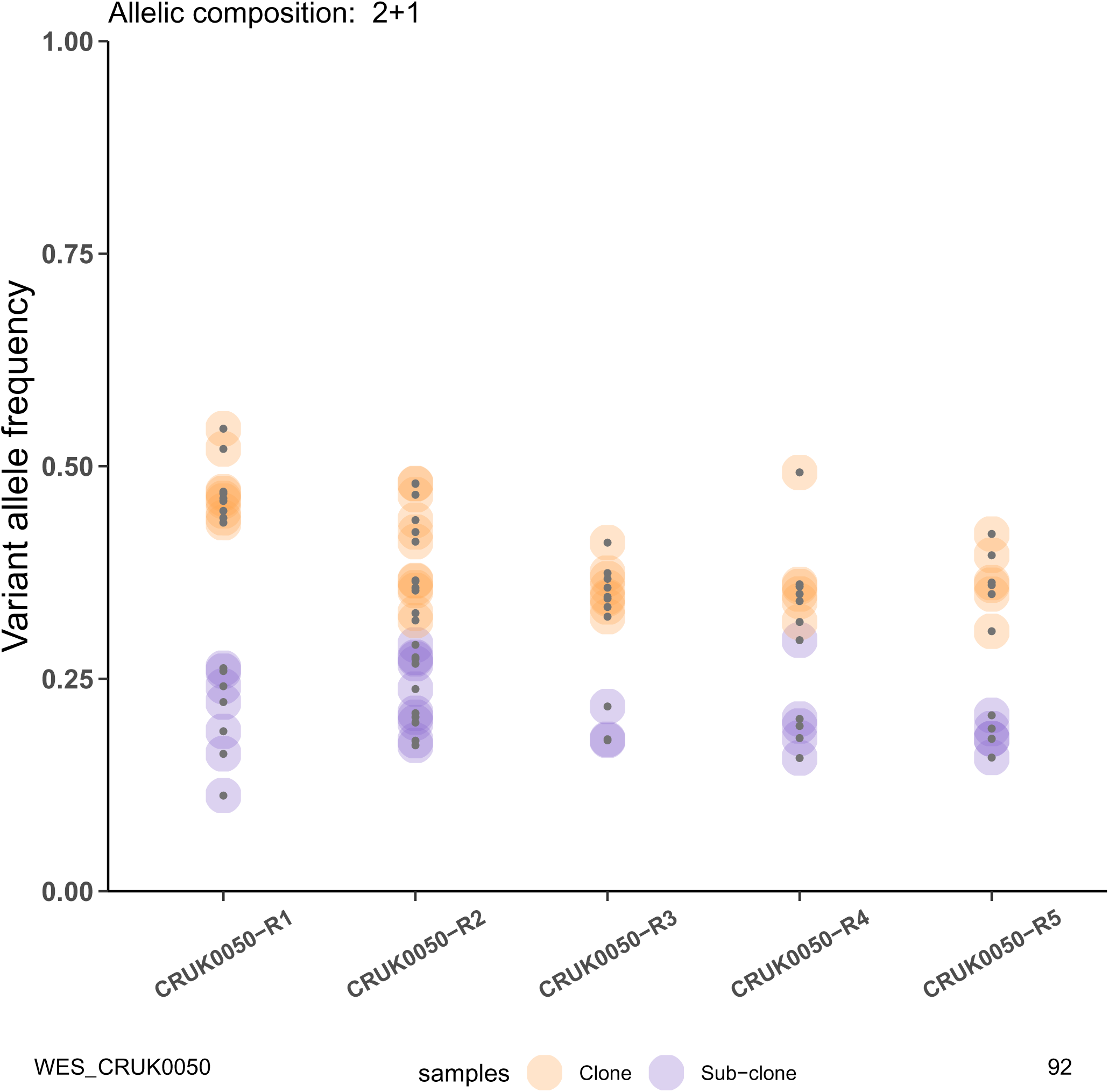
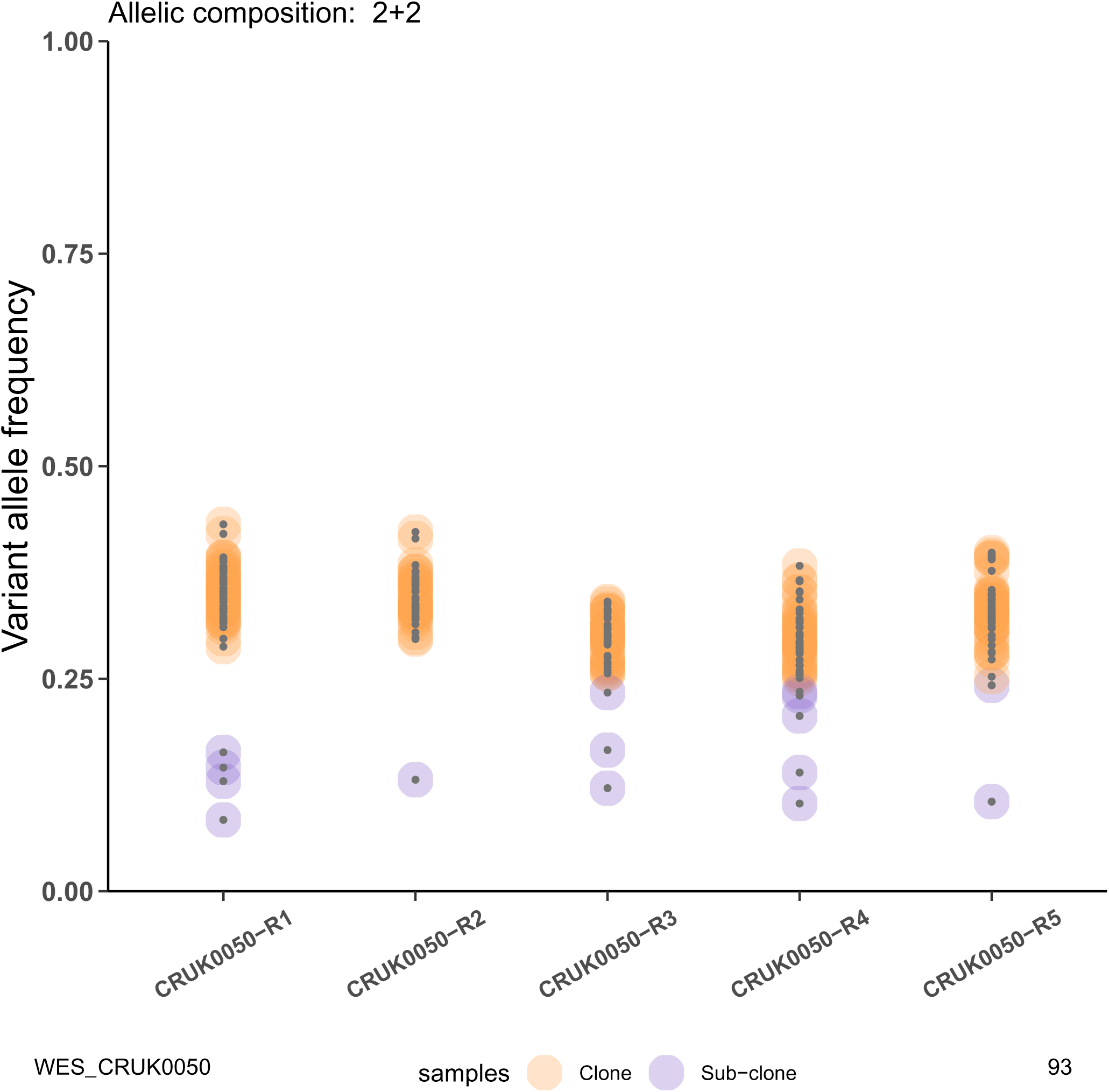
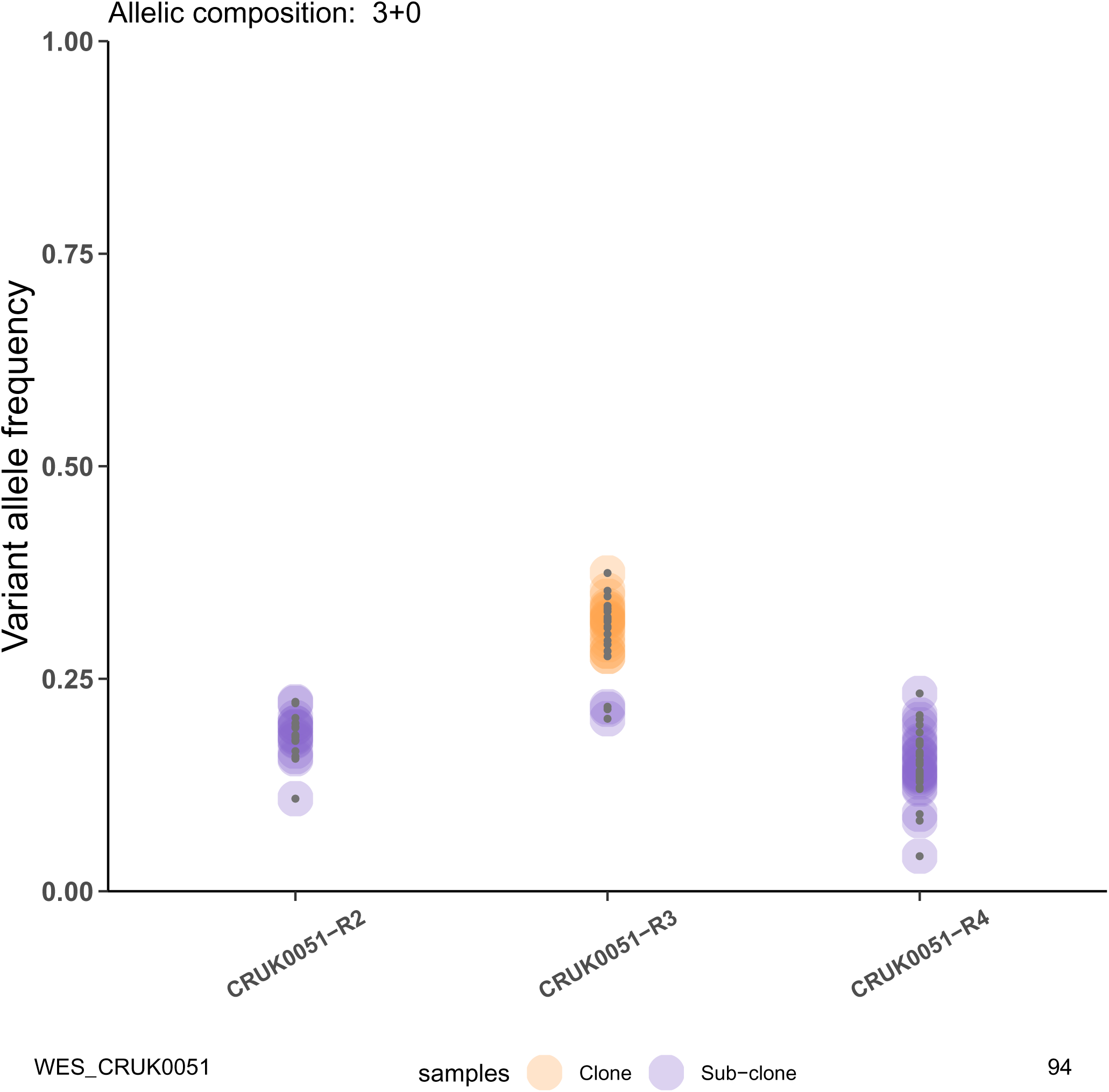
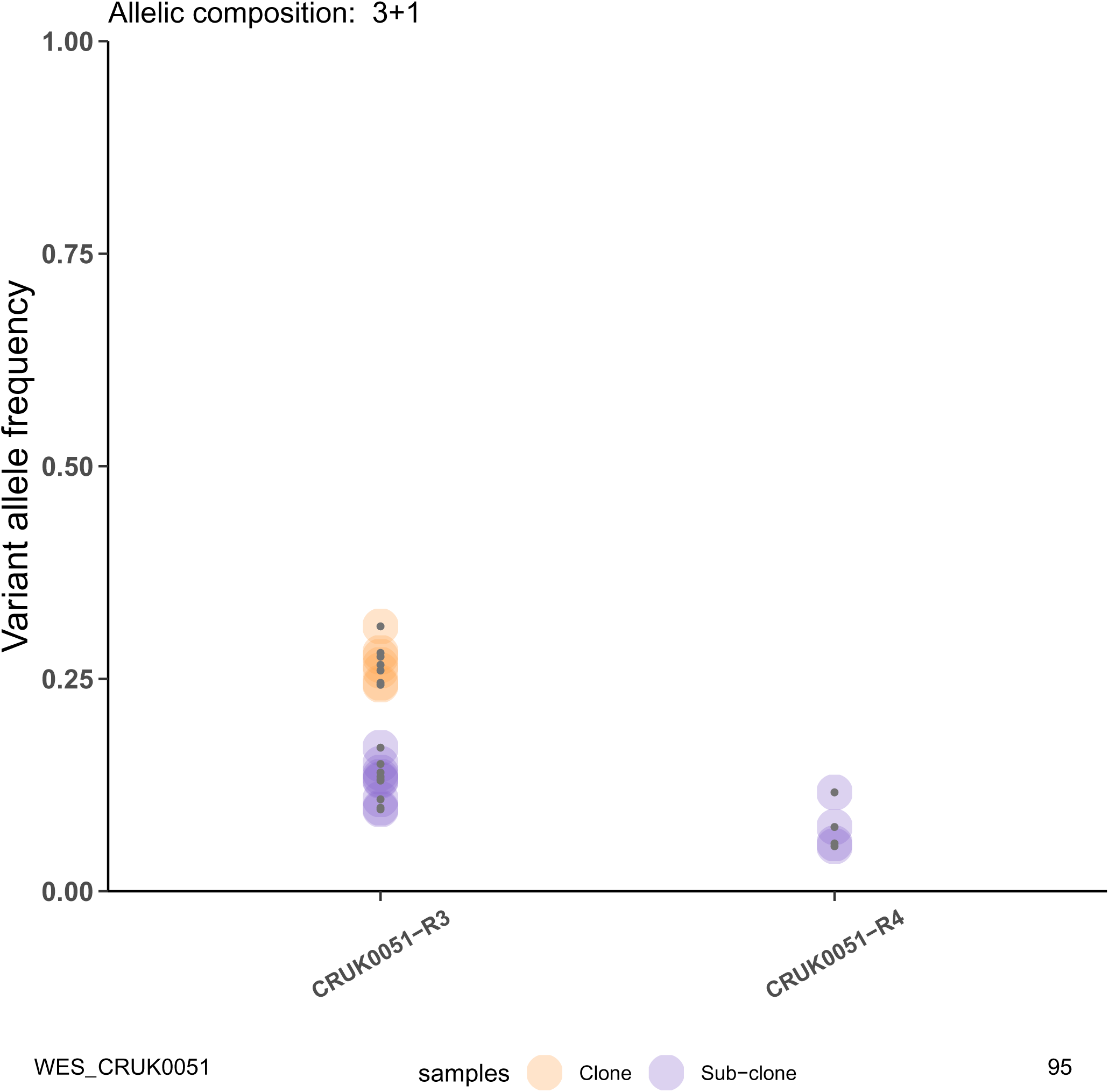
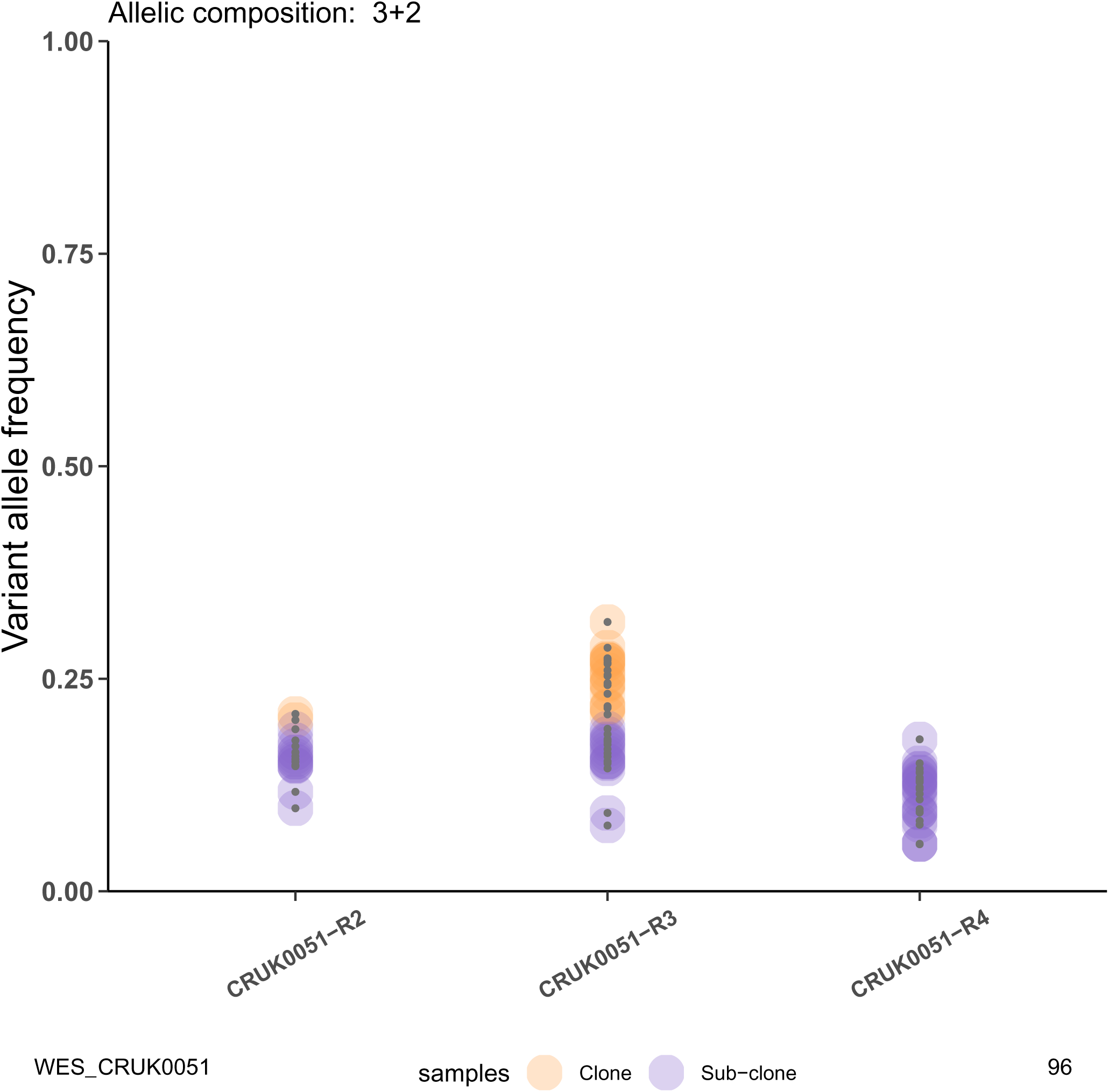
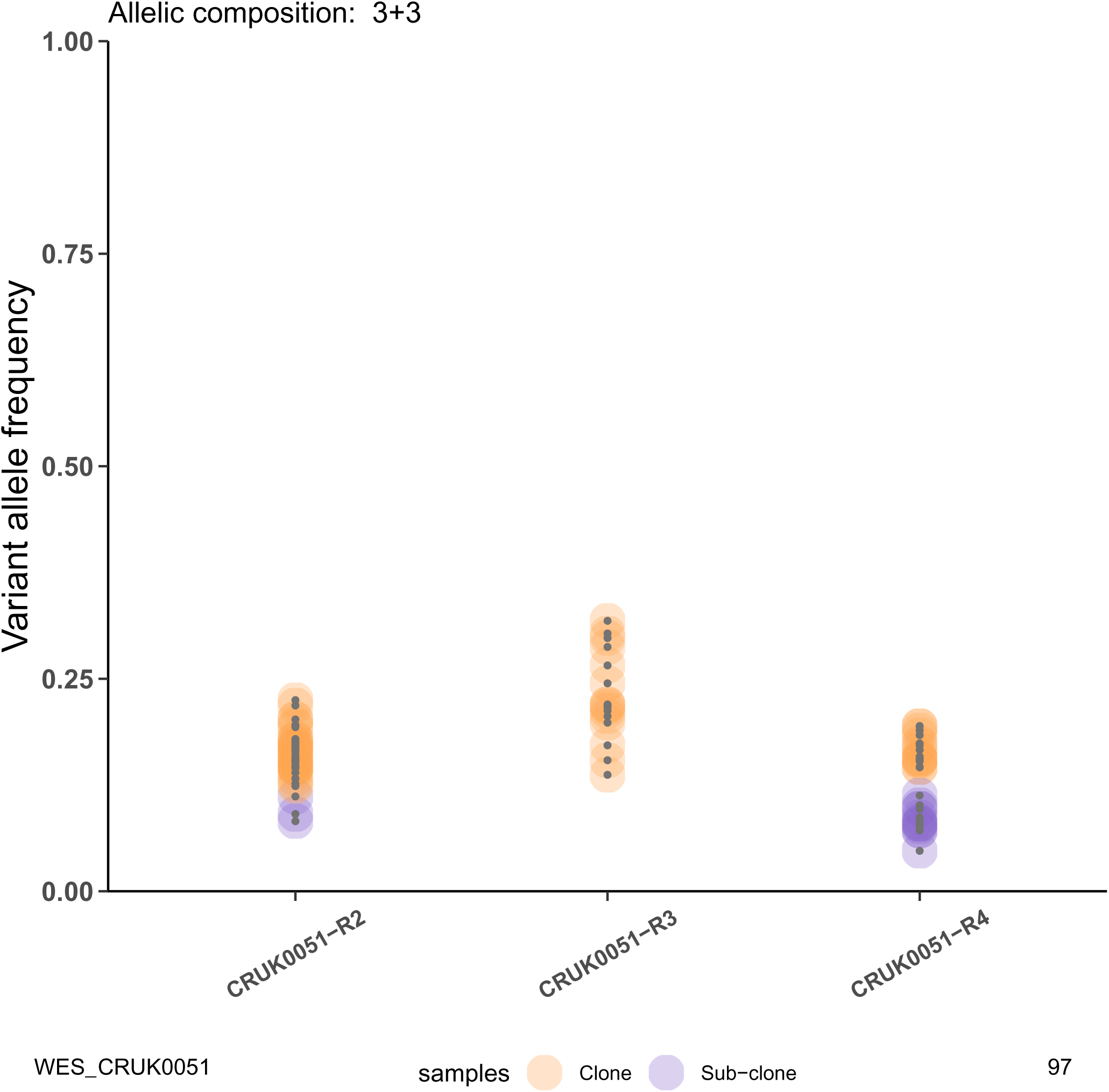
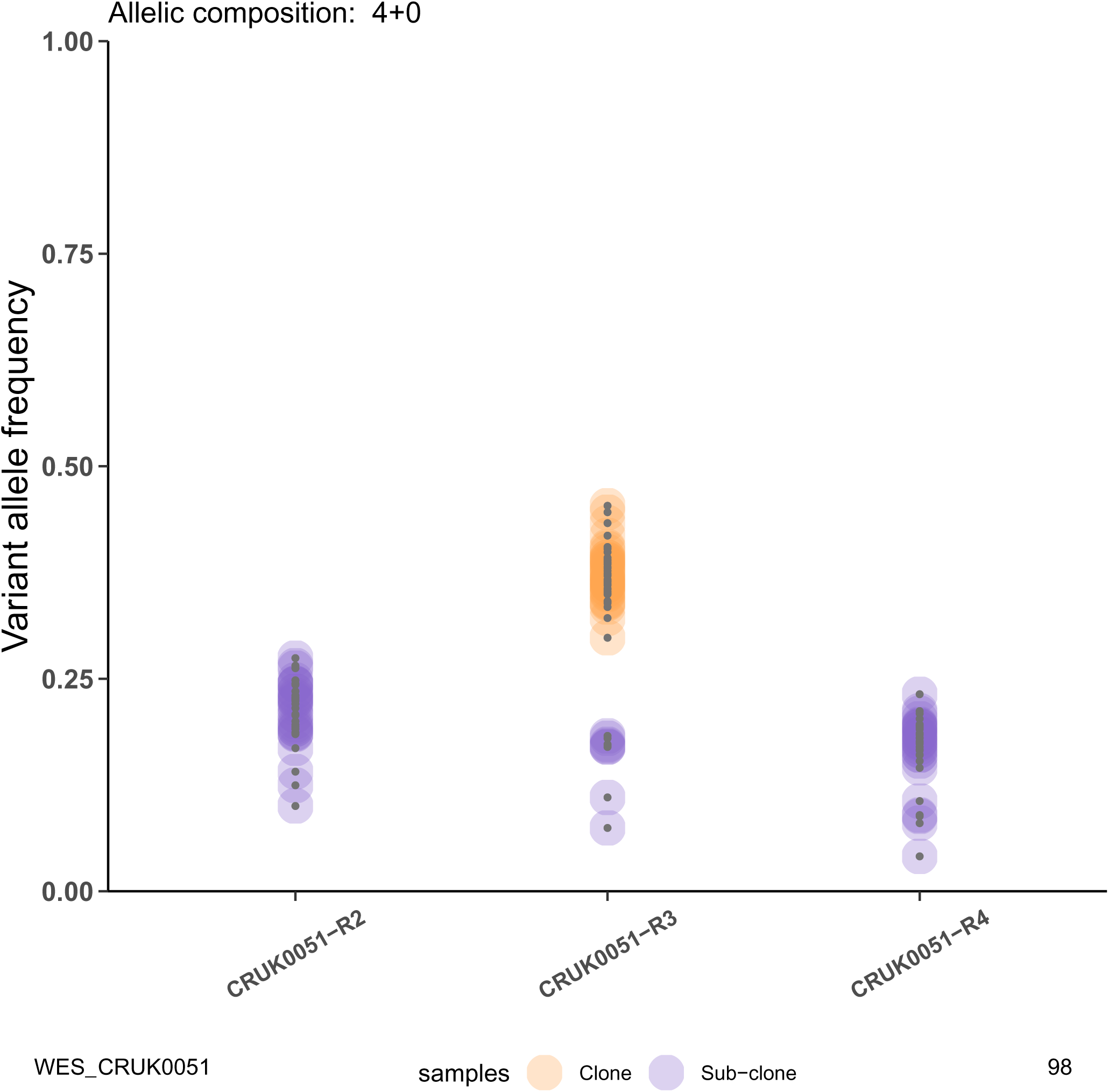
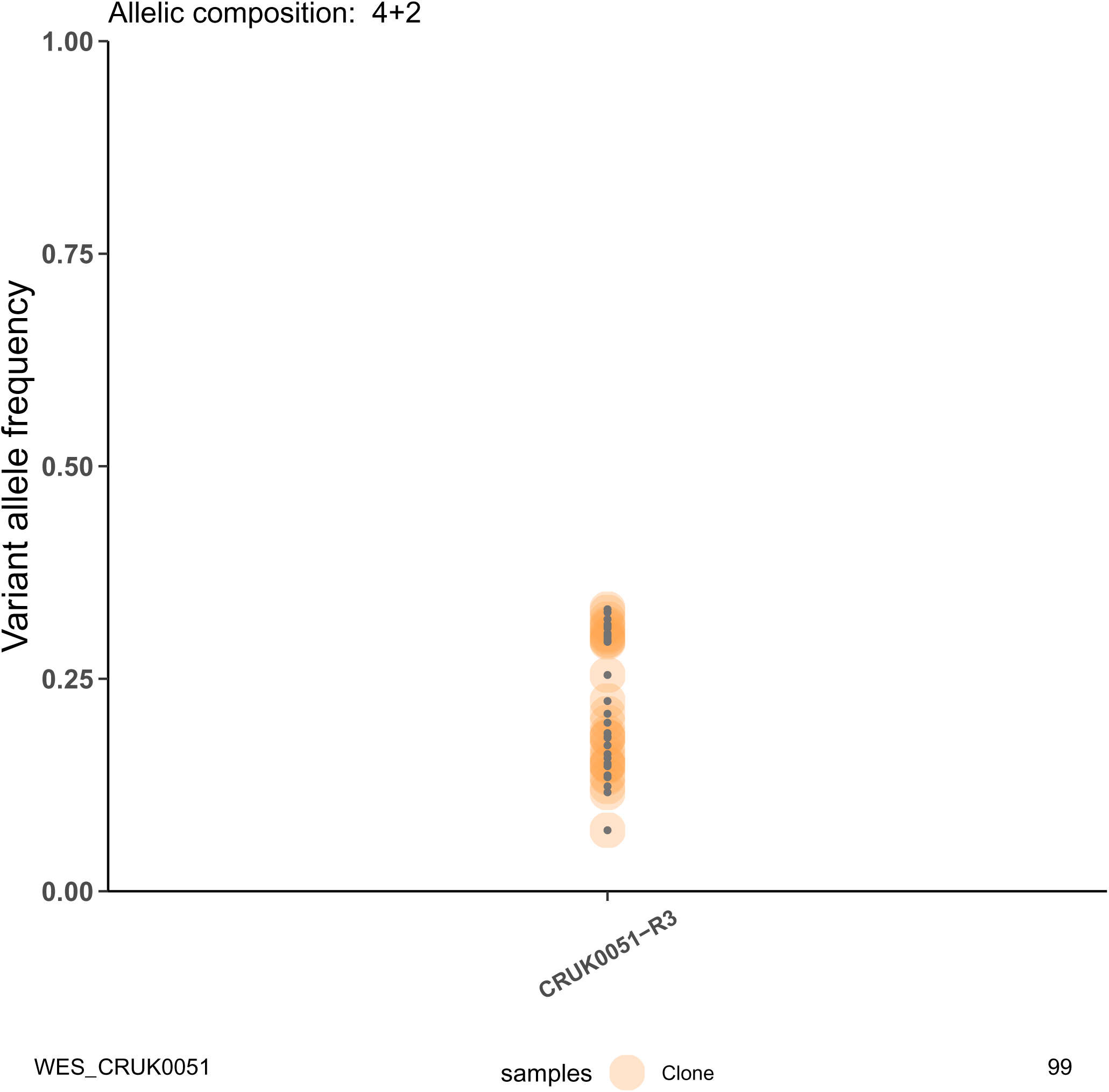
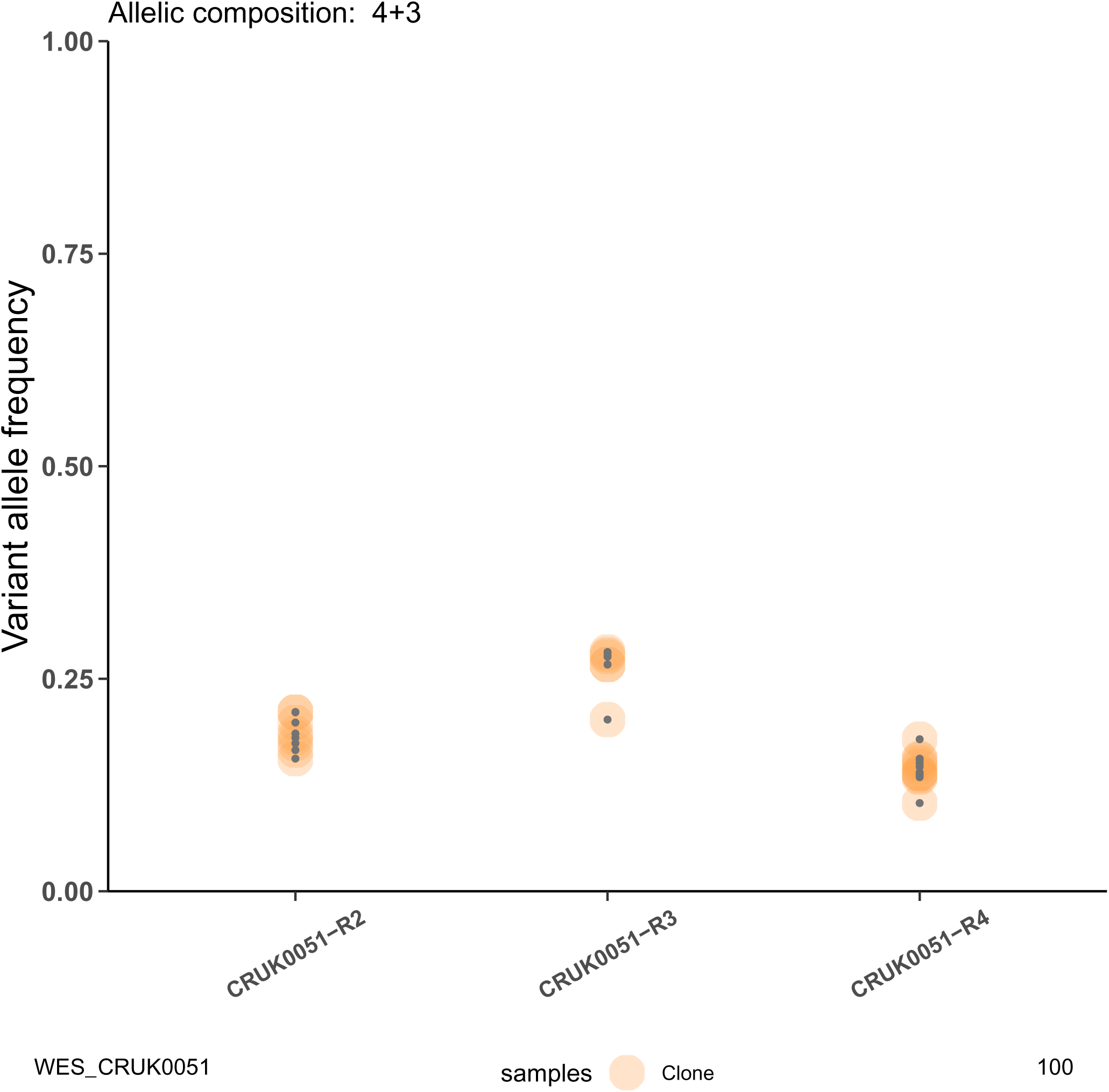
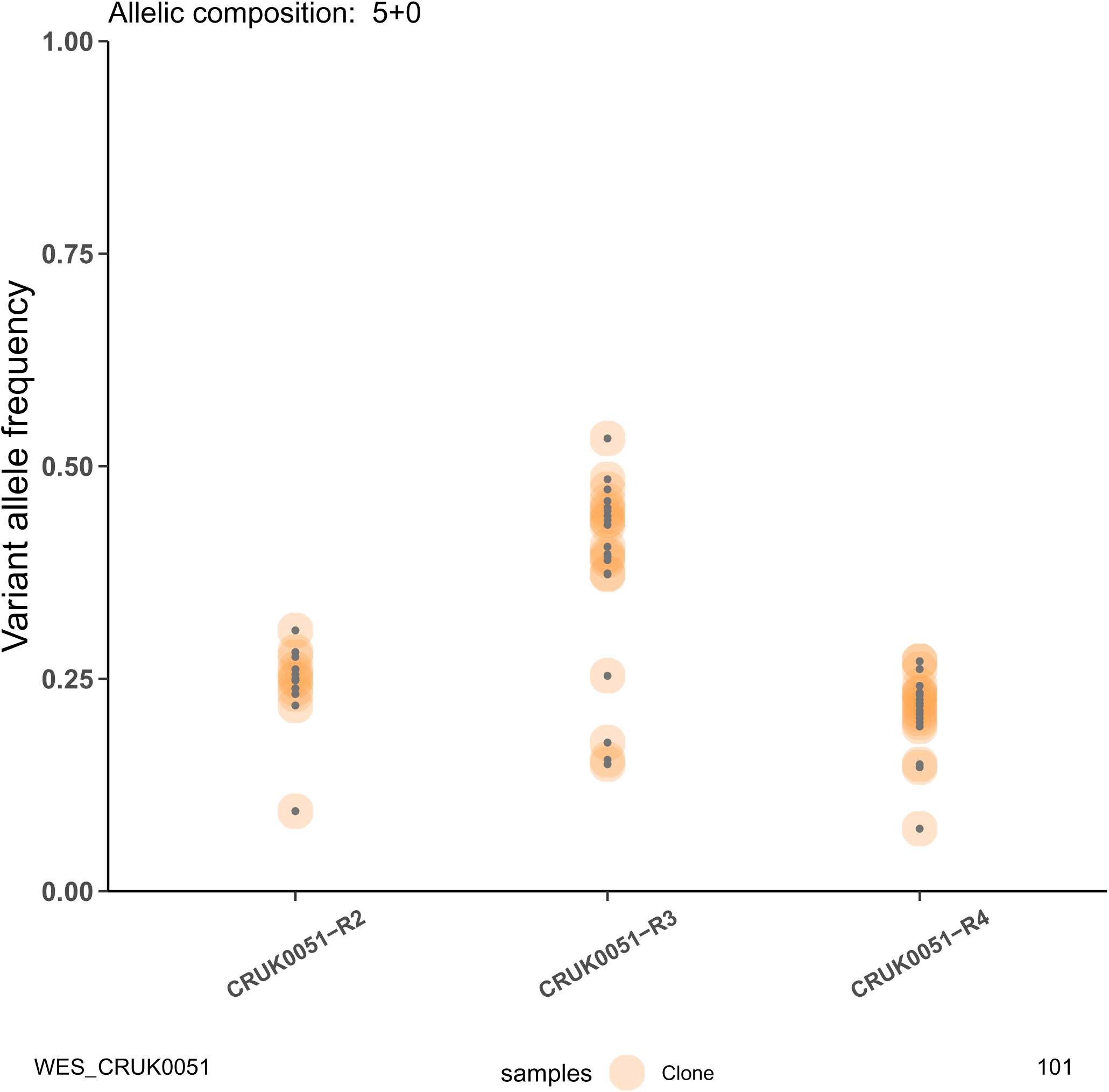
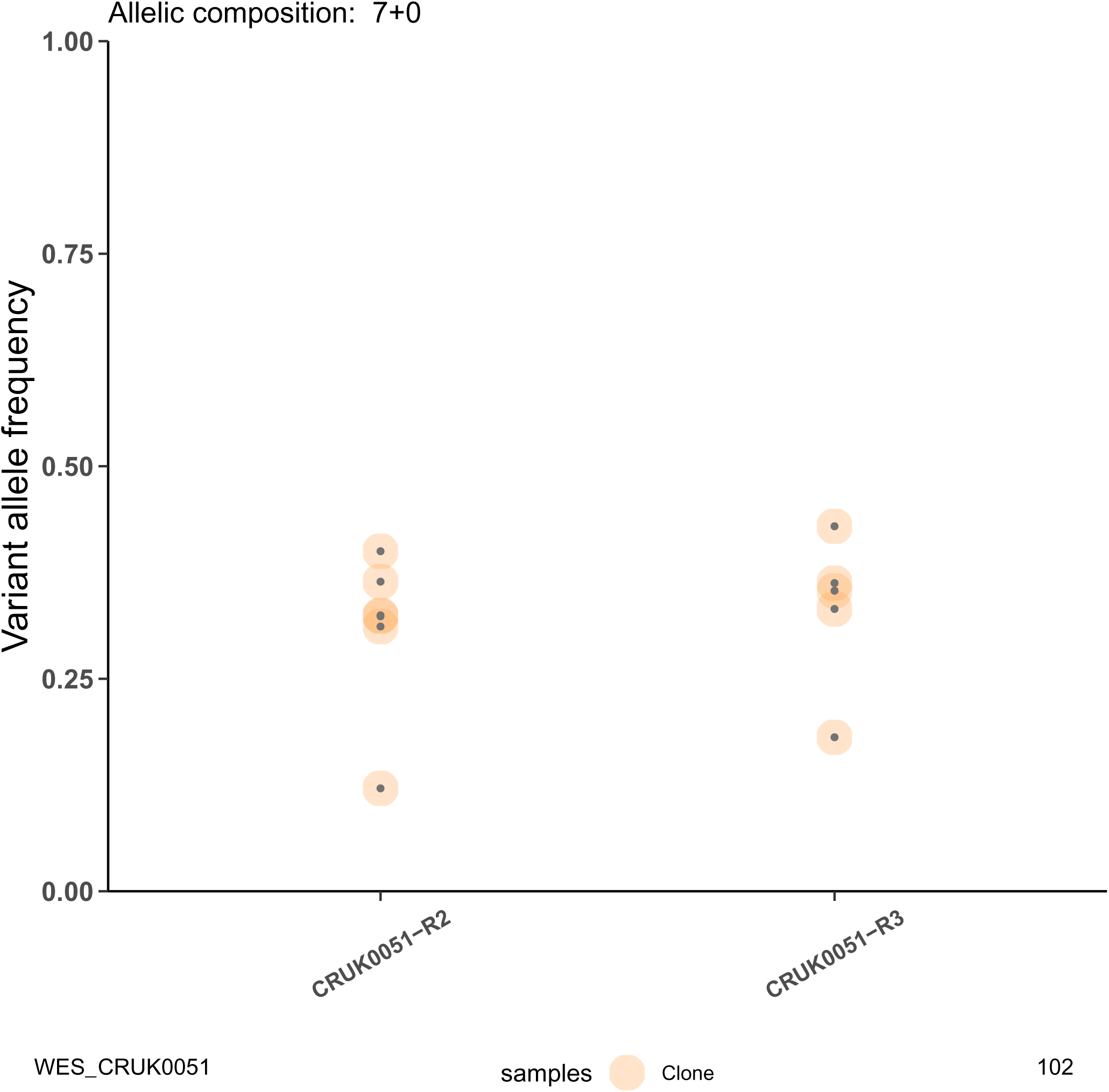
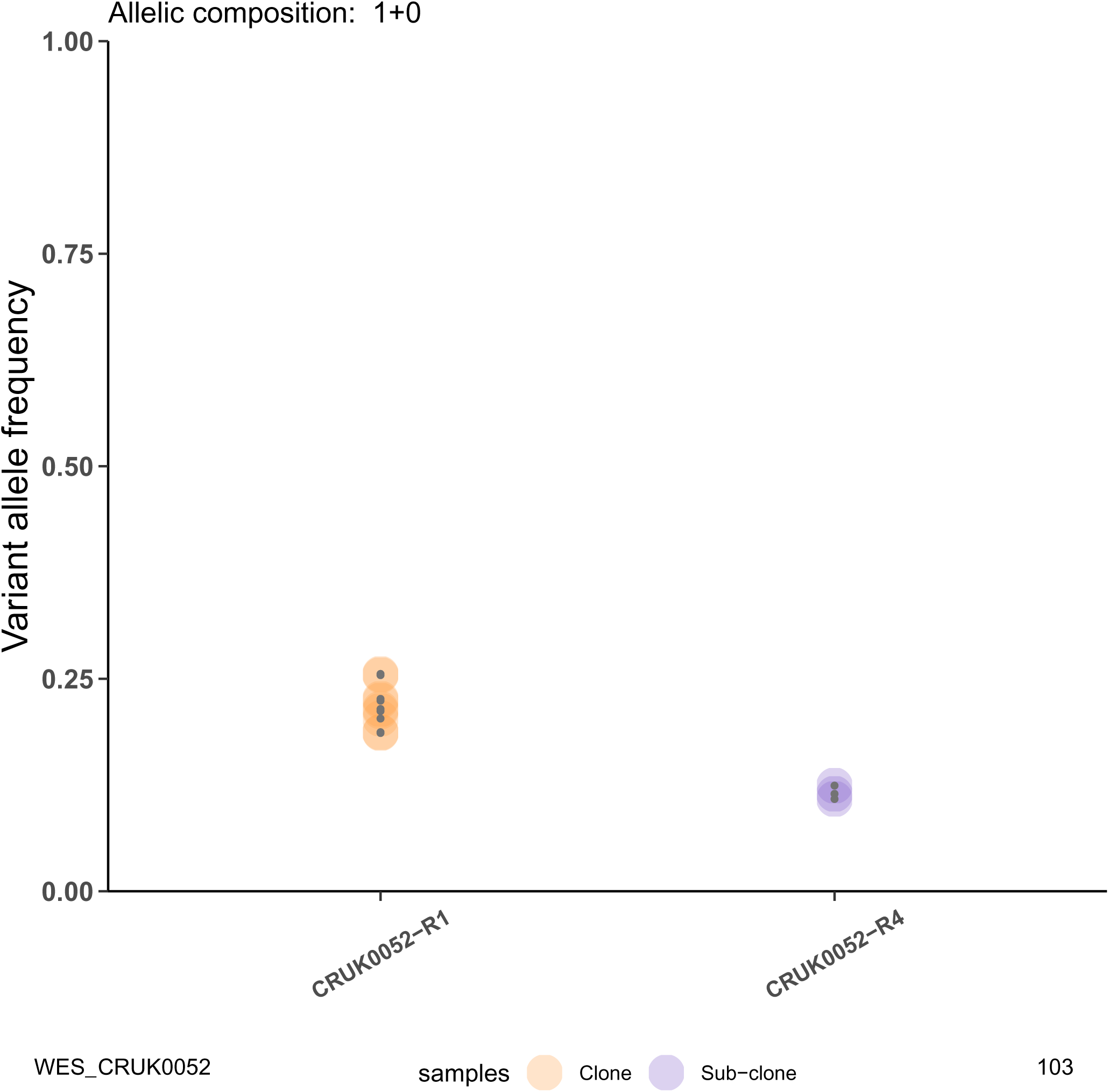
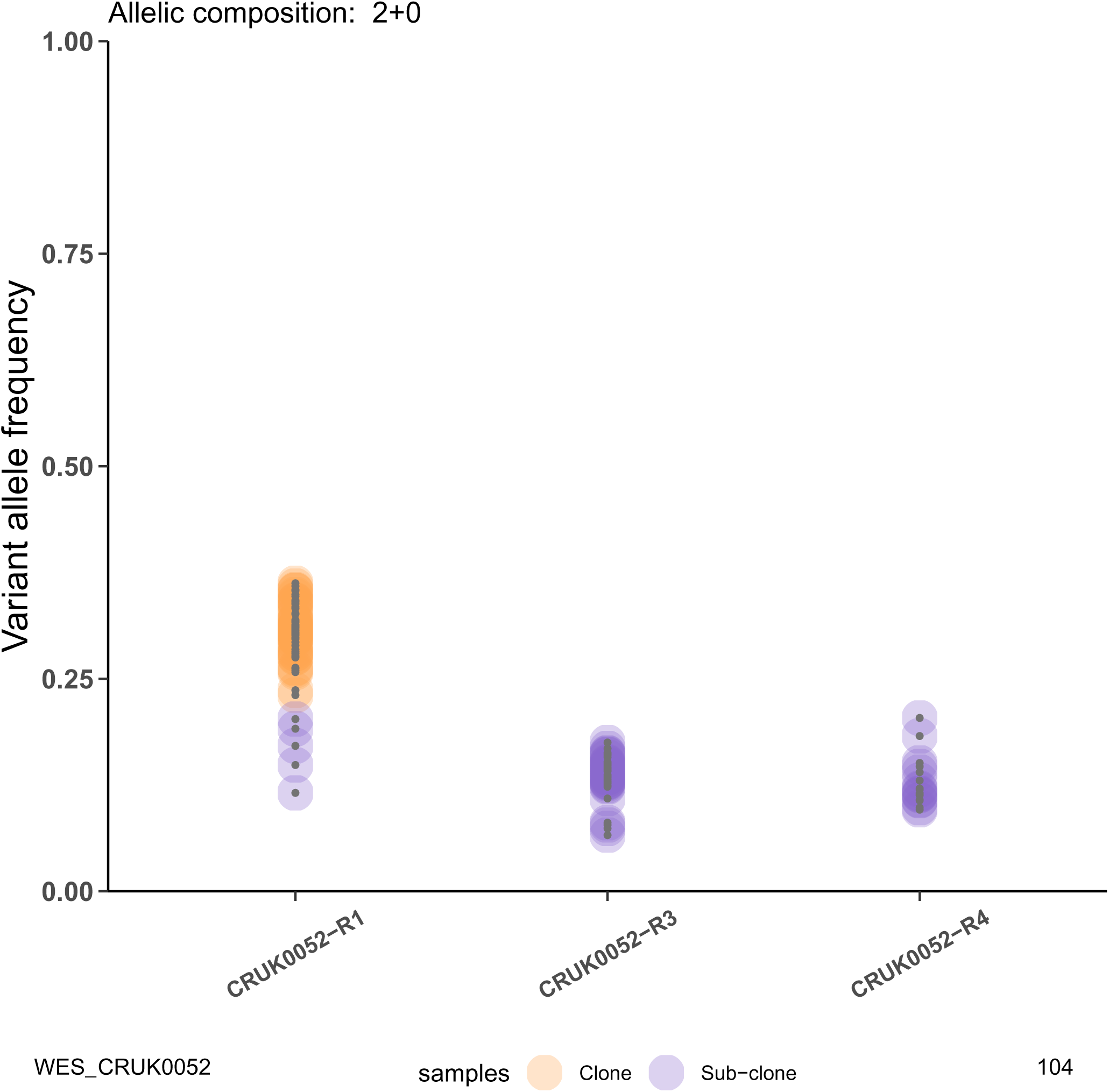
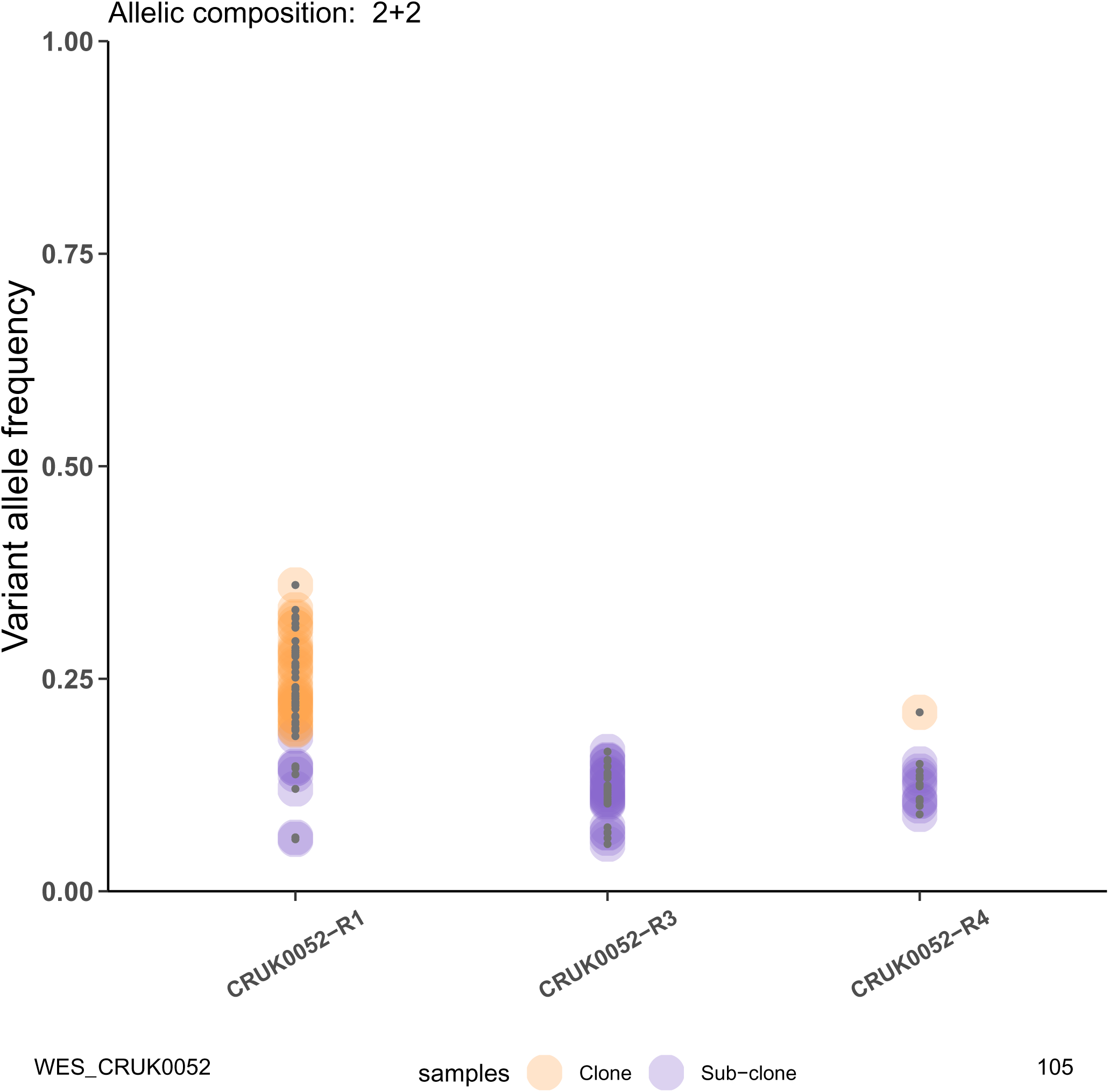
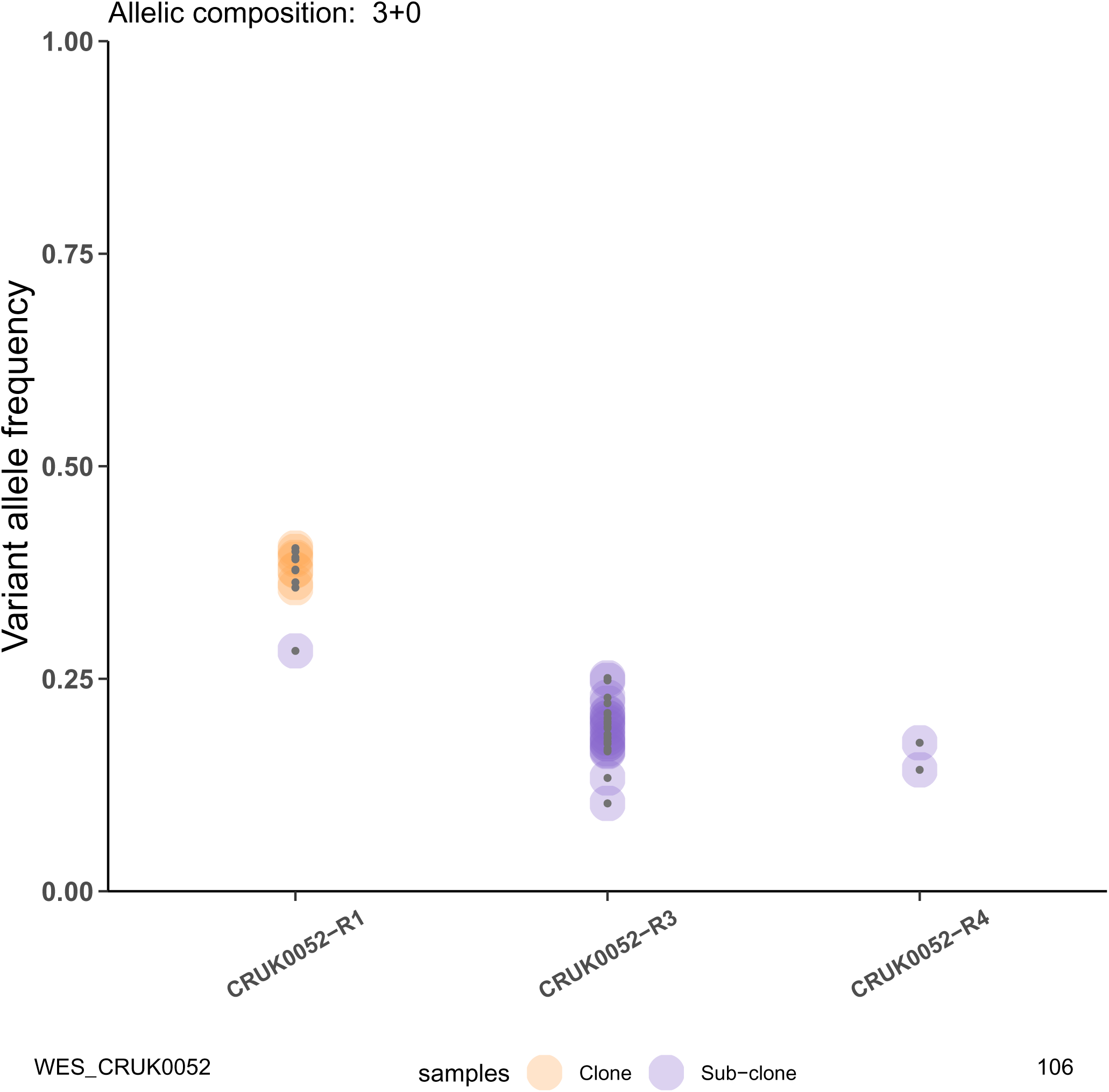
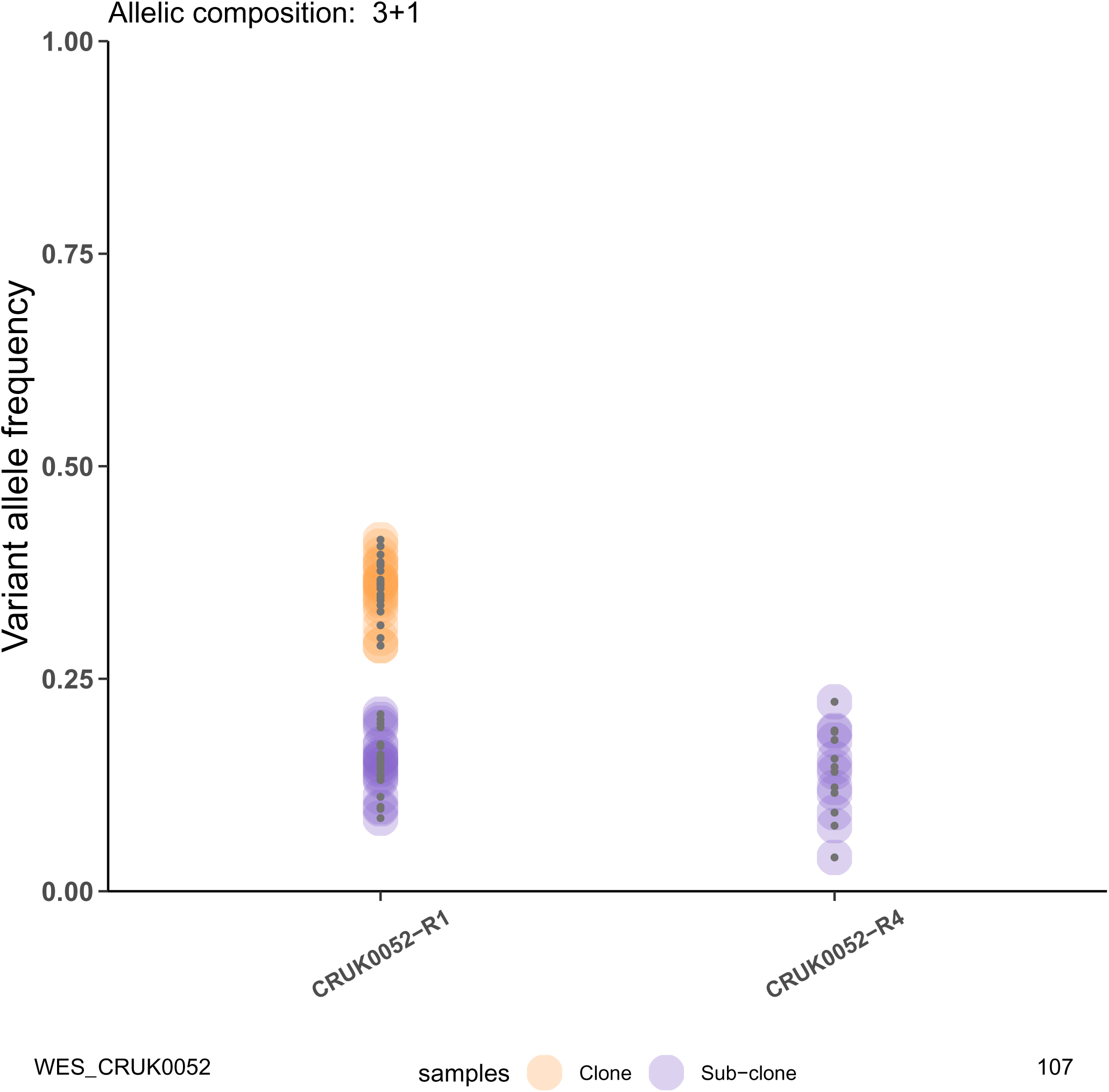
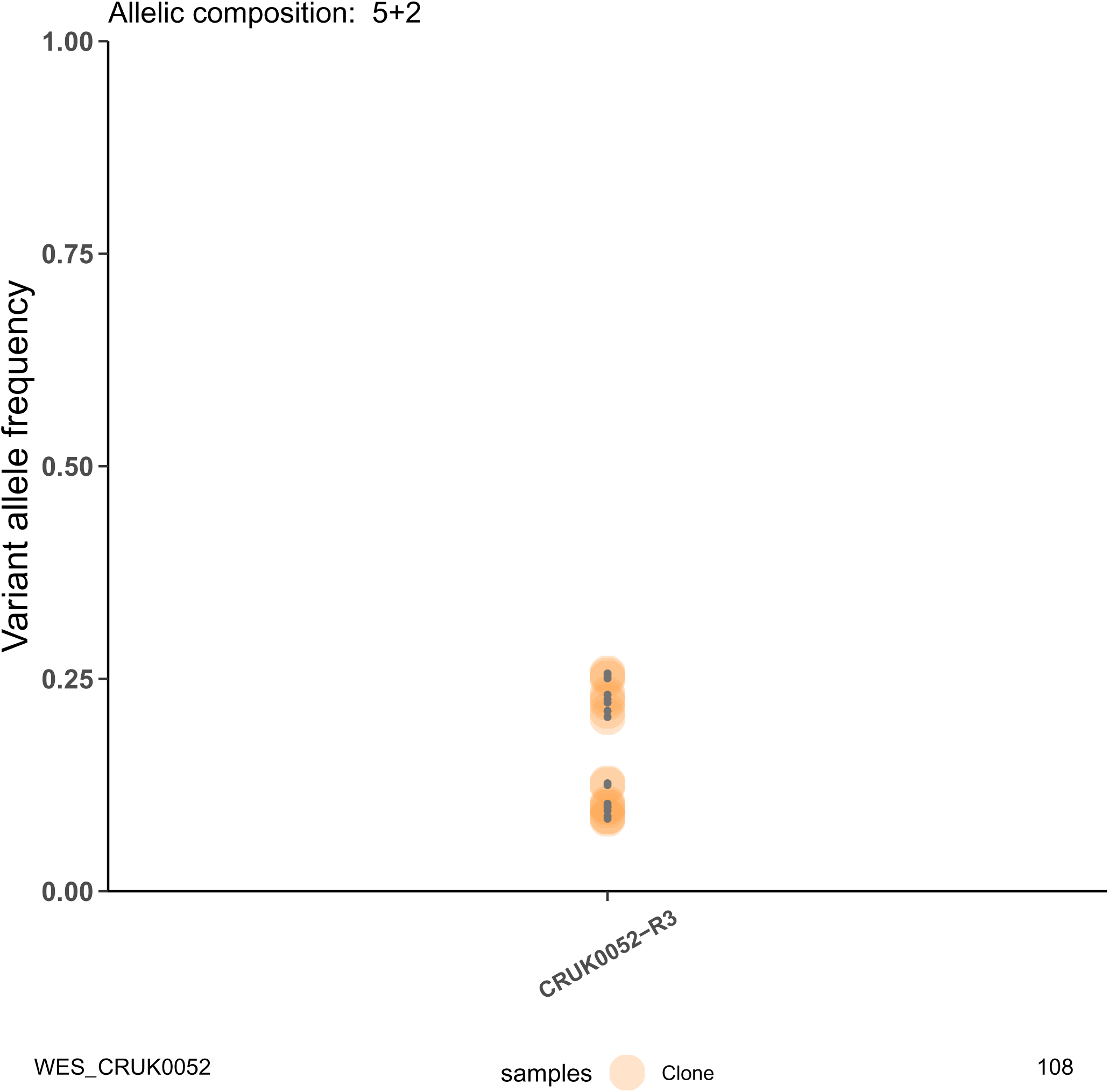
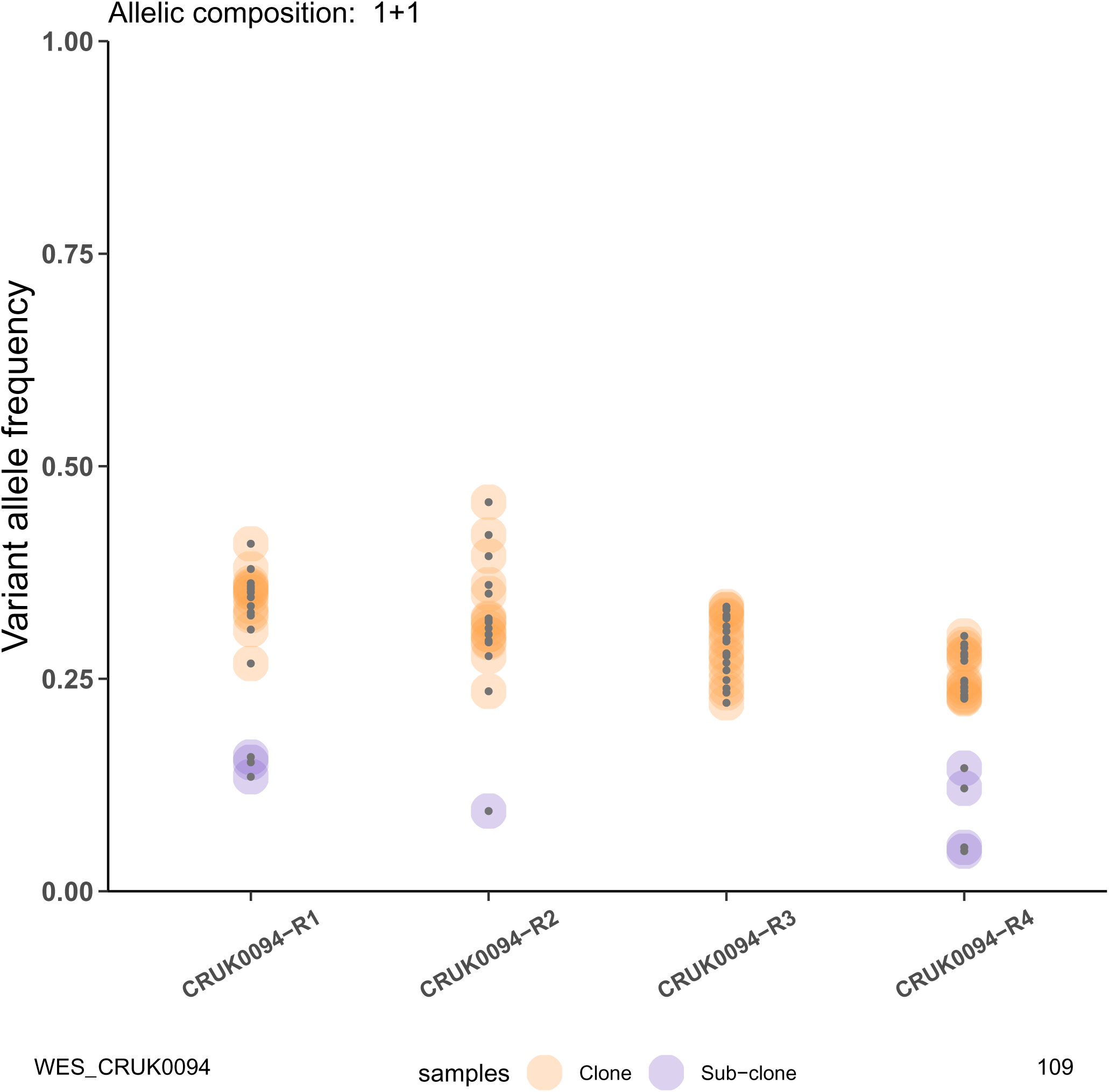
Clonal deconvolution performed by CRUST for all 20 tumors. All variants across all samples from each tumor were segregated according to their respective allelic composition. Separate analyses were performed for each distinct allelic composition. The variants are annotated as clonal or subclonal after analysis according to which populations they were inferred to belong to.

## Supplementary table legends

**Supplementary Table 1**

*Scaling* demonstrates scaled and unscaled clonal deconvolution of the three samples of pediatric neuroblastoma tumor NB12. The differences in clonality predictions are highlighted with conflicting colors.

*NB22_copynumber* lists the estimated segmental copy numbers from whole exome calls on the four samples from pediatric neuroblastoma sample NB22.

*phs00159_seq* includes summary statistics of sequencing on the tumor sample AML31 with whole genome sequencing, exome capture, custom capture and custom ion torrent as provided in the study dbgap: phs000159

*phs00159_iontorrent* are the summary statistics of sequencing on AML31 with custom ion torrent provided in phs000159 additionally quality controlled to include variants only with at least 15x coverage at the mutated allele.

*CN_comparision* is a comparison of predicted allelic composition estimated with CRUST with published (Karlsson et al., 2018) SNP array data with the pediatric neuroblastoma tumor NB22.

**Supplementary Table 2**

Summary statistics of clonal predictions provided by CRUST on the TRACERx data is provided along with comparisons with the published results.

*Deconvolution results* enlists all distinct variants that remain after quality control steps performed on the samples from 20 NSCLC tumors (a total of 9443). The predicted allelic composition and the clonality status are appended along with the WES summaries of these variants.

*Clonality predictions* provides an array map of all annotated genes in the CRUST analysis. Each cell correspond to data on all variants (if present) of a gene (primary row name) in a specific sample (secondary column name). The clonality predictions are in secondary row names (Clone/ Sub-clone). The numbers in each cell shows the frequency of a clonality prediction that relates to the gene and the sample. (example: 1 in the cell corresponding to CRUK0001-R1, *AASS* gene, Sub-clone means: there is one variant in the sample CRUK0001-R1 belonging to the AASS gene that was predicted to be subclonal.) This table helps visualizing clonal crossovers in genes (example: *AAK1* in CRUK0017).

*Clonal crossover* gives a frequency-based summary of the samples, variants and corresponding genes analyzed by CRUST. This table provides insight on the quality of the samples, the heterogeneity in allelic compositions, the prediction tendencies of CRUST as well as a comparison of the analysis with the published data. *Primary samples selected* is the number of samples extracted per tumor (row name); *distinct copy number states* means the number of unique copy number states observed per tumor; The *crossover variants* indicate clonal crossover events and *dropped variants* are those that were excluded from analysis in the published study. The variants analyzed are summarized at gene level with number of distinct genes retained per sample (H). The number of genes (I) and variants (J-K) corresponding to crossover events are denoted to demonstrate the commonality of it across samples. To assess prediction sensitivity of CRUST percentages of clonality prediction, crossover variants, genes were reported along with that of dropped variants from the published data (L-P).

*Crossover genes* enlist all the genes that harbored variants undergoing clonal crossovers in the corresponding tumors (column name).

*Jamal-Hanjani et al., 2017* includes the clonality and gene status predictions originally provided with the published data on all 100 TRACERx tumors. This table also includes the following added columns: *Driver (original)*, if the gene was predicted to be a driver in the published study; *subclonal driver (original)* if it was predicted to be a subclonal driver; *Retained in CRUST*, if the variants from these genes were retained to be analyzed with CRUST after quality control; *Crossover predicted by CRUST*, if the gene had observed clonal crossover. Columns G through DB enlists the sample-aggregated published clonality predictions for all 100 tumors. This table helps identify if a crossover gene was predicted to be driver or not (if yes, was it clonal or subclonal driver).

*Clonesize estimation* includes all analyzed variants and indicate the base ploidy level of the corresponding sample (diploid, tetraploid etc.) and the estimated unscaled clonesize.

*Clonal separation* is a table summarizing scaled clone sizes estimated per chromosome for each sample. The distinct size of clonal and subclonal clusters are computed by aggregating several variants that correspond to similar clone sizes (range of 15%). Assuming that, 1) the unknown purity of the sample would affect the inferred sizes and 2) the largest estimated clone size can be scaled up to be 100%, all sample specific estimates were rescaled.

*Clonal nesting* includes a pivot table that can enlist sample specific clonal and subclonal clusters discerned from the previous table. On right there are three tumors for which subclonal nesting is inferred. As a rule of thumb cluster sizes within 15% of each other across samples were inferred to originate from same population. These three examples are also featured in Figure 5.

